# Rational design of selective bispecific EPO-R/CD131 agonists

**DOI:** 10.1101/2025.04.10.648227

**Authors:** Kailyn E. Doiron, Jeffrey C. Way, Pamela A. Silver

## Abstract

Erythropoietin (EPO) suppresses apoptosis and promotes survival by signaling through EPO-R/EPO-R on hematopoietic progenitors or EPO-R/CD131 on non-hematopoietic cells. However, EPO signaling through EPO-R/CD131 is controversial and there is no solved structure of a complex. Here, we constructed a structural model of EPO-R/CD131 and designed several anti-EPO-R, anti-CD131 bispecific proteins that selectively activate EPO-R/CD131. Treatment with these fusion proteins is sufficient to activate STAT5 phosphorylation downstream of EPO-R/CD131 without engaging EPO-R/EPO-R. We demonstrated that proteins with a tandem scFv or bispecific antibody format activate EPO-R/CD131, in contrast to an equimolar mixture of the individual scFvs. Finally, we explored the effect of modifications to binding domain arrangement and linker length and found results consistent with our structural model of an EPO-R/CD131 complex. These findings highlight the utility of bispecific scaffolds in the development of cytokine receptor agonists and provide a foundation for the study of EPO-R/CD131 biology and future clinical development.

**Graphical Abstract:** 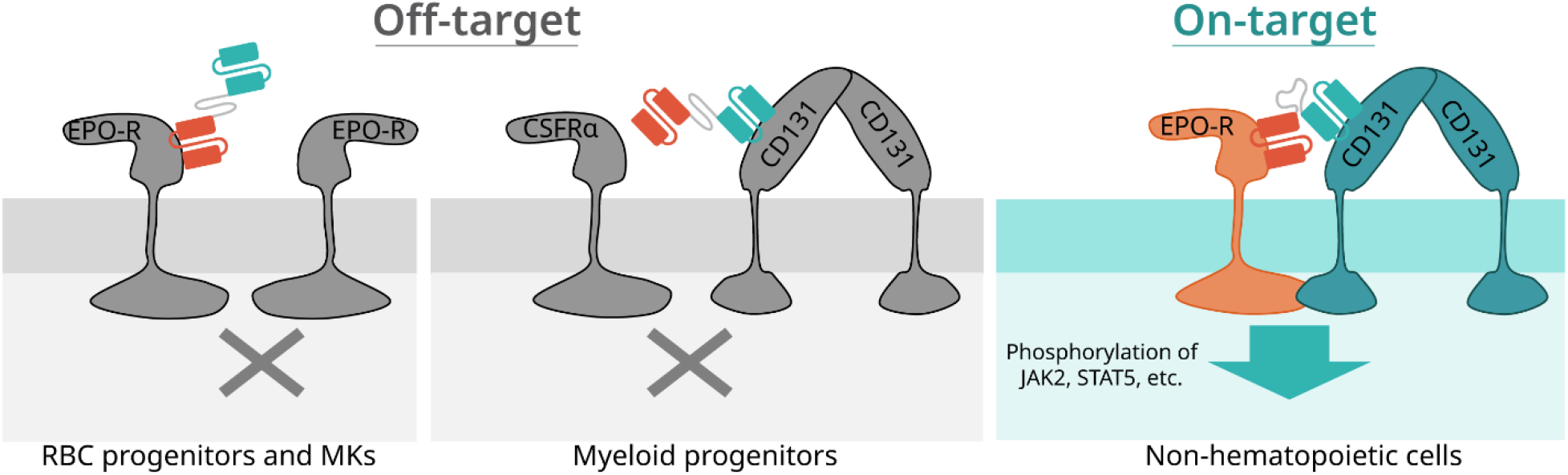

## Introduction

Erythropoietin (EPO) is a cytokine that is essential for red blood cell (RBC) maturation. EPO promotes erythropoiesis by suppressing apoptosis in a subset of erythroid progenitors, enabling their survival and subsequent differentiation (Watowich, 2011). EPO signaling in these cells is well-characterized and begins through binding of EPO to the homodimeric EPO receptor (EPO-R/EPO-R) (Broudy *et al*., 1991; Philo *et al*., 1996; Watowich, 1999). Ligand binding induces structural changes in the cytoplasmic domains of EPO-R, triggering transactivation of JAK2 and then STAT5 (Witthuhn *et al*., 1993; Chin *et al*., 1996; Klingmüller *et al*., 1996). Once STAT5 is phosphorylated by JAK2, it translocates to the nucleus where it activates expression of genes necessary for the survival, proliferation and differentiation of erythroid cells, including the anti-apoptotic proteins Bcl-2, Bcl-XL (Wojchowski *et al*., 1999; Watowich, 2011). Due to its ability to promote erythropoiesis, EPO and its derivatives (e.g., Epoetin alfa, Darbepoetin alfa) are FDA-approved drugs for the treatment of anemia resulting from chronic kidney disease or chemotherapy.

EPO signaling also drives activated platelet generation, expression of pro-thrombotic proteins, protection of non-hematopoietic cells against hypoxic stress, and nitric oxide production by the vascular epithelium (Lippi *et al*., 2010; Sautina *et al*., 2010; Lee *et al*., 2020; Sergio and Rolando, 2022). To explain these diverse responses, we have proposed previously that EPO mediates the body’s response to hemorrhage, simultaneously protecting tissues against reduced oxygen and promoting cessation of bleeding and the generation of new RBCs (Burrill *et al*., 2016; Lee *et al*., 2021). The protection against stress is not limited to hypoxia; EPO reduces cell death responses to a variety of stimuli. Due to its anti-apoptotic properties, there has been interest in using EPO in disease states such as traumatic brain injury, myocardial infarction, and neurodegenerative diseases such as ALS (Ghezzi and Brines, 2004; Grunfeld *et al*., 2007; Gan *et al*., 2012; Patel *et al*., 2012; Hernández *et al*., 2017; Robinson *et al*., 2018). However, the dose of EPO needed to activate protective signaling (∼1-20 nM) is generally higher than the dose necessary for its effects on erythroid and megakaryocytic cells (100-200 pM) (Anagnostou *et al*., 1990; Masuda *et al*., 1993; Brines and Cerami, 2008). At these doses, the likelihood of off-tissue EPO-R homodimer-driven effects, such as RBC and platelet generation, increases (Brines and Cerami, 2008).

Tissue-protective signaling by EPO is thought to be mediated, at least in part, by a heterodimer of EPO-R and CD131; however, the details of this signaling mode are unclear and controversial (Brines *et al*., 2004; Cheung Tung Shing *et al*., 2018). CD131, the common beta receptor (βc), is the beta subunit of the GM-CSF, IL-3, and IL-5 receptors and is crucial for the differentiation of a diverse set of myeloid cell types (Scott and Begley, 1999). There is substantial evidence that EPO-R and CD131 interact. Jubinsky *et al*. (Jubinsky *et al*., 1997) first demonstrated that CD131 functionally associates with EPO-R through co-immunoprecipitation experiments, and Hanazono *et al*. (Hanazono *et al*., 1995) observed phosphorylation of CD131 in response to EPO. Moreover, Sautina *et al*. (Sautina *et al*., 2010) found that co-immunoprecipitation of EPO-R with anti-CD131 was enhanced by EPO. However, the structure of the full signaling complex and how they interact is still unknown. Sautina *et al*. also found that VEGF receptor was co-immunoprecipitated with EPO-R and anti-CD131 and may be part of the full complex (Sautina *et al*., 2010), and Cheung Tung Shing *et al*. (Cheung Tung Shing *et al*., 2018) failed to find evidence of an interaction between the extracellular domains of EPO-R and CD131 with and without EPO using several biophysical binding assays (including surface plasmon resonance, microscale thermophoresis and analytical size-exclusion chromatography). The findings of the aforementioned studies are not necessarily in conflict with each other as an interaction could still happen between the transmembrane segments of these receptors. This mechanism is supported by work from He *et al*. that found that the extracellular domains of each receptor are not required for downstream signaling through EPO-R/CD131 (He *et al*., 2019). In sum, the composition and structure of an EPO-R/CD131 complex and how these proteins interact to induce downstream signaling is still controversial.

To selectively activate EPO-R/CD131 many groups have developed EPO derivatives that putatively reduce or eliminate activation of homodimeric EPO-R, such as carbamylated EPO (cEPO) (Leist *et al*., 2004), EPO mutants with lower affinity for EPO-R (Leist *et al*., 2004; Gan *et al*., 2012; Lee *et al*., 2021), smaller peptide derivatives of EPO (Brines *et al*., 2008; Pankratova *et al*., 2012; Ercan *et al*., 2018; Cho *et al*., 2022) or small EPO-mimetic peptides (Wrighton *et al*., 1996; He *et al*., 2019). The most studied EPO alternative is carbamylated EPO, in which all amino groups on the ten lysines have a carbamyl modification. cEPO has been proposed as a neuroprotective therapeutic that does not produce red blood cells or platelets and is claimed to signal only through EPO-R/CD131 and not through EPO-R homodimers (Leist *et al*., 2004).

However, to the best of our knowledge, the quantitative effect of carbamylation on EPO’s binding to EPO receptor or CD131 has not been characterized (e.g., by surface plasmon resonance), nor has its activity been rationalized in terms of solved structures of CD131 or EPO and EPO-R.

Additionally, cEPO has the same plasma half-life as EPO itself – about eight hours (Macdougall *et al*., 1989). In the treatment of anemia, EPO can be used intermittently because treatment produces a burst of erythroid progenitor activity that produces red blood cells lasting 100 days. However, neuroprotection by EPO signaling may require continuous exposure, which may require genetically fusing a hormone to a module that extends half-life such as albumin or the Fc region of an antibody. Rational design of such an engineered protein would require a structural understanding of how EPO, EPO-R and CD131 interact. The focus of the present work is to explore the structure of the EPO-R/CD131 signaling complex at a level sufficient to construct engineerable fusion proteins that will activate EPO-R/CD131 selectively over EPO-R/EPO-R.

## Results

### Rational design of tandem scFv proteins for selective activation of EPO-R/CD131

We aimed to develop selective agonists of EPO-R/CD131 that do not activate EPO-R/EPO-R. Given the mechanisms of complex formation of the GM-CSF receptor (Carr *et al*., 2001; Hansen *et al*., 2008; Caveney *et al*., 2024) and EPO-R/EPO-R (Watowich *et al*., 1994, 1999; Syed *et al*., 1998), we hypothesized that binding EPO-R and CD131 using a bispecific molecule would be sufficient for pathway activation through induced proximity. To explore how binding flexibility and orientation may affect receptor activation, we designed three fusion proteins with distinct structural configurations. Each protein is bispecific and designed to bind both EPO-R and CD131. We first generated a tandem scFv, whereby two scFvs are joined by a linker to generate a single-chain bispecific binder. This protein (denoted as TscFv) consists of two previously reported high-affinity scFvs (Borges *et al*., 2008; Moraga *et al*., 2015; Panousis *et al*., 2016) connected with a flexible glycine, serine, and glutamic acid linker (Figure 1A). We then fused either the entire TscFv or the individual scFvs to the fragment crystallizable domain (Fc) of mouse IgG2a (Figure 1A) since Fc fusion can improve the stability, manufacturability, and serum half-life profiles of biologics. The first Fc fusion protein is a bispecific antibody with each scFv fused to one half of Fc (denoted as BsAb), and the second Fc fusion consists of the entire TscFv linked to one half of Fc (denoted as TscFv-Fc) (Figure 1A). To ensure both halves of the bispecific Fc fusion proteins are present, we used a previously published heterodimeric Fc variant that favors heterodimer formation through electrostatic steering (Wang *et al*., 2019). This improves the likelihood of retaining both halves of Fc after purification. The desired signaling of these three molecules is depicted in contrast to endogenous signaling of EPO in Figure 1B.

**Figure 1.**
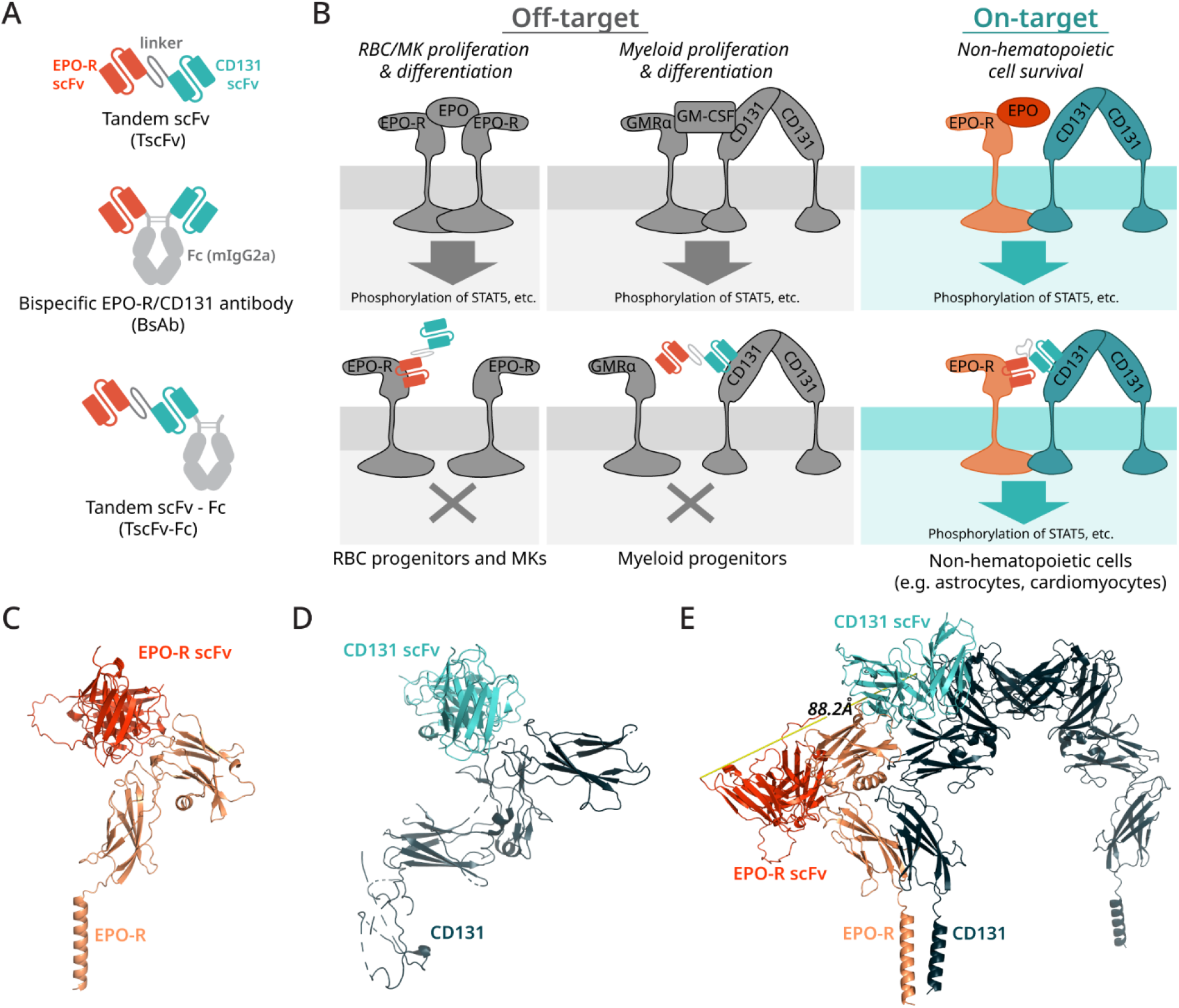
Design rationale for tandem scFvs that selectively bind EPO-R/CD131. **A**. Schematic of the three bispecific fusion proteins explored in this work. Each consists of a previously published EPO-R binding scFv (Borges *et al*., 2008; Moraga *et al*., 2015) and a previously published CD131 binding scFv (Panousis *et al*., 2016). The first protein (TscFv) is a tandem scFv whereby the EPO-R scFv and the CD131 scFv are joined by a long linker to make a single chain, bispecific binder. The other two proteins consist of the binding moieties fused to the fragment crystallizable (Fc) region of mouse IgG2a. Fusion to Fc is expected to enhance half-life due to an increase in size (and therefore a decrease in the rate of renal clearance) and FcRn-mediated recycling (Kontermann, 2011). One Fc fusion consists of the scFvs individually fused to the N terminus of Fc, forming a bispecific antibody (BsAb), and the other consists of the entire TscFv fused to one half of Fc (TscFv-Fc). **B**. Schematic of endogenous signaling through EPO-R and CD131 compared to the desired pathway activation by our fusion proteins. EPO naturally signals through either homodimeric EPO-R (which mediates RBC and platelet generation) or EPO-R/CD131. Our fusion proteins are designed to eliminate the activation of EPO-R homodimers to mimic EPO-induced activation of EPO-R/CD131 with enhanced selectivity. **C**. Solved structure (PDB 4y5y) of the EPO-R scFv bound to EPO-R (Moraga *et al*., 2015) ***D***. Solved structure of the CD131 scFv (PDB 5dwu) bound to one monomer of CD131 (Panousis *et al*., 2016). ***E***. Alphafold 3 (Abramson *et al*., 2024) and PyMOL model of the potential structure of the tandem scFv in complex with EPO-R/CD131. We used Alphafold 3 to predict the interaction of each scFv with hEPO-R and (CD131)_2_. Then we aligned both structures using the common structural elements (i.e., hEPO-R and (CD131)_2_) using PyMOL. The resulting structure is one possible prediction of how EPO-R and CD131 may interact. In this model the C-terminus of the EPO-R scFv is 88 angstroms away from the N-terminus of the CD131 scFv. We used this model to design the linker to our tandem scFv, ensuring it could span that distance.

Since there is no structural information for a hypothetical EPO-R/CD131 complex, we used available structural data of the scFvs binding to their respective receptors (**Figure 1C, 1D**) and AlphaFold 3 (Abramson *et al*., 2024) to model the potential interaction of EPO-R/CD131 and TscFv (**Figure 1E**). Specifically, we first generated one model of anti-EPO-R/EPO-R/(CD131)_2_, and a separate model of anti-CD131/EPO-R/(CD131)_2_ using AlphaFold. We included two copies of CD131 since it naturally forms a non-signaling dimer in which the two transmembrane segments are separated by 90-140 Å (Carr *et al*., 2001). In the two predicted complexes, EPO-R and (CD131)_2_ have a similar geometry as in these structures. We aligned the complexes using PyMol so that the EPO-R and (CD131)_2_ molecules in each model were maximally aligned, and then we inferred the relationship of the anti-EPO-R and anti-CD131 scFvs as a hypothetical geometry to be tested. Using this structure, we measured an approximate distance the scFvs could be in the fully formed complex, which informed our linker design. In this model, the C terminus of the EPO-R scFv is positioned 88.2 Å from the N terminus of the CD131 scFv (**Figure 1E**). Assuming an average amino acid contour length of 4 Å, (Ainavarapu *et al*., 2006) this distance could be covered by a ≥22 amino acid (aa) linker. Therefore, our first TscFv design connects the EPO-R and CD131 scFvs with a 23 aa linker consisting of glycine, serine and glutamic acid to allow unimpeded long-range movement of each scFv and to increase overall solubility. The spacing of the scFvs in BsAb also should be sufficient to allow simultaneous binding since the distance between the Fabs of wildtype mouse IgG2a, is 117 ± 37 Å (Sosnick *et al*., 1992).

The proposed model has features that are consistent with past observations about a putative EPO-R/CD131 interaction. In the structural model, EPO-R associates with CD131 along a face away from the EPO binding site, such that if EPO were bound to EPO-R in this complex, EPO would not contact CD131. Such an interaction might nonetheless be promoted by EPO if we envision that unliganded EPO-R exists primarily in a self-inhibited dimeric state implied by the crystal structure (Livnah *et al*., 1999) in equilibrium with an unliganded monomer. In the unliganded dimer, EPO-R interacts with itself by the same surface that binds to EPO, and the geometry of this interaction would force the EPO-R dimer to lie flat against the cell membrane. Additionally, EPO would bind to the unliganded monomer and prevent re-formation of the inactive dimer so the EPO/EPO-R complex could then interact with CD131. This model would explain how high concentrations of EPO variants with a mutation in the “weak face” surface that interacts with the second EPO-R could still signal through EPO-R and CD131. We explore this model further in the Supporting Information (Figure S1, S2). Concurrently, we also generated models using AlphaFold 3 for interacting extracellular domains of EPO-R and CD131, with and without EPO or the TscFv (Figure S3). These models all showed EPO-R in proximity to the most membrane-proximal domain of CD131, but differed in specific contacts. This suggests that the extracellular domains of EPO-R and CD131 might interact weakly or not at all, and that their interaction may require contacts between the transmembrane and/or intracellular domains.

We first assessed whether fusion of the scFvs to each other or to Fc would hinder their binding. We expressed and purified each protein and verified the presence of heterodimers in the case of the Fc fusions (Figure S4E and S4F). Then, we determined whether the constructs bind EPO-R and CD131 by elisa. TscFv bound both receptors with similar EC_50_s (Figure 2A, 2B), demonstrating that TscFv folds properly and the tandem scFv format is flexible enough to allow proper receptor binding.

**Figure 2.**
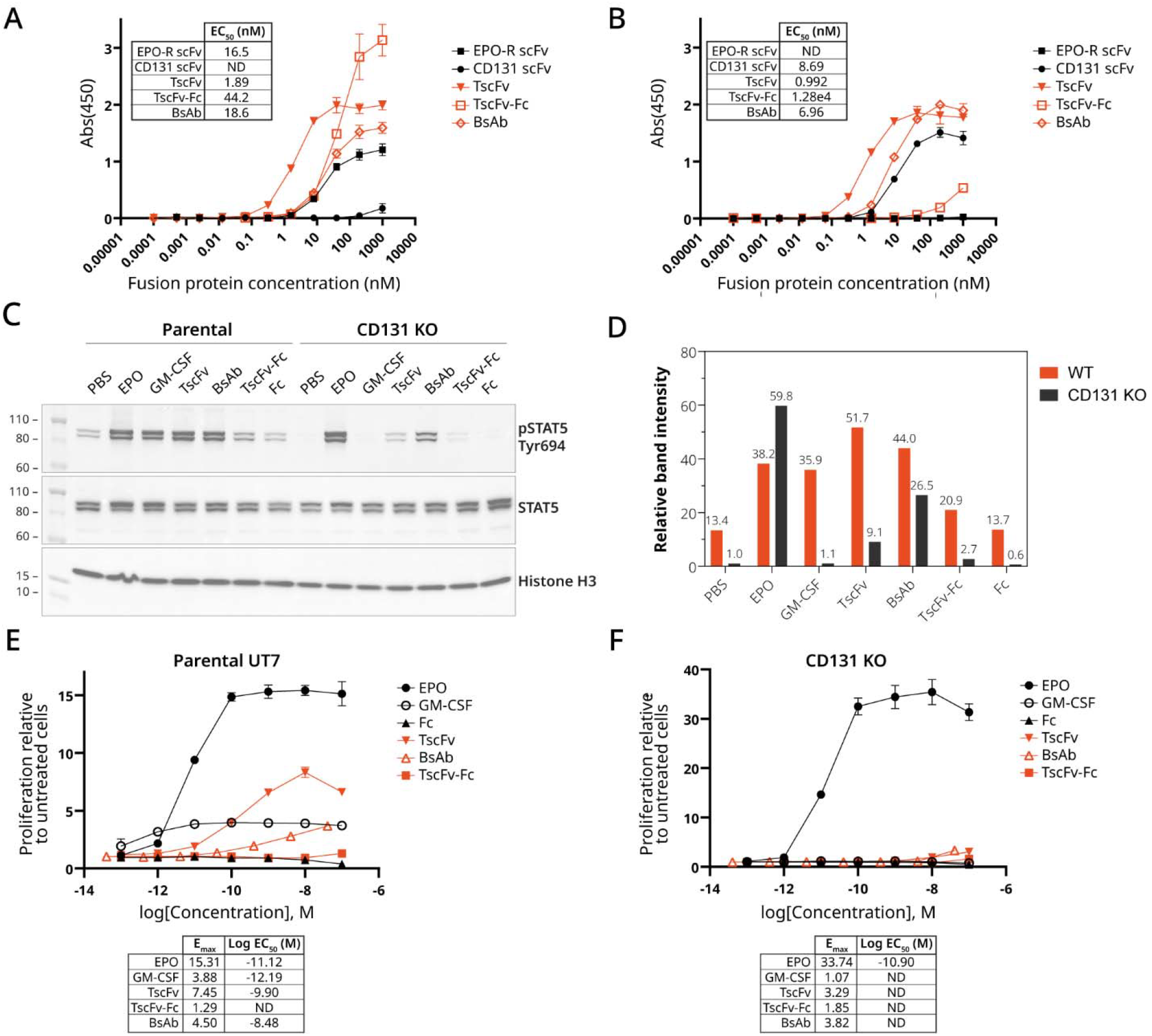
Tandem scFv and BsAb bind EPO-R and CD131 and activate downstream CD131-dependent STAT5 phosphorylation and proliferation in UT-7-ES1. **A-B**. Elisa of each of the bispecific proteins binding to EPO-R or CD131. Recombinant **A**. EPO-R or **B**. CD131 were used to coat elisa plates and relative binding of each protein to both receptors was determined. Elisa data are reported as mean A_450_ ± SD of three replicates. EC_50_ values were calculated using GraphPad Prism. **C**. All fusion proteins activated the phosphorylation of STAT5 at Tyr694. UT-7 EPO S1 (UT-7-ES1) is a cell line that is dependent on signaling through EPO-R/EPO-R, EPO-R/CD131, or GMCSFRα/CD131 to survive. Parental UT-7-ES1 or a UT-7-ES1 clonal CD131 KO line were FBS starved overnight, treated with 100 nM proteins for 15 min, and lysed in the presence of protease and phosphatase inhibitors. Proteins were run by SDS-PAGE and visualized by western blot with the designated antibodies. **D**. Quantification of the density of sample bands in **C** relative to PBS treated sample from the CD131 KO line. **E**. TscFv and BsAb support growth of parental UT-7-ES1. UT-7-ES1 were cytokine starved overnight, treated with the designated proteins and viability was measured 72 hrs later. **F**. TscFv and BsAb do not support growth of UT-7-ES1 with CD131 KO. A clonal CD131 KO UT-7-ES1 line was cytokine starved overnight, treated with the designated proteins and viability was measured 72 hrs later. Proliferation data are reported as proliferation relative to untreated cells and are represented by mean luminescence ± SD of three replicates. EC_50_ values were calculated using GraphPad Prism.

In contrast, both Fc fusion constructs (BsAb and TscFv-Fc) had at least an order of magnitude lower apparent binding affinity for either receptor when compared to TscFv.

TscFv-Fc and BsAb exhibited similar binding to EPO-R (**Figure 2A**), with EC_50_s of 44.2 nM and 18.6 nM respectively. However, TscFv-Fc has severely impaired binding (12,800 nM) to CD131 compared to the other proteins (**Figure 2B**), suggesting steric hindrance from the Fc or the lack of flexibility of that scFv due to its fusion to the C terminus to Fc. The diminished binding of the BsAb and TscFv-Fc molecules is explained further in **Supplementary Figure 5**. Overall, TscFv and BsAb are expressed and purified to titers of 5-10 mg/L and bind both EPO-R and CD131 as expected.

### EPO-R/CD131 tandem scFv and BsAb induce CD131-dependent STAT5 signaling and proliferation in UT-7-ES1 cells

To determine if our proteins could activate EPO-R signaling, we first assessed their ability to support the growth of an EPO-responsive cell line, UT-7 EPO S1 (herein referred to as UT-7-ES1). UT-7-ES1 (Shimomura *et al*., 1992; Komatsu *et al*., 1993) expresses both EPO-R and CD131 and is dependent on either EPO or GM-CSF for growth. Consequently, signaling from EPO-R/EPO-R, EPO-R/CD131, or GM-CSFRα/CD131 could be sufficient to support proliferation of these cells. Western blotting to detect phosphorylated STAT5 (Tyr694) revealed faint bands at the correct molecular weight for STAT5A and STAT5B in the unstimulated cell line, indicating some level of basal STAT5 activation in UT-7-ES1.

As expected, the natural cytokines that support cell proliferation, EPO or GM-CSF, induced approximately 3-fold higher STAT5 phosphorylation than both negative controls (PBS and Fc) (**Figure 2C, 2D**). Both TscFv and BsAb also activated STAT5 phosphorylation, inducing 3 to 4-fold higher phosphorylation of STAT5 when compared to the PBS or Fc treated samples respectively (**Figure 2C, 2D**). TscFv-Fc elicited less STAT5 phosphorylation than TscFv and BsAb (**Figure 2C, 2D**), possibly due to its impaired CD131 binding at this concentration compared to TscFv and BsAb (**Figure 2B**). In sum, EPO, GM-CSF, TscFv, and BsAb all induced STAT5 phosphorylation in the parental UT-7-ES1 cell line.

Since STAT5 phosphorylation can be activated through either EPO-R/EPO-R or EPO-R/CD131, we assessed whether CD131 expression was required. We used standard CRISPR-Cas9 gene-editing to generate a clonal UT-7-ES1 CD131 knockout (Clone C; **Figure S6**). This knockout line has a 25-bp deletion in exon 4 (**Figure S6D**) which is in the first domain of CD131, the main extracellular domain that interacts with GM-CSF. A western blot using an anti-CD131 antibody against parental UT-7-ES1 revealed three bands (**Figure S6A**), and the two high-molecular weight bands are knocked out in the UT-7-ES1 Clone C cell line (**Figure S6B**). In this knockout line, CD131 KO eliminated the basal STAT5 phosphorylation of untreated samples that we observed in parental UT-7-ES1 (**Figure 2C, 2D**).

As expected, treatment with GM-CSF no longer induced STAT5 phosphorylation in UT-7-ES1 clone C (**Figure 2C, 2D**), verifying functional knockout of CD131 in this cell line. Also as expected, STAT5 phosphorylation induced by EPO is not reduced in the CD131 KO cell line since signaling can still occur through homodimeric EPO-R (**Figure 2C, 2D**). Each engineered bispecific construct induced some level of STAT5 phosphorylation in this line despite the lack of functional CD131. However, STAT5 phosphorylation was markedly reduced, and all three bispecific constructs induced less STAT5 phosphorylation than EPO (**Figure 2C, 2D**). The weak remaining induction of STAT5 phosphorylation by our proteins could be due to binding of the anti-CD131 scFv element to a fragment of CD131 in the clone C cell line (**Figure S6B, S6D**).

Next, we determined whether the stimulation provided by our fusion proteins was able to support proliferation of UT-7-ES1. As expected, cytokine-starved parental UT-7-ES1 cells showed dose-dependent survival after treatment with EPO or GM-CSF (**Figure 2E**). TscFv induced strong dose-dependent survival of UT-7-ES1, although the potency and efficacy was less than EPO (**Figure 2E**). This could be because EPO can induce signaling through both EPO-R homodimers and EPO-R/CD131, while TscFv should only be able to signal through EPO-R/CD131. BsAb also supported dose-dependent survival of UT-7-ES1. However, BsAb is less potent than TscFv and only supports modest growth at the three highest concentrations tested. Finally, consistent with the binding and signaling data, TscFv-Fc does not support proliferation of UT-7-ES1. These data demonstrate that the natural ligands, EPO and GM-CSF, and our proteins, TscFv and BsAb, are able to support dose-dependent proliferation of UT-7-ES1.

In the UT-7-ES1 CD131 KO line, EPO, but not GM-CSF, supported proliferation (**Figures 2F, S6C**), further verifying the functional knockout of CD131. However, in contrast to the parental UT-7-ES1 cell line, CD131 KO cells treated with TscFv and BsAb did not proliferate at any dose tested (**Figure 2F**). TscFv-Fc, did not support proliferation of either Parental or CD131 KO UT-7-ES1 (**Figure 2E, 2F**) which is consistent with the STAT5 phosphorylation (**Figure 2C, 2D**) and CD131 binding (**Figure 2B**) data. In sum, TscFV and BsAb induce CD131-dependent STAT5 phosphorylation and proliferation in UT-7-ES1.

### EPO-R/CD131 tandem scFv and BsAb selectively activate EPO-R/CD131 but not EPO-R/EPO-R

To further investigate the pathway selectivity of the tandem scFv proteins, we next compared treatment responses in engineered human embryonic kidney (HEK293T) cell lines overexpressing EPO-R, CD131 or both EPO-R and CD131 (**Figures 3A, S7**). Parental HEK293T do not express either EPO-R or CD131. We transduced HEK293T cells with lentiviral expression constructs encoding EPO-R, CD131 or both receptors to yield pooled cell lines with stable receptor expression. Each cell line was transduced independently, and they are not lineally related (i.e., the cell line expressing both EPO-R and CD131 is not a derivative of either the EPO-R or CD131 only line). Both receptors were FLAG-tagged to verify expression of each receptor (**Figures 3B, 3C, S7**). The cell line expressing EPO-R can only signal through homodimeric EPO-R, while the line that expresses both EPO-R and CD131 can signal through both homodimeric EPO-R and EPO-R/CD131 signaling. Comparing the treatment responses of these two cell lines enables the determination of selectivity by showing whether CD131 is required for activation (**Figure 3A**).

**Figure 3.**
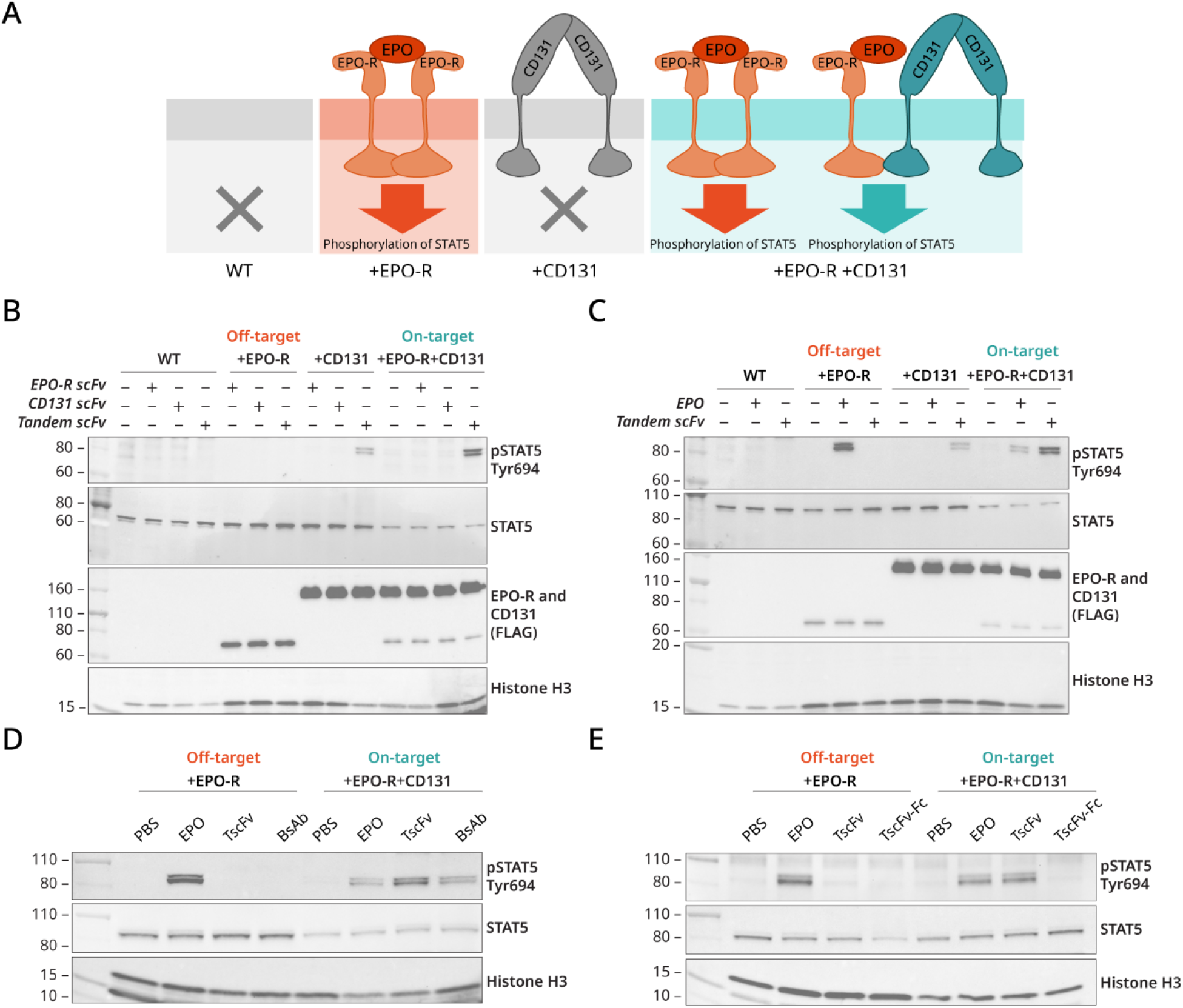
Tandem scFv and BsAb selectively activate STAT5 phosphorylation downstream of EPO-R/CD131 but not homodimeric EPO-R. **A**. Schematic of signaling in each overexpression line. HEK293T cells were stably transduced with FLAG-tagged expression constructs of either hEPO-R, hCD131, or both. Comparing the response between cells with both EPO-R and CD131 (which will have EPO-R/EPO-R and EPO-R/CD131 signaling) to cells with only EPO-R (which only can signal through EPO-R/EPO-R) can illuminate EPO-R/CD131-specific signaling. **B-E**. Cells were FBS starved overnight, treated with the designated proteins at 100 nM for 15 min, and lysed in the presence of protease and phosphatase inhibitors. Proteins were run by SDS-PAGE and visualized by western blot with the designated antibodies. **B**.TscFv, but not the individual scFvs, triggers CD131-dependent STAT5 phosphorylation. **C**. TscFv has a different signaling profile than EPO and does not activate EPO-R/EPO-R. **D**. BsAb is also only able to induce STAT5 phosphorylation in cells that express CD131. **E**. TscFv-Fc does not activate STAT5 phosphorylation downstream of EPO-R/EPO-R nor EPO-R/CD131.

Additionally, unlike UT-7-ES1, HEK293T do not express the alpha subunit of any CD131 receptor, so treatment response in these lines cannot be complicated by activation of related CD131-containing receptors.

As expected, the unfused anti-EPO-R and anti-CD131 scFvs did not induce STAT5 phosphorylation in any cell line (**Figure 3B**), most likely due to the inability of a monomeric scFv to induce receptor dimerization. Also as expected, EPO induced STAT5 phosphorylation in both EPO-R and EPO-R/CD131 expressing cell lines (**Figure 3C**), confirming functional EPO-R signaling in these lines. However, in contrast, TscFv induced strong STAT5 phosphorylation only in cells that express both EPO-R and CD131, but not the EPO-R-only cell line (**Figure 3B**), demonstrating specificity to heterodimeric EPO-R signaling. TscFv also induced weak STAT5 phosphorylation in the CD131-only overexpression line (**Figure 3B**), which remains unexplained, but we believe it is a cell-line specific effect. To determine if this effect was cell-line dependent, we separately constructed a HEK293T cell line expressing a V5-tagged CD131 that expresses less CD131 (**Figure S7**). In this line which only expresses CD131, TscFv did not induce detectable STAT5 phosphorylation (**Figure S8**). Finally, BsAb also demonstrated specificity as treatment only induced STAT5 phosphorylation in the cell line expressing EPO-R and CD131 (**Figure 3D**).

To determine if simply the presence of both scFvs was sufficient for signaling, we treated cells with an equimolar mix of the EPO-R and CD131 scFvs. An equimolar mix of the scFvs did not induce STAT5 signaling in either the EPO-R only or EPO-R and CD131 overexpression cell lines (**Figure S9**), suggesting simultaneous binding and linking of both receptors is necessary. Finally, consistent with the signaling and binding data, TscFv-Fc was unable to activate STAT5 phosphorylation in either line at the same dose (**Figure 3E**). In conclusion, TscFv and BsAb are most active in the EPO-R/CD131 cell line and not active in cells expressing only EPO-R, and the bispecific nature of TscFv and BsAb is necessary for activity. Therefore, induced proximity of EPO-R and CD131 by a single bispecific molecule appears to be sufficient for EPO-R/CD131 activation.

### Decreasing linker length between anti-EPO-R and anti-CD131 scFvs in TscFv modifies signaling

Based on the EPO-R/CD131/tandem scFv structural model (**Figure 1E**), we hypothesized that shortening the inter-scFv linker would reduce or eliminate downstream signaling. Specifically, in our model, the C-terminus of the anti-EPO-R scFv and the N-terminus of the anti-CD131 scFv are separated by 88 Å. The 23 aa linker in TscFv (denoted GS-L, **Figure 4**) can cover this distance when fully extended. If this model is correct, shortening the linker would likely reduce or eliminate signaling because the proper geometry of EPO-R and CD131 would be entropically unfavored or sterically infeasible.

**Figure 4.**
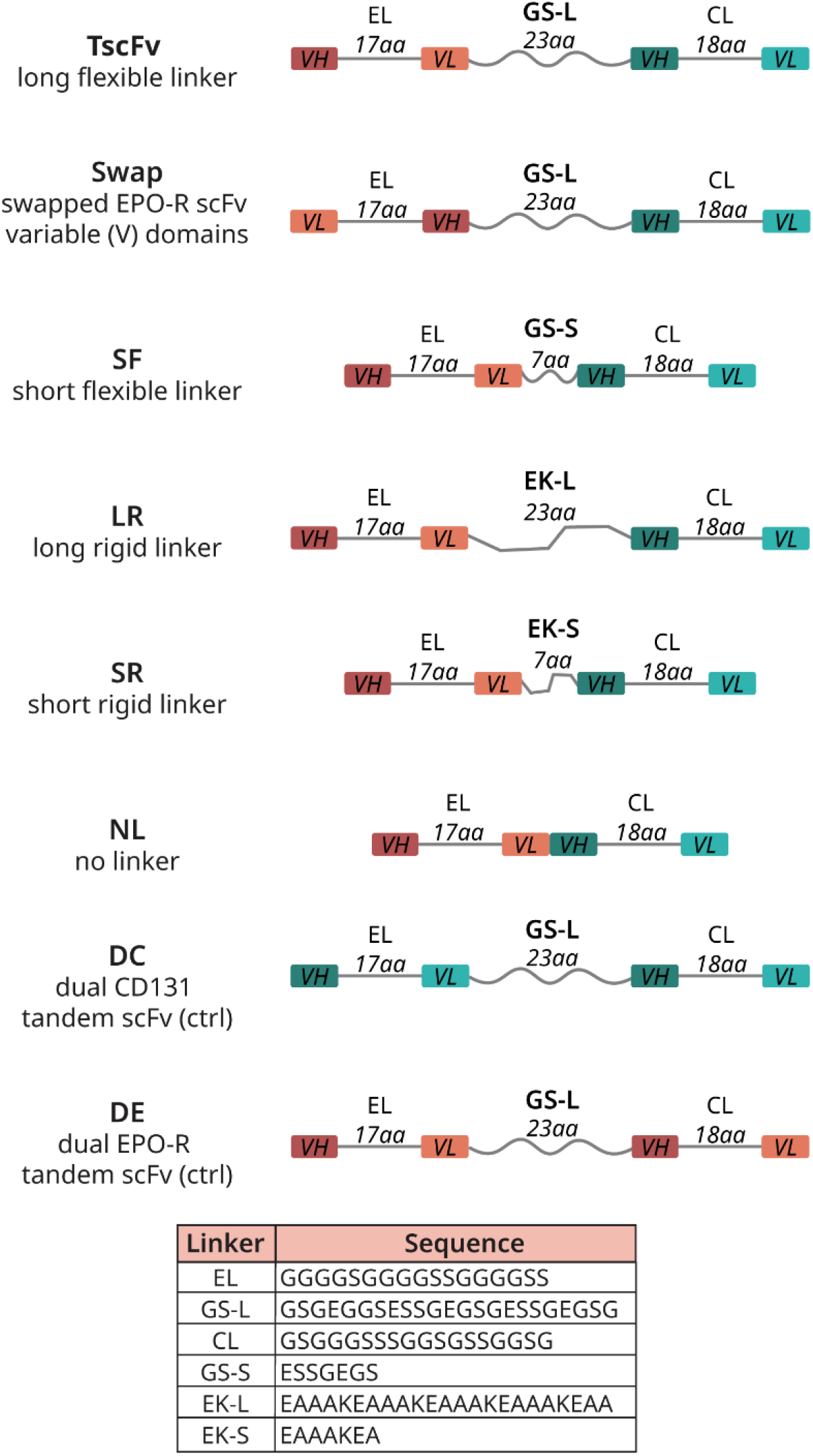
Design of tandem scFv linker and domain variants. **A**. Variants were made of TscFv with changes to the linker connecting the two scFvs or the variable (V) domain order of the EPO-R scFv. Two control tandem scFvs (TscFv-DE and TscFv-DC) were also constructed to test whether signaling was due to the bispecific binding of TscFv, not just the presence of two scFvs. The linkers connecting the variable domains of each scFv (EL and CL) were kept the same in all constructs. The sequence of each linker is listed. Abbreviations: VH=heavy chain, VL=light chain, GS=Glycine and serine linker, EK=Glutamic acid and lysine linker, L=long, S=short.

We designed several new TscFv variants with linkers of differing length and rigidity (Chen *et al*., 2013). Specifically, a tandem scFv with a 7-amino acid linker between the anti-EPO-R scFv and the anti-CD131 scFv (SF for short, flexible); a tandem scFv with a 23-amino acid rigid linker between the scFvs (LR for long, rigid); a tandem scFv with a 7-amino acid rigid linker between scFvs (SR for short, rigid); and a tandem scFv with no linker (NL) (**Figure 4**). We also rearranged the variable domains of the anti-EPO-R scFv in one variant so that its linker attachment point is further away from the anti-CD131 scFv (“Swap”) (**Figure 4**).

The linker variants with no linker, shorter linkers, or more rigid linkers (NL, SF, SR, LR) induced two to four-fold less STAT5 phosphorylation in parental UT-7-ES1 compared to the original TscFv (which has a long, flexible linker) (**Figure 5A, 5B**). STAT5 phosphorylation in the UT-7-ES1 CD131 KO cell line was greatly reduced or completely eliminated for all of the linker variants (except “Swap”), suggesting CD131 dependent signaling (**Figure 5C, 5D**). Residual activity of these constructs could be explained according to the model depicted in **Supplementary Figure 10**. Unexpectedly, the “Swap” tandem scFv induced more STAT5 phosphorylation in comparison to TscFv in the UT-7-ES1 CD131 KO line (**Figure 5C, 5D**). We believe this is due to the linker length between the two scFvs, which may force dimerization of the anti-EPO-R scFv and enable some level of homodimeric EPO-R signaling (diagrammed in **Supplementary Figure 11**).

**Figure 5.**
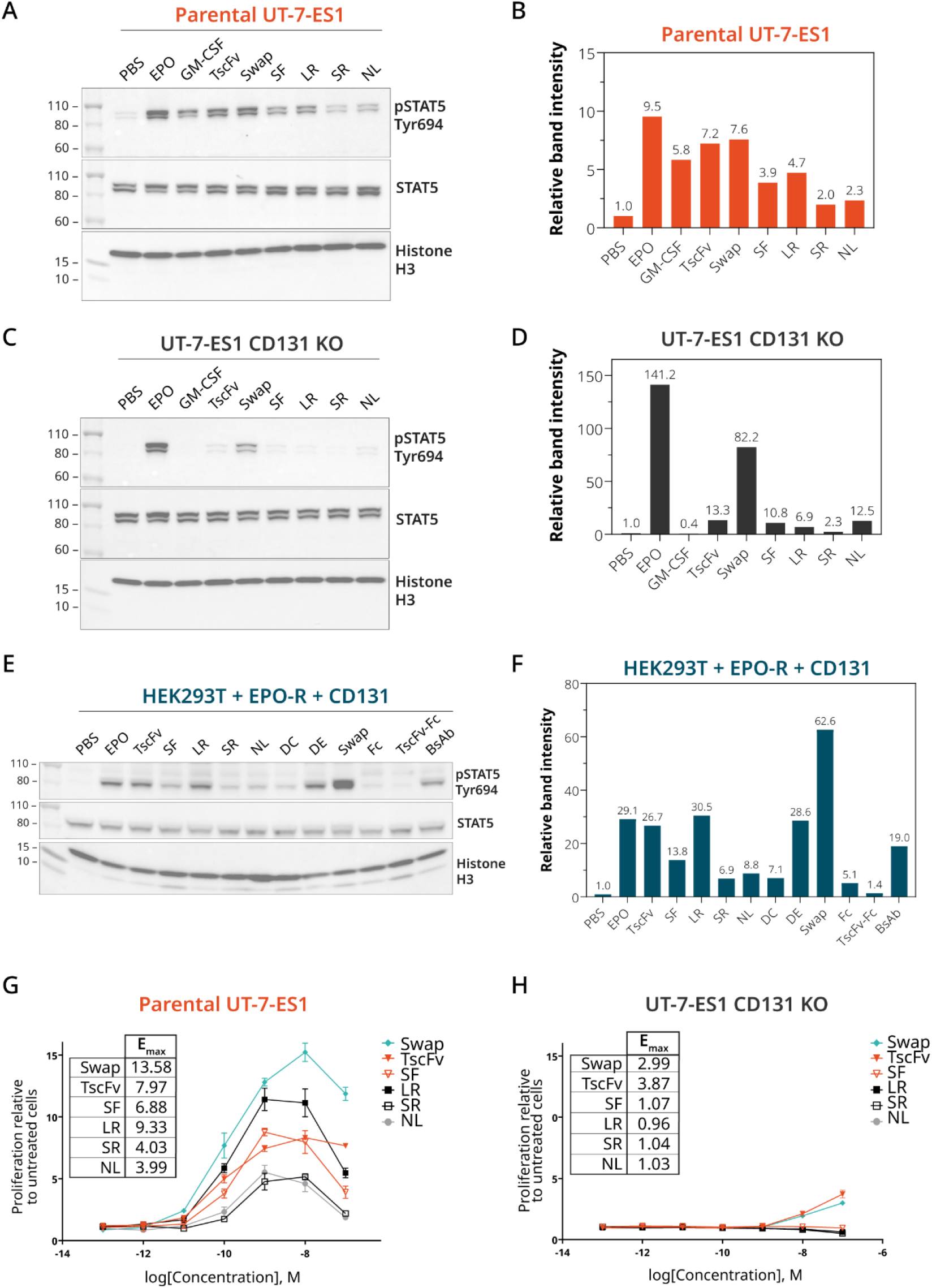
Decreased linker length between EPO-R and CD131 scFvs decreases signaling of tandem scFv variants. **A**, C. UT-7-ES1 or UT-7-ES1 CD131 KO cells were FBS starved overnight, treated with the designated proteins at 100 nM for 15 min, and lysed in the presence of protease and phosphatase inhibitors. Proteins were run by SDS-PAGE and visualized by western blot with the designated antibodies. **B, D**. Band densities of STAT5 phosphorylated at Tyr694 in **A** and **C** were quantified and plotted relative to the density of the PBS-treated control bands. **E**. HEK293T overexpressing EPO-R and CD131 were FBS starved overnight, treated with the designated proteins at 100 nM for 15 min, and lysed in the presence of protease and phosphatase inhibitors. Proteins were run by SDS-PAGE and visualized by western blot with the designated antibodies. **F**. Band densities of E relative to the PBS-treated control sample. **G**. Parental UT-7-ES1 lines were cytokine starved overnight, treated with the designated proteins at 100 nM and viability was measured 72 hrs later. TscFv and its variants are able to support UT-7-ES1 proliferation, but variants with decreased linker length induce less proliferation. H. UT-7-ES1 CD131 KO cells were treated as in G and viability was measured 72 hrs later. TscFv and linker variants are unable to support proliferation of UT-7-ES1 when CD131 is knocked out. Proliferation data are reported as proliferation relative to untreated cells and are represented by mean luminescence ± SD of three replicates. E_max_ values were calculated using GraphPad Prism.

All linker variants also induced STAT5 phosphorylation in the HEK293T line that expresses EPO-R and CD131 (**Figure 5E, 5F**). TscFv variants with longer linkers (TscFv, LR) were more efficacious than those with shorter linkers, consistent with the UT-7-ES1 results. We also designed two tandem scFv controls, DC (dual anti-CD131 scFv) and DE (dual anti-EPO-R scFv)), which each consist of two of the same scFv connected by a long, flexible linker (**Figure 4**). Since it has two EPO-R binding moieties, DE should signal through EPO-R/EPO-R, while the dual anti-CD131 scFv (DC) should not. As expected, DE induced a level of STAT5 phosphorylation that is similar to EPO, and DC did not induce strong STAT5 phosphorylation (**Figure 5E, 5F**). Comparison of the response induced by TscFv, DE, and DC further suggests that the induced STAT5 phosphorylation is not due to the presence of two linked scFvs but is specific to the presence of both EPO-R and CD131 binding moieties in the same molecule (**Figures 5E, 5F, S12**).

Finally, we tested the ability of the TscFv linker variants to support proliferation of UT-7-ES1. The maximum response (E_max_) of the variants varies markedly (**Figure 5G**). TscFv variants with shorter linkers induced less proliferation compared to the original (**Figure 5G**). As expected, DE supported cell proliferation since it can signal through homodimeric EPO-R in the UT-7-ES1 CD131 KO cell line (**Figure S12**). However, all of the linker variants induced little or no proliferation in the UT-7-ES1 CD131 KO cell line (Figure 5H), further demonstrating their specificity. Altogether, these data indicate that TscFv and its variants require CD131 to support UT-7-ES1 proliferation, suggesting they are more selective for EPO-R/CD131 than natural EPO.

## Discussion

Signaling through a combination of EPO-R and CD131 is thought to promote resistance to cell stress, such as neurons exposed to hypoxia or traumatic brain injury. In this study, we developed the first selective EPO-R/CD131 agonists. Previous work identified antibodies that bind to EPO-R (Lim *et al*., 2010; Moraga *et al*., 2015) or CD131 (Panousis *et al*., 2016) and solved the structure of these scFvs in complex with their target antigen. We used AlphaFold 3 (Abramson *et al*., 2024) to construct a model of an EPO-R/CD131 signaling complex, determined where the V regions from the anti-EPO-R and anti-CD131 antibodies would bind, and then designed and expressed a tandem scFv (TscFv) consisting of an anti-EPO-R scFv, an anti-CD131 scFv, and a linker long enough to connect them. Additionally, we constructed a bispecific antibody (BsAb) by fusing the scFvs to a heterodimeric Fc. TscFv and BsAb induced STAT5 phosphorylation through activation of EPO-R/CD131, without engaging EPO-R/EPO-R (**Figure 2, Figure 3**) and also induced cell proliferation of UT-7-ES1 in a CD131-dependent manner (**Figure 2, Figure 4**). Finally, we demonstrate that shortening the linker between the scFvs reduced signaling in a manner consistent with the predicted structural model of EPO-R/CD131. These proteins provide proof of concept for design of selective EPO-R/CD131 agonists and lay a foundation for the development of a clinical-grade EPO-R/CD131 biologic drug.

EPO and EPO variants can prevent cell death of non-hematopoietic cells in the face of cytotoxic agents or hypoxia in in vitro and in vivo models of neurodegeneration and trauma (Ghezzi and Brines, 2004; Grunfeld *et al*., 2007; Patel *et al*., 2012; Ponce *et al*., 2013; Noh *et al*., 2014; Robinson *et al*., 2018; Hemani *et al*., 2021). However, the concentration needed to generate this cell protective signaling (1-20 nM vs.100-200 pM for hematopoietic effects) would lead to undesired effects, such as RBC and prothrombotic platelet generation, due to activation of homodimeric EPO-R (Brines and Cerami, 2012). Previous efforts to make specifically tissue-protective EPOs have centered around mutated or modified EPO, such as carbamylated EPO or proteins with mutations in the face of EPO that mediates the weak interaction with EPO-R (Leist *et al*., 2004; Brines *et al*., 2008; Gan *et al*., 2012; Pankratova *et al*., 2012; Cho *et al*., 2022). These molecules fortuitously happened to have desirable properties, without a clear mechanism by which the EPO-R/CD131 signaling properties derive from the structure.

In this work, our strategy has been to construct molecules that bring EPO-R and CD131 together via scFvs that bind to each receptor. The best selective EPO-R/CD131 activator is a single polypeptide chain tandem scFv consisting of an N-terminal scFv that binds to EPO-R (Moraga *et al*., 2015), a central 23-amino acid linker, and a C-terminal scFv that binds to CD131 (Panousis *et al*., 2016), denoted TscFv. The design of this molecule was driven by a structurally plausible AlphaFold 3 model of interacting EPO-R and CD131. However, the 23-amino acid linker has a maximum extended length of about 100 Angstroms, so the molecule could accommodate many possible configurations of EPO-R and CD131.

TscFv stimulated the growth of UT-7-ES1 in a CD131-dependent manner and was not active in cells that only express EPO-R (**Figures 2E, 2F, 5G, 5H**). This is in contrast to a tandem scFv consisting of two anti-EPO -R scFvs (denoted DE; **Figure 4, S12** (**“DE”**)), which can still stimulate the growth of cells without CD131. These observations also suggest that TscFv assembles as diagrammed, and does not form dimers in which two anti-EPO-R scFvs form a diabody that could link two EPO-Rs. Specific signaling through EPO-R/CD131 was observed in two paired cell lines: UT-7-ES1 parental cells compared to UT-7-ES1 CD131-knockout cells (**Figures 2, 5**), and HEK293T cells engineered to express EPO-R only or both EPO-R and CD131 (**Figures 3, 5**). When parental UT-7-ES1 cells were treated with TscFv, the level of STAT5 phosphorylation was comparable to that induced by EPO, while in the UT-7-ES1 CD131 knockout cells, only a trace of STAT5 phosphorylation was observed. The UT-7-ES1 parental cells proliferated in response to TscFv and its variants (**Figures 4, 5G, 5H**), while the UT-7-ES1 CD131 knockout cells did not proliferate at all in response to any of these proteins (**Figures 2E, 2F, 5G, 5H**). Similarly, TscFv stimulated STAT5 phosphorylation in HEK293T cells expressing both EPO-R and CD131, but not cells expressing only EPO-R (**Figure 3**).

Based on our results, we envision that a practical therapeutic protein that specifically activates EPO-R/CD131 complexes could be constructed. Such proteins could be used to treat neurological disorders, such as traumatic brain injury (Robinson *et al*., 2018), ALS (Noh *et al*., 2014; Lauria *et al*., 2015), and possibly also Alzheimer’s disease (Kalluru *et al*., 2024). To be useful in these chronic conditions, such a molecule should not induce autoantibodies to endogenous EPO and should have a long plasma half-life. Repeated administration of EPO-based scaffolds introduces the risk of triggering an autoimmune response to endogenous EPO (Casadevall *et al*., 2005; Tamilvanan *et al*., 2010). In contrast, our constructs will not induce autoantibody production to EPO as they differ significantly in structure.

Pharmacokinetics represents a further challenge. Human EPO has a plasma half-live of only a few hours, but nonetheless has been a successful drug since the major use of EPO is to treat kidney failure patients who are on dialysis (Kalantar-Zadeh, 2017). Typically, such patients undergo dialysis three times per week, and IV administration of EPO through the line already set up for the dialysis is practical in this context. Moreover, in treating anemia it is not essential to have continuous exposure to EPO, as long as there is production of red blood cells, which have a lifetime of about 100 days. However, in treating a neurodegenerative disorder, continuous EPO signaling may be necessary. Boesch *et al*. (Boesch *et al*., 2007, 2008) performed a pilot clinical trial on 12 patients with Friedreich’s ataxia and saw a favorable effect on biomarkers; the patients received EPO three times per week, the same dose that a dialysis patient would receive, but a dose that likely enhances thrombotic events due to induction of EPO-R/EPO-R signaling. In subsequent trials of EPO and carbamylated EPO for Friedreich’s ataxia, patients were dosed once every two to three weeks (Mariotti *et al*., 2012; Boesch *et al*., 2014). Additionally, Lauria *et al*. (Lauria *et al*., 2015) clinically tested EPO as a treatment for ALS, also dosing once per two weeks. Given EPO’s plasma half-life in humans of about 8 hrs, the patients’ nervous systems would be unprotected for most of each dosing period. Our anti-EPO-R/anti-CD131 tandem scFv has a much larger Stoke’s radius than EPO and carbamylated EPO, which should lead to a superior plasma half-life. Moreover, current fusion protein technologies allow for profound extension of plasma half-life, such that high levels of a protein drug can be maintained for several weeks. With this in mind, we designed BsAb, which consists of the EPO-R and CD131 scFvs fused to Fc. BsAb is also selective for EPO-R/CD131 and will likely have a superior plasma half-life due to its size and Fc receptor (FcRn)-mediated recycling (Kontermann, 2011). However, before such technologies can be applied successfully, it is necessary to have a structural understanding of how the active protein element binds to its targets, as we have sought to reveal in the present experiments.

In conclusion, we report the rational design of several bispecific selective protein agonists of EPO-R/CD131. These fusion proteins serve as a foundation for future studies of EPO-R/CD131 biology and the development of next-generation cytokine mimetics with improved specificity and reduced off-target effects.

## Methods

### DNA construct design and generation

The anti-EPO-R scFv sequence was obtained from Moraga et. al (Moraga *et al*., 2015) (PDB 4y5y) and the anti-CD131 scFv sequence was obtained from Panousis et. al (Panousis *et al*., 2016) (PDB 5dwu). These sequences were codon-optimized and cloned into pSecTag2a. An engineered heterodimeric fragment crystallizable (Fc) domain of mouse IgG2a (H4A, H4B) (Wang *et al*., 2019) was used in the Fc fusion constructs. The Fc fusions were tagged with His6 on one monomer (H4B) to increase the likelihood of purification of only heterodimeric proteins. All protein expression constructs included a secretion signal to enable purification in mammalian cells. Sequences for the spacers of the gRNAs for CD131 KO were designed using the Custom Alt-R™ CRISPR-Cas9 guide RNA tool from IDT against the human CSFR2B locus. Complete sequences of human EPO-R and human CD131 were taken from UniProtKB ID P19235 and UniProtKB ID P32927 respectively and human codon-optimized using Benchling.

Plasmids were constructed using multiple fragment isothermal assembly using NEBuilder® HiFi DNA Assembly Master Mix (NEB E2621L) or were ordered as Twist clonal genes. Protein expression plasmids were assembled into pSecTag2a or pOptivec (with added secretion signal from the V-J2-C region of the mouse Ig kappa-chain) to enable protein secretion into the media. All secreted proteins were His6-tagged to enable purification. Overexpression constructs and CRISPR gene editing plasmids were assembled into third-generation lentiviral plasmid backbones for packaging into lentivirus. All plasmids were sequence-verified before use via whole plasmid long-read sequencing (Plasmidsaurus). See Supporting Information for more information on each plasmid (**Supplementary Table 1**), protein sequences (**Supplementary Table 6**), full plasmid sequence files and more detailed information on cloning procedures.

### AlphaFold structural prediction

The final predicted structure of the tandem scFv, hEPO-R and hCD131 (**Figure 1E**) was constructed using a combination of AlphaFold 3 (Abramson *et al*., 2024) and the pyMOL structural align feature. Initially, we generated an AlphaFold 3 prediction of the entire complex (tandem scFv/EPO-R/CD131). However, the resultant predicted structure did not seem biologically plausible. In this model, the transmembrane domain of hEPO-R was sticking perpendicular to the transmembrane domain of hCD131, and the structures did not align well with the solved PDB structures (PDB: 4y5y and 5dwu) of each scFv bound to its respective receptor. Instead, AlphaFold 3 sequences of the EPO-R scFv, hEPO-R and two copies of hCD131 were input in AlphaFold 3 to generate a predicted structure. Then, the same was done for the CD131 scFv, hEPO-R and two copies of hCD131. Using the pyMOL align feature, these two structures were combined using the common elements of hEPO-R and 2 hCD131 to align the structures to yield the final structure reported in **Figure 1E**. The measurement feature in PyMOL was used to measure the distance between the termini of the scFvs to design the tandem scFv linker and linker variant proteins (**Figure 1, 4**). See the Supporting Information (**Supplementary Figures 1-3**) for a more detailed discussion of model generation and features.

### Cell culture

HEK293T (ATCC CRL-11268) were obtained from Dr. Rui Tong Quek (Harvard Medical School) and cultured in DMEM (ATCC 30-2002) supplemented with 10% (v/v) fetal bovine serum (FBS; Thermo 10082147). Stable HEK293T overexpression lines expressing either hEPO-R, hCD131 or both hEPO-R and hCD131 were grown in HEK293T complete media as described above supplemented with 10 μg/mL blasticidin (Thermo A1113903) and/or 2.5 μg/mL puromycin (Thermo A1113803). FreeStyle 293-F cell lines (Thermo R79007) were obtained from Dr. Timothy Chang (Harvard Medical School) and cultured in FreeStyle 293 Expression Medium (Thermo 12338026) shaking at 130 rpm in sterile, vented cap Erlenmeyer cell culture shaker flasks (Corning CLS431143-50EA or CLS431145-25EA). UT-7 EPO S1 (UT-7-ES1) were obtained from Dr. R Rogers Yocum and were grown in IMDM (no phenol red) (Thermo 21056023), 10% HI FBS (Thermo 10082147), 1X GlutaMAX™ Supplement (Thermo 35050079), and 20ng/mL human EPO (Peprotech 100-64). UT-7-ES1 with stable CD131 KO were cultured in complete UT-7-ES1 media supplemented with 10 μg/mL blasticidin (Thermo A1113903) and 2.5 μg/mL puromycin (Thermo A1113803).

HEK293T cell lines were passaged every 3-4 days by gently washing the cell monolayer once with DPBS (Thermo 14190144) followed by lifting with TrypLE (Thermo 12605010) and reseeding at 1/10 – 1/20 dilution (∼10% confluence). FreeStyle 293-F cells and UT-7-ES1 cell lines were passaged every 3-4 days or when density reached 1 – 2 million cells/mL and directly reseeded in new flasks at 0.3 – 0.5 million cells/mL. All cell lines were maintained at 37^°^C and 5% CO_2_ in a humidified incubator. A full list of cell lines generated in this work and the corresponding figures they are used in is located in **Supplementary Table 5**.

### Lentivirus and stable cell line generation

Briefly, stable HEK293T cell lines expressing hEPO-R or hCD131 or both hEPO-R and hCD131 and the UT-7-ES1 CD131 KO line were generated by lentiviral transduction followed by selection with 10 μg/mL blasticidin (Thermo A1113903) and/or 2.5 μg/mL puromycin (Thermo A1113803) depending on the construct. Cells were transduced, incubated for 24-48 hrs and then changed into selection medium. Cell samples were collected once cells recovered to >95% viability to analyze transgene expression or gene KO via SDS-PAGE and western blot or sequencing. See the Supporting Information for more details on the transduction procedure for each of these lines. A full list of cell lines generated in this work and the corresponding figures they are used in is located in **Supplementary Table 5**.

### SDS-PAGE and Immunostaining

Protein samples or cell lysate were diluted in 4X Laemmli SDS sample buffer (Thermo J60015.AD) and diH_2_O water to yield a final 1X Laemmli concentration. Samples were boiled at 95^°^C for 5 minutes to denature proteins and then loaded on 4 to 20% Tris-Glycine gradient gels (Thermo XP04205BOX) submerged in 1X Tris-glycine running buffer (Thermo LC2675). Between 100 ng and 10 μg total protein (depending on application) were run alongside 5-10 μL of Novex™ Sharp Pre-stained Protein Standards (Thermo LC5800) for 1 hr at 200V. Gels were rinsed once with diH_2_O, stained using SimplyBlue™ SafeStain (Thermo LC6065) following the manufacturer’s protocol, and imaged using the Coomassie setting on a ChemiDoc MP (Bio-rad 12003154).

Gels for immunostaining were instead removed from the cassette, rinsed once with diH_2_O in a dish and transferred to a nitrocellulose membrane (Thermo IB23002) according to the manufacturer’s protocol using the P0 setting on the iBlot 2 gel transfer device (Invitrogen IB21001). Blots were washed twice while shaking in TBST (TBS (Thermo J60877.K3) + 0.05% Tween-20 (Millipore Sigma P9416-50ML)) and blocked for 1 hr at room temperature or overnight at 4^°^C while shaking in TBST + 1% BSA (GoldBio A-420-10). Primary antibody was added directly to the blocking solution and incubated shaking for 1 hr at room temperature or overnight at 4^°^C. Residual primary antibody was removed by washing three times in TBST. Blots were then incubated in secondary antibody diluted in TBST + 1% BSA for 1 hr at room temperature or overnight at 4^°^C while shaking. Blots were washed three times in TBST to remove unbound secondary antibody, and developed and imaged using SuperSignal West Dura Extended Duration Substrate (Thermo Scientific 34075) and the Chemi setting on a ChemiDoc MP. To quantify western blot signal, band intensities for phospho-STAT5, total STAT5, and the loading control (histone H3) were measured using the ImageLab densitometry software. Bands were background corrected individually by the software, and background-corrected intensities were normalized to the loading control from the same well to account for variations in sample loading. The corrected, normalized phospho-STAT5 signal was then normalized to the corresponding STAT5 signal to account for differences in protein expression levels across samples. Finally, all normalized values were expressed relative to the samples treated with the negative control (PBS), which was set to 1. More detail about the immunostaining protocol and full gel or blot images and analyses are provided in the Supporting Information. Antibody catalog numbers and dilutions are listed in **Supplementary Table 3**.

### Protein expression and purification

Proteins were expressed transiently in HEK293-F cells using 293-fectin (Thermo 12347019) following the manufacturer’s protocol. Flasks were incubated for 4-6 days at 37^°^C, 5% CO2 and shaking at 130 rpm to allow expression and secretion of proteins into the medium. Cells were pelleted and the cell supernatant containing the expressed proteins was isolated. His-tagged proteins were purified from the cleared cell supernatant using His60 Ni Superflow Resin and buffers (Takara 635677) following the manufacturer’s protocol. Fractions were taken at each step for later analysis by SDS-PAGE and western blotting using anti-6X His-HRP antibody (Abcam ab1187). Eluted proteins were concentrated and buffer exchanged into endotoxin-free PBS (TekNova P0300) using Amicon Ultra-15 Centrifugal Filters (Millipore-Sigma UFC901008 or UFC901024 depending on protein size) according to the manufacturer’s protocol. Protein concentration was assessed using a BCA Protein Assay Kit (Thermo 23227) following the manufacturer’s protocol. Purified, concentrated proteins were aliquoted, flash frozen in liquid nitrogen, and stored at -80^°^C until use. Aliquots were not freeze-thawed more than once to protect protein integrity. A list of all proteins generated in this study is located in **Supplementary Table 4**.

### Cell lysate generation

3-8 million HEK293T or HEK293T overexpression lines or 4 million UT-7-ES1/UT-7-ES1 CD131 KO clone C cells were washed three times using 1X TBS (Thermo J60877.K3) to remove excess cytokine and/or FBS and plated in 60mm dishes or 6-well dishes in serum and cytokine starved media (DMEM only (ATCC 30-2002) for HEK293T lines; IMDM only (Thermo 21056023) for UT-7-ES1). The next day, lysis buffer (1X RIPA (CST 9806S) supplemented with protease and phosphatase inhibitors (Thermo A32961) was prepared and chilled to 4^°^C. Wash buffer was prepared and chilled to 4^°^C by supplementing 1X TBS with protease and phosphatase inhibitors. Proteins were diluted in media without serum and cytokines to 1 μM, added to cells to yield a final in-plate concentration of 100 nM, and incubated for exactly 15 minutes at 37^°^C and 5% CO_2._. Cells were washed in wash buffer once and moved to 1.5mL tubes and pelleted by centrifugation for 2 min at 300xg. The cell pellet was resuspended in 200 – 400 μL of lysis buffer and lysed according to the manufacturer’s protocol. Protein concentration of the lysates was taken by BCA. Lysates were normalized to the same concentration before analysis by western blotting. See Supporting Information methods for more details.

### UT-7-ES1 cell proliferation assays

UT-7-ES1 or UT-7-ES1 CD131 KO clone C cell lines were washed three times using TBS to remove residual cytokines and seeded in white, flat-bottom, clear 96-well plates (Millipore Sigma CLS3610) at 9.0 x 10^3^ cells per well in 90 μL of growth medium with no growth factors (IMDM + 10% FBS only). The following day, purified proteins were serially diluted 10-fold (10^−6^ M to 10−^12^ M) in growth medium with no growth factors in 96-well plates (Corning 3641) and 10 μL of protein dilution or medium was added to the cells. Plates were gently swirled and then incubated at 37^°^C in 5% CO2 for 72 h. Cell proliferation was determined by Cell Titer Glo (Promega G7572) following the manufacturer’s protocol. Luminescence was read on a BioTek Synergy Neo HTS microplate reader using the Luminescence fiber default protocol. Background (media only wells) luminescence was subtracted from each sample and each well’s corrected luminescence was divided by the luminescence of untreated cells. Data are reported as proliferation relative to untreated cells and are represented by mean luminescence ± SD of three replicates. EC_50_ and E_max_ were calculated using GraphPad Prism using nonlinear regression (log(agonist) vs. response, variable slope (four parameters).

### Enzyme-linked immunosorbent assay (Elisa)

Recombinant human EPO-R (RND systems 963-ER) or human CD131 (AcroBio CD1-H5256-100 μg) was diluted to 1 μg/mL in PBS and 50 μL per well was added to coat MaxiSorp 96-well ELISA plates (Sigma M9410). Plates were incubated shaking overnight at 4^°^C and then washed five times in wash buffer (PBS + 0.05% Tween-20 (PBST) (Millipore Sigma P9416-50ML)). Plates were blocked with 200 μL per well of blocking buffer (PBS + 3% BSA (GoldBio A-420-10)) for 2 hrs shaking at RT and then washed five times with PBST. His-tagged proteins were serially diluted in PBS, 50 μL per well of each dilution was added, and the plate was incubated for 3 hrs shaking at RT or overnight shaking at 4^°^C. Plates were washed five times with PBST to remove unbound proteins. Detection antibody (anti-His HRP (Abcam ab1187)) was diluted 1/10,000 in PBS + 3% BSA, and 50 μL was added to each well and incubated at RT shaking for 1hr. Plates were washed five times with PBST to remove unbound antibody. 100 μL of room temperature TMB Substrate solution (Thermo 34028) was added to each well and absorbance was monitored until A_620_ reached >0.8 OD. 2M sulfuric acid was added to stop the reaction, and absorbance was measured at 450 nm. Data are reported as mean A_450_ ± SD of three replicates. EC_50_ and E_max_ were calculated using GraphPad Prism using nonlinear regression (log(agonist) vs. response, variable slope (four parameters).

## Acknowledgements

The authors thank Professor Timothy Mitchison who provided thoughtful advice on experimental direction. K.E.D. would like to thank Drs. Ethan Jones, Neil Dalvie, Rui Tong Quek, and Kasia Kready for their helpful comments on the manuscript. K.E.D would like to additionally acknowledge the Harvard Systems, Synthetic, and Quantitative Biology PhD program, the Harvard Therapeutics Graduate Program (TGP), and the FujiFilm Fellowship for funding and support.

## Author Contributions

K.E.D designed the project direction, conceptualized, designed and performed the experiments, made the structural model, and analyzed the data. K.E.D and J.C.W. created the figures and wrote the manuscript. K.E.D., J.C.W, and P.A.S. edited and approve the manuscript.

