## Supporting Information for "Rational design of selective bispecific EPO-R/CD131 agonists"

**EPO-R/CD131 agonists**

**TABLE OF CONTENTS**

Supplementary Figures	2

Extended methods	19

**Plasmid construct generation**	19

**Cell culture**	20

**Lentiviral generation**	21

**Lentiviral Transduction**	21

**SDS-PAGE**	22

**Western Blot**	23

**Protein expression**	24

**His-tag purification**	25

**Cell lysate treatment and generation**	25

Supplementary Tables	28

**Supplementary Table 1**: Plasmid list and additional sequences	28

**Supplementary Table 2:** Reagent catalog numbers	31

**Supplementary Table 3**: Antibody catalog numbers and dilutions	37

**Supplementary Table 4**: Protein name mappings	39

**Supplementary Table 5**: Generated cell lines & the figures they were used in	41

**Supplementary Table 6**: Protein sequences	43

References	56

### **Supplementary Figures**

**Supplementary discussion of the generation of a EPO-R/CD131/tandem scFv complex**

AlphaFold 3 (Abramson *et al.*, 2024) was used to generate the structure of the tandem scFv bound to a complex of EPO-R and CD131. The amino acid sequences of the extracellular domain and transmembrane segment of hEPO-R and two copies of hCD131 were submitted to AlphaFold 3 along with the sequence for the EPO-R scFv to generate a predicted structure of the scFv binding to the receptor complex (**Supplementary Figure 1A**). Then, the same was done for the CD131 scFv, hEPO-R and two copies of hCD131 (**Supplementary Figure 1B**). Using the pyMOL superimpose (super) feature, these two structures were aligned using the common structural elements of hEPO-R and two copies of hCD131. The model 0 structure was used in both cases. This yields a predicted structure of the scFvs in complex with hEPO-R and two copies of hCD131 which we present as a potential model of the entire complex (**Figure 1E**). Confidence metrics of each half of this model are shown in **Supplementary Figure 1**. To ensure biological plausibility of the our model of the complex, our structure was overlaid on the solved structures of each scFv binding its respective receptor (PDB IDs: 4y5y and 5dwu; Lim *et al.*, 2010; Panousis *et al.*, 2016) and examined for alignment. These solved structures aligned well with our predicted complex structure, indicating that AlphaFold 3 had not significantly distorted any of the proteins.

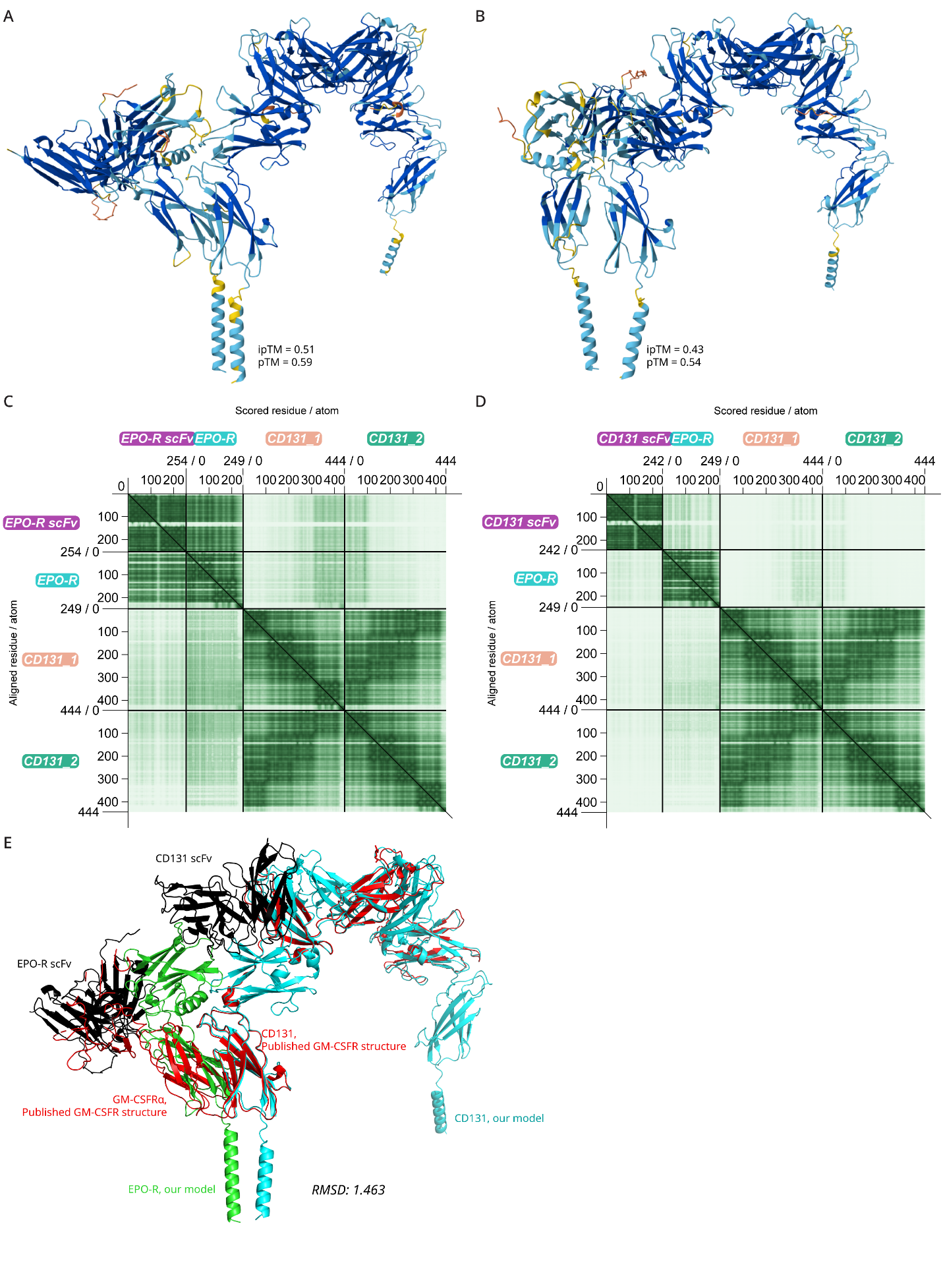

**Supplementary Figure 1.** **Confidence metrics for the proposed model of an EPO-R/CD131 interaction.** Structures of the AlphaFold models that were overlaid in pyMOL to generate the model in **Figure 1E. A.** Predicted structure of the anti-EPO-R/EPO-R/(CD131)_2_ complex. **B.** Predicted structure of the anti-CD131/EPO-R/(CD131)_2_ complex. Chains are colored by AlphaFold pLDDT (per-atom) confidence values. iPTM and PTM values from AlphaFold are also reported. PTM values for both complexes are over 0.5, indicating that the predicted folds in the EPO-R/CD131 complex corresponds to the true structures. **C-D.** AlphaFold PAE matrix from the models in **A.** and **B.**, respectively. The PAE scores corresponding to known interactions are high (dark green). The PAE scores for the putative interaction between EPO-R and CD131 is higher than background for the C-terminal domain of one CD131 subunit and for the N-terminus of the other subunit, corresponding to the predicted interaction, particularly in **A**. **Figure 1E** was generated by superimposing the CD131 elements of these two models in pyMOL, and this model was then used to estimate the linker length needed to attach the two scFvs. This model may or may not be more accurate than the model in **Supplementary Figure** **3F-I. E.** pyMOL structural overlay of the published GM-CSF receptor (GMCSFR) on our structural model from **Figure 1E**. The CD131 elements from both structures align well, and the homologous GM-CSFRα or EPO-R subunits occupy a similar 3D location.

The resulting model of the complex has several features that make it plausible, but also has one region with a steric clash that could be rectified by adjustment of a loop in EPO-R. For purposes of discussion of the hypothetical EPO-R-CD131 interaction, two views of the putative complex along with only the anti-EPO-R scFv are shown (**Supplementary Figure 2**, same complex as shown in **Supplementary Figure 1A**). We also explored several other models in this work prior to generating the model shown in **Figure 1E**. Below, we discuss key features of each of these models (**Supplementary Figure 3**).

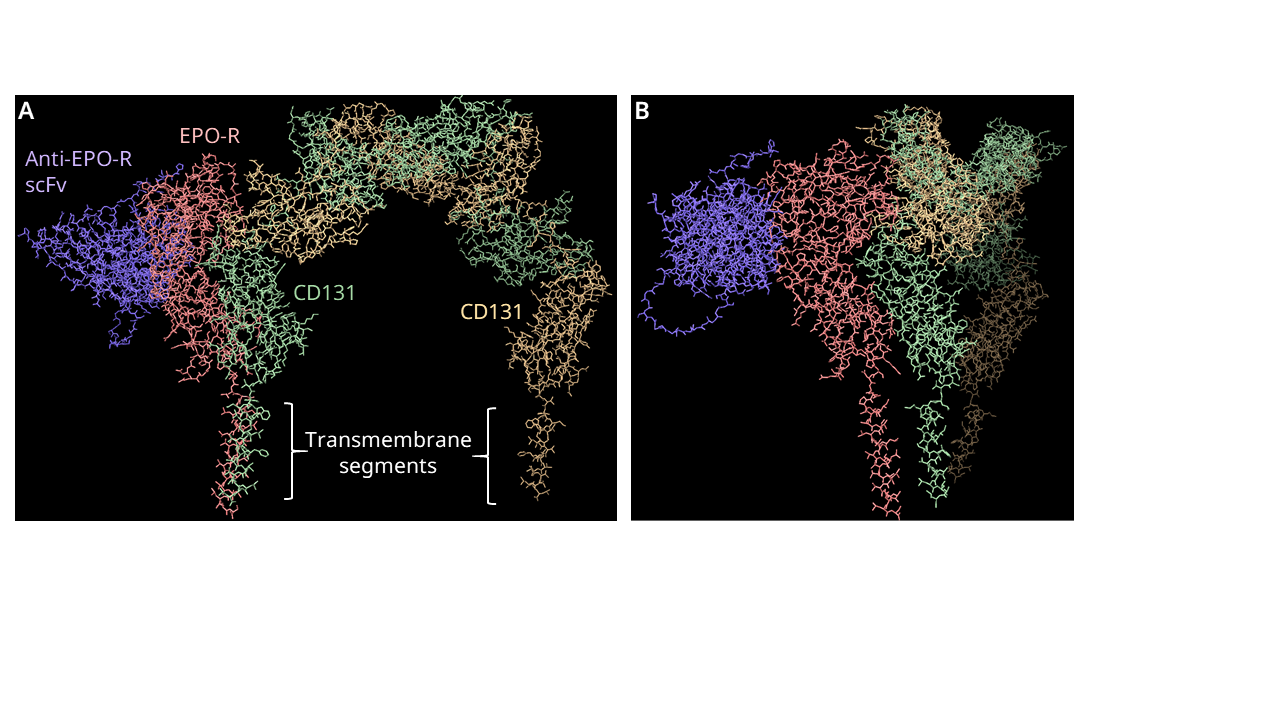

**Supplementary Figure 2.** A hypothetical model of an EPO-R-CD131 interaction generated by AlphaFold 3 (Abramson *et al.*, 2024). The model is built from a solved structure of an EPO-R-binding scFv complexed with EPO-R and the dimeric CD131 found in the 3CXE structure of GM-CSF interacting with its alpha-receptor subunit and CD131. The anti-EPO-R scFv is placed roughly where EPO binds to EPO-R. In the latter structure, CD131 appears to form a stable dimer that forms an arch over the cell membrane and places its transmembrane segments sufficiently far apart that they likely do not signal together. In this model, EPO-R and CD131 interact along a surface of EPO-R that is opposite from the EPO binding site. **A.** View facing CD131. **B.** View facing the EPO-R/CD131 complex, rotated about 75 degrees relative to **A.**

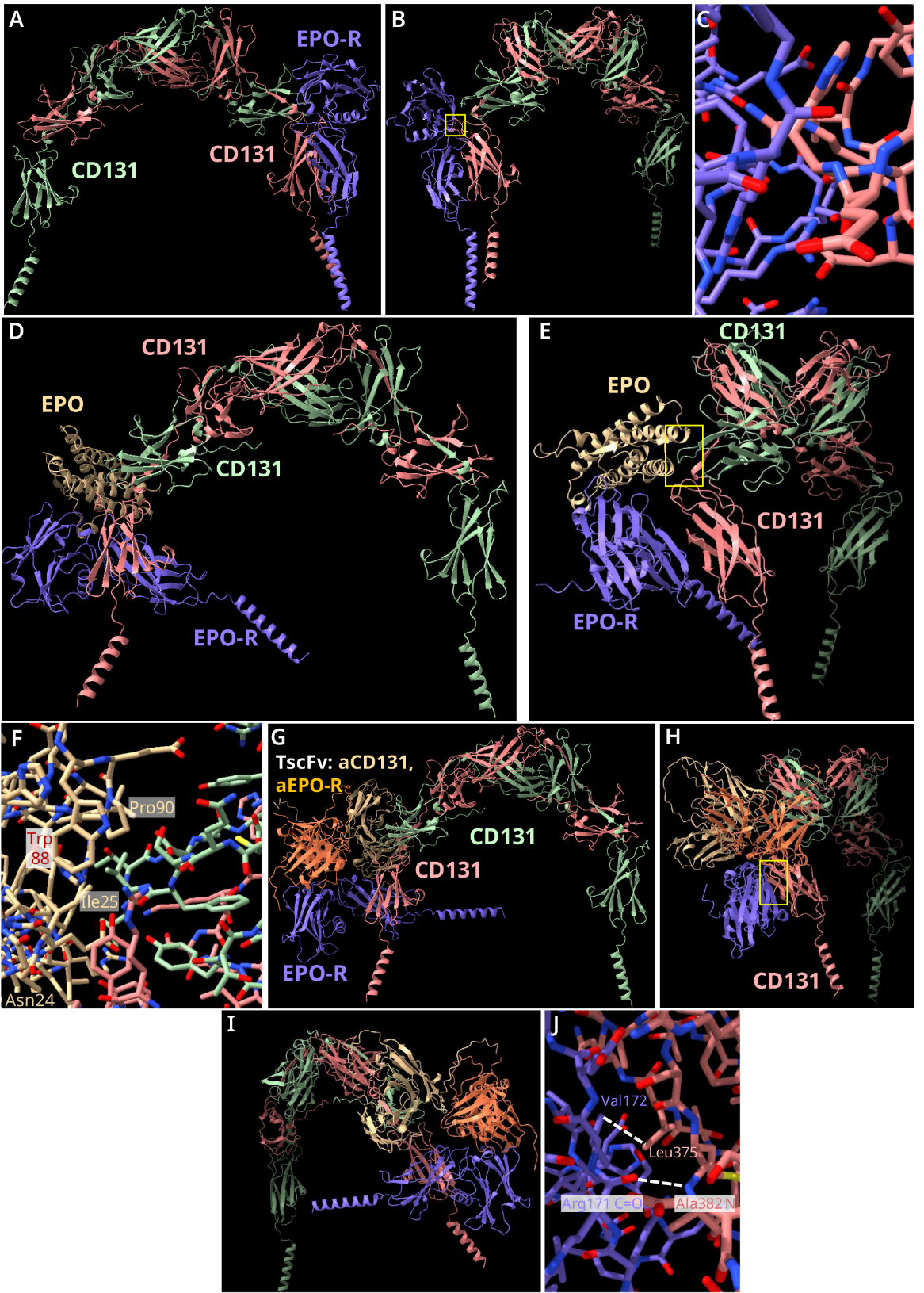

**Supplementary Figure 3. Analysis of related structures predicted by AlphaFold 3.** We also used AlphaFold 3 to predict the following structures of additional complexes: 1. EPO-R, CD131 (two copies), 2. EPO-R, CD131 (two copies), and EPO itself, and 3. EPO-R, CD131 (two copies), and the tandem scFv. 3. Differs from the model in **Figure 1E** as it is an AlphaFold prediction of the entire complex including the full tandem scFv.

**Structure 1: EPO-R with CD131**

In the AlphaFold3-predicted structure of EPO-R with a CD131 dimer and no other molecules (**Supplementary Figure 3A-3C**), EPO-R is associated with one of the CD131 subunits. The transmembrane segments of EPO-R and this subunit are roughly parallel, and the membrane-proximal domains of each protein are within a few Angstroms of each other. However, neither of these interactions show contacts that are close enough to allow for hydrogen bonding or van der Waals interactions. In this predicted structure, there is a strong steric clash between the membrane-distal domain of EPO-R and CD131, with several carbon, nitrogen and oxygen atoms. We therefore consider this model to be obviously incorrect.

To understand why this happens, it is important to recognize that AlphaFold 3 is based on machine learning and not physics. From a distance, this model does look like other transmembrane receptor complexes.

**Structure 2: EPO-R and CD131 with erythropoietin**

In this AlphaFold3 predicted structure (**Supplementary Figure 3C-3E**), EPO interacts with EPO receptor in the same way as seen in the structure 1eer. EPO also interacts with CD131 through residues including Leu17, Glu21, Asn24, Ile25, Trp88, Pro90, and His94. There are no overt steric clashes in this model, and EPO-R and CD131 are not in direct contact. In this structure, the EPO receptor extracellular domain is at an angle, lying almost flat against where the cell membrane would be. This conformation is reminiscent of the 1ERN structure of inactive dimeric EPO-R, and quite different from the 1EER structure of (EPO-R)_2_ with EPO.

The transmembrane segments of EPO-R and CD131 are not parallel and would be about 50 Angstroms apart. However, the fact that the transmembrane segments are not parallel is not an argument against the model overall. Both subunits of CD131 participate in this hypothetical interaction, via residues Ser86, Val88, Val89, and Thr90 on one subunit, and Arg348, Tyr349, His 351, and Tyr405 on the other. We note that EPO residues Tyr88 and Pro90 are not conserved between mouse and human EPO, and that Asn24 is actually N-glycosylated (although this fact was not used in modeling) (Way et al., 2004). In CD131, Ser86, Val88 and Val89 are arginine, serine, and isoleucine in the mouse. The lack of conservation suggests that this model may not be correct.

**Structure 3. EPO-R, CD131, and the tandem scFv**

In this Alpha Fold 3-predicted structure (**Supplementary Figure 3F-3I**), the EPO-R and CD131 interact through the middle of their most membrane-proximal domains. These domains are oriented at about 90^o^ and show extensive contacts including (for EPO-R and CD131, respectively) Arg 171(backbone carbonyl)-Ala382(backbone nitrogen), Val172-Leu375, Leu183-Leu375, Val182(alpha carbon)-Glu376(alpha carbon), Val182(backbone carbonyl-Glu376(backbone nitrogen), Glu173(backbone nitrogen)-Ser380, Leu175-Asn377, Leu175-His379, Arg178-Asp353, and Arg178-Asn357. Cheung Tung Shing *et al.* (Cheung Tung Shing *et al.*, 2018) found no biochemical evidence of interaction between the isolated extracellular domains of EPO-R and CD131 in the presence or absence of EPO, which is consistent with our modeling (**Supplementary Figures 3A-3E**). We hypothesize that if this interaction actually occurs in nature, it is not particularly strong and needs to be stabilized by other interactions, such as those that may occur inside the cell.

In this structure, the scFvs make the same contacts with their receptors as seen in the structures 1EER and 5DWU. If the TscFv is removed from this model and EPO is docked with EPO-R, there are no meaningful contacts between EPO and CD131, but no steric clashes either.

The transmembrane segments of EPO-R and CD131 are not parallel and would be about 40 Angstroms apart, which may reflect that AlphaFold 3 does not recognize the geometric constraints of transmembrane segments.

We consider this model to be plausible but far from proven. The model could be tested by site-directed mutagenesis; for example, removing putative contacts in EPO receptor such that EPO-R homodimeric signaling would be maintained but interaction with CD131 would be lost. More broadly, we found that AlphaFold predictions of the EPO-R/CD131 interaction are generally different from each other, depending on what other molecules are included in the modeling. This fact suggests that if the extracellular domains of EPO-R and CD131 do interact, this interaction may be weak and require stabilization through intracellular contacts between these proteins.

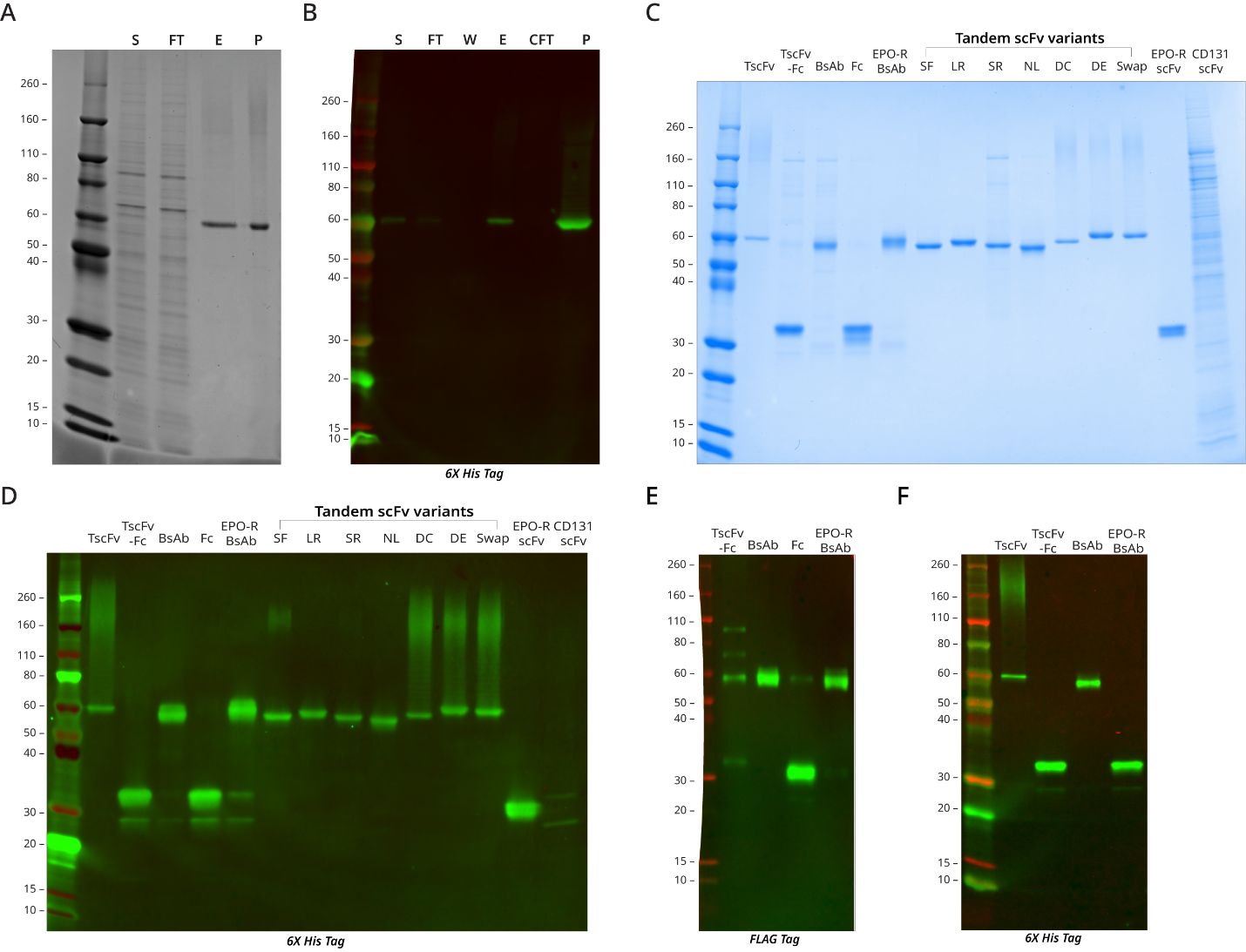

**Supplementary Figure 4. Purification gels of all proteins used in this study.
A.** Representative SDS-PAGE and Coomassie stain or **B.** anti-His tag western blot of TscFv purification. **A-B.** S= cell supernatant, FT= flowthrough, W= wash, E=elution, CFT= concentrator flow through, P=purified (final) protein. **C.** SDS-PAGE and Coomassie stain or **D.** anti-His tag western blot of all expressed proteins. **E. and F.** anti-FLAG tag or anti-His tag western blots demonstrating both halves of the heterodimeric Fc constructs are present after purification. See **Supplementary Table 3** for antibody catalog numbers and dilutions and **Supplementary Table 4** for protein construct name mappings.

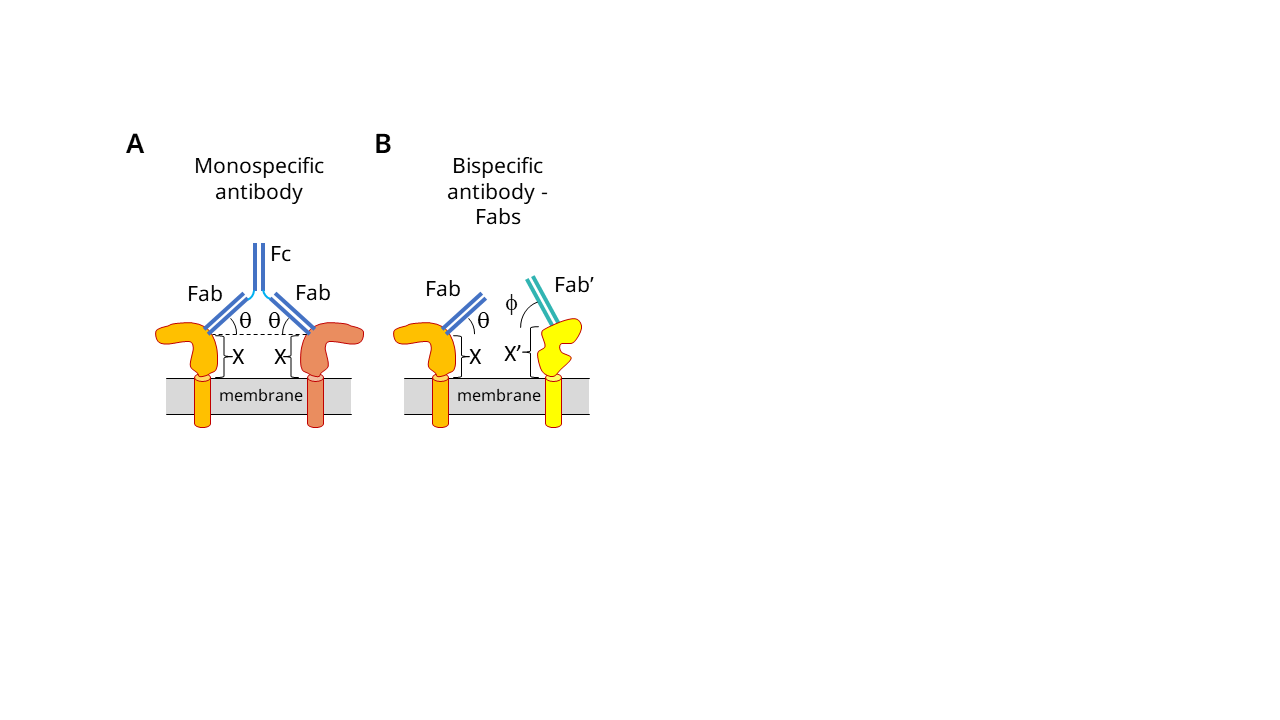

**Supplementary Figure 5. Model for the decrease in activity of the Bispecific Antibody (BsAb, Figure 2).** **A.** Binding of a monospecific antibody (blue) to two identical antigens (tan, light brown) on a cell surface. **B.** Binding of Fabs to two different antigens on a cell surface. The distance of the epitopes from the cell surface on each target and the angle of binding will generally be different, such that the attachment points to the Fc region will be different heights off the membrane. This can lead to decreased or loss of activity.

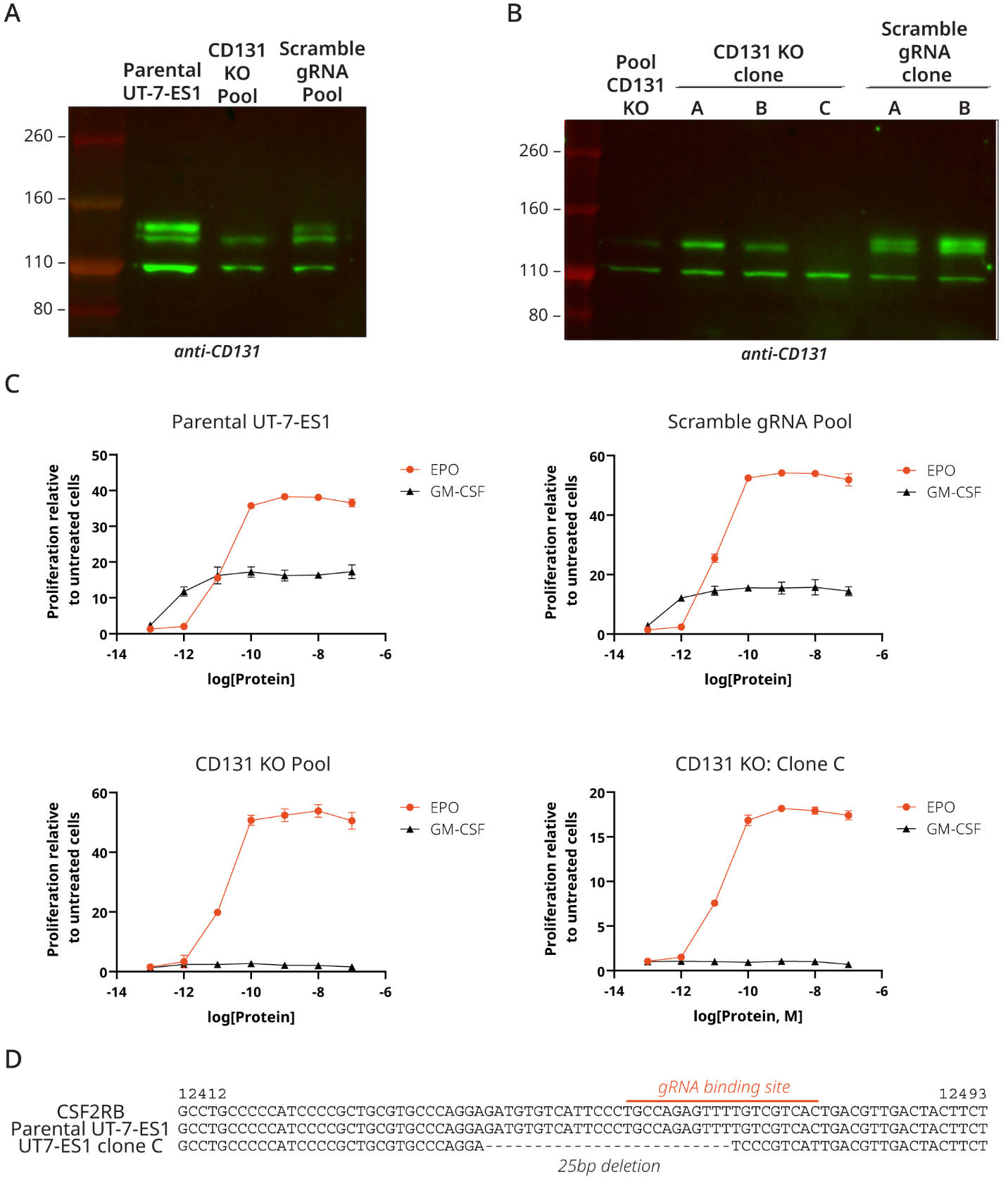

**Supplementary Figure 6. CD131 is functionally knocked out in pooled and clonal UT-7-ES1 CD131 KO cell lines. A.** Parental UT-7 EPO S1 cells (UT-7-ES1), a pool of UT-7-ES1 cells expressing spCas9 and two gRNAs that targeted the CD131 locus, and a pool of UT-7-ES1 expressing spCas9 and two scrambled gRNAs were pelleted and lysed. Then, the lysates were subjected to SDS-PAGE and proteins were visualized by anti-CD131 western blot. One band corresponding to CD131 is knocked out in CRISPR KO pool. **B.** Pooled CD131 KO and scramble gRNA UT-7-ES1 cell lines were used to generate clonal cell lines by limiting dilution. Clonal lines were processed and visualized via anti-CD131 western blot as in **A**. Two bands corresponding to CD131 are knocked out in clone C. **C.** Parental UT-7-ES1 or UT-7-ES1 KO cell lines were cytokine starved overnight, treated with the indicated proteins and viability was measured 72 hrs later. CD131 is functionally knocked out in clone C. CD131 KO Clone C was the cell line used in all main text UT-7-ES1 proliferation experiments.

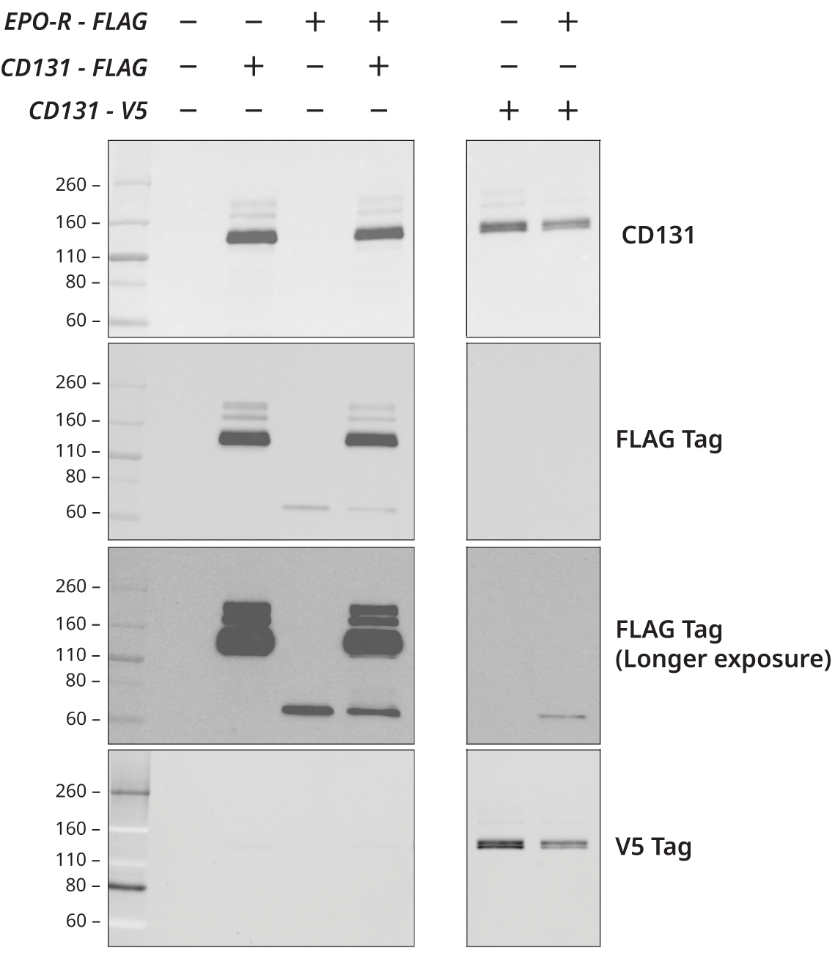

**Supplementary Figure 7. EPO-R and/or CD131 are overexpressed in stably transduced HEK293T cell lines.** HEK293T were transduced with lentiviral overexpression constructs expressing FLAG tagged EPO-R and/or FLAG or v5 tagged CD131. Cells were pelleted, lysed, and the lysates were subjected to SDS-PAGE. Proteins were visualized by western blot with the listed antibodies. Multiple cell lines were established using varied lentiviral titers during transduction to find a pool with the highest expression levels. Titers of the viruses differed and may have led to the observed expression differences between CD131 and EPO-R within and across the cell lines. See **Supplementary Table 5** for specific information on which cell lines were used in which main text figures.

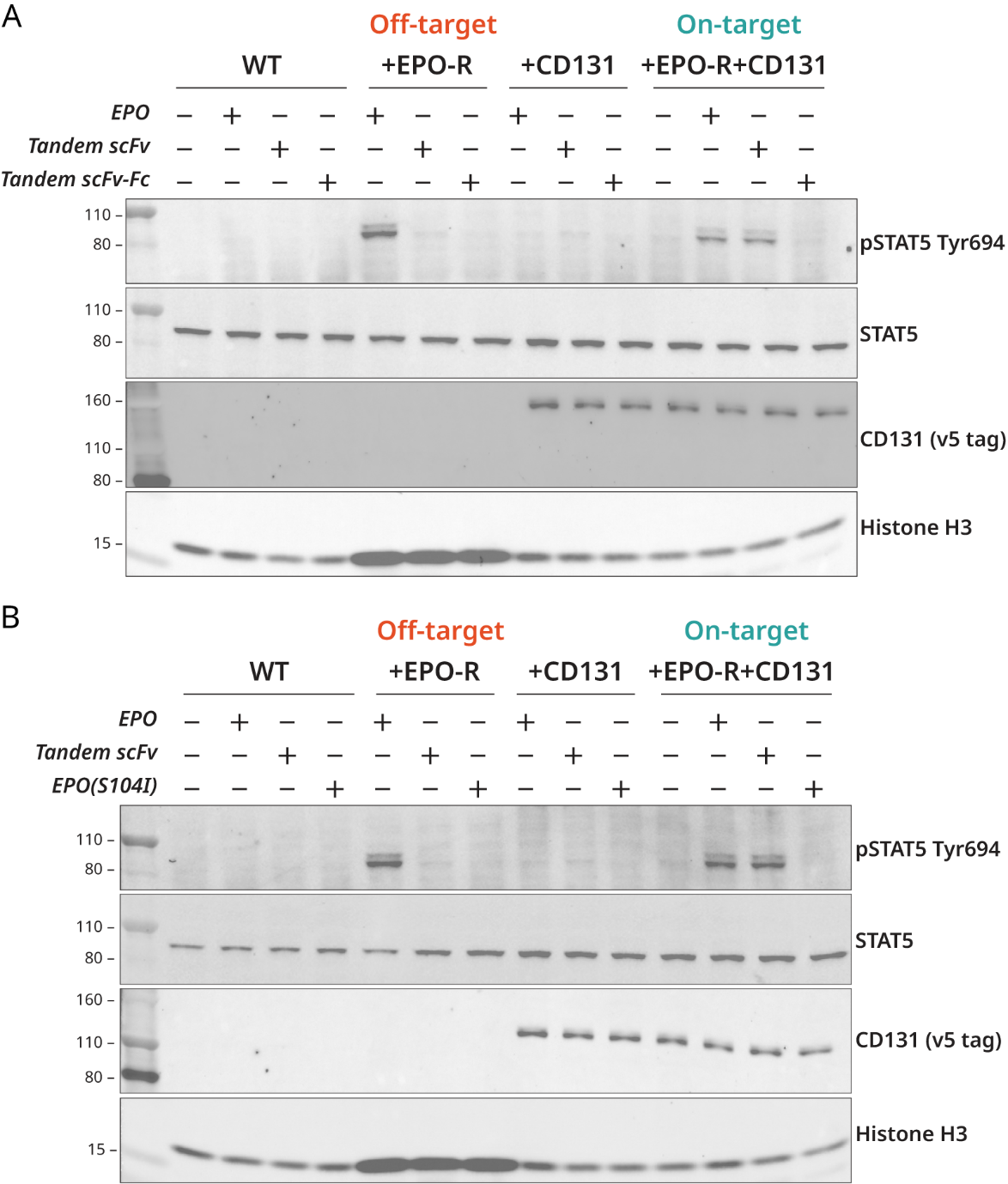

**Supplementary Figure 8. Tandem scFv does not induce STAT5 phosphorylation in a CD131-expressing cell line with low CD131 expression.** A HEK293T CD131-v5 tag cell line was established in addition to the HEK293T CD131-FLAG tag line from **Figure 3B, 3C**. The CD131-v5 tag cell line showed lower CD131 expression than the CD131-FLAG tag line, based on a western blot with an anti-CD131 antibody (**Supplementary Figure 7**). HEK293T overexpression cell lines were pelleted, lysed, the lysates were subjected to SDS-PAGE and proteins were visualized western blot with the indicated antibodies. **A.** and **B.** represent independent experiments. Unlike in the FLAG-tagged CD131 overexpression line (**Figure 3B, 3C**), TscFv did not induce STAT5 phosphorylation in the v5-tagged CD131 overexpression line. CD131 expression of each cell line is shown in **Supplementary
Figure 7** (third lane (CD131-FLAG) and penultimate lane (CD131-v5)). The mechanism by which TscFv induces some STAT5 phosphorylation in one CD131-expressing, EPO-R- null cell line and not the other is unclear.

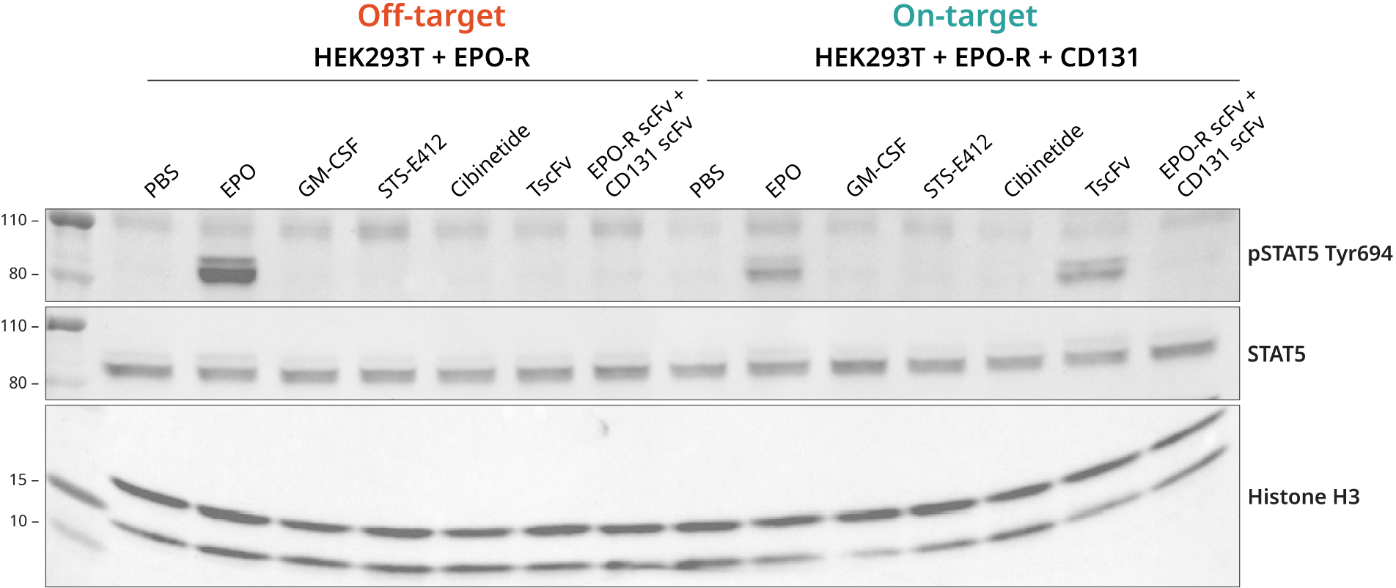

**Supplementary Figure 9. An equimolar mixture of the EPO-R and CD131 scFvs does not induce STAT5 phosphorylation in cells expressing only EPO-R or both EPO-R and CD131.** HEK293T overexpression cell lines were treated with 100nM of the specified molecules for 15 minutes at 37°C. The EPO-R scFv + CD131 scFv sample was an equimolar mix of each protein (100nM final concentration for both). Cells were then pelleted, lysed, the lysates were subjected to SDS-PAGE and proteins were visualized via western blot with the indicated antibodies. No STAT5 phosphorylation is induced by the equimolar mixture, suggesting STAT5 phosphorylation is dependent on the single chain, bispecific format of TscFv. Additionally, STS-E412 and Cibinetide, two literature-reported EPO-R/CD131 agonists, do not agonize EPO-R/CD131 in our assay.

**Discussion of how TscFv with shorter linkers could still induce signaling**

Short-linker variants of the TscFv were constructed to test the structural model proposed in **Figure 1E**, which predicted that a linker of at least 22 amino acids between the two scFv domains would be required for binding of each scFv to its receptor. As observed in **Figure 5**, short-linker variants are less potent than the 23-amino acid linker version of the TscFv in inducing EPO-R/CD131 heterodimeric signaling, but this signaling is not eliminated. Residual signaling could come about if the short-linker TscFv constructs can bind across two signaling complexes, as depicted below (**Supplementary Figure 10**). Alternatively, it may be sufficient to bring EPO-R and CD131 into general proximity, without any interaction between the extracellular domains. Formation of a signaling complex could be accommodated by flexibility at the junction between the extracellular domains and the transmembrane segments.

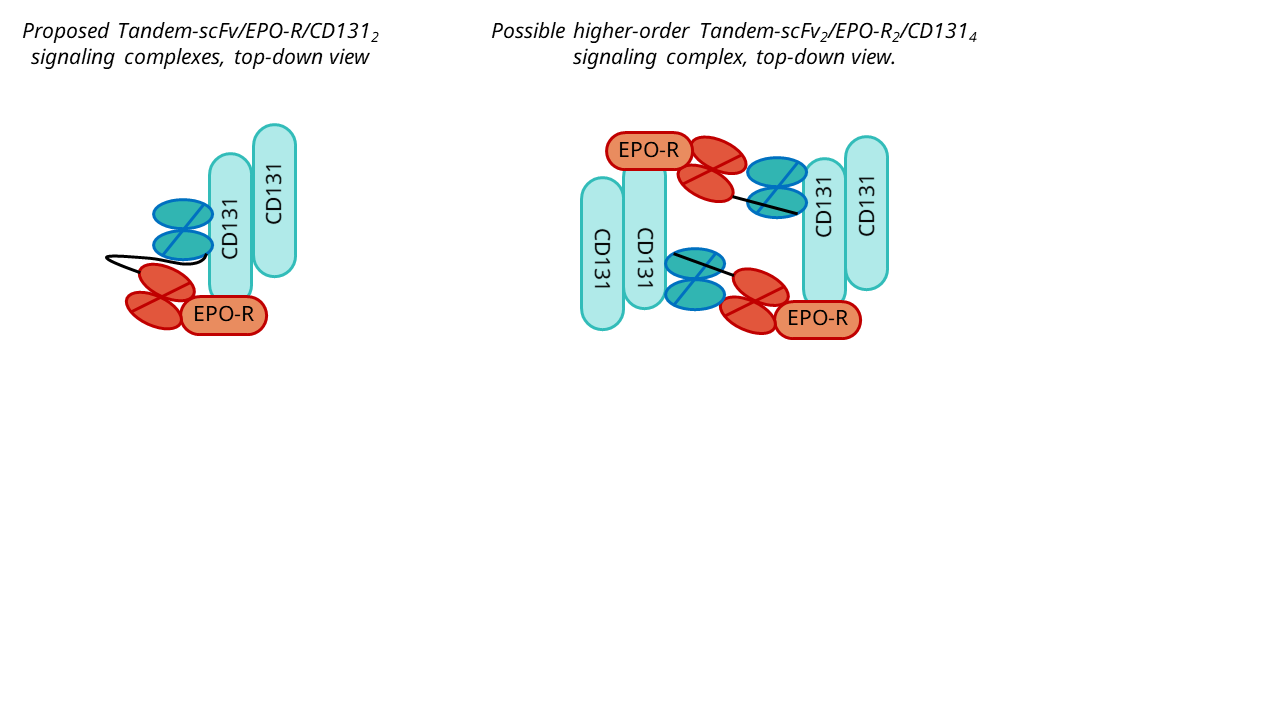

**Supplementary Figure 10. Model for residual activity of Tandem-scFv constructs with short linkers.** Our linker variant data demonstrates that tandem scFvs with linkers less than 88 angstroms still signal through EPO-R/CD131. This may be due to the formation of higher order structures (Tandem scFv_2_/EPO-R_2_/CD131_4_). In this complex, two tandem scFvs could be closer in space than 88 angstroms and the linker length may still be able to span the distance between receptors. **Left.** Diagrammatic representation of a Tandem scFv binding to EPO-R and CD131, with the inter-scFv linker shown in black. **Right.** A dimeric version of the proposed signaling complex could overcome the geometric constraint imposed by a short inter-scFv linker.

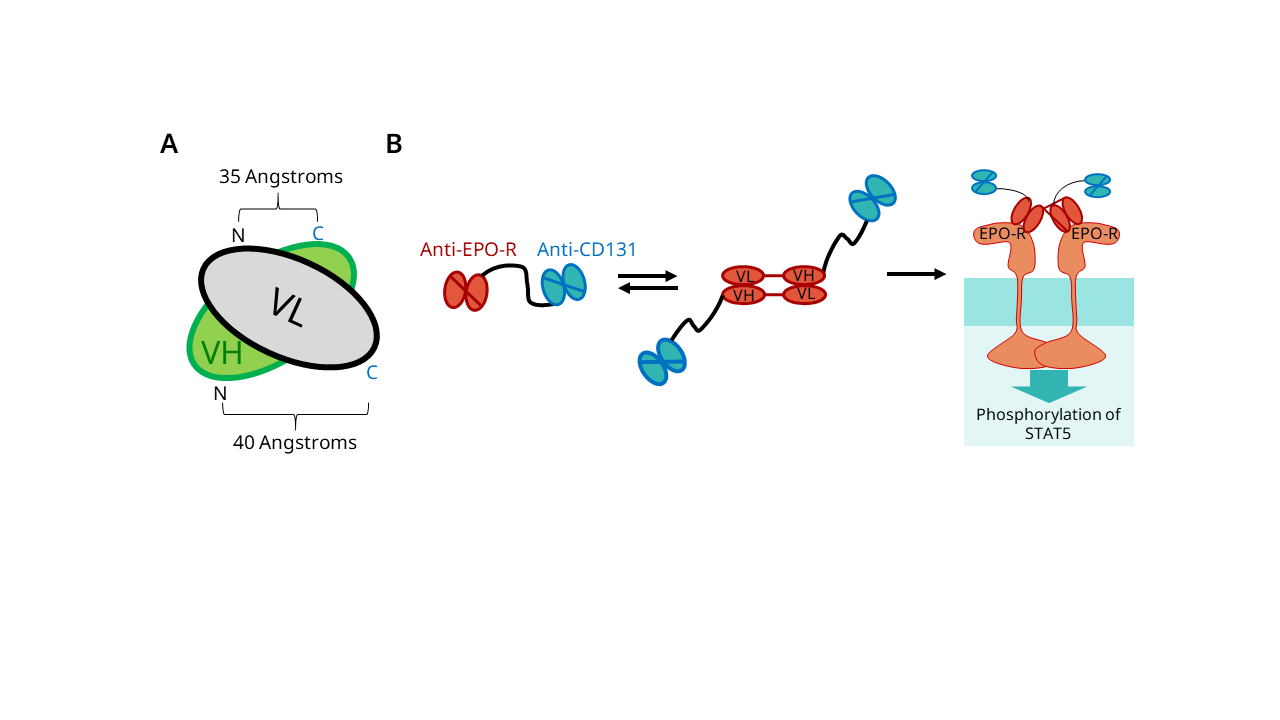

**Supplementary Figure 11. Model for why the “Swap” construct induces STAT5 phosphorylation in UT-7(EPO-S1) CD131-KO cells while the Tandem scFv does not.** **A.** Diagram of a typical scFv, showing the geometry of the N- and C-termini of the VH and VL domains. The VH and VL domains are approximately rotationally symmetric around an axis running between them, but the orientations differ in detail. The distance between the VH C-terminus and VL N-terminus is about 35 Angstroms and relatively un-occluded, while the distance from the VL C-terminus and the VH N-terminus is about 40 Angstroms, and a straight line between these termini runs through the VL. Thus, to properly attach the termini in the VL-VH configuration requires a longer linker than in the VH-VL configuration. i.e., a linker of more than 17 amino acids may be required to ensure that an scFv in the VL-VH orientation is entirely in a monomeric state. **B.** Proposed monomer-dimer equilibrium of the Swap configuration and signaling via EPO receptor homodimers. We hypothesize that the short linker in the anti-EPO-R VH-VL scFv promotes the formation of a dimeric state, which can then cross-link EPO-R and induce STAT5 phosphorylation. The dimerized form of anti-EPO-R V regions used in these experiments are known to induce EPO-R signaling (Lim *et al.*, 2010; Moraga *et al.*, 2015). We note that while the Swap construct induces STAT5 phosphorylation in UT-7(EPO-S1) CD131-KO cells, the stimulation is not strong enough to induce proliferation of these cells. This may indicate that the signaling process induced by SWAP is somehow defective.

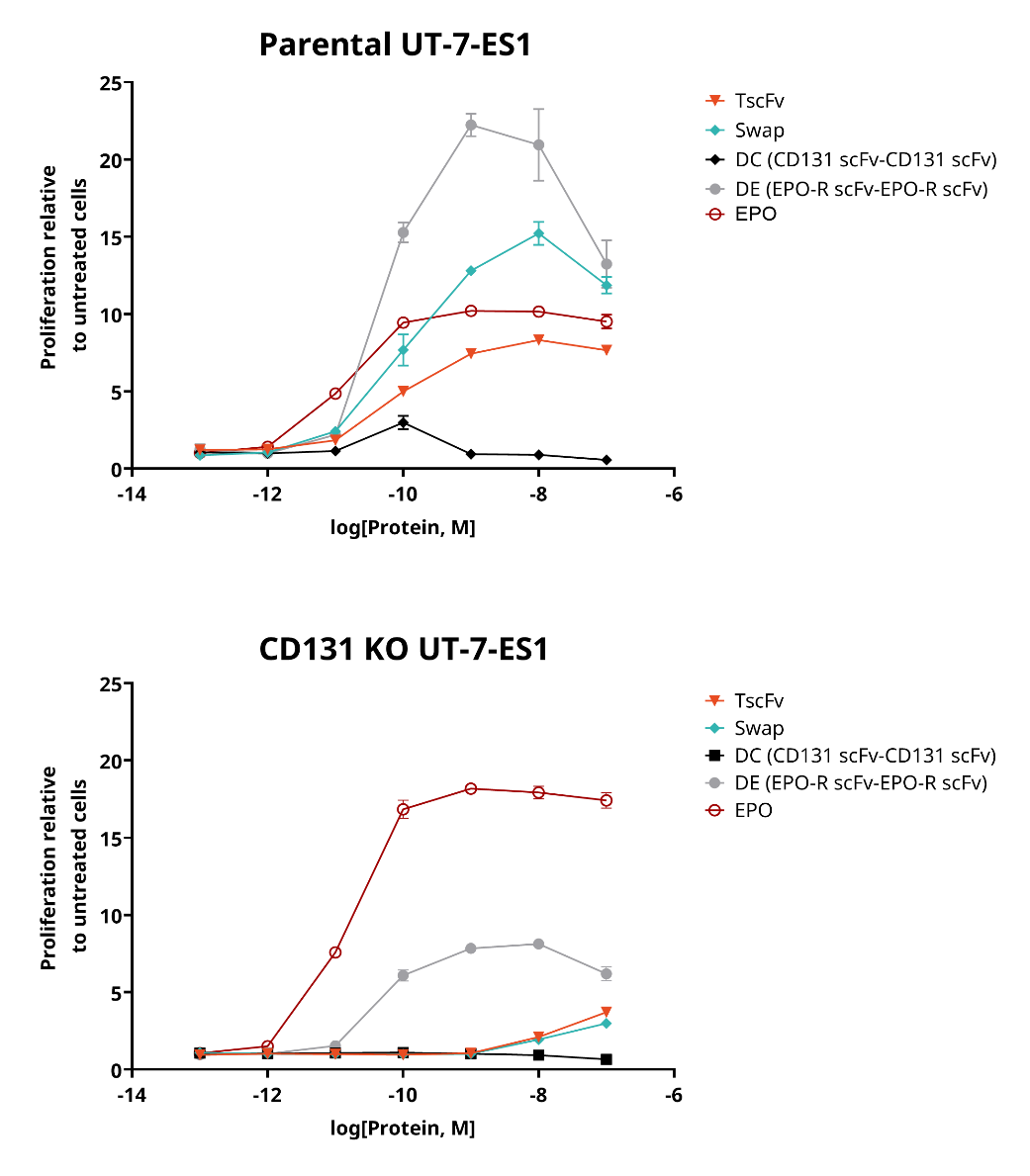

**Supplementary Figure 12. EPO and an anti-EPO-R – anti-EPOR-R tandem scFv (DE), but not TscFv, are able to support the growth of UT-7-ES1 CD131 KO. A.** Parental UT-7-ES1 or a UT-7-ES1 KO cell line were cytokine starved overnight, and treated with the indicated proteins: TscFv (antiEPO-R-antiCD131), “Swap” (antiEPO-R-antiCD131 in which the order of the anti-CD131 VH and VL domains was reversed without a compensating increase in the length of the intervening linker); DC (double-antiCD131, consisting of tandem antiCD131 scFvs); DE (double-antiEPO-R, consisting of tandem antiEPO-R scFvs); and erythropoietin itself. Viability was measured 72 hrs later. The EPO-R/CD131 tandem scFv (TscFv) and Swap are unable to support UT-7-ES1 proliferation in the absence of CD131, but DE and EPO are able to support proliferation in the absence of CD131.

| 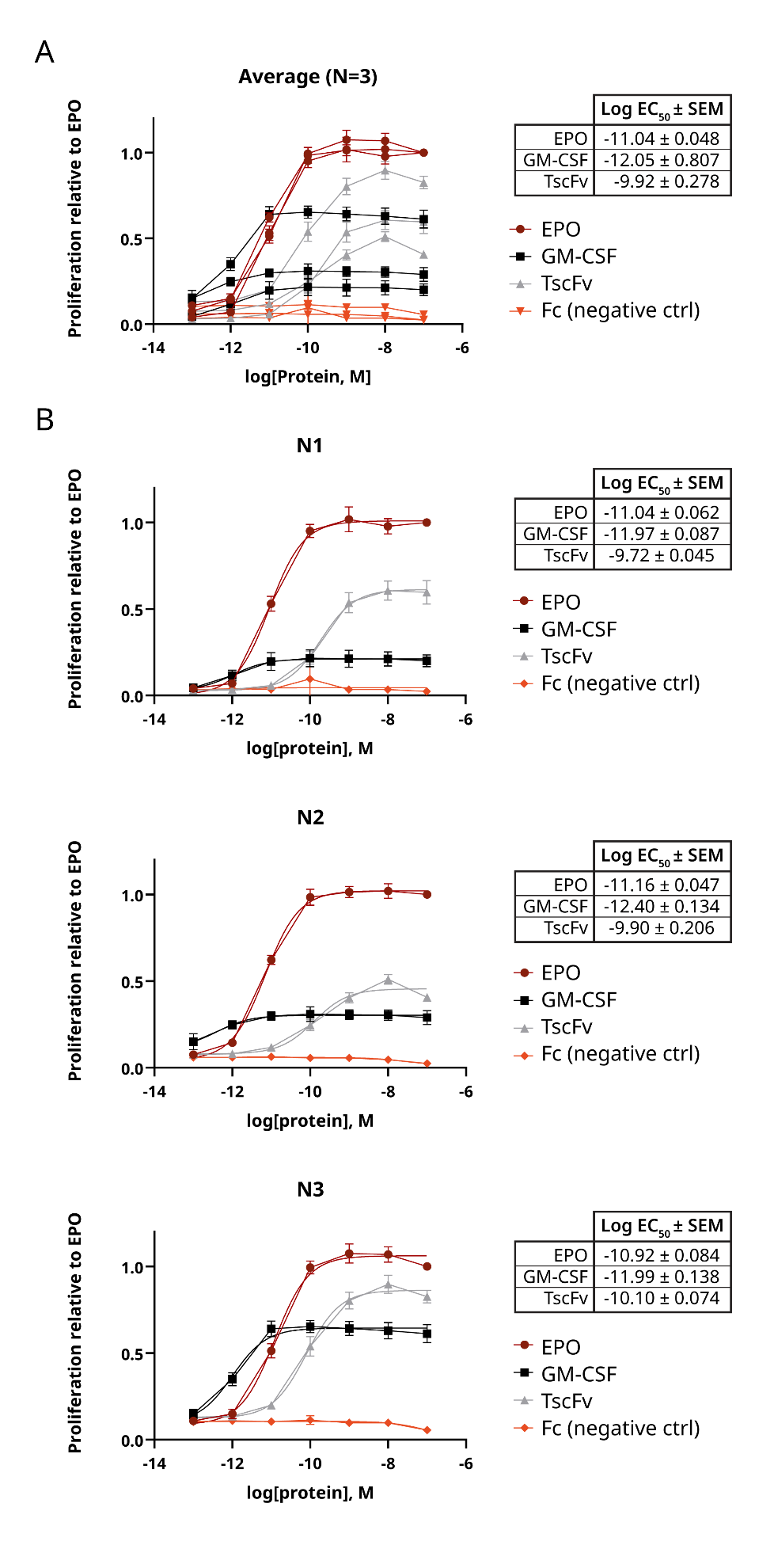 | **Supplementary**  **Figure 13. Proliferative response profiles to proteins in this study replicate across experiments.** Parental UT-7-ES1 were cytokine starved overnight, treated with the indicated proteins and viability was measured 72 hrs later. **A.** Aggregate of N=3 independent experiments shown in **B.** performed on separate days and cell aliquots. Data are reported as proliferation relative to EPO treated cells at the same dose and are represented by mean luminescence ± SD of three replicates. Log EC_50_s were calculated using GraphPad Prism using nonlinear regression (log(agonist) vs. response, variable slope (four parameters). E_max_ (Max proliferation relative to EPO samples) may change depending on experiment, but Log EC_50_s remain similar. |
| --- | --- |

### **Extended methods**

#### **Plasmid construct generation**

A PCR amplicon of the vector backbone (either pSecTag2a or pOptivec) or inserts of interest were amplified using Q5 High Fidelity 2X Master Mix (NEB M0492S) diluted to 1X, 0.5 µM each of forward and reverse primers, 4 ng of plasmid and water to 25 µL. Oligonucleotide primers were ordered from IDT which were checked for thermodynamic parameters using IDT OligoAnalyzer. PCR was run on a thermal cycler (BioRad C1000) with the following parameters: 30s of 98°C denaturation, 30 cycles of 30s denaturation at 98°C, annealing at the primer annealing temperature as calculated by the NEB Tm calculator for 30s, extension at 72°C for 30-50 s/kb, and final extension at 72°C for 2 minutes. PCR products were run in 1x sample buffer (NEB B7024S) on a 1% agarose gel (Lonza 50004 dissolved in 1X TAE buffer) with a ladder (Thermo SM1333) for analysis. Following the confirmation of a single band, products were treated with Dpn1 (NEB R0176S) following the manufacturer’s instructions, PCR cleaned (Promega A9281) and DNA concentration was determined using NanoDrop One (Thermo ND-ONE-W). Inserts were prepared by either PCR amplification from previously cloned vectors as above or by annealing self-complementary oligonucleotides (IDT) for insertions for shorter insertions. Self-complementary oligonucleotides were resuspended in Duplex buffer (IDT 11-05-01-03), combined in an equimolar ratio, heated to 94°C for 2 minutes and cooled gradually at room temperature to anneal. Assembly was performed according to a miniaturized version of the manufacturer’s protocol where each reaction had a total volume of 5 µL and between 0.01 - 0.25 pmol of total DNA for 1-2 Fragments, and 0.1 - 0.5 pmol of total DNA for 3+ fragments. For a 1-2 insert assembly, the insert:vector ratio of pmol should be 2-3:1, for 3+ inserts the ratio should be 1:1, and for small fragments (< 200 bp) the ratio should be 5:1. Assembly reactions were incubated at 50°C for 1 hr, and 1 µL of each reaction was transformed into NEB 5-alpha E. coli (NEB C2987H) following the manufacturer’s protocol. 50 µL of transformation mix was plated on ampicillin agar plates (100 µg/mL) and incubated overnight at 37°C. Individual colonies were picked the following day into 10 µL nuclease free water. 5 µL was sent for colony sequencing (Plasmidsaurus) and 5 µL was used to start 150 mL overnight cultures. Cultures were maxiprepped with an endotoxin-free maxi kit (Qiagen 12963), and DNA concentration was taken by NanoDrop. Occasionally purified DNA was sent for sequencing rather than colonies.

In some cases, plasmids were modified using KLD enzyme mix (NEB M0554S) to make small insertions, deletions and substitutions, such as changing protein tags, following the manufacturer’s instructions rather than isothermal assembly.

All constructs used for protein expression are in the pSecTag2a plasmid backbone obtained from Dr. Timothy Chang (Harvard Medical School) or the pOptivec plasmid backbone with added secretion signal from the V-J2-C region of the mouse Ig kappa-chain (sequence from pSecTag2a plasmid) for secreted protein expression. All secreted proteins were His6-tagged to enable purification. Notably, the Fc fusion proteins were His6-tagged only on one half of Fc to favor purification of heterodimers.

hEPO-R and hCD131 receptor overexpression constructs and tandem scFv linker variant constructs were ordered as clonal genes from Twist using the codon-optimized sequences as detailed above. Overexpression constructs (plasmids KD42, KD53 and KD66) were ordered in the pHAGE-CMV puro second-generation lentiviral backbone (a gift from Dr. Rui Tong Quek). The plasmid encoding hEPO-R FLAG blasticidin (plasmid KD53), was cloned into the pHAGE-CMV puro second-generation lentiviral backbone (a gift from Dr. Rui Tong Quek), and the puromycin resistance cassette was swapped for *ble* (blasticidin resistance) using isothermal assembly as described below. The hCD131 HA, v5 tag construct (KD66) was cloned using isothermal assembly of self-complimentary oligonucleotides encoding the HA and v5 tags with overlaps to replace the FLAG tag in KD42. CSF2RB g1 and CSF2RB g2 plasmids (plasmids KD51 and 52 respectively) were cloned using isothermal assembly of oligonucleotides encoding CSF2RB-targeting gRNA spacers Custom Alt-R™ CRISPR-Cas9 guide RNA tool from IDT into a third generation lentiviral gRNA plasmid (obtained from Dr. Rui Tong Quek, originally a gift from John Doench & David Root (Addgene plasmid # 76370 ; http://n2t.net/addgene:76370 ; RRID:Addgene_76370)(Doench *et al.*, 2016). Oligonucleotides were designed so that the CSF2RB spacers replaced the existing GSK3A spacer. Tandem scFv linker and domain variant plasmids (plasmids KD67 – KD80) were ordered as clonal genes in pSecTag2a from Twist.

#### **Cell culture**

HEK293T cell lines were passaged every 3-4 days by washing the cell monolayer once with DPBS (Thermo 14190144), lifting by treatment with trypLE (Thermo 12605010) for 3-5 minutes, neutralizing with complete growth medium and directly passaged into new flasks at a 1/10 – 1/20 dilution.

All cell lines were maintained at 37°C and 5% CO_2_ in a humidified incubator. All complete media were prepared by combining ingredients and then filtering by 0.2 µm filter (Corning CLS431153). The exception to this is FreeStyle 293 Expression Medium which comes ready-to-use and was not filtered. All cell lines were cryopreserved by resuspending cells to 10 million cells/aliquot in 90-95% complete medium supplemented with 5-10% (v/v) cell-culture grade DMSO (Sigma D2650-100ML). Cells were frozen in a Mr. Frosty Freezing Container (Thermo 5100-0001) overnight at -80°C. Vials were then moved to liquid nitrogen for long term storage the next day. All cell lines were thawed by incubating cryovials at 42°C in a bead bath until all ice melted then moved directly into pre-warmed media in a new flask. A full list of cell lines generated in this work and the corresponding figures they are used in is located in **Supplementary Table 5**.

#### **Lentiviral generation**

HEK293T (Low passage P<10, viability ≥90%) were plated so they would reach 60-80% confluence the following day in complete medium in T75 flasks. The next day, 12 µg total divided amongst each overexpression construct in the pHAGE lentiviral plasmid backbone was co-transfected with 9 µg of psPAX2 (a gift from Didier Trono, Addgene plasmid # 12260 ; http://n2t.net/addgene:12260 ; RRID:Addgene_12260) and 3 µg of pMD2.G (a gift from Didier Trono, Addgene plasmid # 12259 ; http://n2t.net/addgene:12259 ; RRID:Addgene_12259) packaging plasmids with Lipofectamine 3000 (Thermo L3000015) following the manufacturer’s protocol. After 18-24 hr, transfection medium was aspirated and replaced with complete medium. Culture medium was collected after 24 and 48 hrs and combined, and cells were pelleted by centrifugation at 300xg for 5 minutes. Virus-containing supernatant was filtered through a 0.45 µm Durapore PVDF membrane steriflip (Millipore SE1M003M00). Lenti-X Concentrator (TaKaRa 631232) was then diluted to 1x in the filtered virus-containing medium, the tube was gently mixed and then incubated at 4°C for at least 1h (but as long as 3 days). After incubation, tubes were centrifuged for 45 min at 1,500xg to pellet virus. Supernatant was removed carefully, and the viral pellet was resuspended in complete medium without antibiotics in 1/10th the original volume of harvested supernatant. Virus was aliquoted and frozen at -80°C until use or used right away.

#### **Lentiviral Transduction**

All stable cell lines generated in this study are listed in **Supplementary Table 5**. One day before transduction, 500,000 low passage (P<10) HEK293T cells per well (viability ≥90%) were plated in 6 well plates. Transductions were performed the following day by replacing media with 0.4 mL of 10X polybrene (Millipore TR1003-G), varying volumes of virus (0.5 – 2mL) and complete medium without antibiotics to 4 mL total. Mock wells were transduced with 3.6 mL of medium and 0.4 mL of polybrene only to monitor toxicity of the transduction mix. Multiple viral volumes were tried in each experiment since differing viral loads can lead to differences in survival and/or expression levels.

Cells were transduced, incubated for 24-48 hrs, virus-containing media was aspirated, and selection medium was added. Puromycin selection was allowed to proceed for at least 4 days while blasticidin selection proceeded for 14 days before cryopreservation of cell aliquots and collection of samples to analyze transgene expression via SDS-PAGE and western blot. HEK293T overexpression cell lines used in this study were chosen based on highest transgene expression level. Several versions of each HEK293T + hEPO-R, HEK293T + hCD131 and HEK293T + hEPO-R + hCD131 were established from differing viral titer during transduction and the tags on the overexpression construct (See table below).

UT7 EPO S1 (UT-7-ES1) knockout lines were generated by lentiviral transduction as described above. 4 µg of lenti-Cas9 blasticidin plasmid obtained from Dr. Rui Tong Quek (originally from Feng Zhang (Addgene plasmid # 52962 ; http://n2t.net/addgene:52962 ; RRID:Addgene_52962)(Sanjana *et al.*, 2014) and 4 µg each of KD51 and KD52 plasmids were co-transfected into HEK293T as described above for virus production. For control line generation, 4 µg of the lenti-Cas9 blasticidin plasmid and 4 µg each of 2 non-targeting scramble gRNA plasmids (non-targeting control gRNA (BRDN0001149198), Addgene plasmid # 80248 ; http://n2t.net/addgene:80248 ; RRID:Addgene_80248) and non-targeting control gRNA (BRDN0001148862) Addgene plasmid # 80263 ; http://n2t.net/addgene:80263 ; RRID:Addgene_80263 which were a gift from John Doench & David Root) (Doench *et al.*, 2016) were co-transfected. UT-7-ES1 pooled CD131 KO and pooled scramble gRNA lines were selected for 14 days before cryopreservation of cell aliquots and collection of samples to analyze the KO via SDS-PAGE and western blot for human CD131 (CST 3432S, 1/1000). Parental UT-7-ES1 stain for 3 CD131 bands, CD131 KO pooled line stains for 2; however, pooled line no longer respond to GM-CSF for growth, which is indicative of a functional KO of CD131. Pooled lines were then subcloned by limiting dilution into 96 well plates to generate ~1 cell per well. Wells with single cells were grown until the well was 60 – 80% confluent, and then each line was moved to T25 flasks for expansion and eventual cryopreservation. Samples were taken and analyzed via SDS-PAGE and western blot for human CD131.

#### **SDS-PAGE**

4X Laemmli SDS sample buffer (Thermo J60015.AD) was diluted with the protein samples or cell lysate and diH_2_O water to 1X. Samples were boiled at 95°C for 5 minutes to denature proteins. 4 to 20% Tris-Glycine gradient gels (Thermo XP04205BOX) were loaded into the XCell *SureLock*™ Mini-Cell (Invitrogen EI0001) gel electrophoresis system and 1X Tris-glycine running buffer (Thermo LC2675) was added to the chamber. Combs were removed and 10-20 µL of sample (between 100 ng and 10 µg total protein depending on application) were run alongside 5-10 µL of Novex™ Sharp Pre-stained Protein Standards (Thermo LC5800) for 1 hr at 200V or until dye front reached the bottom of the gel. Gels were carefully removed from the cassette using a gel knife (Thermo EI9010), rinsed once with diH_2_O in a dish and then stained using SimplyBlue™ SafeStain (Thermo LC6065) and destained with diH_2_O following the manufacturer's protocol. Stained gels were imaged with a ChemiDoc MP (Bio-rad 12003154) using the Coomassie setting.

#### **Western Blot**

Gels were carefully removed from the cassette using a gel knife (Thermo EI9010), rinsed once with diH_2_O in a dish and transferred to a nitrocellulose membrane (Thermo IB23002) using the iBlot 2 gel transfer device (Invitrogen IB21001). The transfer stack was set up according to the manufacturer’s protocol. Care was taken to not touch the membrane with anything but clean tweezers and to roll air bubbles out gently with a roller (Thermo 84747) between each layer of the stack. Transfer was run using the default P0 transfer protocol (20 V for 1 minute, 23 V for 4 minutes, 25 V for 2 minutes). The blot was transferred to a western blot box (Southern Labware B1200-7BK or B1300-8BK) and washed twice while shaking in TBS (Thermo J60877.K3) + 0.05% Tween-20 (Millipore Sigma P9416-50ML). Blots were blocked for 1 hr at room temperature or overnight at 4°C while shaking in TBST + 1% BSA (GoldBio A-420-10). Primary antibody was then added directly to the blocking solution and incubated shaking for 1 hr at room temperature or overnight at 4°C. Blots were washed three times in TBST to remove unbound primary antibody and either directly developed for conjugated primary antibodies or stained with secondary antibodies. Secondary antibodies were diluted in TBST + 1% BSA, added to the blot box, and incubated for 1 hr at room temperature or overnight at 4°C shaking. Blots were washed three times more in TBST to remove unbound secondary antibody. Blots were developed using SuperSignal West Dura Extended Duration Substrate (Thermo Scientific 34075) by either rolling 200-500 µL of substrate over the blot on the imaging tray or by quickly dunking the blot in substrate then transferring it to the imaging tray. Blots were imaged using the chemi setting on a ChemiDoc MP. All steps for fluorescent conjugated antibodies were done in covered boxes to prevent photobleaching. All antibodies and dilutions used are listed in **Supplementary Table 3.**

Bands were background corrected individually by the ImageLab software (local background correction feature), and background-corrected intensities were normalized to the loading control from the same well to account for variations in sample loading.

$Normalized band intensity= \frac{intensity of background-corrected sample band}{intensity of background-corrected loading control band}$

The corrected, normalized phospho-STAT5 signals were then normalized to the corresponding total STAT5 signal to account for differences in protein expression levels across samples.

$$STAT5 phosphorylation= \frac{intensity of normalized phosphorylated STAT5 band}{intensity of normalized total STAT5 band}$$

Finally, all normalized values were expressed relative to the samples treated with the negative control (PBS), which was set to 1. More detail about the immunostaining protocol and full gel or blot images and analyses are provided in the Supporting Information. Antibody catalog numbers and dilutions are listed in **Supplementary Table 3**.

#### **Protein expression**

Proteins were expressed transiently in HEK293-F cells following the manufacturer’s protocol. 24 hrs prior to transfection, low passage (P<10) HEK293-Fs (viability ≥90%) were seeded in sterile, vented cap Erlenmeyer cell culture shaker flasks (Corning CLS431143-50EA or CLS431145-25EA) at 600,000 cells/mL in Freestyle 293 Expression medium (Thermo 12338018) in no more than 1/3 of the total flask volume overnight. 1 µg of plasmid DNA/mL of culture of each plasmid (if using two plasmids, DNA was split evenly between them) was added to Opti-MEM (Thermo 31985062). 2 µL of 293-fectin (Thermo 12347019) per 1 µg of DNA was added to Opti-MEM in a 15 mL falcon tube and incubated for 5 minutes at RT. DNA:optimem and 293fectin:optimem mixtures were combined and incubated for 20-30 minutes at RT. Mixture was added to cells while gently swirling. Flasks were incubated for 4-6 days at 37°C, 5% CO_2_ and shaking at 130 rpm to allow expression and secretion of proteins into the medium. Only endotoxin-free, sterile reagents were used. Cells were pelleted and the cell supernatant containing the expressed proteins was isolated.

#### **His-tag purification**

His-tagged proteins were purified from the cleared cell supernatant using His60 Ni Superflow Resin and buffers (Takara 635677). 1-3 mL of resin was centrifuged at 100xg for 2 minutes and supernatant was removed. Resin was washed two times in 10mL of equilibration buffer, spun at 100xg for 2 minutes and the supernatant was removed. Equilibrated resin was moved into 50mL falcon tubes with cell supernatant containing the expressed proteins and incubated rotating or for at least 2 hrs at 4°C. If supernatant volume was >100mL, supernatant and resin was incubated in 500mL conical flasks (Corning CLS431145-25EA) and shaken at 4°C overnight. Supernatant:resin slurry was moved to a column (Thermo 89898) and allowed to run through into a 50mL falcon collection tube. The column was washed with 10mL of equilibration buffer, 10 mL of wash buffer and proteins were eluted in 15mL of elution buffer. Eluted proteins were concentrated and buffer exchanged into endotoxin-free PBS (TekNova P0300) using Amicon Ultra-15 Centrifugal Filters (Millipore-Sigma UFC901008 or UFC901024 depending on protein size) according to the manufacturer’s protocol. Concentration continued until elution buffer represented <0.01% of the buffer composition and proteins were concentrated to 1 – 1.5mL. All fractions of purification were collected and saved. Aliquots of the cell supernatant, flowthrough, washes, elution and pure fractions were taken for later SDS-PAGE analysis. Purified, concentrated proteins were aliquoted into SafeLock tubes (Eppendorf 22600028), flash frozen in liquid nitrogen and stored at -80°C until use. Aliquots were not freeze-thawed more than once to protect protein integrity. Only endotoxin-free, sterile reagents were used to ensure compatibility with cell culture.

Protein concentration was assessed using a BCA Protein Assay Kit (Thermo 23227) following the manufacturer’s protocol in 96 well plates. Briefly, 10 µL of each sample was added to wells in a 96 well plate (Corning 3641) along with a standard curve of BSA (125 µg/mL – 2000 µg/mL) (Thermo 23209). 200 µL of prepared BCA reagent was added and incubated at 37°C for 30 minutes. Absorbance was read at 562 nm. Standard curve was fit with a linear equation and sample absorbances were used to calculate concentrations if they fit within the linear range.

#### **Cell lysate treatment and generation**

Cells were cell counted using Trypan Blue (BioRad 1450021), and 3-8 million cells were pelleted by centrifugation at 250xg for 5 min. The media was aspirated, and the pellets were washed three times using 1X TBS (Thermo J60877.K3) to remove residual cytokines and/or serum. 3-8 million HEK293T or HEK293T overexpression lines or 4 million UT-7-ES1 cells were plated in 60mm dishes or 6-well dishes. Cells were plated in serum and cytokine starved media (DMEM only (ATCC 30-2002) for HEK293T lines; IMDM only (Thermo 21056023) for UT-7-ES1 lines) to limit basal phosphorylation and incubated overnight at 37°C and 5% CO_2_ . The next day, lysis buffer (1X RIPA (CST 9806S) supplemented with protease and phosphatase inhibitors (Thermo A32961) was prepared and chilled to 4°C. Wash buffer was prepared and chilled to 4°C by supplementing 1X TBS with protease and phosphatase inhibitors. Proteins were diluted in serum and cytokine starved media to 1 µM, added to cells to yield a final in-plate concentration of 100 nM, swirled gently to mix and incubated for exactly 15 minutes at 37°C and 5% CO_2_. However, because they are suspension cells, UT-7-ES1 plates were taken out of the incubator at 10 min, moved to 15 mL falcon tubes and centrifuged for the remaining 5 minutes at 250xg. Then, media was aspirated and pellets were resuspended in wash buffer. For HEK293T lines, media is aspirated at 15mins and cells were lifted with trypLE treatment for 5 minutes or gently scraped (Greiner Bio-One 07-000-553) into 1-3 mL of wash buffer. Wash buffer was used to move the cell suspension into 15 mL falcon tubes. Cells were pelleted, wash buffer aspirated and the pellet was resuspended in 200 – 400 µL of lysis buffer. Samples were incubated on ice for 5 min and then centrifuged at 14,000xg for 15 min. Lysate was pipetted into a new tube, being careful not to disturb the pellet, and the pellet was discarded. Protein concentration of the lysates were taken by BCA. Gel samples were made in reducing Laemmli sample buffer as described above. Lysates were normalized to the same concentration to keep protein load consistent across samples.

**UT7-E-S1 proliferation assays**

UT-7-ES1 or UT-7-ES1 CD131 KO cell lines were pelleted by centrifugation at 250xg for 5 min, media was aspirated and the pellet was washed three times using TBS to remove residual cytokines. Cells were seeded in white, flat-bottom, clear 96-well plates (Millipore Sigma CLS3610) at 9.0 x 10^3^ cells per well in 90 μL of growth medium with no growth factors (IMDM + 10% FBS only). 90 µL of growth medium was added to control wells. Plates were swirled gently and then incubated overnight at 37°C in 5% CO_2_. The following day, purified proteins were serially diluted 10-fold (10^−6^ M to 10−^12^ M) in growth medium with no growth factors in 96-well plates (Corning 3641) and 10 µL of protein dilution or medium was added to the cells. Plates were gently swirled, and cells were incubated at 37°C in 5% CO_2_ for 72 h. Cell proliferation was determined by Cell Titer Glo (Promega G7572) following the manufacturer’s protocol. Briefly, 100 µL of room temperature Cell Titer Glo was added to plates. Plates were mixed by slow rotation for 2 minutes each then dark adapted for 10 minutes by covering with tinfoil or by putting the plate inside the reader for 10 minutes before reading. Luminescence was read on a BioTek Synergy Neo HTS microplate reader using the Luminescence fiber default protocol. In the main text figures, background (media only wells) luminescence was subtracted from each sample well and each well’s corrected luminescence was divided by the luminescence of untreated cells. Reported data are proliferation relative to untreated cells and are presented as mean luminescence ± SD of three replicates. In **Supplementary Figure 13**, corrected luminescence was divided by the luminescence of EPO treated samples at the same dose to compare the positive control to all other samples. Reported data are proliferation relative to EPO treated cells presented as mean luminescence ± SD of three replicates.

**Enzyme-linked immunosorbent assay (elisa)**

Recombinant human EPO-R (RND systems 963-ER) or human CD131 (AcroBio CD1-H5256-100 µg) were resuspended to 100 µg/mL in PBS following the manufacturer’s instructions, aliquoted into single-use aliquots and stored at -20°C until use. Receptors were thawed and diluted to 1µg/mL in PBS. 50 µL per well was added to coat MaxiSorp 96-well ELISA plates (Sigma M9410). Plates were incubated shaking overnight at 4°C. Plates were washed five times in wash buffer (PBS + 0.05% Tween-20 (PBST) (Millipore Sigma P9416-50ML)) by dipping the plate in a 5L container of wash buffer and tossing the liquid out. Plates were blocked with 200 µL per well of blocking buffer (PBS + 3% BSA (GoldBio A-420-10)) for 2 hrs shaking at RT. Plates were washed five times with PBST. Proteins to be tested were diluted in a ten-fold dilution series in PBS in multi-well strip plates (Corning 4413), and 50 µL per well was added. Proteins were incubated with serially diluted proteins for 3 hrs shaking at RT or overnight shaking at 4°C. Plates were washed five times with PBST to remove unbound proteins. anti-His HRP detection antibody (Abcam ab1187) was diluted 1/10,000 in PBS + 3% BSA and 50 µL was added to each well and incubated at RT shaking for 1hr. Plates were washed five times with PBST to remove unbound antibody. 100 µL of room temperature TMB Substrate solution (Thermo 34028) was added to each well and plate was monitored until the absorbance at 620nm reached >0.8 OD. Then, 2M sulfuric acid was added to stop the reaction, and the absorbance was measured at 450 nm. Data is reported as mean A_450_ ± SD of three replicates. EC_50_ and E_max_ were calculated using GraphPad Prism using nonlinear regression (log(agonist) vs. response, variable slope (four parameters).

**Genomic DNA isolation**

Genomic DNA of UT-7-ES1 cells was isolated to sequence verify the CD131 KO pool and clones. Briefly, 50,000 cells of each clonal line or the Parental UT-7-ES1 were pelleted, washed 1x with DPBS (Thermo 14190144) and frozen at -80°C until analysis. Lysis buffer was composed of 10mM Tris-HCl pH 8.0 (TekNova T1115) supplemented with 0.05% SDS and was filter sterilized with a 0.2 µm filter (Corning CLS431153). Lysis buffer was stored in aliquots until use. Immediately before cell lysis, proteinase K (NEB P8107S) was added to an aliquot of lysis buffer and diluted to 1X. Pelleted cells were thawed and 35 µL of lysis buffer was added to each sample. Tubes were incubated at 37°C for 1 hr. 25 µL was transferred to a new PCR plate and proteinase K was heat inactivated by 20 min incubation at 80°C. Lysed samples were stored either at 4°C for next day use or used immediately as template in PCR. Primers to sequence the CSF2RB locus (see primer table) were designed around the targeted cut sites in exon 4 and exon 5 so the generated PCR product would be >500 bp for sequencing. PCR was carried out as previously described with the following changes: 28 cycles, 30s/kb amplicon. After amplification, reaction was run on an agarose gel as above to verify a single band, and unpurified PCR products were sent to Plasmidsaurus for sequencing.

### **Supplementary Tables**

Supplementary Tables contain the following information:

- **Supplementary Table 1**: Plasmid list and additional sequences
- **Supplementary Table 2**: Reagent catalog numbers
- **Supplementary Table 3**: Antibody catalog numbers and dilutions
- **Supplementary Table 4:** Protein name mappings
- **Supplementary Table 5**: Generated cell lines & the figures they were used in
- **Supplementary Table 6**: Protein sequences
- **Supplementary Table 7**: Linker sequences in TscFv variants

##### **Supplementary Table 1**: Plasmid list and additional sequences

*Full plasmid sequence files provided in the supporting information data folder*

| **Short name** | **Construct** | **Backbone** | **Purpose** | **Tags** |
| --- | --- | --- | --- | --- |
| **KD29** | 4y5y-H4B | pOptivec | Secreted expression in mammalian cells | His6 |
| **KD33** | Fc5 H4A | pSecTag2A | Secreted expression in mammalian cells | His6 |
| **KD42** | human CD131-FLAG | rt42a | Overexpression of membrane protein in mammalian cells | c-myc, FLAG |
| **KD51** | CSF2RB_guide RNA 2 | rt139a | CRISPR KO of human CD131 in mammalian cells | - |
| **KD52** | CSF2RB_guide RNA 3 | rt139a | CRISPR KO of human CD131 in mammalian cells | - |
| **KD53** | human EPO-R-FLAG | rt192a | Overexpression of membrane protein in mammalian cells | FLAG |
| **KD56** | H4B FLAG tag | pOptivec (parent is KD29) | Secreted expression in mammalian cells | FLAG |
| **KD57** | 4y5y-H4B FLAG tag | pOptivec (parent is KD29) | Secreted expression in mammalian cells | FLAG |
| **KD58** | H4A his-myc | pSecTag2A | Secreted expression in mammalian cells | His6, c-myc |
| **KD59** | 4y5y H4A his myc | pSecTag2A | Secreted expression in mammalian cells | His6, c-myc |
| **KD60** | H4A FC5 his myc | pSecTag2A | Secreted expression in mammalian cells | His6, c-myc |
| **KD61** | FC5 H4A 4y5y his myc | pSecTag2A | Secreted expression in mammalian cells | His6, c-myc |
| **KD62** | KD62_4y5x_ACN_dualscfv | pSecTag2A | Secreted expression in mammalian cells | His6, c-myc |
| **KD63** | EPO-R scfv (4y5y) linked to anti-CD131 (5dwu) scfv his tag | pSecTag2A | Secreted expression in mammalian cells | His6, c-myc |
| **KD64** | EPO-R scfv (4y5y) linked to anti-CD131 (5dwu) scfv-H4B flag | pOptivec | Secreted expression in mammalian cells | FLAG |
| **KD65** | anti-CD131 (4dwu) scfv- H4A his myc | pSecTag2A | Secreted expression in mammalian cells | His6, c-myc |
| **KD66** | human CD131- V5, HA | rt42a | Overexpression of membrane protein in mammalian cells | v5, HA tag |
| **KD67** | tandem scfv, 16aa-same linker, short 2 repeats | pSecTag2A | Secreted expression in mammalian cells | His6 |
| **KD68** | tandem scfv, 7aa-same linker 1 repeat | pSecTag2A | Secreted expression in mammalian cells | His6 |
| **KD69** | tandem scfv, 3aa-shortest | pSecTag2A | Secreted expression in mammalian cells | His6 |
| **KD72** | tandem scfv, 23aa-rigid EAAAK | pSecTag2A | Secreted expression in mammalian cells | His6 |
| **KD73** | tandem scfv, 7aa-rigid EAAAK | pSecTag2A | Secreted expression in mammalian cells | His6 |
| **KD74** | tandem scfv, 7aa-rigid AP | pSecTag2A | Secreted expression in mammalian cells | His6 |
| **KD75** | tandem scfv, 23aa-semi rigid EAAAK | pSecTag2A | Secreted expression in mammalian cells | His6 |
| **KD76** | tandem scfv, 23aa-semi rigid AP | pSecTag2A | Secreted expression in mammalian cells | His6 |
| **KD77** | tandem scfv, no linker | pSecTag2A | Secreted expression in mammalian cells | His6 |
| **KD78** | dual CD131 tandem scfv control, 5dwu-5dwu | pSecTag2A | Secreted expression in mammalian cells | His6 |
| **KD79** | dual EPO-R scfv control, 4y5y-4y5y | pSecTag2A | Secreted expression in mammalian cells | His6 |
| **KD80** | EPO-R scFv (VL VH) - CD131 scFv (VH VL) tandem scFv | pSecTag2A | Secreted expression in mammalian cells | His6 |
| **KD81** | KD55 (H4B-empty- no tags) + kozak | pOptivec (parent is KD29) | Secreted expression in mammalian cells | - |
| **KD82** | KD56 (H4B FLAG tag) + kozak | KD56 | Secreted expression in mammalian cells | FLAG |
| **KD83** | KD57 (4y5y-H4B FLAG) + kozak | KD57 | Secreted expression in mammalian cells | FLAG |
| **KD84** | KD64 (EPO-R scfv (4y5y) linked to anti-CD131 (5dwu) scfv-H4B flag) + kozak | KD64 | Secreted expression in mammalian cells | FLAG |
| **KD85** | anti CD131 scfv (pdb=5dwu) myc his | KD65 | Secreted expression in mammalian cells | His6, c-myc |
| **rt42a** | S2_N-EGFP_08 | - | Overexpression in mammalian cells | - |
| **rt139a** | sgRNA_GSK3A-1 | - | Expression of gRNA in mammalian cells | - |
| **rt192a** | pHAGE_S2N-Nluc_nanoBRET_blast_1_04 | pHAGE | Overexpression in mammalian cells | - |
| **JML42** | DA330 EPO-R scFv (pdb=4y5y) myc his | pSecTag2A | Secreted expression in mammalian cells | His6, c-myc |
| **JMLN9** | EPO(S104I) | N/A | Secreted expression in mammalian cells | His6 |

**Additional sequences**

| **Abbreviated Name** | **Spacer Sequence** | **Target** |
| --- | --- | --- |
| **CSF2RB_gRNA_2** | ACAAGCGGCTTCAGGACTCT | CSF2RB Exon 5 |
| **CSF2RB_gRNA_3** | GTGACGACAAAACTCTGGCA | CSF2RB Exon 4 |

##### **Supplementary Table 2:** Reagent catalog numbers

| **Product Name** | **Vendor** | **Catalog no.** | **Notes** |
| --- | --- | --- | --- |
| Nalgene™ General Long-Term Storage Cryogenic Tubes | Thermo Scientific | 5000-1020 | Mammalian cell cryopreservation |
| DPBS | Gibco (Thermo) | 14190144 |  |
| TrypLE™ Express Enzyme (1X), phenol red | Gibco (Thermo) | 12605010 | Faster and maintains surface receptor expression better than trypsin |
| Lonza MycoAlert Mycoplasma Detection Kit | Lonza | LT07-118 |  |
| RIPA Buffer (10X) #9806 | Cell Signaling Technologies | 9806S |  |
| Pierce™ Protease Inhibitor Mini Tablets, EDTA-free | Thermo Scientific | A32955 |  |
| Amicon Ultra-15 Centrifugal Filter Unit 10kDa | Millipore Sigma | UFC901024 |  |
| Pierce™ Disposable Columns, 5 mL | Thermo Scientific | 29922 |  |
| Pierce™ BCA Protein Assay Kit | Thermo Scientific | 23227 |  |
| ZymoPURE Plasmid Miniprep Kit | Zymo Research | D4210 | for ultra-pure endotoxin-free plasmid DNA for mammalian cultures |
| Novex™ Sharp Pre-stained Protein Standard | Thermo Scientific | LC5800 |  |
| iBlot™ 2 Transfer Stacks, nitrocellulose, mini | Thermo Scientific | IB23002 |  |
| Anti-6X His tag® antibody (HRP) (ab1187) | Abcam | ab1187 |  |
| SuperSignal West Dura Extended Duration Substrate | Thermo Scientific | 34075 |  |
| Pierce™ Bovine Serum Albumin Standard Ampules, 2 mg/mL | Thermo Scientific | 23209 | For BCA protein quantification. This is supplied in BCA kit, but can order extra if you run out |
| 6X TriTrack DNA Loading Dye | Thermo Scientific | R1161 |  |
| GeneRuler 1 kb Plus DNA Ladder, ready-to-use | Thermo Scientific | SM1333 |  |
| Seakem LE Agarose | Lonza | 50004 |  |
| UltraPure™ Ethidium Bromide, 10 mg/mL | Thermo Scientific | 15585011 |  |
| Nuclease free water | NEB | B1500S |  |
| Q5 High Fidelity 2X Master Mix | NEB | M0492S |  |
| Q5® Site-Directed Mutagenesis Kit | NEB | E0554S |  |
| FreeStyle™ 293 Expression Medium | Thermo Scientific | 12338018 |  |
| 293fectin™ Transfection Reagent | Thermo Scientific | 12347019 |  |
| GlutaMAX™ Supplement | Gibco | 35050061 |  |
| NEBuilder® HiFi DNA Assembly Master Mix | NEB | E2621L |  |
| 300 mL Nuclease Free Duplex Buffer | IDT | 11-05-01-12 | for annealing complementary oligonucleotides isothermal assembly fragments |
| 1X PBS, Endotoxin Tested | TekNova | P0300 | All proteins resuspended or buffer-exchanged into this PBS |
| Pierce™ Centrifuge Columns, 10 mL | Thermo Scientific™ | 89898 |  |
| His60 Ni Superflow Resin & Buffer Set Bundle | Takara | 635677 |  |
| Corning® 1L Baffled Polycarbonate Erlenmeyer Flask with Vent Cap | Corning (Fisher Scientific) | 431403 |  |
| 500 mL Erlenmeyer Flask w/ Vent Cap, polycarbonate, sterile, 25/cs | Corning (Fisher Scientific) | CLS431145-25EA |  |
| 125 mL Polycarbonate Erlenmeyer Flask with Vent Cap | Corning (Fisher Scientific) | CLS431143-50EA | Shaker flasks for HEK293-F |
| Eppendorf Safe-Lock Tubes, 1.5 mL, Biopur®, colorless, 100 tubes, individually wrapped | Eppendorf | 22600028 | tubes for endotoxin free storage of proteins |
| Recombinant Human Erythropoietin/EPO Protein, CF | R&D systems | 11264-TC-050 |  |
| Recombinant Human Common beta Chain His-tag Protein, CF | R&D systems | 9960-CB-050 |  |
| Anti-Myc tag antibody [9E10] (HRP) (ab62928) | abcam | ab62928 |  |
| rabbit Phospho-EPO-R (Tyr426) | Thermo | PA5-106138 |  |
| rabbit anti Human Erythropoietin R Antibody | thermo | MA5-29248 | noted as "EPO-R antibody 1". Very non-specific, don't recommend use |
| rabbit anti EPO-R Monoclonal Antibody (38409) | Thermo | MA5-23824 | notes as "EPO-R antibody 2", this is the preferred EPO-R antibody, rest are too non-specific |
| Cytokine Receptor Common beta-Chain Antibody | CST | 3432S |  |
| Phospho-CSF2RB (Tyr593) Polyclonal Antibody | thermo | PA5-36654 |  |
| Phospho-Stat5 (Tyr694) Antibody | CST | 9351S |  |
| GAPDH (D16H11) XP® Rabbit mAb #5174 | CST | 5174T |  |
| Histone H3 (D1H2) XP® Rabbit mAb | CST | 4499S |  |
| Pierce Protease and Phosphatase Inhibitor Mini Tablets,EDTA-free (A32961) | Thermo | A32961 |  |
| Recombinant Human Erythropoietin R Fc Chimera Protein, CF | R&D systems | 963-ER |  |
| Recombinant Human EPO | Peprotech | 100-64 |  |
| Recombinant Human GM-CSF | Peprotech | 300-03 |  |
| CellTiter-Glo® Luminescent Cell Viability Assay | Promega | G7572 |  |
| Cas9 (S. pyogenes) (7A9-3A3) Mouse mAb | CST | 14697 |  |
| QIAGEN Plasmid Plus Maxi Kit (25) | Qiagen | 12963 |  |
| Polybrene Infection / Transfection Reagent | Millipore Sigma | TR-1003-G |  |
| Lipofectamine™ 3000 Transfection Reagent | Thermo | L3000008 |  |
| TRIS-buffered saline (TBS, 20X) pH 7.4 | Thermo | J60877.K3 |  |
| IMDM, no phenol red | Thermo | 21056023 |  |
| HRP Anti-DDDDK tag (Binds to FLAG® tag sequence) antibody [M2] | Abcam | ab49763 |  |
| 1-Step™ Ultra TMB-ELISA Substrate Solution | Thermo | 34028 |  |
| Wizard® SV Gel and PCR Clean-Up System | Promega | A9281 |  |
| 96 well MicroWell™ MaxiSorp™ flat bottom plate, pinchbar design | Sigma-Aldrich | M9410-1CS |  |
| TWEEN® 20 | Sigma-Aldrich | P9416-50ML |  |
| Plasmid PLUS endotoxin free maxi Kit | Qiagen | 12963 |  |
| KLD Enzyme Mix | NEB | M0554S |  |
| Iblot2 gel transfer device | Invitrogen | IB21001 |  |
| XCell SureLock™ Mini-Cell | Invitrogen | EI0001 |  |
| Laemmli SDS sample buffer, reducing (4X) | Thermo | J60015.AD |  |
| Tris-glycine running buffer | Thermo | LC2675 |  |
| Western blot roller | Thermo | 84747 |  |
| Gel Knife | Thermo | EI9010 |  |
| SimplyBlue™ SafeStain | Thermo | LC6065 |  |
| HyClone™ Water, Cell Culture Grade (Endotoxin-Free), Cytiva | Fisher Scientific | SH3052903 |  |
| Western Blot Box, 3 1/2" x 2 9/16" x 1", Black, 5/Pack | Southern Labware | B1200-7BK |  |
| Western Blot Box, Removable Lid, Opaque Black, for Novex Minigel, 8.6 x 8.6 x 2.8cm, 10/Pack | Southern Labware | B1300-8BK |  |
| ChemiDoc MP | Bio-Rad | 12003154 |  |
| Bovine Serum Albumin (BSA), Fraction V, Protease Free | GoldBio | A-420-10 |  |
| Blasticidin | Thermo | A1113903 |  |
| Puromycin | Thermo | A1113803 |  |
| TekNova 1X Tris HCl (pH 8.0, endotoxin free) | TekNova | T1115 |  |
| Greiner Bio-One Cell Scrapers | Greiner Bio-One | 07-000-553 |  |
| Proteinase K | NEB | NEB P8107S |  |
| 0.2 µm media filter, 500mL | Corning | CLS431153 |  |
| Stat5 (D2O6Y) Rabbit mAb | CST | 94205 |  |
| β-Actin (8H10D10) Mouse mAb #3700 | Cell Signaling Technology (CST) | 3700S |  |
| EPO-R Polyclonal Antibody | Thermo | PA591874 | noted as "EPO-R antibody 3", non-specific staining, don't recommend use |
| EPO receptor antibody | Genetek | GTX37704 | noted as "EPO-R antibody 4", works but only at very high concentrations so it becomes expensive |
| 6x-His Tag Monoclonal Antibody (4E3D10H2/E3), Alexa Fluor™ 488 | Thermo | MA1-135-A488 |  |
| 6x-His Tag Monoclonal Antibody (HIS.H8), Alexa Fluor™ 488 | Thermo | MA1-21315-A488 |  |
| Alexa Fluor® 647 Streptavidin | BioLegend | 405237 |  |
| Plate lids | Corning | 3931 |  |
| Corning® 96-well Polypropylene Cluster Tubes, 8-Tube Strip Format, Sterile, 12 Strips/Rack | Corning | 4413 |  |
| Trypan Blue | BioRad | 1450021 |  |
| Goat anti-Rabbit IgG (H+L) Secondary Antibody, HRP | Thermo | 31460 |  |
| Goat anti-Mouse IgG (H+L) Secondary Antibody, HRP | Thermo | 31430 |  |

##### **Supplementary Table 3**: Antibody catalog numbers and dilutions

*All antibodies can be used for either 1 hr shaking at room temperature or shaking at 4°C overnight unless otherwise specified. All EPO receptor antibodies stain many of non-specific bands and should be avoided if possible or used only with appropriate controls.*

| **Antibody** | **Brand** | **Catalog no.** | **Dilution** | **Used in Figure(s)** |
| --- | --- | --- | --- | --- |
| Anti-6X His tag® antibody (HRP) (ab1187) | Abcam | ab1187 | 1/10,000  (could probably use even less) | Supplementary Figure 4B |
| rabbit Phospho-EPO-R (Tyr426) | Thermo | PA5-106138 | 1/1000 |  |
| rabbit anti Human Erythropoietin R Antibody | Thermo | MA5-29248 | 1/1000 |  |
| EPO-R Monoclonal Antibody (38409) | Thermo | MA5-23824 | 1/500 |  |
| Cytokine Receptor Common beta-Chain Antibody | CST | 3432S | 1/1000 | Supplementary Figure 6A-6B, Supplementary Figure 7 |
| Phospho-CSF2RB (Tyr593) Polyclonal Antibody | Thermo | PA5-36654 | 1/1000 |  |
| Stat5 (D3N2B) Rabbit mAb #25656 | CST | 25656S | 1/1000 | Figure 3B, Figure 3C, Supplementary Figure 8 |
| Stat5 (D2O6Y) Rabbit mAb #94205 (the above antibody works fine, but this one is preferred) | CST | 94205S | 1/1000 overnight at 4°C | Figure 2C, 3D, 3E, 5A, 5C, 5E |
| Phospho-Stat5 (Tyr694) Antibody | CST | 9351S | 1/1000 | Figure 2, 3, 5, Supplementary Figure 8, Supplementary Figure 9 |
| GAPDH (D16H11) XP® Rabbit  mAb #5174 | CST | 5174T | 1/1000 |  |
| Histone H3 (D1H2) XP® Rabbit mAb | CST | 4499S | 1/1000 | Figure 2, Figure 3, Figure 5, Supplementary Figure 8, Supplementary Figure 9 |
| Cas9 (S. pyogenes) (7A9-3A3) Mouse mAb | CST | 14697 | 1/1000 |  |
| HRP Anti-DDDDK tag (Binds to FLAG® tag sequence) antibody [M2] | Abcam | ab49763 | 1/1000 | Figure 3, Supplementary Figure 7 |
| EPO-R Polyclonal Antibody | Thermo | PA591874 | 1/500 |  |
| EPO receptor antibody | Genetek | GTX37704 | 1/200 |  |
| β-Actin (8H10D10) Mouse mAb #3700 | CST | 3700S | 1/10,000 |  |
| V5 Tag Monoclonal Antibody (E10/V4RR), DyLight™ 488 | Thermo | MA5-15253-D488 | 1/1000 (1 hr at RT; do not use overnight) | Supplementary Figure 7, 8 |
| 6x-His Tag Monoclonal Antibody (HIS.H8), Alexa Fluor™ 488 | Thermo | MA1-21315-A488 | 1/1000 (1 hr at RT; do not use overnight) | Supplementary Figure 4D, 4F |
| 6x-His Tag Monoclonal Antibody (4E3D10H2/E3), Alexa Fluor™ 488 | Thermo | MA1-135-A488 | 1/1000 (1 hr at RT; do not use overnight) |  |
| Goat Anti-Mouse IgG2a heavy chain (Biotin) (ab97243) | Abcam | ab97243 | 1/1000 |  |
| Alexa Fluor® 647 Streptavidin | BioLegend | 405237 | 1/1000 (1 hr at RT; do not use overnight) |  |
| DyLight® 650 Anti-DDDDK tag (Binds to FLAG® tag sequence) antibody [M2] | Abcam | ab117492 | 1/1000 (1 hr at RT; do not use overnight) | Supplementary Figure 4E |
| Goat anti-Rabbit IgG (H+L) Secondary Antibody, HRP | Thermo | 31460 | 1/10,000 | All blots requiring secondaries unless otherwise stated |
| Goat anti-Mouse IgG (H+L) Secondary Antibody, HRP | Thermo | 31430 | 1/10,000 | All blots requiring secondaries unless otherwise stated |

##### **Supplementary Table 4**: Protein name mappings

| **Short name** | **Full name** | **Plasmids** | **Notes** |
| --- | --- | --- | --- |
| **Fc** | H4A his myc - H4B Flag | KD56 + KD58 | Modified the H4A/H4B constructs (from Wang et. Al 2019: https://doi.org/10.1080/19420862.2019.1685350) to have tags on both halves (His6 on one, FLAG on the other) to make detection of both halves easier. His tag is for purification. H4A and H4B are a modified mouse IgG2a (electrostatic steering heterodimers). |
| **CD131 scFv** | anti-CD131scfv (PDB: 5dwu) | KD50 | PDB: 5dwu |
| **EPO-R scFv** | Da330 scFv (PDB: 4y5y) | JML42 | PDB: 4y5y |
| **EPO-R BsAb** | 4y5y H4A his myc - 4y5y H4B flag (Bispecific antibody to EPO-R) | KD57+KD59 | Modified the H4A/H4B constructs (from Wang et. Al 2019: https://doi.org/10.1080/19420862.2019.1685350) to have tags on both halves (His6 on one, FLAG on the other) to make detection of both halves easier. His tag is for purification. H4A and H4B are a modified mouse IgG2a (electrostatic steering heterodimers). |
| **TscFv** | Tandem scFv version 1: 4y5y - 23aa linker - 5dwu his (may also be referenced as TscFv-LF) | KD63 | PDB: 4y5y, PDB: 5dwu |
| **TscFv-Fc** | H4A his myc - 4y5y 5dwu H4B flag | KD58 + KD64 | Modified the H4A/H4B constructs (from Wang et. Al 2019: https://doi.org/10.1080/19420862.2019.1685350) to have tags on both halves (His6 on one, FLAG on the other) to make detection of both halves easier. His tag is for purification. H4A and H4B are a modified mouse IgG2a (electrostatic steering heterodimers). |
| **BsAb** | 5dwu H4A his myc - 4y5y H4B flag | KD57 + KD65 |  |
| **SF** | tandem scFv-7aa gs linker | KD68 | Tandem scFv variants, modified from TscFv (plasmid KD63) constructs |
| **LR** | tandem scFv- 23 aa rigid linker | KD72 |  |
| **SR** | Tandem scFv- 7aa rigid linker | KD73 |  |
| **NL** | Tandem scFv- no linker | KD77 |  |
| **DC** | 5dwu-5dwu tandem scFv | KD78 |  |
| **DE** | 4y5y - 4y5y tandem scFv | KD79 |  |
| **Swap** | VL VH VH VL tandem | KD80 |  |
| **P38 (not shown)** | 4y5x - 23aa linker - 5dwu his | KD62 | This construct wouldn't express. PDB:4y5x, PDB:5dwu |

##### **Supplementary Table 5**: Generated cell lines & the figures they were used in

| **Short name** | **Cell Line** | **Transgene(s)** | **Full cell line description** | **Transduced virus (mL)** | **Notes** | **Figures** |
| --- | --- | --- | --- | --- | --- | --- |
| KDsc24 | HEK293T | +hCD131 | HEK293T + KD42 (hCD131 FLAG in puro vector) | 2 | HEK293T overexpression line used in this study (Figure 3) | Figure 3B, 3C |
| KDsc26 | HEK293T | +hEPO-R | HEK293T + KD53 (hEPO-R FLAG in blast vector) | 2 | HEK293T overexpression line used in this study (Figure 3) | Figure 3B, 3C |
| KDsc27 | HEK293T | +hEPO-R + hCD131 | HEK293T + KD42 + KD53 (hCD131 FLAG in puro, hEPO-R FLAG in blast vector) | 1+1 | HEK293T overexpression line used in this study (Figure 3) | Figure 3B, 3C |
| KDsc28 | UT-7-ES1 | spCas9 + 2 gRNAs for CD131 KO | UT-7-ES1 + Cas9 + 2 scrambled gRNAs (knockout pool) | 0.4 | spCas9 and gRNAs were packaged into virus together (not two separate viruses) | Supp Figure 2 |
| KDsc29 | UT-7-ES1 | spCas9 + 2 scrambled gRNAs | UT-7-ES1 + Cas9 + CD131 KO gRNAs (2, one targeting exon 4, one exon 5; knockout pool) | 0.4 | spCas9 and gRNAs were packaged into virus together (not two separate viruses) | Supp Figure 2 |
| KDsc30 | HEK293T | +hCD131 | HEK293T + KD66 lo virus (hCD131 in puro vector with HA and V5 tags). Short name "HCL" | 0.333 | HCL= HEK293T parental line + hCD131 lo virus. HCH= HEK293T parental line + hCD131 hi virus. HCB= HEK293T parental line + both hCD131 + hEPO-R. These cell lines are Parental HEK293T cells infected with KD66 virus or KD66 virus + KD53 virus. Hi or lo virus refers to how much I infected with originally. sometimes lower titers can lead to higher expression | Figure 3D, 3E, 5E, Supp Figure 3, Supp Figure 4, Supp Figure 5 |
| KDsc32 | KDsc26 | +hEPO-R + hCD131 | HEK293T + KD53 (HEPO-R FLAG in blast vector) + KD66 (hCD131 HA V5 tags in puro vector). Short name "HCB" | 0.333 + 0.333 |  | Figure 3D, 3E, 5E, Supp Figure 3, Supp Figure 4,  Supp Figure 5 |
| KDsc35 | KDsc29 | single cell clones of KDsc29 | Clone A, derived from KDsc29 (UT-7-ES1 + spCas9 + CD131 KO gRNAs (2, one targeting exon 4, one exon 5)) | - | Single cell clonal lines of KDsc29 isolated by limiting dilution. Clone C is the one I used in this study. Has an extra CD131-staining band knocked out compared to all other clones and the pool (Parental UT-7-ES1 has 3 bands, KO pool and most clones have 2, clone c has one). | Supp Figure 2 |
| KDsc36 | KDsc29 |  | Clone B, derived from KDsc29 (UT-7-ES1 + spCas9 + CD131 KO gRNAs (2, one targeting exon 4, one exon 5)) | - |  | Supp Figure 2 |
| KDsc37 | KDsc29 |  | Clone C, derived from KDsc29 (UT-7-ES1 + spCas9 + CD131 KO gRNAs (2, one targeting exon 4, one exon 5)) | - |  | Figure 2, 5, Supp Figure 2, Supp Figure 6 |
| KDsc43 | KDsc28 | single cell clones of KDsc28 | Clone A, derived from KDsc28 (UT-7-ES1 + spCas9 + 2 scrambled gRNAs | - | Single cell clonal lines of KDsc28 isolated by limiting dilution. | Supp Figure 2 |
| KDsc44 | KDsc28 |  | Clone B, derived from KDsc28 (UT-7-ES1 + spCas9 + 2 scrambled gRNAs | - |  | Supp Figure 2 |

##### **Supplementary Table 6**: Protein sequences

*Full plasmid sequence files provided in the supporting information data folder*

| **Abbreviated Name** | **Protein** | **Protein sequence** |
| --- | --- | --- |
| **KD29** | 4y5y-H4B | METDTLLLWVLLLWVPGSTGDHSAFAGSEVQLVESGGGLVQPGGSLRLSCAVSGFTFSKYWMTWVRQAPGKGLEWVANIKPDGSEKYYVESVKGRFTISRDNAKNSVYLQMNSVRAEDTAVYYCARVSRGGSFSDWGQGTLVTVSGGGGSGGGGSSGGGGSSQSALTQPPSASGSPGQSVTISCTGTSSDVGAYNYVSWYQQHPGKAPKLMIYEVARRPSGVPDRFSGSKSGNTASLTVSGLQAEDEADYYCSSYAGSNNFAVFGRGTKLTVLGPEPRGPTIKPCPPCKCPAPNLLGGPSVFIFPPKIKDVLMISLSPIVTCVVVDVSEDDPDVQISWFVNNVEVHTAQTQTHREDYDSTLRVVSALPIQHQDWMSGKEFKCKVNNKDLPAPIERTISKPKGSVRAPQVYVLPPPEEEMTKKQVSLTCLVKDFMPEDIYVEWTNNGKTELNYKNTEPVLDSDGSYFMYSELTVEKKNWVERNSYSCSVVHEGLHNHHTTDSFSRTPGA* |
| **KD33** | Fc5 H4A | METDTLLLWVLLLWVPGSTGDEVQLQASGGGLVQAGGSLRLSCAASGFKITHYTMGWFRQAPGKEREFVSRITWGGDNTFYSNSVKGRFTISRDNAKNTVYLQMNSLKPEDTADYYCAAGSTSTATPLRVDYWGKGTQVTVSSEPRGPTIKPCPPCKCPAPNLLGGPSVFIFPPKIKDVLMISLSPIVTCVVVDVSEDDPDVQISWFVNNVEVHTAQTQTHREDYDSTLRVVSALPIQHQDWMSGKEFKCKVNNKDLPAPIERTISKPKGSVRAPQVYVLPPPEKEMTKKQVSLTCLVKDFMPEDIYVEWTNNGKTELNYKNTEPVLKSDGSYFMYSKLTVEKKNWVERNSYSCSVVHEGLHNHHTTKSFSRTPGAGPEQKLISEEDLNSAVDHHHHHH* |
| **KD42** | human CD131-FLAG | MVLAQGLLSMALLALCWERSLAGAEETIPLQTLRCYNDYTSHITCRWADTQDAQRLVNVTLIRRVNEDLLEPVSCDLSDDMPWSACPHPRCVPRRCVIPCQSFVVTDVDYFSFQPDRPLGTRLTVTLTQHVQPPEPRDLQISTDQDHFLLTWSVALGSPQSHWLSPGDLEFEVVYKRLQDSWEDAAILLSNTSQATLGPEHLMPSSTYVARVRTRLAPGSRLSGRPSKWSPEVCWDSQPGDEAQPQNLECFFDGAAVLSCSWEVRKEVASSVSFGLFYKPSPDAGEEECSPVLREGLGSLHTRHHCQIPVPDPATHGQYIVSVQPRRAEKHIKSSVNIQMAPPSLNVTKDGDSYSLRWETMKMRYEHIDHTFEIQYRKDTATWKDSKTETLQNAHSMALPALEPSTRYWARVRVRTSRTGYNGIWSEWSEARSWDTESVLPMWVLALIVIFLTIAVLLALRFCGIYGYRLRRKWEEKIPNPSKSHLFQNGSAELWPPGSMSAFTSGSPPHQGPWGSRFPELEGVFPVGFGDSEVSPLTIEDPKHVCDPPSGPDTTPAASDLPTEQPPSPQPGPPAASHTPEKQASSFDFNGPYLGPPHSRSLPDILGQPEPPQEGGSQKSPPPGSLEYLCLPAGGQVQLVPLAQAMGPGQAVEVERRPSQGAAGSPSLESGGGPAPPALGPRVGGQDQKDSPVAIPMSSGDTEDPGVASGYVSSADLVFTPNSGASSVSLVPSLGLPSDQTPSLCPGLASGPPGAPGPVKSGFEGYVELPPIEGRSPRSPRNNPVPPEAKSPVLNPGERPADVSPTSPQPEGLLVLQQVGDYCFLPGLGPGPLSLRSKPSSPGPGPEIKNLDQAFQVKKPPGQAVPQVPVIQLFKALKQQDYLSLPPWEVNKPGEVCGSEQKLISEEDLGGGSDYKDDDDK* |
| **KD53** | human EPO-R-FLAG | MDHLGASLWPQVGSLCLLLAGAAWAPPPNLPDPKFESKAALLAARGPEELLCFTERLEDLVCFWEEAASAGVGPGNYSFSYQLEDEPWKLCRLHQAPTARGAVRFWCSLPTADTSSFVPLELRVTAASGAPRYHRVIHINEVVLLDAPVGLVARLADESGHVVLRWLPPPETPMTSHIRYEVDVSAGNGAGSVQRVEILEGRTECVLSNLRGRTRYTFAVRARMAEPSFGGFWSAWSEPVSLLTPSDLDPLILTLSLILVVILVLLTVLALLSHRRALKQKIWPGIPSPESEFEGLFTTHKGNFQLWLYQNDGCLWWSPCTPFTEDPPASLEVLSERCWGTMQAVEPGTDDEGPLLEPVGSEHAQDTYLVLDKWLLPRNPPSEDLPGPGGSVDIVAMDEGSEASSCSSALASKPSPEGASAASFEYTILDPSSQLLRPWTLCPELPPTPPHLKYLYLVVSDSGISTDYSSGDSQGAQGGLSDGPYSNPYENSLIPAAEPLPPSYVACSGSEQKLISEEDLGGGSDYKDDDDK* |
| **KD56** | H4B FLAG tag | METDTLLLWVLLLWVPGSTGDEPRGPTIKPCPPCKCPAPNLLGGPSVFIFPPKIKDVLMISLSPIVTCVVVDVSEDDPDVQISWFVNNVEVHTAQTQTHREDYDSTLRVVSALPIQHQDWMSGKEFKCKVNNKDLPAPIERTISKPKGSVRAPQVYVLPPPEEEMTKKQVSLTCLVKDFMPEDIYVEWTNNGKTELNYKNTEPVLDSDGSYFMYSELTVEKKNWVERNSYSCSVVHEGLHNHHTTDSFSRTPGADYKDDDDK* |
| **KD57** | 4y5y-H4B FLAG tag | METDTLLLWVLLLWVPGSTGDHSAFAGSEVQLVESGGGLVQPGGSLRLSCAVSGFTFSKYWMTWVRQAPGKGLEWVANIKPDGSEKYYVESVKGRFTISRDNAKNSVYLQMNSVRAEDTAVYYCARVSRGGSFSDWGQGTLVTVSGGGGSGGGGSSGGGGSSQSALTQPPSASGSPGQSVTISCTGTSSDVGAYNYVSWYQQHPGKAPKLMIYEVARRPSGVPDRFSGSKSGNTASLTVSGLQAEDEADYYCSSYAGSNNFAVFGRGTKLTVLGPEPRGPTIKPCPPCKCPAPNLLGGPSVFIFPPKIKDVLMISLSPIVTCVVVDVSEDDPDVQISWFVNNVEVHTAQTQTHREDYDSTLRVVSALPIQHQDWMSGKEFKCKVNNKDLPAPIERTISKPKGSVRAPQVYVLPPPEEEMTKKQVSLTCLVKDFMPEDIYVEWTNNGKTELNYKNTEPVLDSDGSYFMYSELTVEKKNWVERNSYSCSVVHEGLHNHHTTDSFSRTPGADYKDDDDK* |
| **KD58** | H4A his-myc | METDTLLLWVLLLWVPGSTGDEPRGPTIKPCPPCKCPAPNLLGGPSVFIFPPKIKDVLMISLSPIVTCVVVDVSEDDPDVQISWFVNNVEVHTAQTQTHREDYDSTLRVVSALPIQHQDWMSGKEFKCKVNNKDLPAPIERTISKPKGSVRAPQVYVLPPPEKEMTKKQVSLTCLVKDFMPEDIYVEWTNNGKTELNYKNTEPVLKSDGSYFMYSKLTVEKKNWVERNSYSCSVVHEGLHNHHTTKSFSRTPGAGPEQKLISEEDLNSAVDHHHHHH* |
| **KD59** | 4y5y H4A his myc | METDTLLLWVLLLWVPGSTGDHSAFAGSEVQLVESGGGLVQPGGSLRLSCAVSGFTFSKYWMTWVRQAPGKGLEWVANIKPDGSEKYYVESVKGRFTISRDNAKNSVYLQMNSVRAEDTAVYYCARVSRGGSFSDWGQGTLVTVSGGGGSGGGGSSGGGGSSQSALTQPPSASGSPGQSVTISCTGTSSDVGAYNYVSWYQQHPGKAPKLMIYEVARRPSGVPDRFSGSKSGNTASLTVSGLQAEDEADYYCSSYAGSNNFAVFGRGTKLTVLGPEPRGPTIKPCPPCKCPAPNLLGGPSVFIFPPKIKDVLMISLSPIVTCVVVDVSEDDPDVQISWFVNNVEVHTAQTQTHREDYDSTLRVVSALPIQHQDWMSGKEFKCKVNNKDLPAPIERTISKPKGSVRAPQVYVLPPPEKEMTKKQVSLTCLVKDFMPEDIYVEWTNNGKTELNYKNTEPVLKSDGSYFMYSKLTVEKKNWVERNSYSCSVVHEGLHNHHTTKSFSRTPGAGPEQKLISEEDLNSAVDHHHHHH* |
| **KD60** | H4A FC5 his myc | METDTLLLWVLLLWVPGSTGDEPRGPTIKPCPPCKCPAPNLLGGPSVFIFPPKIKDVLMISLSPIVTCVVVDVSEDDPDVQISWFVNNVEVHTAQTQTHREDYDSTLRVVSALPIQHQDWMSGKEFKCKVNNKDLPAPIERTISKPKGSVRAPQVYVLPPPEKEMTKKQVSLTCLVKDFMPEDIYVEWTNNGKTELNYKNTEPVLKSDGSYFMYSKLTVEKKNWVERNSYSCSVVHEGLHNHHTTKSFSRTPGAEVQLQASGGGLVQAGGSLRLSCAASGFKITHYTMGWFRQAPGKEREFVSRITWGGDNTFYSNSVKGRFTISRDNAKNTVYLQMNSLKPEDTADYYCAAGSTSTATPLRVDYWGKGTQVTVSSGPEQKLISEEDLNSAVDHHHHHH* |
| **KD61** | FC5 H4A 4y5y his myc | METDTLLLWVLLLWVPGSTGDEVQLQASGGGLVQAGGSLRLSCAASGFKITHYTMGWFRQAPGKEREFVSRITWGGDNTFYSNSVKGRFTISRDNAKNTVYLQMNSLKPEDTADYYCAAGSTSTATPLRVDYWGKGTQVTVSSEPRGPTIKPCPPCKCPAPNLLGGPSVFIFPPKIKDVLMISLSPIVTCVVVDVSEDDPDVQISWFVNNVEVHTAQTQTHREDYDSTLRVVSALPIQHQDWMSGKEFKCKVNNKDLPAPIERTISKPKGSVRAPQVYVLPPPEKEMTKKQVSLTCLVKDFMPEDIYVEWTNNGKTELNYKNTEPVLKSDGSYFMYSKLTVEKKNWVERNSYSCSVVHEGLHNHHTTKSFSRTPGAGPEQKLISEEDLNSAVDHHHHHH* |
| **KD62** | KD62_4y5x_ACN_dualscfv | METDTLLLWVLLLWVPGSTGDQPPSVSEAPGQRVTISCSGSSSNIGNNAVSWYQQLPGKAPTLLIYYDNLLPSGVSDRFSGSKSGTSASLAISGLQSEDEADYYCAAWDDSLNDWVFGGGTKVTVNGSGGGSSSGGSGSSGGSGGGSSGEVQLLESGGGLVQPGGSLRLSCAASGFTFSSYAMSWVRQAPGKGLEWVSAISGSGGSTYYADSVKGRFTISRDNSKNTLYLQMNSLRAEDTAVYYCVKDRVAVAGKGSYYFDSWGRGTTVTVSSGSGEGGSESSGEGSGESSGEGSGEVQLLESGGGLVQPGGSLRLSCAASGFTFPWYRVHWVRQAPGKGLEWVSSIRSSGGFPYYADSVKGRFTISRDNSKNTLYLQMNSLRAEDTAVYYCARFYDSFFDIWGQGTTVTVSSGSGGGSSSGGSGSSGGSGDIQMTQSPSSVSASVGDRVTITCRASQGISSWLAWYQQKPGKAPKLLIYAASSLQSGVPSRFSGSGSGTDFTLTISSLQPEDFATYYCQQANSFPITFGQGTKLEIKHHHHHH* |
| **KD63 (TscFv)** | EPO-R scfv (4y5y) linked to anti-CD131 (5dwu) scfv his tag | METDTLLLWVLLLWVPGSTGDHSAFAGSEVQLVESGGGLVQPGGSLRLSCAVSGFTFSKYWMTWVRQAPGKGLEWVANIKPDGSEKYYVESVKGRFTISRDNAKNSVYLQMNSVRAEDTAVYYCARVSRGGSFSDWGQGTLVTVSGGGGSGGGGSSGGGGSSQSALTQPPSASGSPGQSVTISCTGTSSDVGAYNYVSWYQQHPGKAPKLMIYEVARRPSGVPDRFSGSKSGNTASLTVSGLQAEDEADYYCSSYAGSNNFAVFGRGTKLTVLGPGSGEGGSESSGEGSGESSGEGSGEVQLLESGGGLVQPGGSLRLSCAASGFTFPWYRVHWVRQAPGKGLEWVSSIRSSGGFPYYADSVKGRFTISRDNSKNTLYLQMNSLRAEDTAVYYCARFYDSFFDIWGQGTTVTVSSGSGGGSSSGGSGSSGGSGDIQMTQSPSSVSASVGDRVTITCRASQGISSWLAWYQQKPGKAPKLLIYAASSLQSGVPSRFSGSGSGTDFTLTISSLQPEDFATYYCQQANSFPITFGQGTKLEIKHHHHHH* |
| **KD64** | EPO-R scfv (4y5y) linked to anti-CD131 (5dwu) scfv-H4B flag | METDTLLLWVLLLWVPGSTGDHSAFAGSEVQLVESGGGLVQPGGSLRLSCAVSGFTFSKYWMTWVRQAPGKGLEWVANIKPDGSEKYYVESVKGRFTISRDNAKNSVYLQMNSVRAEDTAVYYCARVSRGGSFSDWGQGTLVTVSGGGGSGGGGSSGGGGSSQSALTQPPSASGSPGQSVTISCTGTSSDVGAYNYVSWYQQHPGKAPKLMIYEVARRPSGVPDRFSGSKSGNTASLTVSGLQAEDEADYYCSSYAGSNNFAVFGRGTKLTVLGPGSGEGGSESSGEGSGESSGEGSGEVQLLESGGGLVQPGGSLRLSCAASGFTFPWYRVHWVRQAPGKGLEWVSSIRSSGGFPYYADSVKGRFTISRDNSKNTLYLQMNSLRAEDTAVYYCARFYDSFFDIWGQGTTVTVSSGSGGGSSSGGSGSSGGSGDIQMTQSPSSVSASVGDRVTITCRASQGISSWLAWYQQKPGKAPKLLIYAASSLQSGVPSRFSGSGSGTDFTLTISSLQPEDFATYYCQQANSFPITFGQGTKLEIKEPRGPTIKPCPPCKCPAPNLLGGPSVFIFPPKIKDVLMISLSPIVTCVVVDVSEDDPDVQISWFVNNVEVHTAQTQTHREDYDSTLRVVSALPIQHQDWMSGKEFKCKVNNKDLPAPIERTISKPKGSVRAPQVYVLPPPEEEMTKKQVSLTCLVKDFMPEDIYVEWTNNGKTELNYKNTEPVLDSDGSYFMYSELTVEKKNWVERNSYSCSVVHEGLHNHHTTDSFSRTPGADYKDDDDK* |
| **KD65** | anti-CD131 (4dwu) scfv- H4A his myc | METDTLLLWVLLLWVPGSTGDEVQLLESGGGLVQPGGSLRLSCAASGFTFPWYRVHWVRQAPGKGLEWVSSIRSSGGFPYYADSVKGRFTISRDNSKNTLYLQMNSLRAEDTAVYYCARFYDSFFDIWGQGTTVTVSSGSGGGSSSGGSGSSGGSGDIQMTQSPSSVSASVGDRVTITCRASQGISSWLAWYQQKPGKAPKLLIYAASSLQSGVPSRFSGSGSGTDFTLTISSLQPEDFATYYCQQANSFPITFGQGTKLEIKEPRGPTIKPCPPCKCPAPNLLGGPSVFIFPPKIKDVLMISLSPIVTCVVVDVSEDDPDVQISWFVNNVEVHTAQTQTHREDYDSTLRVVSALPIQHQDWMSGKEFKCKVNNKDLPAPIERTISKPKGSVRAPQVYVLPPPEKEMTKKQVSLTCLVKDFMPEDIYVEWTNNGKTELNYKNTEPVLKSDGSYFMYSKLTVEKKNWVERNSYSCSVVHEGLHNHHTTKSFSRTPGAGPEQKLISEEDLNSAVDHHHHHH* |
| **KD66** | human CD131- V5, HA | MVLAQGLLSMALLALCWERSLAGAEETIPLQTLRCYNDYTSHITCRWADTQDAQRLVNVTLIRRVNEDLLEPVSCDLSDDMPWSACPHPRCVPRRCVIPCQSFVVTDVDYFSFQPDRPLGTRLTVTLTQHVQPPEPRDLQISTDQDHFLLTWSVALGSPQSHWLSPGDLEFEVVYKRLQDSWEDAAILLSNTSQATLGPEHLMPSSTYVARVRTRLAPGSRLSGRPSKWSPEVCWDSQPGDEAQPQNLECFFDGAAVLSCSWEVRKEVASSVSFGLFYKPSPDAGEEECSPVLREGLGSLHTRHHCQIPVPDPATHGQYIVSVQPRRAEKHIKSSVNIQMAPPSLNVTKDGDSYSLRWETMKMRYEHIDHTFEIQYRKDTATWKDSKTETLQNAHSMALPALEPSTRYWARVRVRTSRTGYNGIWSEWSEARSWDTESVLPMWVLALIVIFLTIAVLLALRFCGIYGYRLRRKWEEKIPNPSKSHLFQNGSAELWPPGSMSAFTSGSPPHQGPWGSRFPELEGVFPVGFGDSEVSPLTIEDPKHVCDPPSGPDTTPAASDLPTEQPPSPQPGPPAASHTPEKQASSFDFNGPYLGPPHSRSLPDILGQPEPPQEGGSQKSPPPGSLEYLCLPAGGQVQLVPLAQAMGPGQAVEVERRPSQGAAGSPSLESGGGPAPPALGPRVGGQDQKDSPVAIPMSSGDTEDPGVASGYVSSADLVFTPNSGASSVSLVPSLGLPSDQTPSLCPGLASGPPGAPGPVKSGFEGYVELPPIEGRSPRSPRNNPVPPEAKSPVLNPGERPADVSPTSPQPEGLLVLQQVGDYCFLPGLGPGPLSLRSKPSSPGPGPEIKNLDQAFQVKKPPGQAVPQVPVIQLFKALKQQDYLSLPPWEVNKPGEVCGSYPYDVPDYAGGGSGKPIPNPLLGLDST* |
| **KD67** | tandem scfv, 16aa-same linker, short 2 repeats | METDTLLLWVLLLWVPGSTGDHSAFAGSEVQLVESGGGLVQPGGSLRLSCAVSGFTFSKYWMTWVRQAPGKGLEWVANIKPDGSEKYYVESVKGRFTISRDNAKNSVYLQMNSVRAEDTAVYYCARVSRGGSFSDWGQGTLVTVSGGGGSGGGGSSGGGGSSQSALTQPPSASGSPGQSVTISCTGTSSDVGAYNYVSWYQQHPGKAPKLMIYEVARRPSGVPDRFSGSKSGNTASLTVSGLQAEDEADYYCSSYAGSNNFAVFGRGTKLTVLGPESSGEGSGESSGEGSGEVQLLESGGGLVQPGGSLRLSCAASGFTFPWYRVHWVRQAPGKGLEWVSSIRSSGGFPYYADSVKGRFTISRDNSKNTLYLQMNSLRAEDTAVYYCARFYDSFFDIWGQGTTVTVSSGSGGGSSSGGSGSSGGSGDIQMTQSPSSVSASVGDRVTITCRASQGISSWLAWYQQKPGKAPKLLIYAASSLQSGVPSRFSGSGSGTDFTLTISSLQPEDFATYYCQQANSFPITFGQGTKLEIKHHHHHH* |
| **KD68** | tandem scfv, 7aa-same linker 1 repeat | METDTLLLWVLLLWVPGSTGDHSAFAGSEVQLVESGGGLVQPGGSLRLSCAVSGFTFSKYWMTWVRQAPGKGLEWVANIKPDGSEKYYVESVKGRFTISRDNAKNSVYLQMNSVRAEDTAVYYCARVSRGGSFSDWGQGTLVTVSGGGGSGGGGSSGGGGSSQSALTQPPSASGSPGQSVTISCTGTSSDVGAYNYVSWYQQHPGKAPKLMIYEVARRPSGVPDRFSGSKSGNTASLTVSGLQAEDEADYYCSSYAGSNNFAVFGRGTKLTVLGPESSGEGSEVQLLESGGGLVQPGGSLRLSCAASGFTFPWYRVHWVRQAPGKGLEWVSSIRSSGGFPYYADSVKGRFTISRDNSKNTLYLQMNSLRAEDTAVYYCARFYDSFFDIWGQGTTVTVSSGSGGGSSSGGSGSSGGSGDIQMTQSPSSVSASVGDRVTITCRASQGISSWLAWYQQKPGKAPKLLIYAASSLQSGVPSRFSGSGSGTDFTLTISSLQPEDFATYYCQQANSFPITFGQGTKLEIKHHHHHH* |
| **KD69** | tandem scfv, 3aa-shortest | METDTLLLWVLLLWVPGSTGDHSAFAGSEVQLVESGGGLVQPGGSLRLSCAVSGFTFSKYWMTWVRQAPGKGLEWVANIKPDGSEKYYVESVKGRFTISRDNAKNSVYLQMNSVRAEDTAVYYCARVSRGGSFSDWGQGTLVTVSGGGGSGGGGSSGGGGSSQSALTQPPSASGSPGQSVTISCTGTSSDVGAYNYVSWYQQHPGKAPKLMIYEVARRPSGVPDRFSGSKSGNTASLTVSGLQAEDEADYYCSSYAGSNNFAVFGRGTKLTVLGPGSGEVQLLESGGGLVQPGGSLRLSCAASGFTFPWYRVHWVRQAPGKGLEWVSSIRSSGGFPYYADSVKGRFTISRDNSKNTLYLQMNSLRAEDTAVYYCARFYDSFFDIWGQGTTVTVSSGSGGGSSSGGSGSSGGSGDIQMTQSPSSVSASVGDRVTITCRASQGISSWLAWYQQKPGKAPKLLIYAASSLQSGVPSRFSGSGSGTDFTLTISSLQPEDFATYYCQQANSFPITFGQGTKLEIKHHHHHH* |
| **KD72** | tandem scfv, 23aa-rigid EAAAK | METDTLLLWVLLLWVPGSTGDHSAFAGSEVQLVESGGGLVQPGGSLRLSCAVSGFTFSKYWMTWVRQAPGKGLEWVANIKPDGSEKYYVESVKGRFTISRDNAKNSVYLQMNSVRAEDTAVYYCARVSRGGSFSDWGQGTLVTVSGGGGSGGGGSSGGGGSSQSALTQPPSASGSPGQSVTISCTGTSSDVGAYNYVSWYQQHPGKAPKLMIYEVARRPSGVPDRFSGSKSGNTASLTVSGLQAEDEADYYCSSYAGSNNFAVFGRGTKLTVLGPEAAAKEAAAKEAAAKEAAAKEAAEVQLLESGGGLVQPGGSLRLSCAASGFTFPWYRVHWVRQAPGKGLEWVSSIRSSGGFPYYADSVKGRFTISRDNSKNTLYLQMNSLRAEDTAVYYCARFYDSFFDIWGQGTTVTVSSGSGGGSSSGGSGSSGGSGDIQMTQSPSSVSASVGDRVTITCRASQGISSWLAWYQQKPGKAPKLLIYAASSLQSGVPSRFSGSGSGTDFTLTISSLQPEDFATYYCQQANSFPITFGQGTKLEIKHHHHHH* |
| **KD73** | tandem scfv, 7aa-rigid EAAAK | METDTLLLWVLLLWVPGSTGDHSAFAGSEVQLVESGGGLVQPGGSLRLSCAVSGFTFSKYWMTWVRQAPGKGLEWVANIKPDGSEKYYVESVKGRFTISRDNAKNSVYLQMNSVRAEDTAVYYCARVSRGGSFSDWGQGTLVTVSGGGGSGGGGSSGGGGSSQSALTQPPSASGSPGQSVTISCTGTSSDVGAYNYVSWYQQHPGKAPKLMIYEVARRPSGVPDRFSGSKSGNTASLTVSGLQAEDEADYYCSSYAGSNNFAVFGRGTKLTVLGPEAAAKEAEVQLLESGGGLVQPGGSLRLSCAASGFTFPWYRVHWVRQAPGKGLEWVSSIRSSGGFPYYADSVKGRFTISRDNSKNTLYLQMNSLRAEDTAVYYCARFYDSFFDIWGQGTTVTVSSGSGGGSSSGGSGSSGGSGDIQMTQSPSSVSASVGDRVTITCRASQGISSWLAWYQQKPGKAPKLLIYAASSLQSGVPSRFSGSGSGTDFTLTISSLQPEDFATYYCQQANSFPITFGQGTKLEIKHHHHHH* |
| **KD74** | tandem scfv, 7aa-rigid AP | METDTLLLWVLLLWVPGSTGDHSAFAGSEVQLVESGGGLVQPGGSLRLSCAVSGFTFSKYWMTWVRQAPGKGLEWVANIKPDGSEKYYVESVKGRFTISRDNAKNSVYLQMNSVRAEDTAVYYCARVSRGGSFSDWGQGTLVTVSGGGGSGGGGSSGGGGSSQSALTQPPSASGSPGQSVTISCTGTSSDVGAYNYVSWYQQHPGKAPKLMIYEVARRPSGVPDRFSGSKSGNTASLTVSGLQAEDEADYYCSSYAGSNNFAVFGRGTKLTVLGPAPAPAPAEVQLLESGGGLVQPGGSLRLSCAASGFTFPWYRVHWVRQAPGKGLEWVSSIRSSGGFPYYADSVKGRFTISRDNSKNTLYLQMNSLRAEDTAVYYCARFYDSFFDIWGQGTTVTVSSGSGGGSSSGGSGSSGGSGDIQMTQSPSSVSASVGDRVTITCRASQGISSWLAWYQQKPGKAPKLLIYAASSLQSGVPSRFSGSGSGTDFTLTISSLQPEDFATYYCQQANSFPITFGQGTKLEIKHHHHHH* |
| **KD75** | tandem scfv, 23aa-semi rigid EAAAK | METDTLLLWVLLLWVPGSTGDHSAFAGSEVQLVESGGGLVQPGGSLRLSCAVSGFTFSKYWMTWVRQAPGKGLEWVANIKPDGSEKYYVESVKGRFTISRDNAKNSVYLQMNSVRAEDTAVYYCARVSRGGSFSDWGQGTLVTVSGGGGSGGGGSSGGGGSSQSALTQPPSASGSPGQSVTISCTGTSSDVGAYNYVSWYQQHPGKAPKLMIYEVARRPSGVPDRFSGSKSGNTASLTVSGLQAEDEADYYCSSYAGSNNFAVFGRGTKLTVLGPSGGGGSEAAAKEAAAKGGGSGGSEVQLLESGGGLVQPGGSLRLSCAASGFTFPWYRVHWVRQAPGKGLEWVSSIRSSGGFPYYADSVKGRFTISRDNSKNTLYLQMNSLRAEDTAVYYCARFYDSFFDIWGQGTTVTVSSGSGGGSSSGGSGSSGGSGDIQMTQSPSSVSASVGDRVTITCRASQGISSWLAWYQQKPGKAPKLLIYAASSLQSGVPSRFSGSGSGTDFTLTISSLQPEDFATYYCQQANSFPITFGQGTKLEIKHHHHHH* |
| **KD76** | tandem scfv, 23aa-semi rigid AP | METDTLLLWVLLLWVPGSTGDHSAFAGSEVQLVESGGGLVQPGGSLRLSCAVSGFTFSKYWMTWVRQAPGKGLEWVANIKPDGSEKYYVESVKGRFTISRDNAKNSVYLQMNSVRAEDTAVYYCARVSRGGSFSDWGQGTLVTVSGGGGSGGGGSSGGGGSSQSALTQPPSASGSPGQSVTISCTGTSSDVGAYNYVSWYQQHPGKAPKLMIYEVARRPSGVPDRFSGSKSGNTASLTVSGLQAEDEADYYCSSYAGSNNFAVFGRGTKLTVLGPGGSGGGGSAPAPAPAGGGGSGGSEVQLLESGGGLVQPGGSLRLSCAASGFTFPWYRVHWVRQAPGKGLEWVSSIRSSGGFPYYADSVKGRFTISRDNSKNTLYLQMNSLRAEDTAVYYCARFYDSFFDIWGQGTTVTVSSGSGGGSSSGGSGSSGGSGDIQMTQSPSSVSASVGDRVTITCRASQGISSWLAWYQQKPGKAPKLLIYAASSLQSGVPSRFSGSGSGTDFTLTISSLQPEDFATYYCQQANSFPITFGQGTKLEIKHHHHHH* |
| **KD77** | tandem scfv, no linker | METDTLLLWVLLLWVPGSTGDHSAFAGSEVQLVESGGGLVQPGGSLRLSCAVSGFTFSKYWMTWVRQAPGKGLEWVANIKPDGSEKYYVESVKGRFTISRDNAKNSVYLQMNSVRAEDTAVYYCARVSRGGSFSDWGQGTLVTVSGGGGSGGGGSSGGGGSSQSALTQPPSASGSPGQSVTISCTGTSSDVGAYNYVSWYQQHPGKAPKLMIYEVARRPSGVPDRFSGSKSGNTASLTVSGLQAEDEADYYCSSYAGSNNFAVFGRGTKLTVLGPEVQLLESGGGLVQPGGSLRLSCAASGFTFPWYRVHWVRQAPGKGLEWVSSIRSSGGFPYYADSVKGRFTISRDNSKNTLYLQMNSLRAEDTAVYYCARFYDSFFDIWGQGTTVTVSSGSGGGSSSGGSGSSGGSGDIQMTQSPSSVSASVGDRVTITCRASQGISSWLAWYQQKPGKAPKLLIYAASSLQSGVPSRFSGSGSGTDFTLTISSLQPEDFATYYCQQANSFPITFGQGTKLEIKHHHHHH* |
| **KD78** | dual CD131 tandem scfv control, 5dwu-5dwu | METDTLLLWVLLLWVPGSTGDEVQLLESGGGLVQPGGSLRLSCAASGFTFPWYRVHWVRQAPGKGLEWVSSIRSSGGFPYYADSVKGRFTISRDNSKNTLYLQMNSLRAEDTAVYYCARFYDSFFDIWGQGTTVTVSSGSGGGSSSGGSGSSGGSGDIQMTQSPSSVSASVGDRVTITCRASQGISSWLAWYQQKPGKAPKLLIYAASSLQSGVPSRFSGSGSGTDFTLTISSLQPEDFATYYCQQANSFPITFGQGTKLEIKGSGEGGSESSGEGSGESSGEGSGEVQLLESGGGLVQPGGSLRLSCAASGFTFPWYRVHWVRQAPGKGLEWVSSIRSSGGFPYYADSVKGRFTISRDNSKNTLYLQMNSLRAEDTAVYYCARFYDSFFDIWGQGTTVTVSSGSGGGSSSGGSGSSGGSGDIQMTQSPSSVSASVGDRVTITCRASQGISSWLAWYQQKPGKAPKLLIYAASSLQSGVPSRFSGSGSGTDFTLTISSLQPEDFATYYCQQANSFPITFGQGTKLEIKHHHHHH* |
| **KD79** | dual EPO-R scfv control, 4y5y-4y5y | METDTLLLWVLLLWVPGSTGDHSAFAGSEVQLVESGGGLVQPGGSLRLSCAVSGFTFSKYWMTWVRQAPGKGLEWVANIKPDGSEKYYVESVKGRFTISRDNAKNSVYLQMNSVRAEDTAVYYCARVSRGGSFSDWGQGTLVTVSGGGGSGGGGSSGGGGSSQSALTQPPSASGSPGQSVTISCTGTSSDVGAYNYVSWYQQHPGKAPKLMIYEVARRPSGVPDRFSGSKSGNTASLTVSGLQAEDEADYYCSSYAGSNNFAVFGRGTKLTVLGPGSGEGGSESSGEGSGESSGEGSGEVQLVESGGGLVQPGGSLRLSCAVSGFTFSKYWMTWVRQAPGKGLEWVANIKPDGSEKYYVESVKGRFTISRDNAKNSVYLQMNSVRAEDTAVYYCARVSRGGSFSDWGQGTLVTVSGGGGSGGGGSSGGGGSSQSALTQPPSASGSPGQSVTISCTGTSSDVGAYNYVSWYQQHPGKAPKLMIYEVARRPSGVPDRFSGSKSGNTASLTVSGLQAEDEADYYCSSYAGSNNFAVFGRGTKLTVLGPHHHHHH* |
| **KD80** | EPO-R scFv (VL VH) - CD131 scFv (VH VL) tandem scFv | METDTLLLWVLLLWVPGSTGDQSALTQPPSASGSPGQSVTISCTGTSSDVGAYNYVSWYQQHPGKAPKLMIYEVARRPSGVPDRFSGSKSGNTASLTVSGLQAEDEADYYCSSYAGSNNFAVFGRGTKLTVLGPGGGGSGGGGSSGGGGSSHSAFAGSEVQLVESGGGLVQPGGSLRLSCAVSGFTFSKYWMTWVRQAPGKGLEWVANIKPDGSEKYYVESVKGRFTISRDNAKNSVYLQMNSVRAEDTAVYYCARVSRGGSFSDWGQGTLVTVSGSGEGGSESSGEGSGESSGEGSGEVQLLESGGGLVQPGGSLRLSCAASGFTFPWYRVHWVRQAPGKGLEWVSSIRSSGGFPYYADSVKGRFTISRDNSKNTLYLQMNSLRAEDTAVYYCARFYDSFFDIWGQGTTVTVSSGSGGGSSSGGSGSSGGSGDIQMTQSPSSVSASVGDRVTITCRASQGISSWLAWYQQKPGKAPKLLIYAASSLQSGVPSRFSGSGSGTDFTLTISSLQPEDFATYYCQQANSFPITFGQGTKLEIKHHHHHH* |
| **KD81** | KD55 (H4B-empty- no tags) + kozak | METDTLLLWVLLLWVPGSTGDEPRGPTIKPCPPCKCPAPNLLGGPSVFIFPPKIKDVLMISLSPIVTCVVVDVSEDDPDVQISWFVNNVEVHTAQTQTHREDYDSTLRVVSALPIQHQDWMSGKEFKCKVNNKDLPAPIERTISKPKGSVRAPQVYVLPPPEEEMTKKQVSLTCLVKDFMPEDIYVEWTNNGKTELNYKNTEPVLDSDGSYFMYSELTVEKKNWVERNSYSCSVVHEGLHNHHTTDSFSRTPGA* |
| **KD82** | KD56 (H4B FLAG tag) + kozak | METDTLLLWVLLLWVPGSTGDEPRGPTIKPCPPCKCPAPNLLGGPSVFIFPPKIKDVLMISLSPIVTCVVVDVSEDDPDVQISWFVNNVEVHTAQTQTHREDYDSTLRVVSALPIQHQDWMSGKEFKCKVNNKDLPAPIERTISKPKGSVRAPQVYVLPPPEEEMTKKQVSLTCLVKDFMPEDIYVEWTNNGKTELNYKNTEPVLDSDGSYFMYSELTVEKKNWVERNSYSCSVVHEGLHNHHTTDSFSRTPGADYKDDDDK* |
| **KD83** | KD57 (4y5y-H4B FLAG) + kozak | METDTLLLWVLLLWVPGSTGDHSAFAGSEVQLVESGGGLVQPGGSLRLSCAVSGFTFSKYWMTWVRQAPGKGLEWVANIKPDGSEKYYVESVKGRFTISRDNAKNSVYLQMNSVRAEDTAVYYCARVSRGGSFSDWGQGTLVTVSGGGGSGGGGSSGGGGSSQSALTQPPSASGSPGQSVTISCTGTSSDVGAYNYVSWYQQHPGKAPKLMIYEVARRPSGVPDRFSGSKSGNTASLTVSGLQAEDEADYYCSSYAGSNNFAVFGRGTKLTVLGPEPRGPTIKPCPPCKCPAPNLLGGPSVFIFPPKIKDVLMISLSPIVTCVVVDVSEDDPDVQISWFVNNVEVHTAQTQTHREDYDSTLRVVSALPIQHQDWMSGKEFKCKVNNKDLPAPIERTISKPKGSVRAPQVYVLPPPEEEMTKKQVSLTCLVKDFMPEDIYVEWTNNGKTELNYKNTEPVLDSDGSYFMYSELTVEKKNWVERNSYSCSVVHEGLHNHHTTDSFSRTPGADYKDDDDK* |
| **KD84** | KD64 (EPO-R scfv (4y5y) linked to anti-CD131 (5dwu) scfv-H4B flag) + kozak | METDTLLLWVLLLWVPGSTGDHSAFAGSEVQLVESGGGLVQPGGSLRLSCAVSGFTFSKYWMTWVRQAPGKGLEWVANIKPDGSEKYYVESVKGRFTISRDNAKNSVYLQMNSVRAEDTAVYYCARVSRGGSFSDWGQGTLVTVSGGGGSGGGGSSGGGGSSQSALTQPPSASGSPGQSVTISCTGTSSDVGAYNYVSWYQQHPGKAPKLMIYEVARRPSGVPDRFSGSKSGNTASLTVSGLQAEDEADYYCSSYAGSNNFAVFGRGTKLTVLGPGSGEGGSESSGEGSGESSGEGSGEVQLLESGGGLVQPGGSLRLSCAASGFTFPWYRVHWVRQAPGKGLEWVSSIRSSGGFPYYADSVKGRFTISRDNSKNTLYLQMNSLRAEDTAVYYCARFYDSFFDIWGQGTTVTVSSGSGGGSSSGGSGSSGGSGDIQMTQSPSSVSASVGDRVTITCRASQGISSWLAWYQQKPGKAPKLLIYAASSLQSGVPSRFSGSGSGTDFTLTISSLQPEDFATYYCQQANSFPITFGQGTKLEIKEPRGPTIKPCPPCKCPAPNLLGGPSVFIFPPKIKDVLMISLSPIVTCVVVDVSEDDPDVQISWFVNNVEVHTAQTQTHREDYDSTLRVVSALPIQHQDWMSGKEFKCKVNNKDLPAPIERTISKPKGSVRAPQVYVLPPPEEEMTKKQVSLTCLVKDFMPEDIYVEWTNNGKTELNYKNTEPVLDSDGSYFMYSELTVEKKNWVERNSYSCSVVHEGLHNHHTTDSFSRTPGADYKDDDDK* |
| **KD85 (CD131 scFv (PDB: 5dwu))** | anti CD131 scfv (pdb=5dwu) myc his | METDTLLLWVLLLWVPGSTGDEVQLLESGGGLVQPGGSLRLSCAASGFTFPWYRVHWVRQAPGKGLEWVSSIRSSGGFPYYADSVKGRFTISRDNSKNTLYLQMNSLRAEDTAVYYCARFYDSFFDIWGQGTTVTVSSGSGGGSSSGGSGSSGGSGDIQMTQSPSSVSASVGDRVTITCRASQGISSWLAWYQQKPGKAPKLLIYAASSLQSGVPSRFSGSGSGTDFTLTISSLQPEDFATYYCQQANSFPITFGQGTKLEIKEQKLISEEDLNSAVDHHHHHH* |
| **rt42a** | S2_N-EGFP_08 | just used as backbone |
| **rt139a** | sgRNA_GSK3A-1 | just used as backbone |
| **rt192a** | pHAGE_S2N-Nluc_nanoBRET_blast_1_04 | just used as backbone |
| **JML42 (EPO-R scFv/3s)** | DA330 EPO-R scFv (pdb=4y5y) myc his | HSAFAGSEVQLVESGGGLVQPGGSLRLSCAVSGFTFSKYWMTWVRQAPGKGLEWVANIKPDGSEKYYVESVKGRFTISRDNAKNSVYLQMNSVRAEDTAVYYCARVSRGGSFSDWGQGTLVTVSGGGGSGGGGSSGGGGSSQSALTQPPSASGSPGQSVTISCTGTSSDVGAYNYVSWYQQHPGKAPKLMIYEVARRPSGVPDRFSGSKSGNTASLTVSGLQAEDEADYYCSSYAGSNNFAVFGRGTKLTVLGP |
