## Supplementary figures and images for "Rational design of selective bispecific EPO-R/CD131 agonists"

### 20231023_ut7epos1_cd131_ko_ntg_09 (Multichannel)_green_edit (2).tif

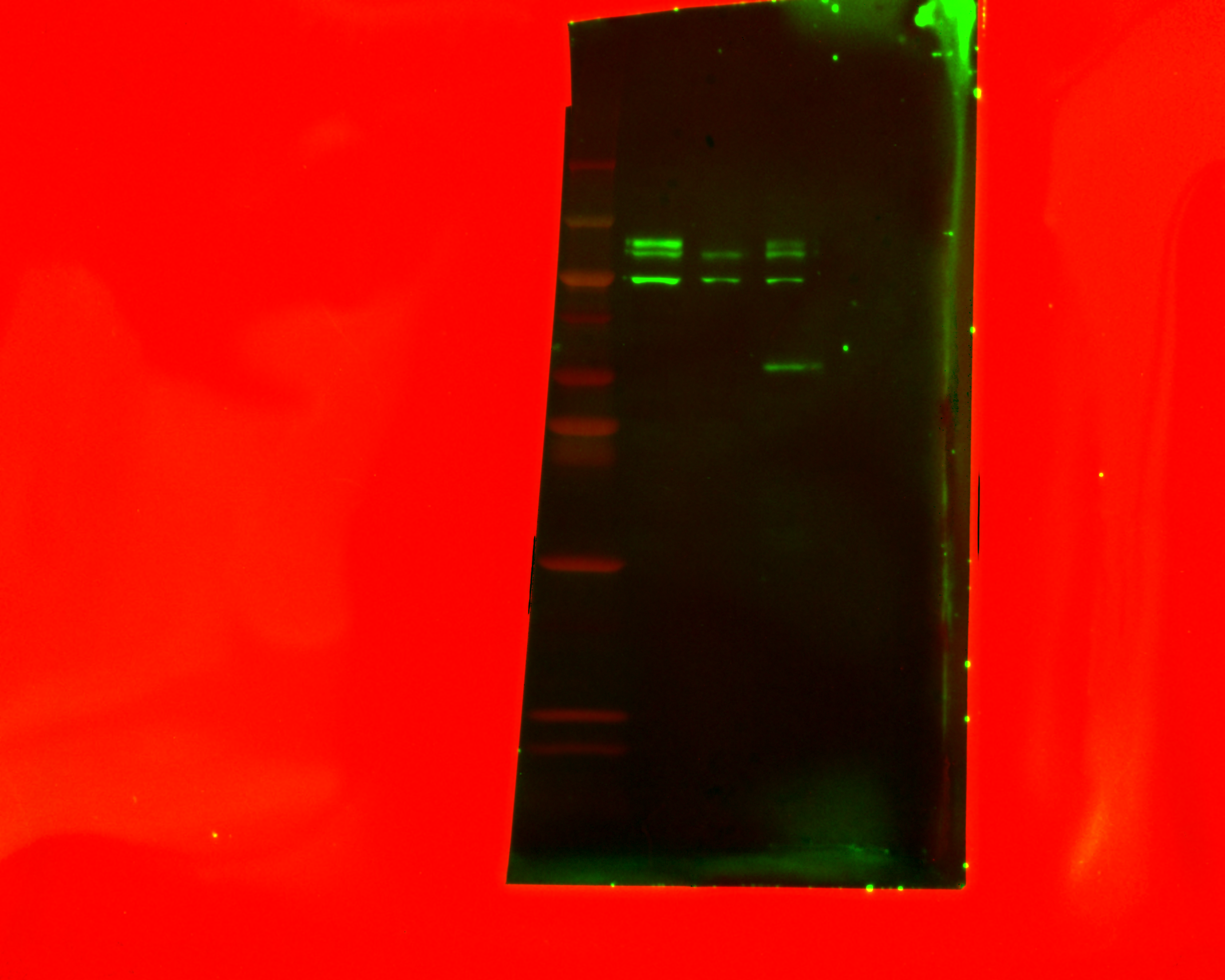

### 20231023_ut7epos1_cd131_ko_ntg_09 (Multichannel)_greyscale_edit_2 - Copy.tif

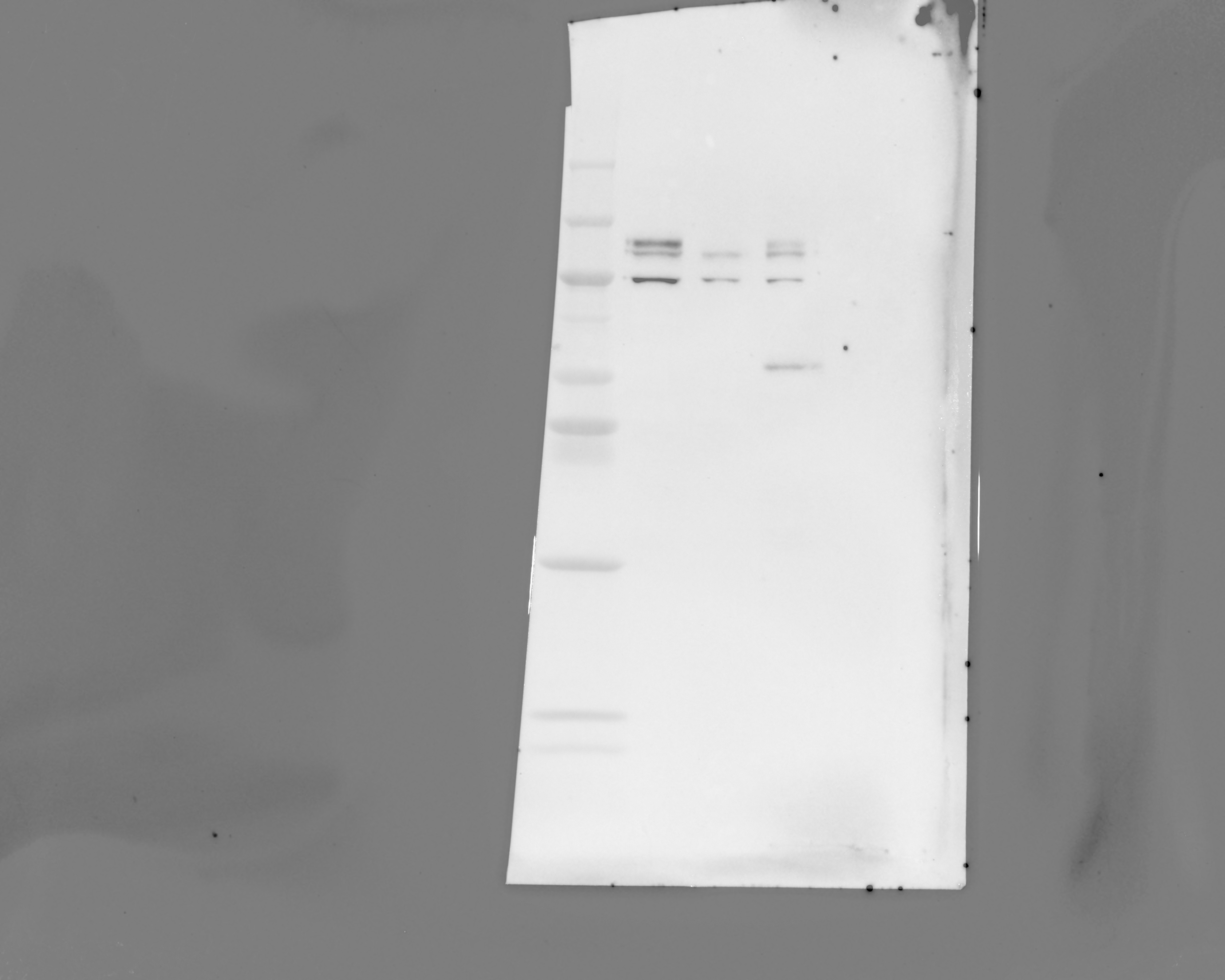

### 20231023_ut7epos1_cd131_ko_ntg_09(Chemiluminescence).tif

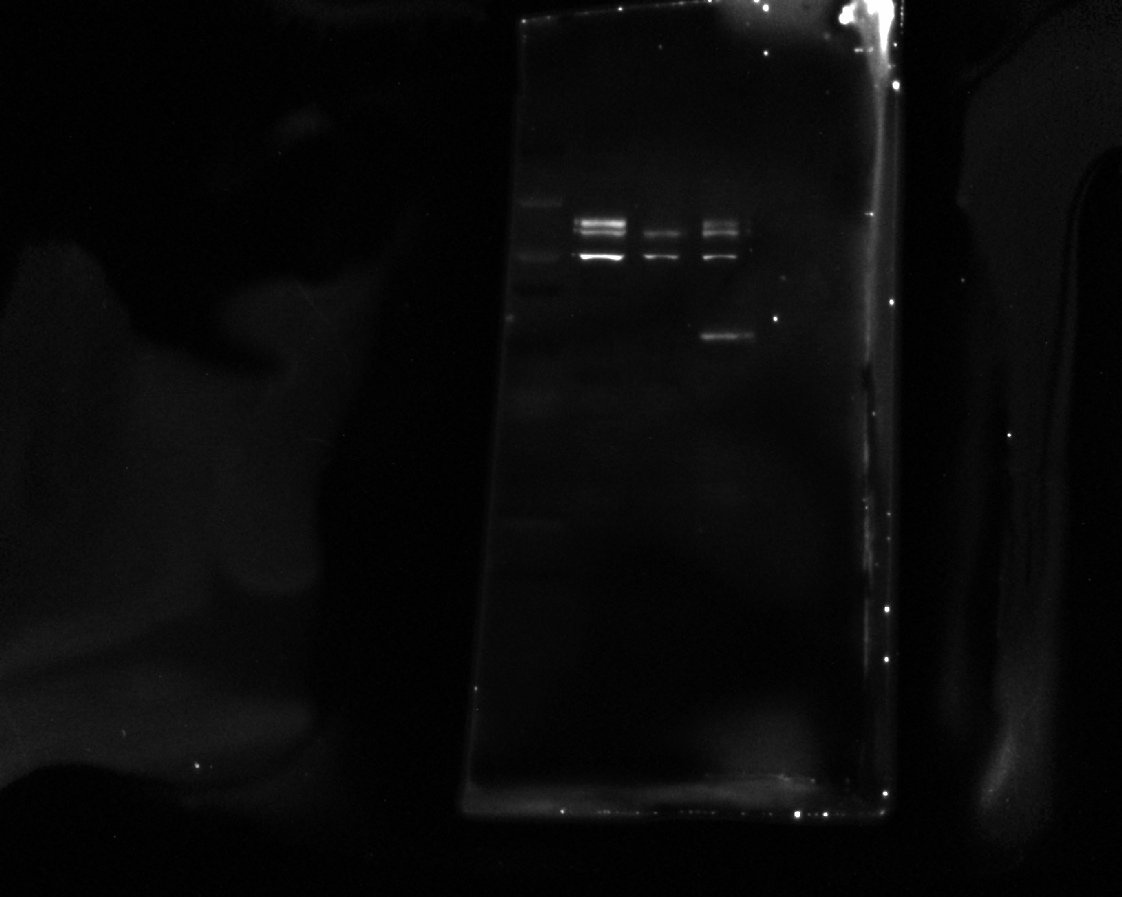

### 20231023_ut7epos1_cd131_ko_ntg_09(Colorimetric).tif

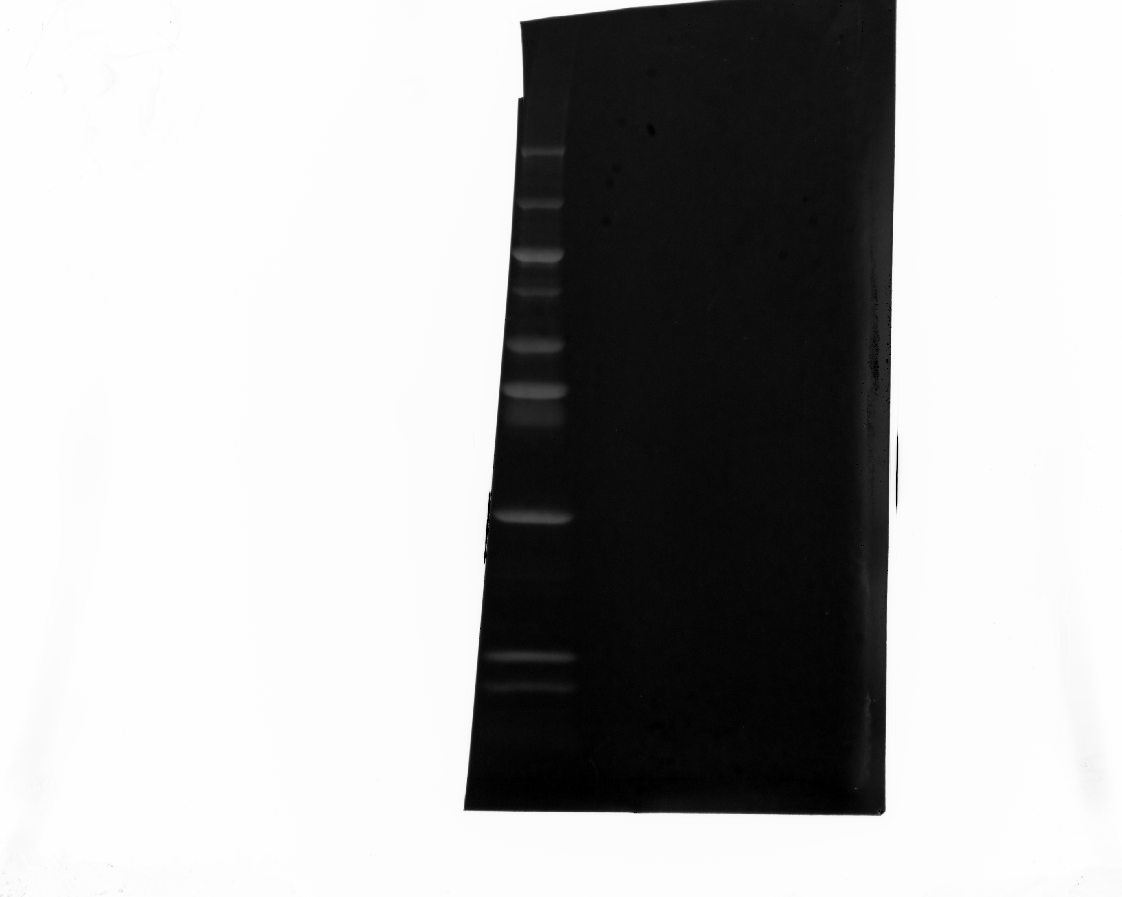

### 20240118_clonal_ut7_cd131_04 (Multi)_edit.tif

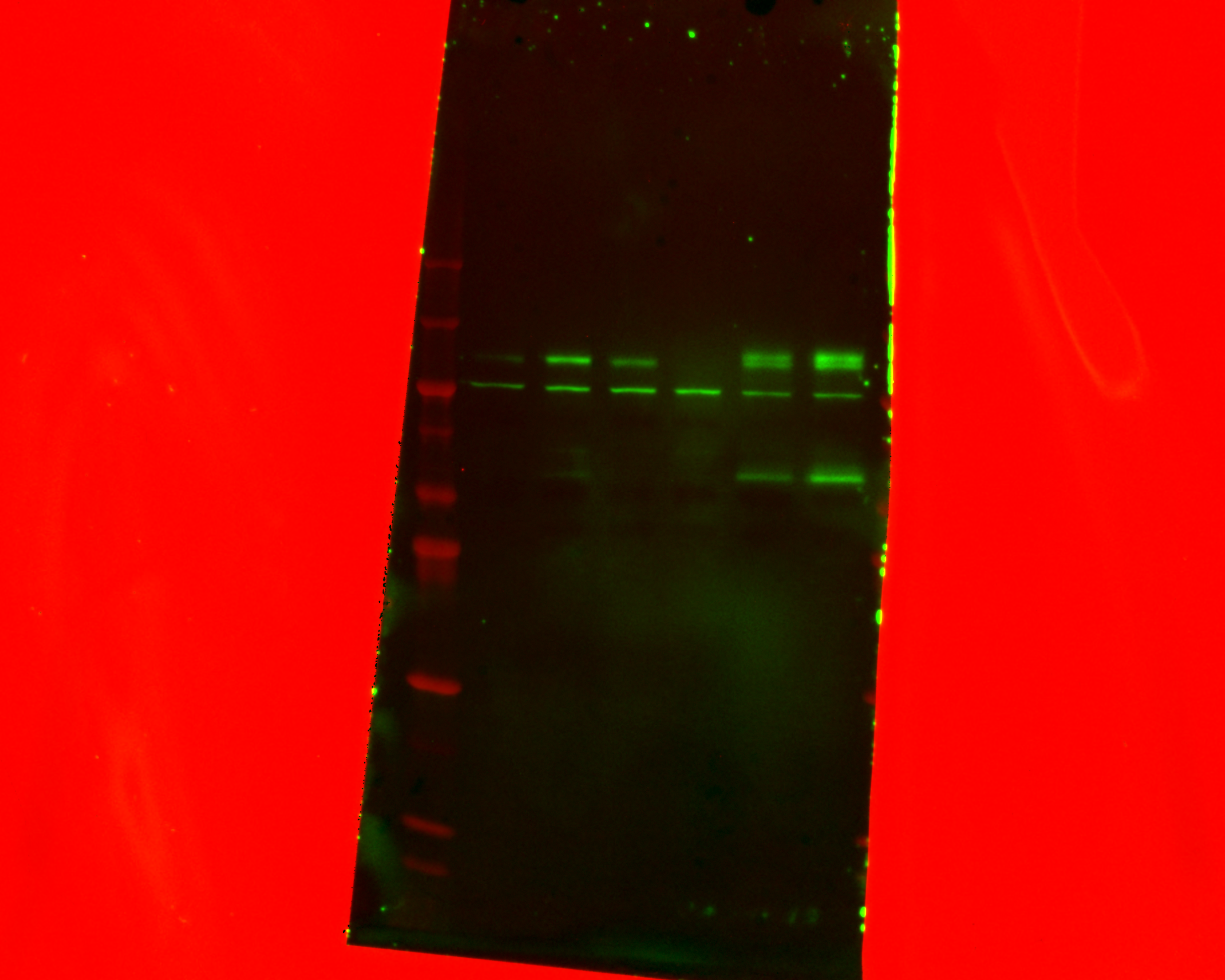

### 20240118_clonal_ut7_cd131_04 (Multichannel)_greyscale_edit2.tif

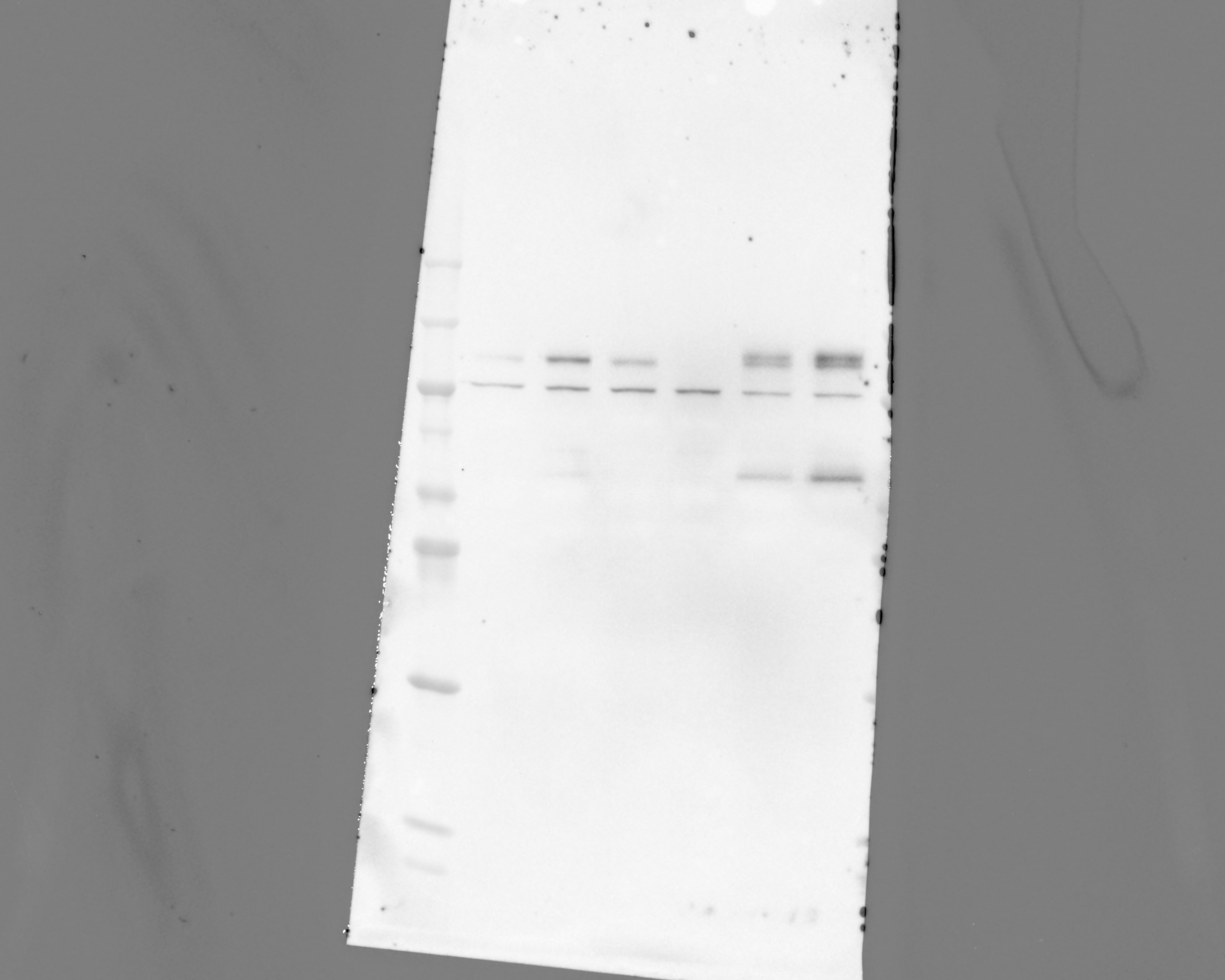

### 20240118_clonal_ut7_cd131_04(Chemiluminescence).tif

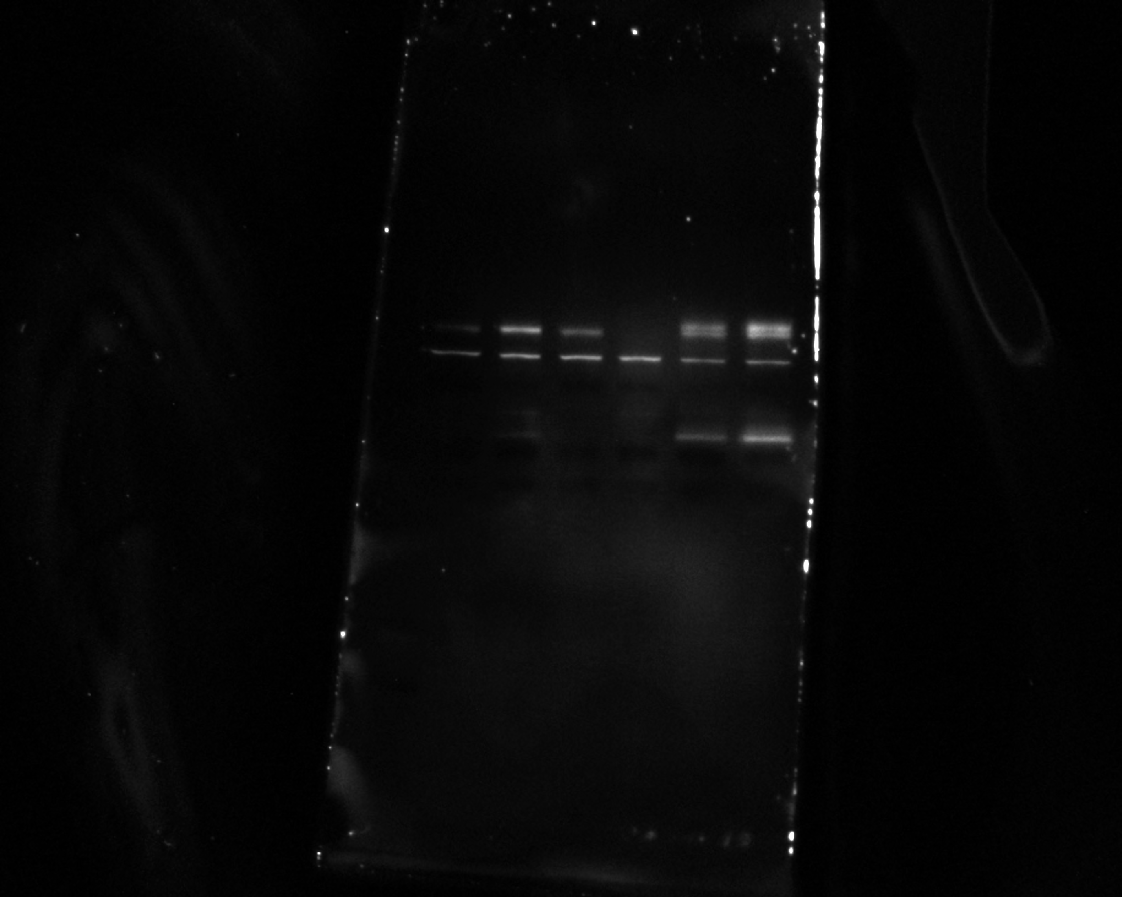

### 20240118_clonal_ut7_cd131_04(Colorimetric).tif

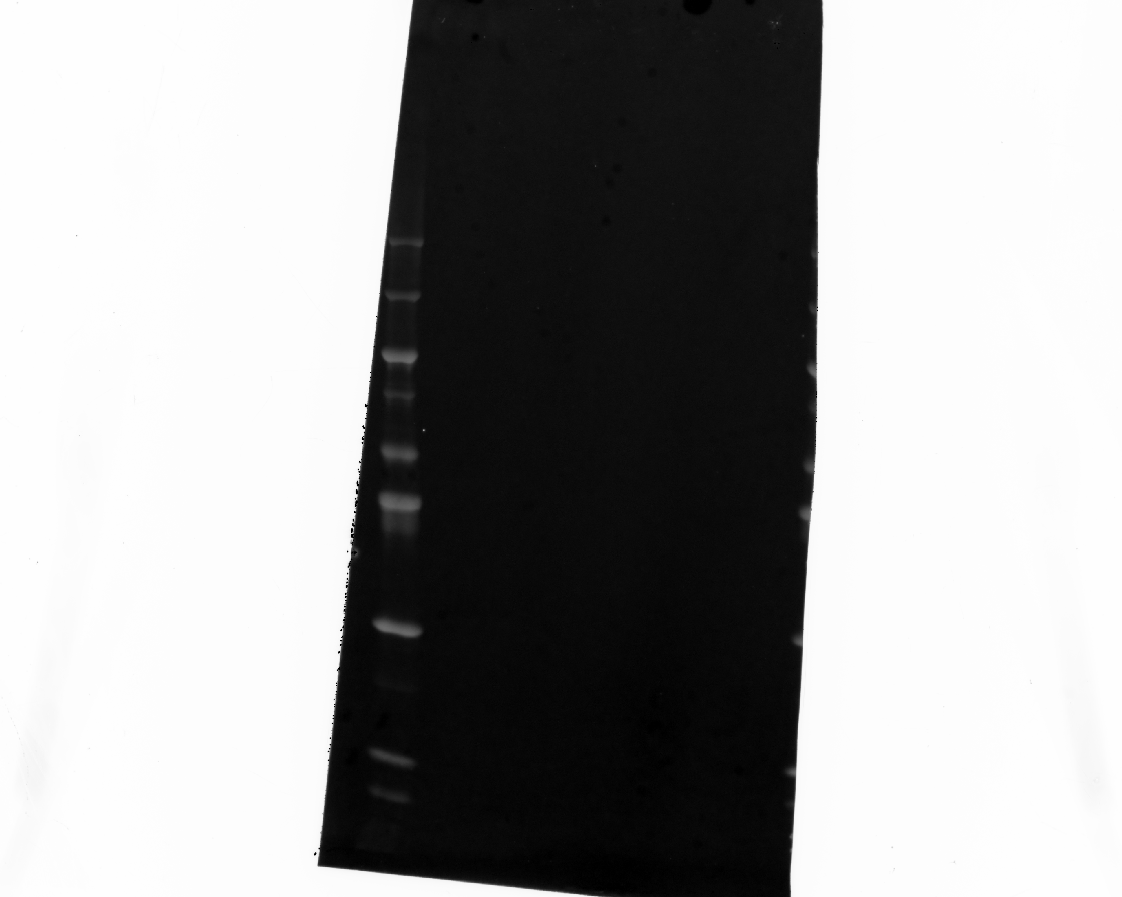

### 20240625_hek_phosphogel_4_rerun_gel_1_hh3_5(Chemiluminescence).tif

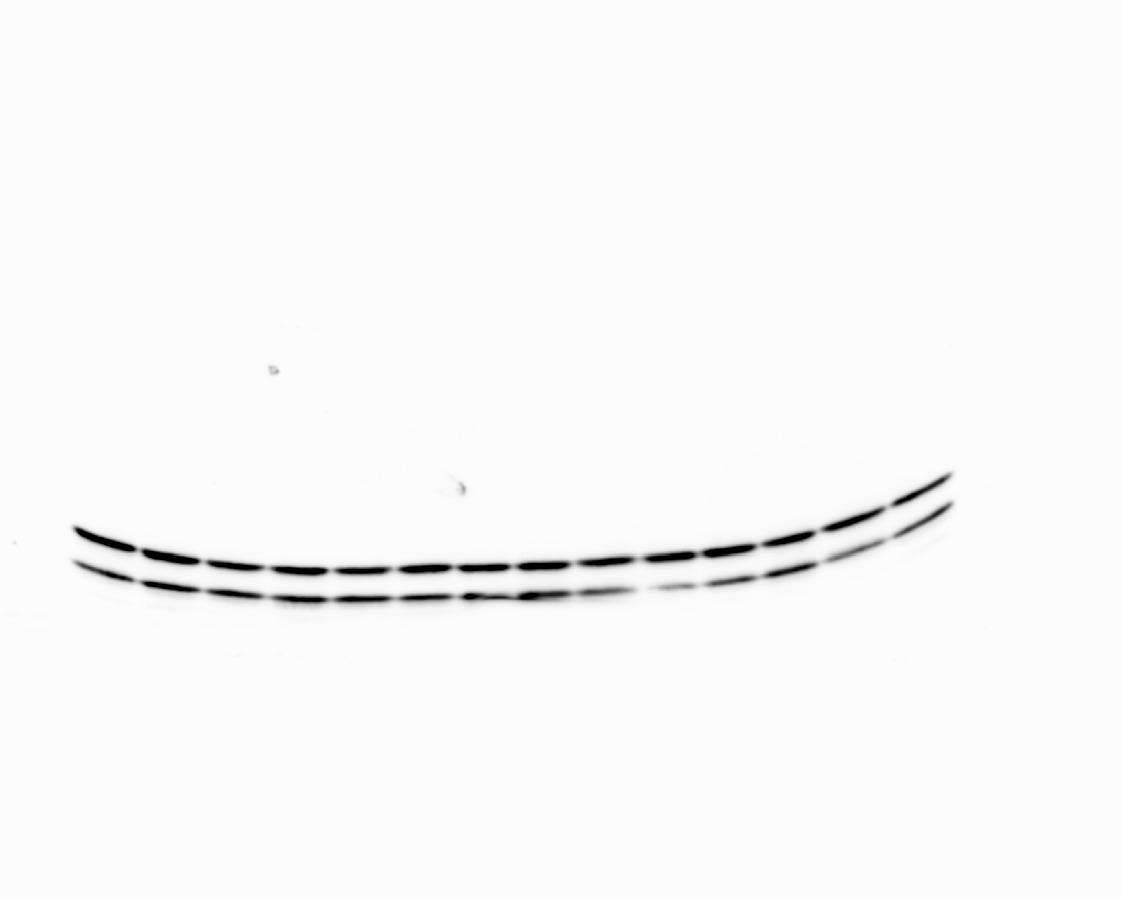

### 20240625_hek_phosphogel_4_rerun_gel_1_hh3_5(Colorimetric).tif

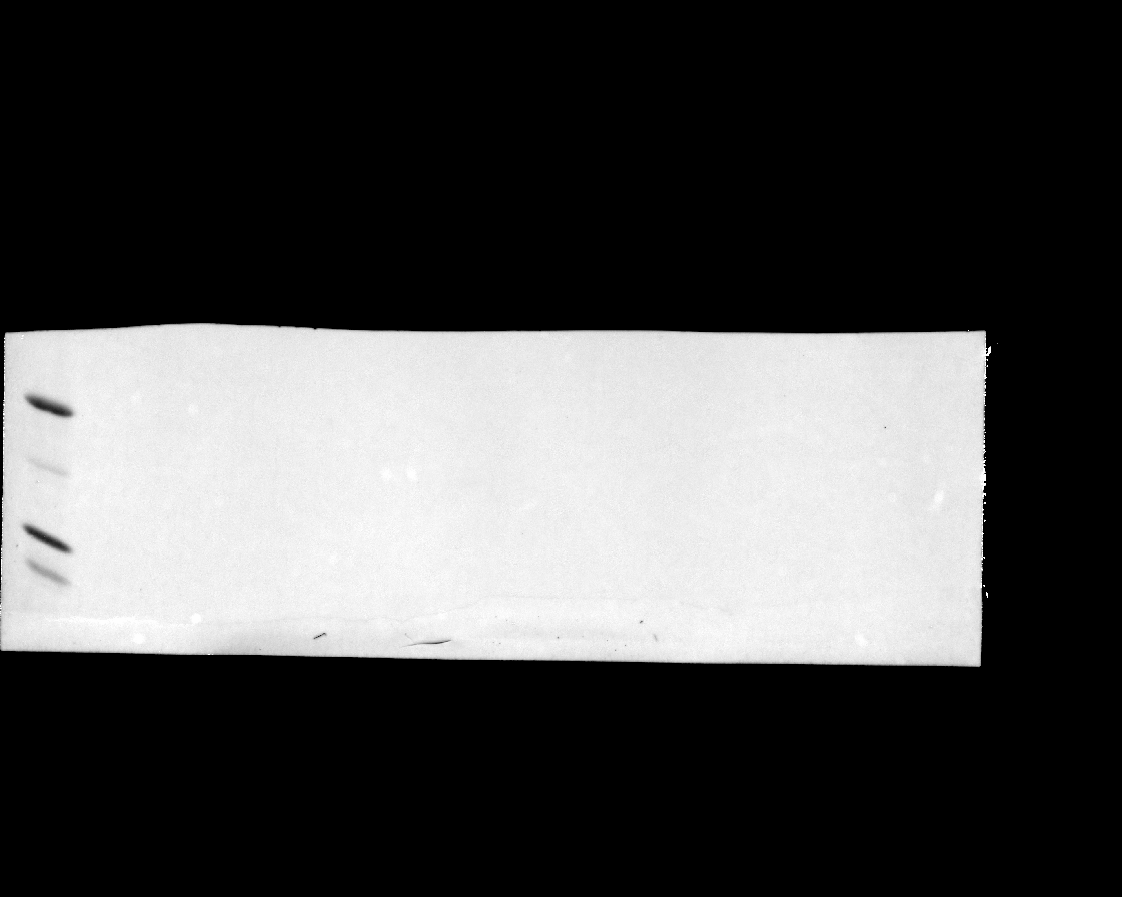

### 20240625_hek_phosphogel_4_rerun_gel_1_hh3_5(Composite).tif

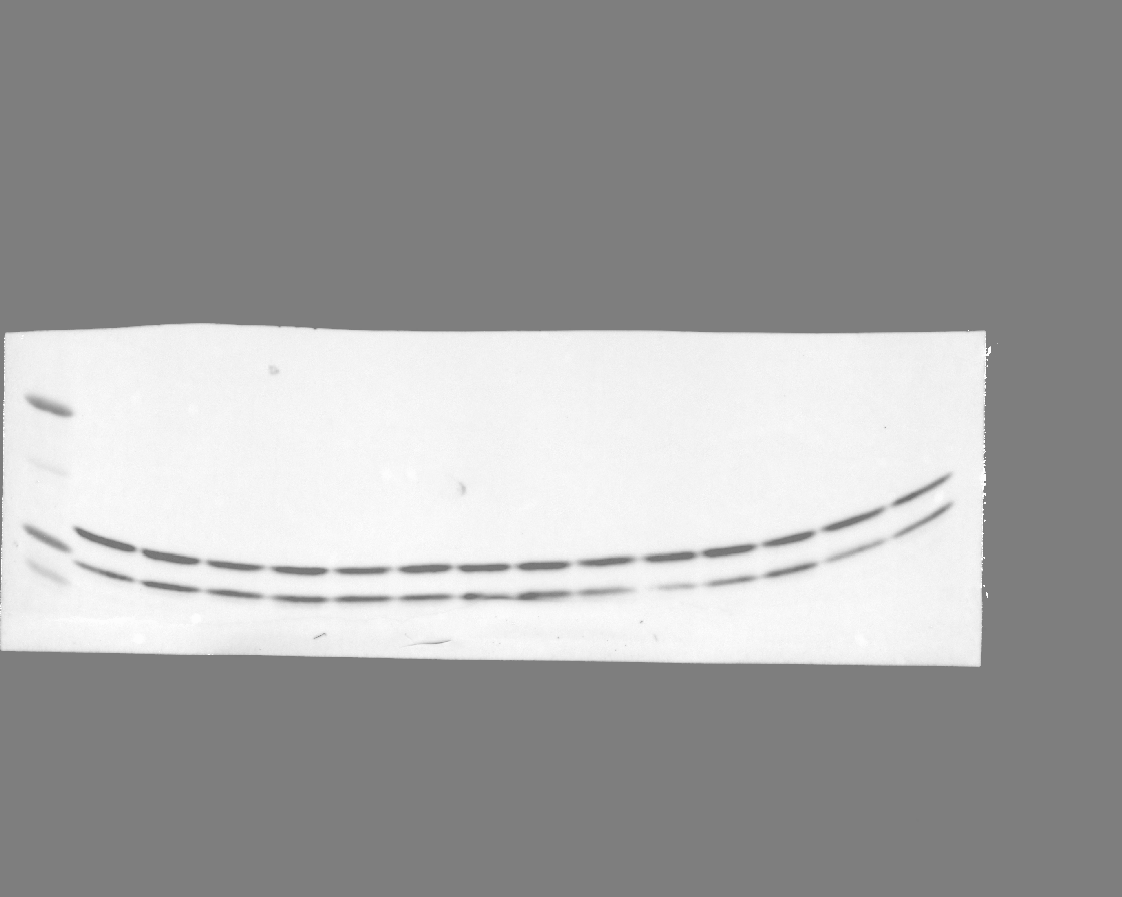

### 20240625_hek_phosphogel_4_rerun_gel_1_stat5_phospho_17(Chemiluminescence).tif

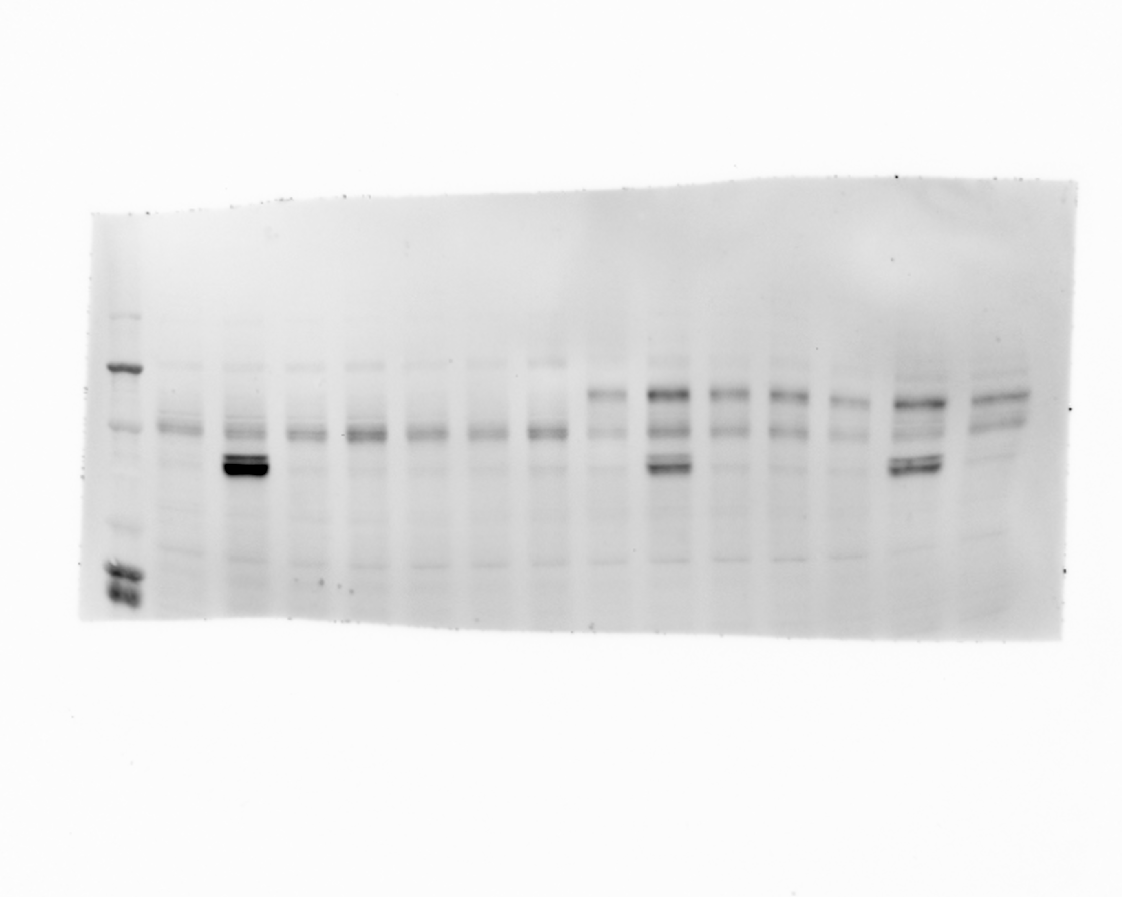

### 20240625_hek_phosphogel_4_rerun_gel_1_stat5_phospho_17(Colorimetric).tif

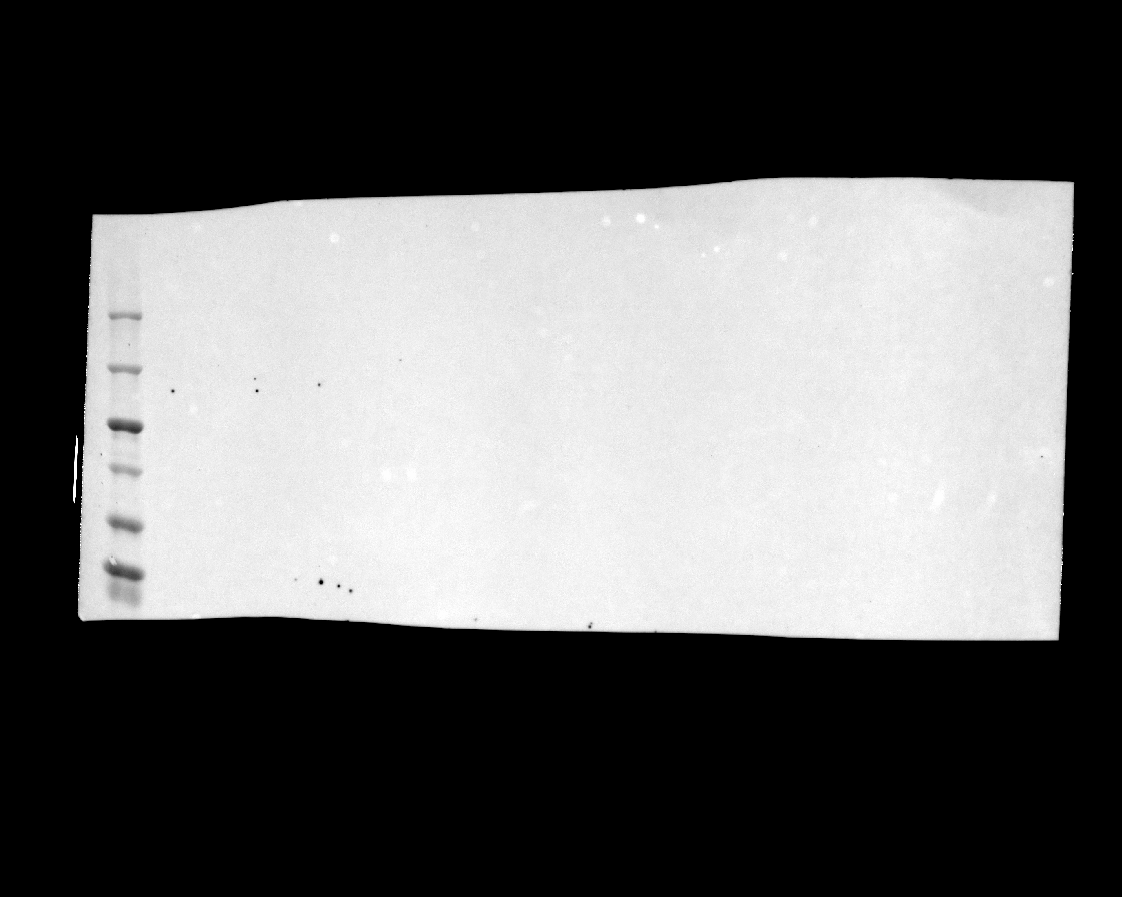

### 20240625_hek_phosphogel_4_rerun_gel_1_stat5_phospho_17(Composite).tif

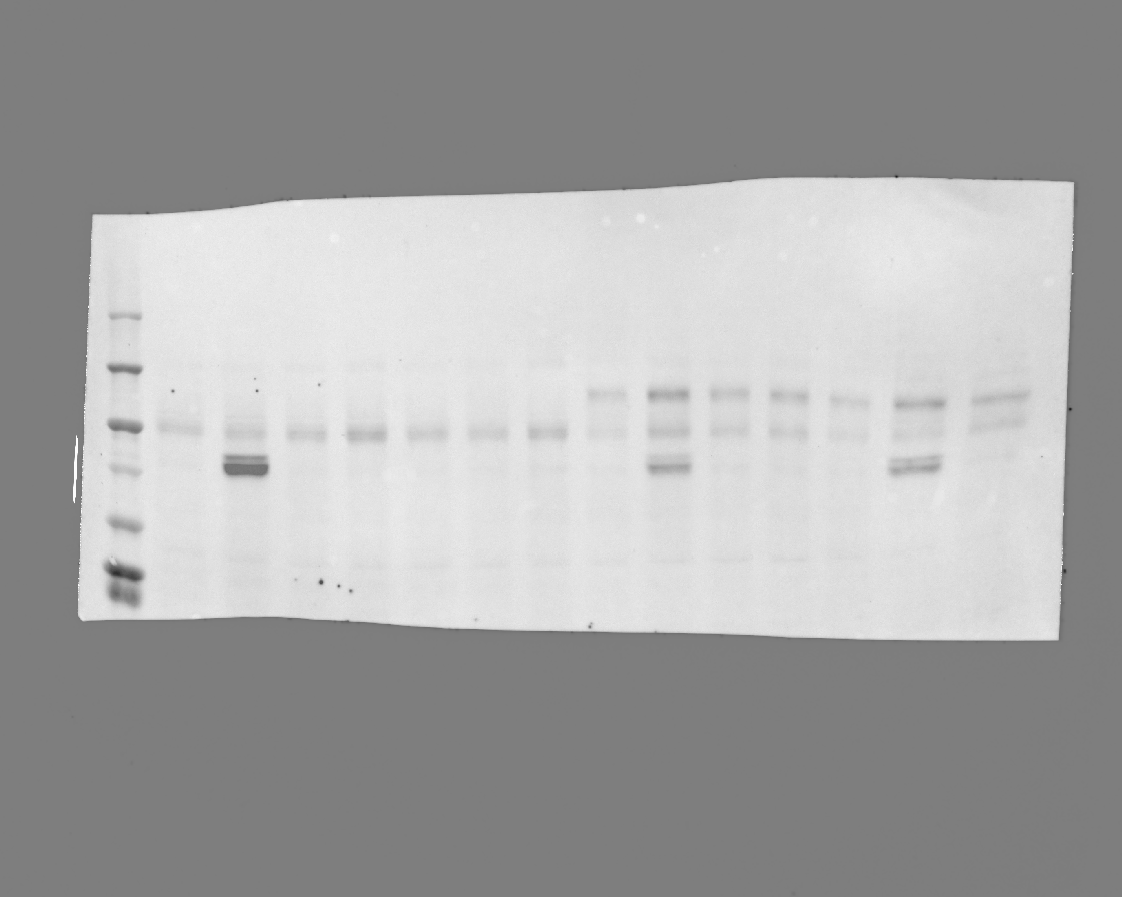

### 20240625_hek_phosphogel_4_rerun_gel_1_stat5_phospho_17.tif

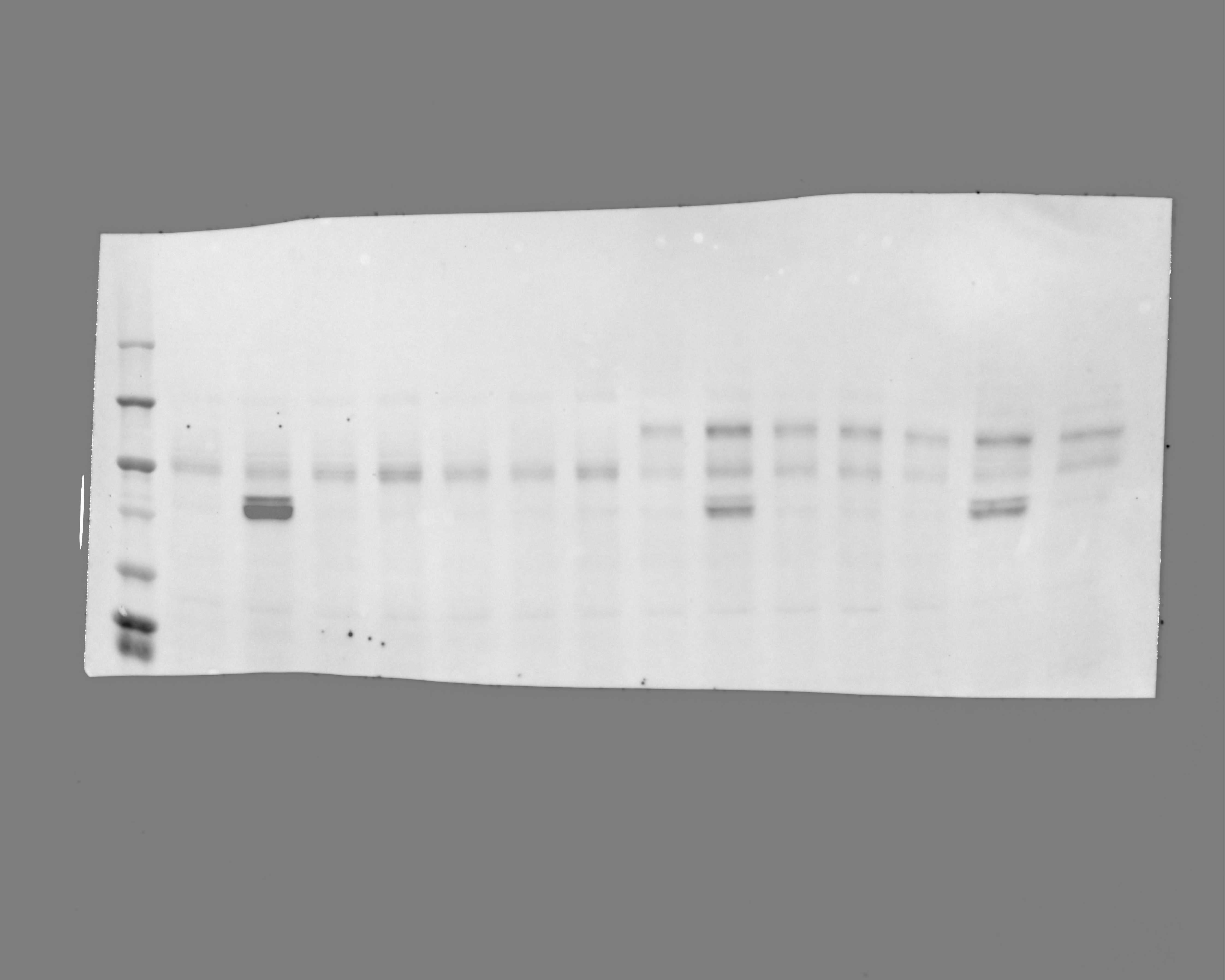

### 20240625_hek_phosphogel_4_rerun_gel_2_stat5_3 (Multichannel).tif

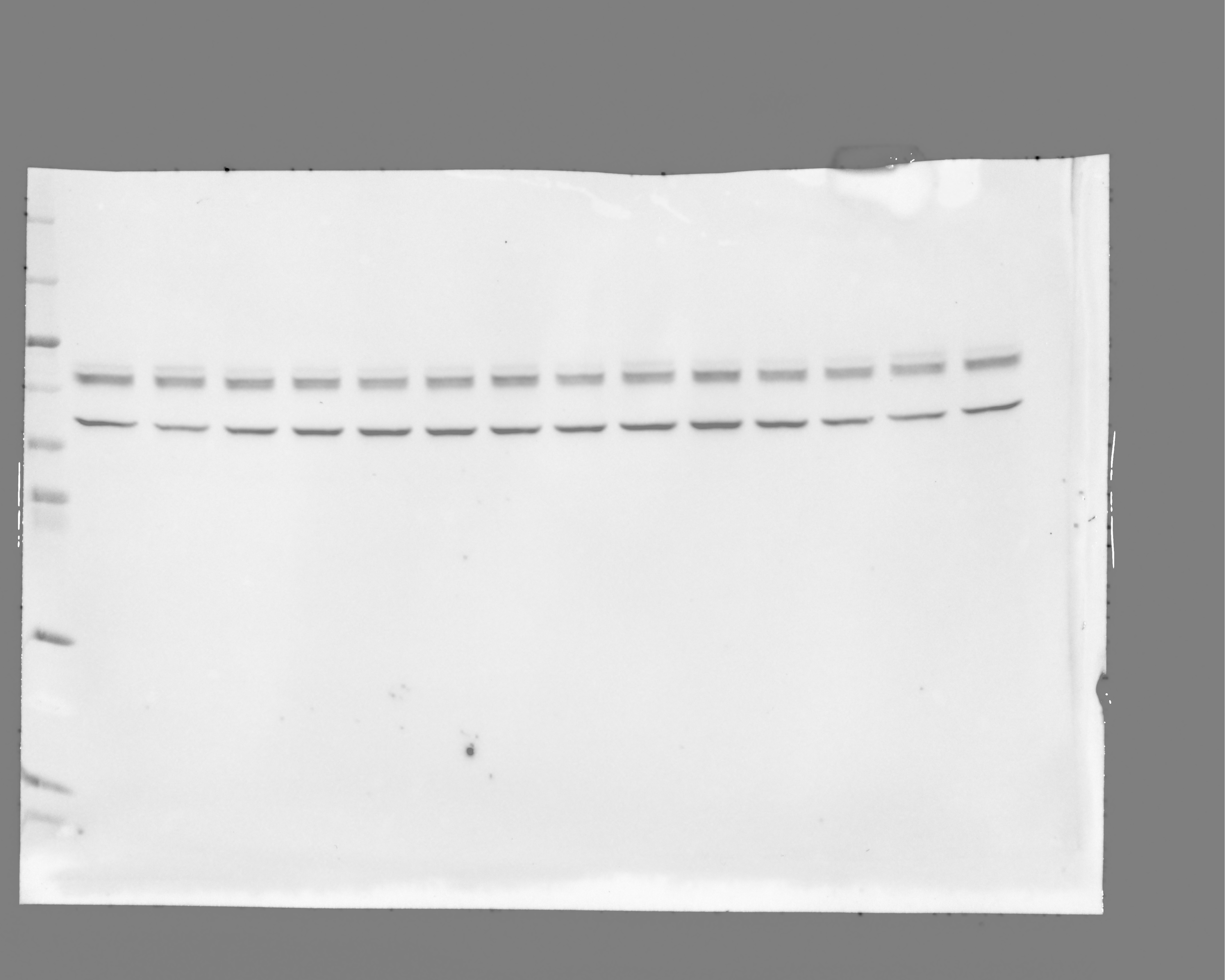

### 20240625_hek_phosphogel_4_rerun_gel_2_stat5_3(Chemiluminescence).tif

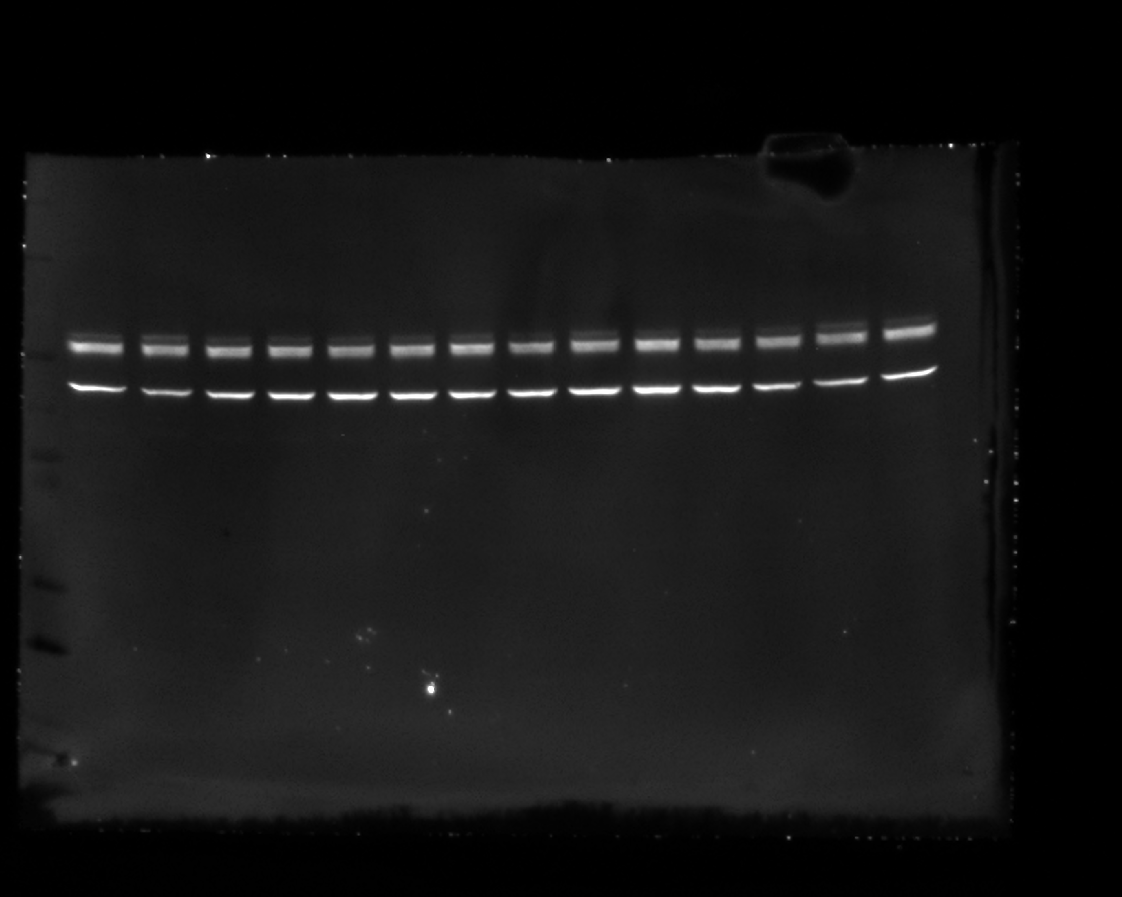

### 20240625_hek_phosphogel_4_rerun_gel_2_stat5_3(Colorimetric).tif

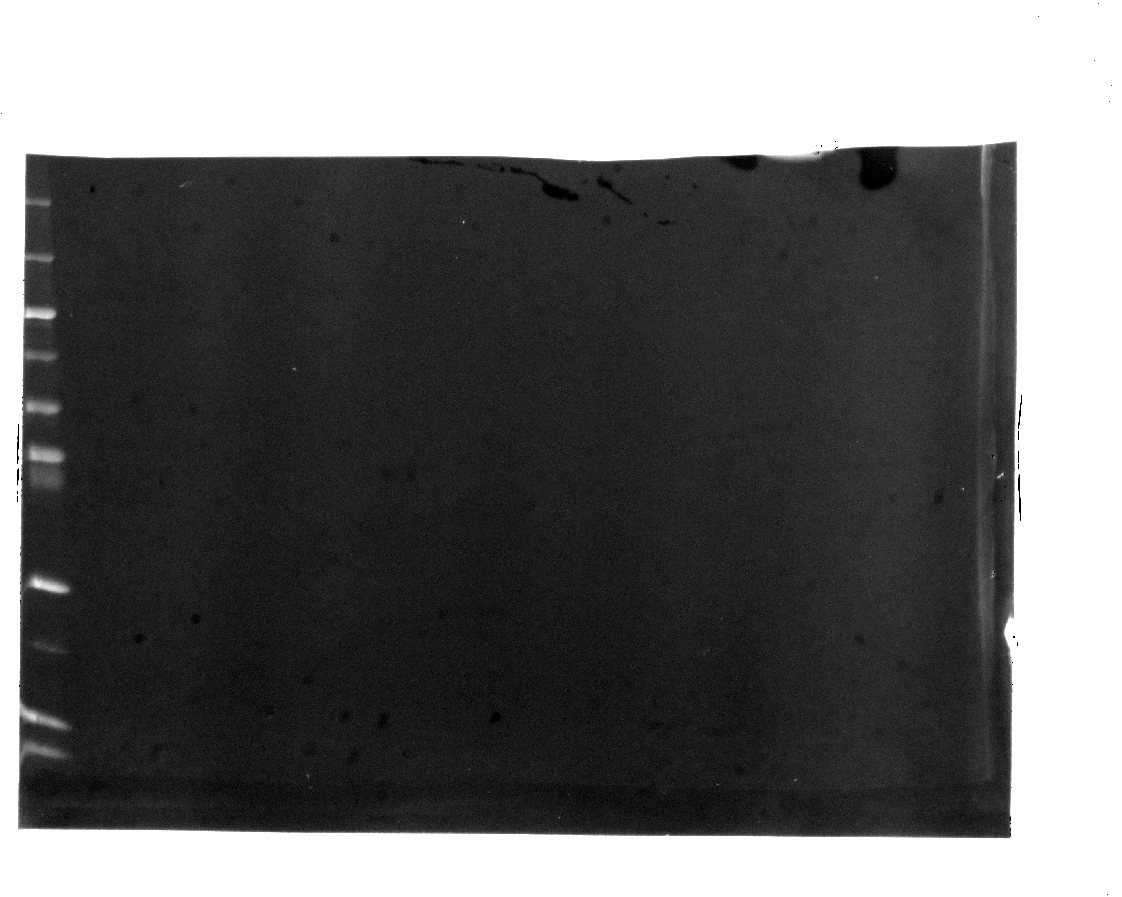

### 20240625_hek_phosphogel_4_rerun_gel_2_stat5_3(Composite).tif

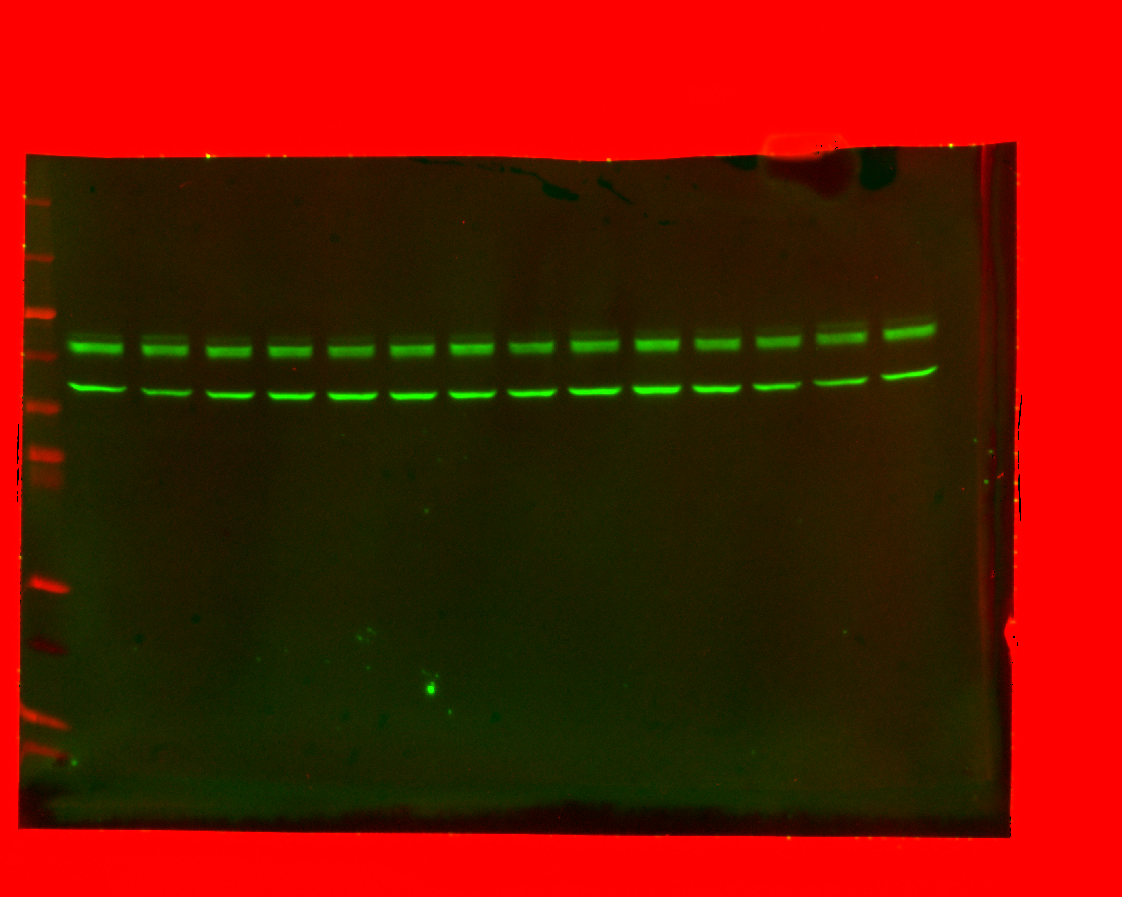

### 20240625_hek_phosphogel_4_rerun_gel_3_hh3_1_7(Chemiluminescence).tif

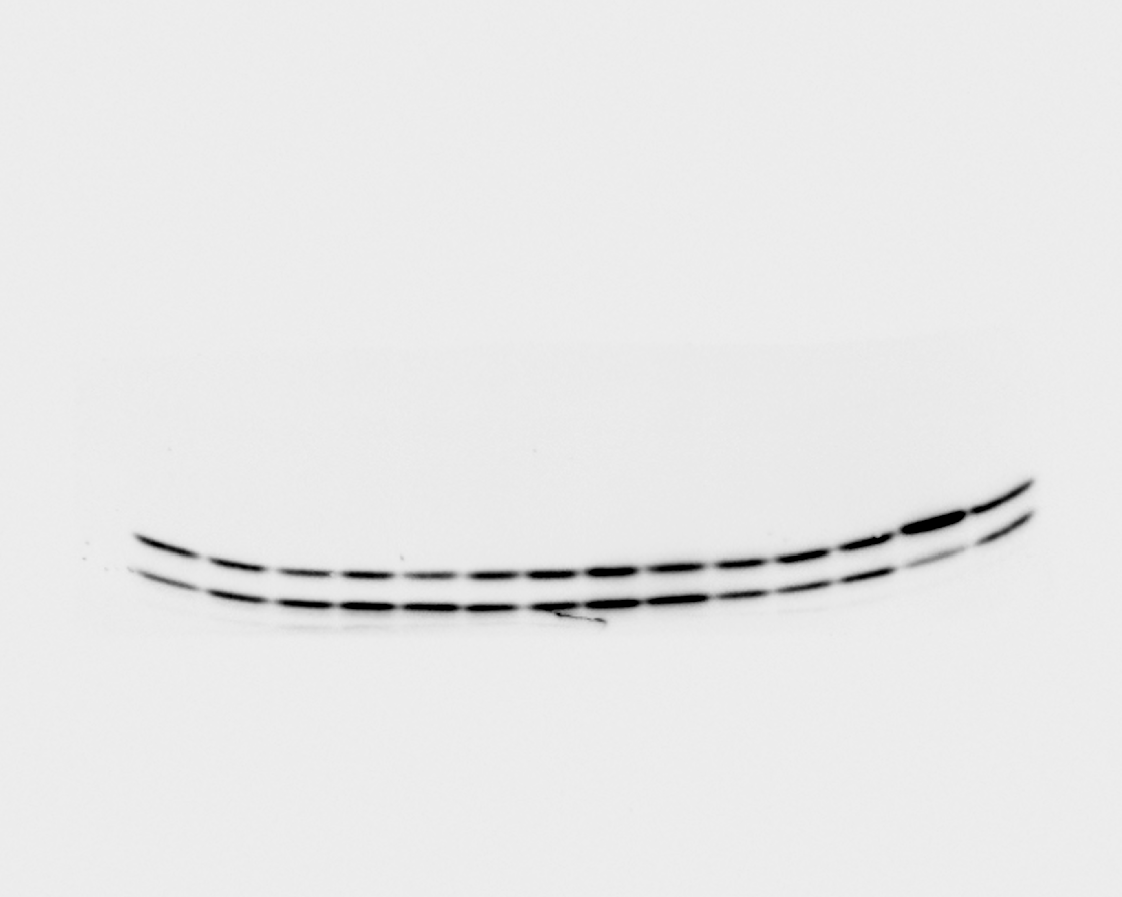

### 20240625_hek_phosphogel_4_rerun_gel_3_hh3_1_7(Colorimetric).tif

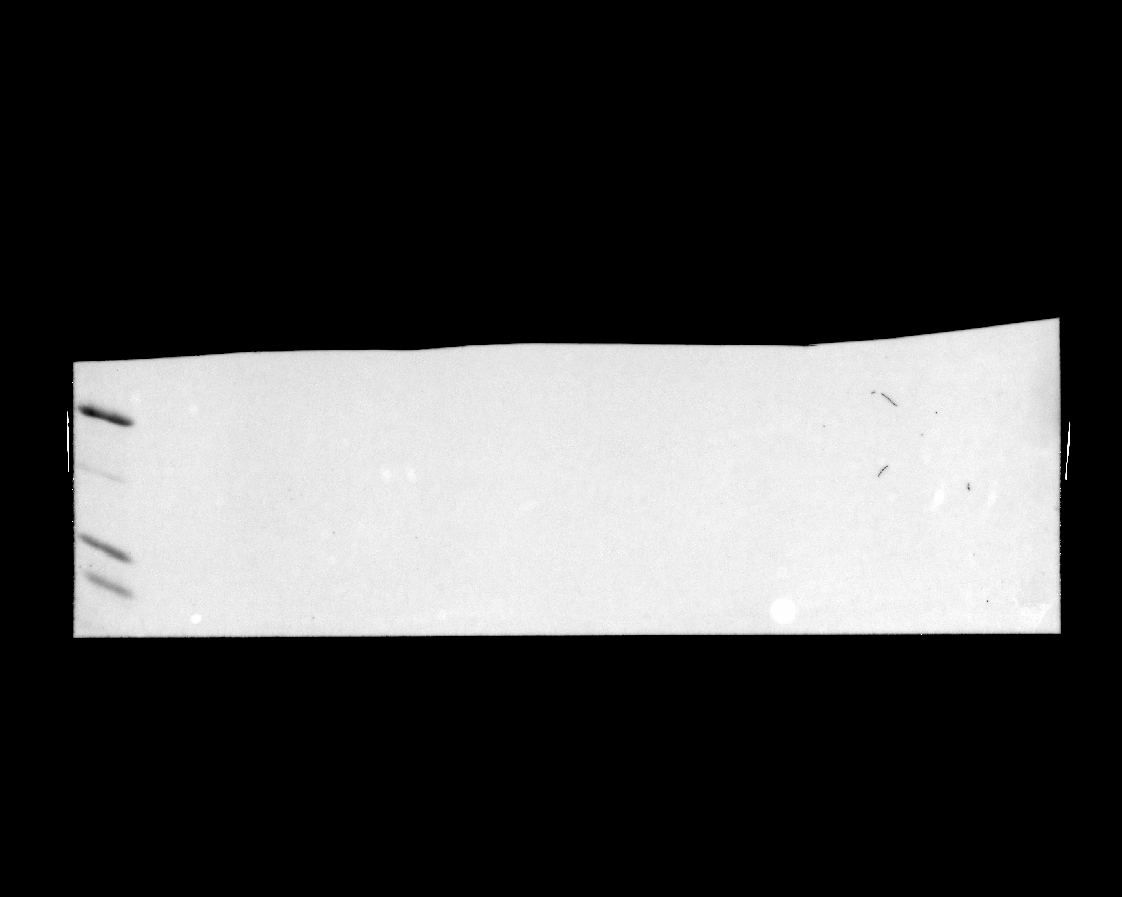

### 20240625_hek_phosphogel_4_rerun_gel_3_hh3_1_7(Composite).tif

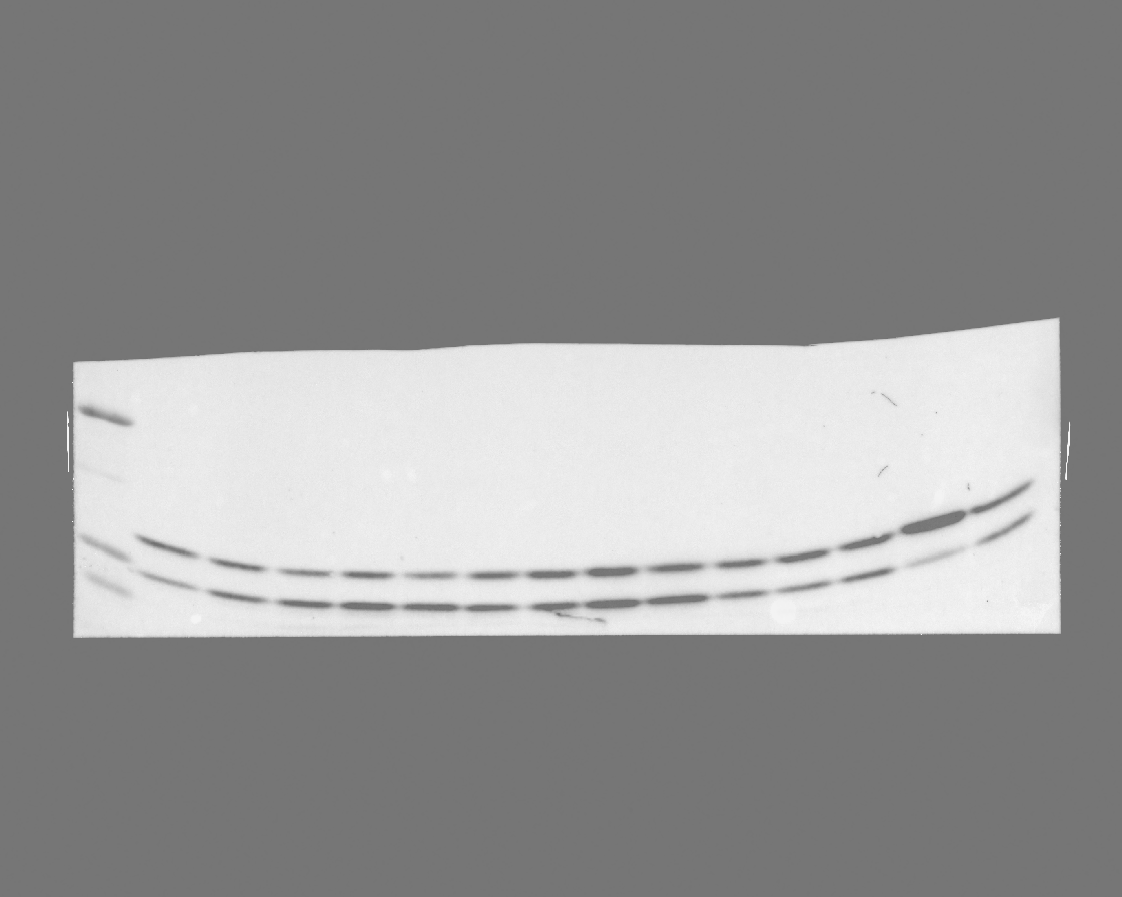

### 20240625_hek_phosphogel_4_rerun_gel_4_reprobe_v5_7(Colorimetric).tif

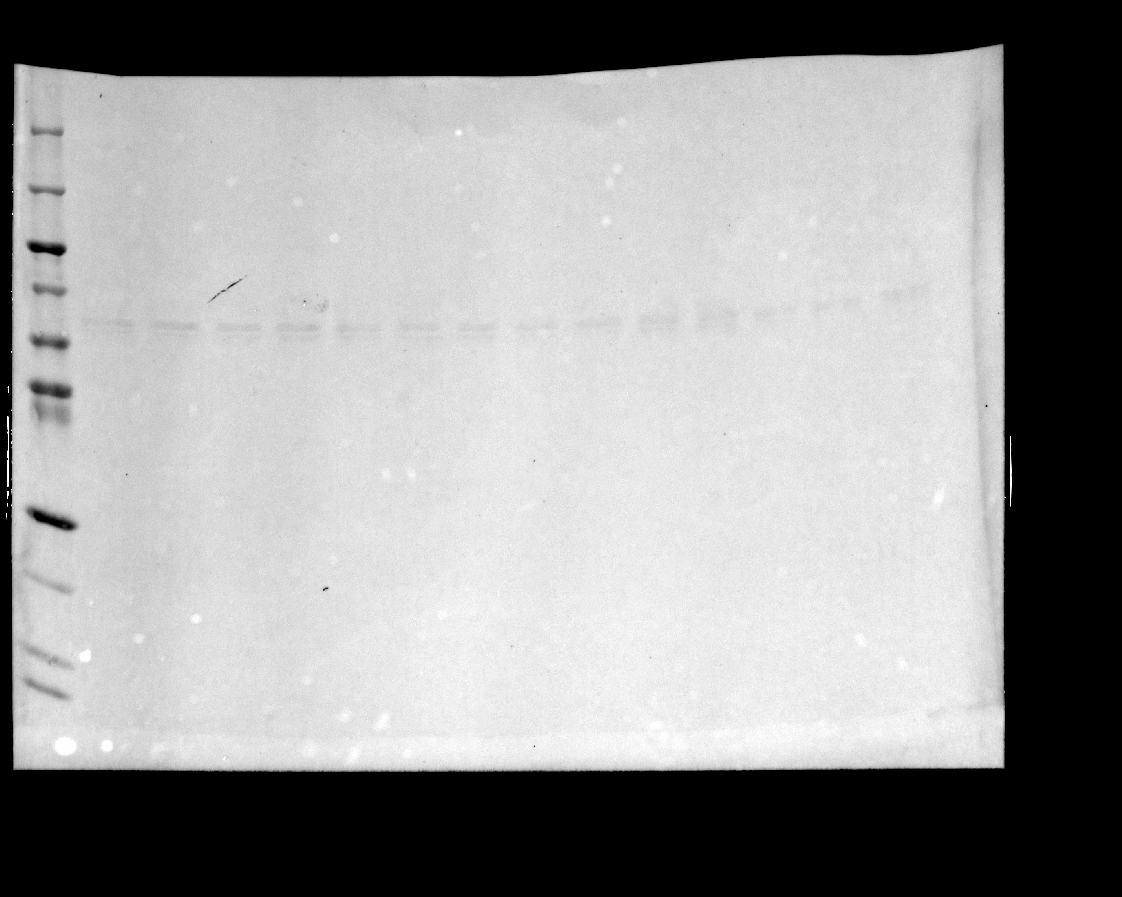

### 20240625_hek_phosphogel_4_rerun_gel_4_reprobe_v5_7(Composite).tif

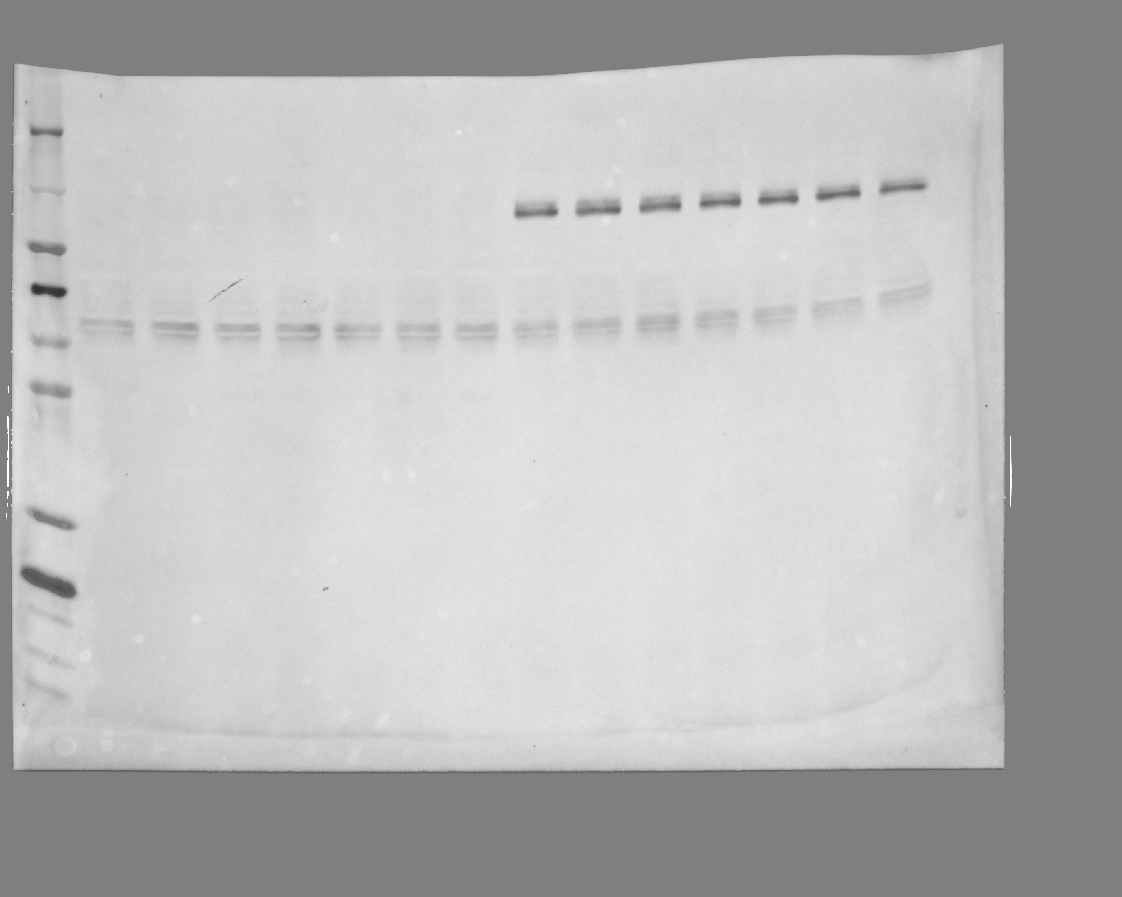

### 20240625_hek_phosphogel_4_rerun_gel_4_reprobe_v5_7(DyLight 488).tif

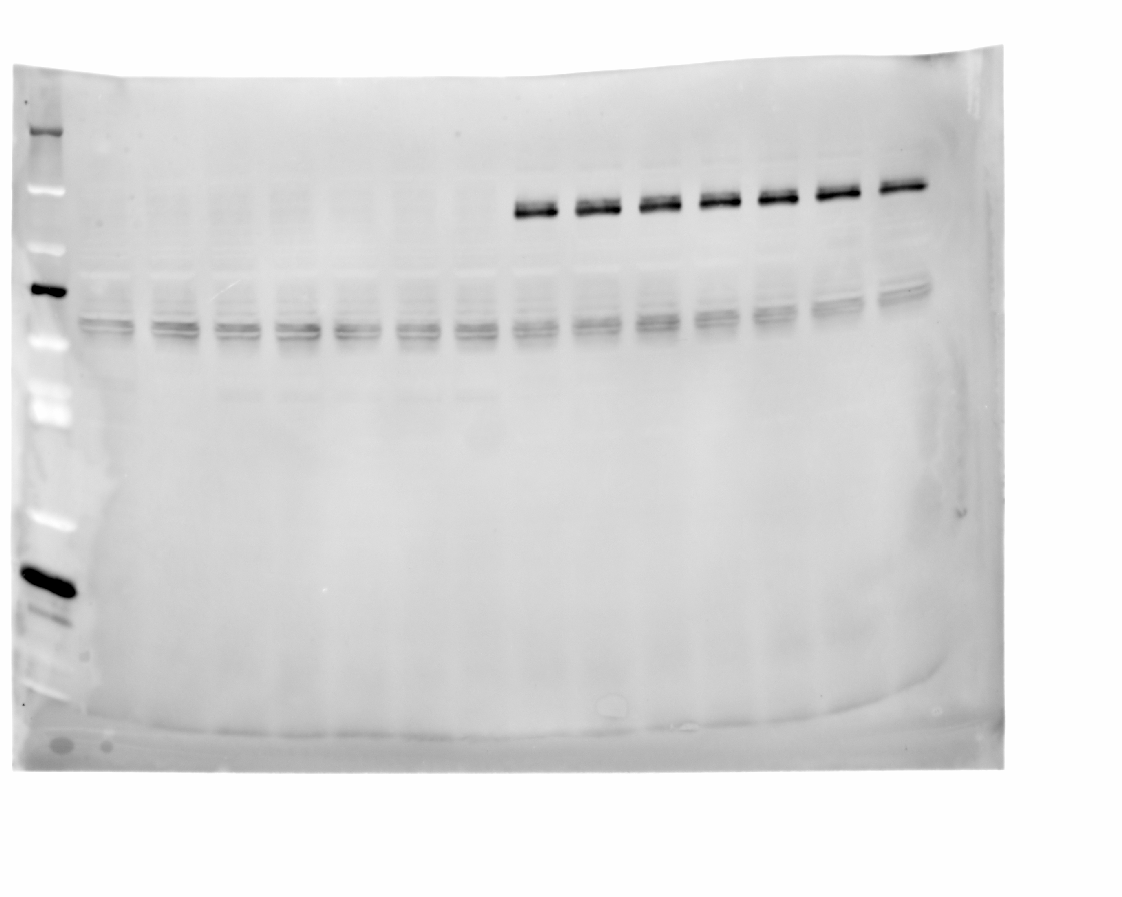

### 20241106_gel1_rerun_coomassie_overnight_destain(Coomassie Blue).tif

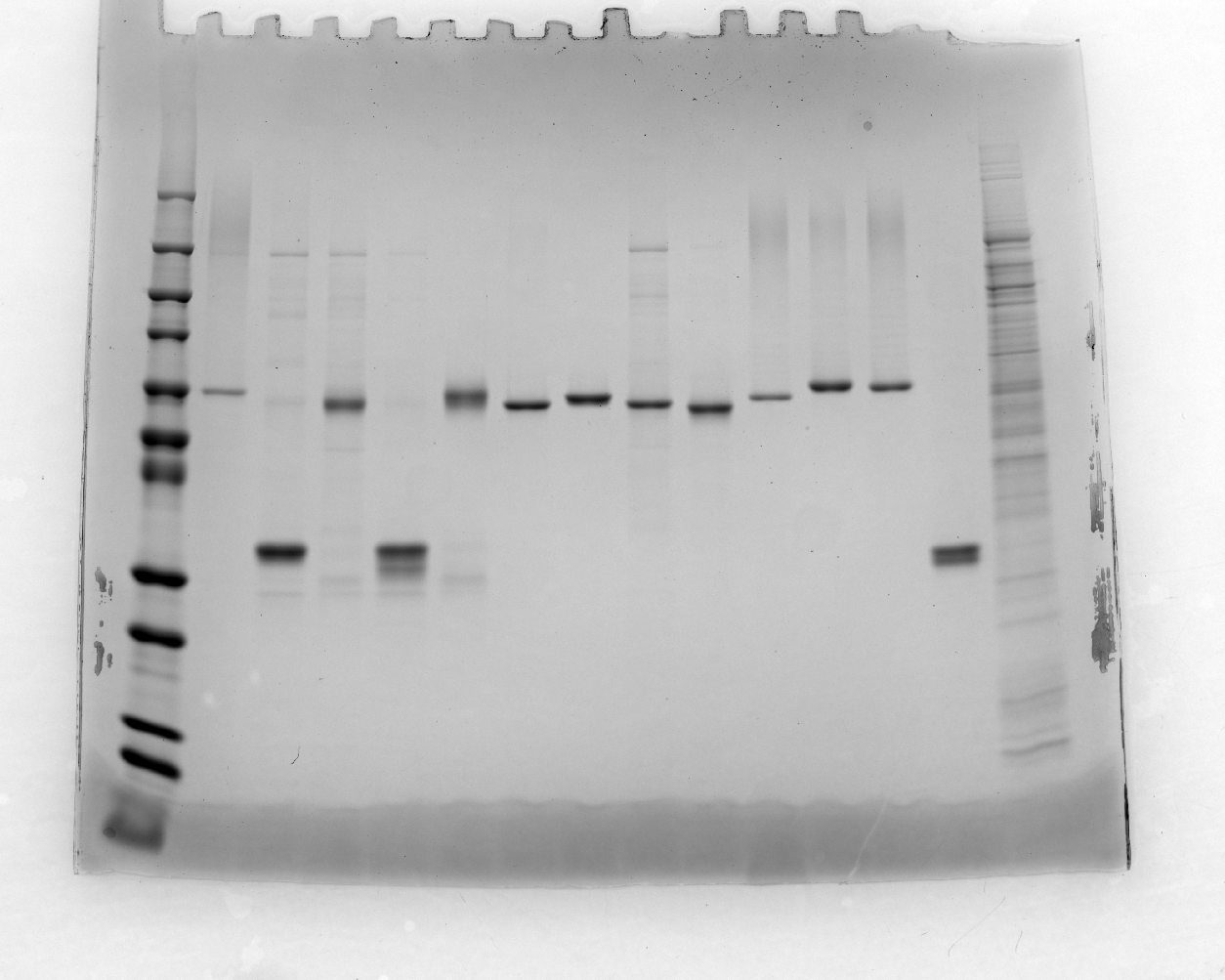

### 20241106_gel1_rerun_coomassie_overnight_destain_blue.tif

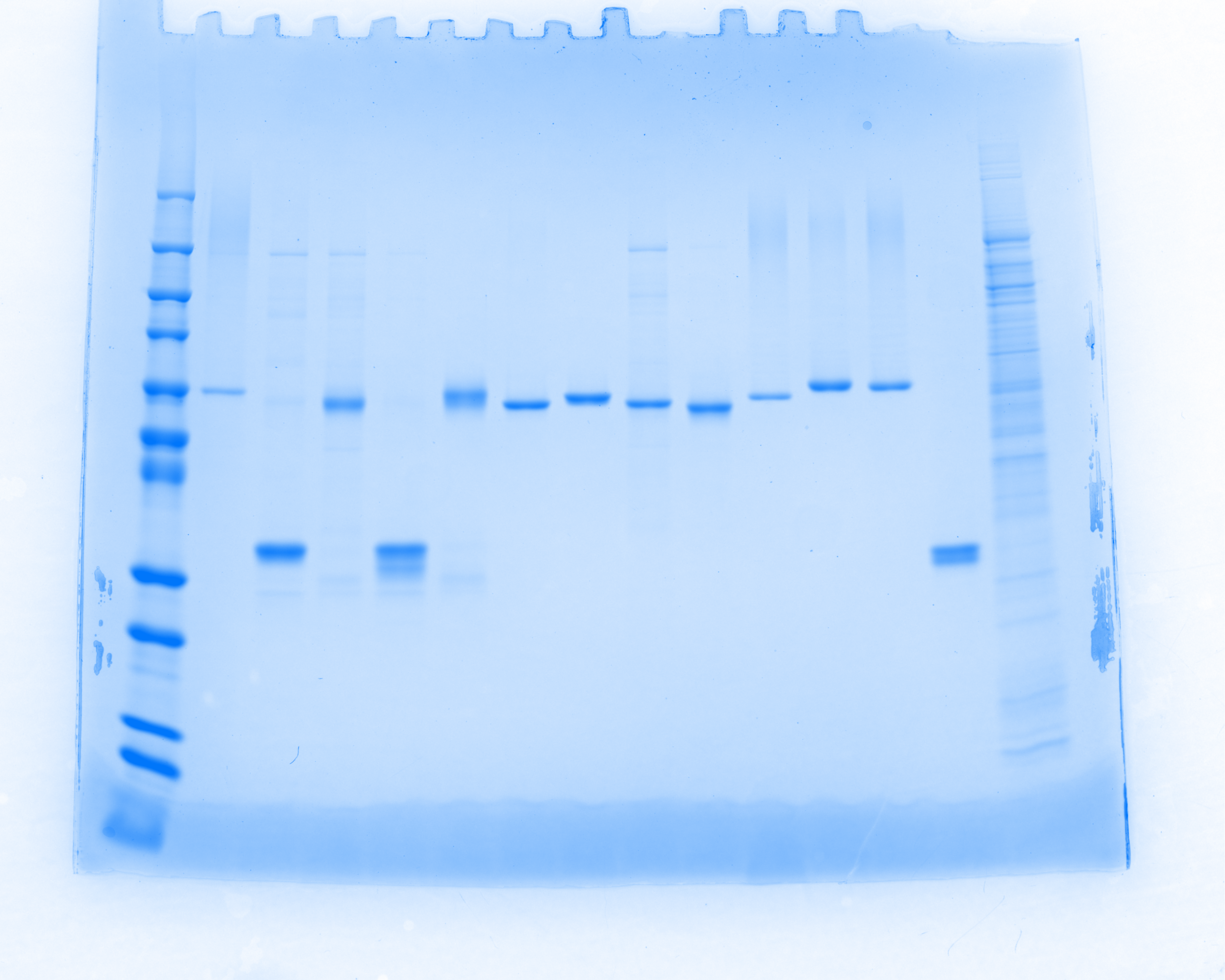

### 20241106_gel1_rerun_reprobe_his488_4 (Alexa 488)_edit.tif

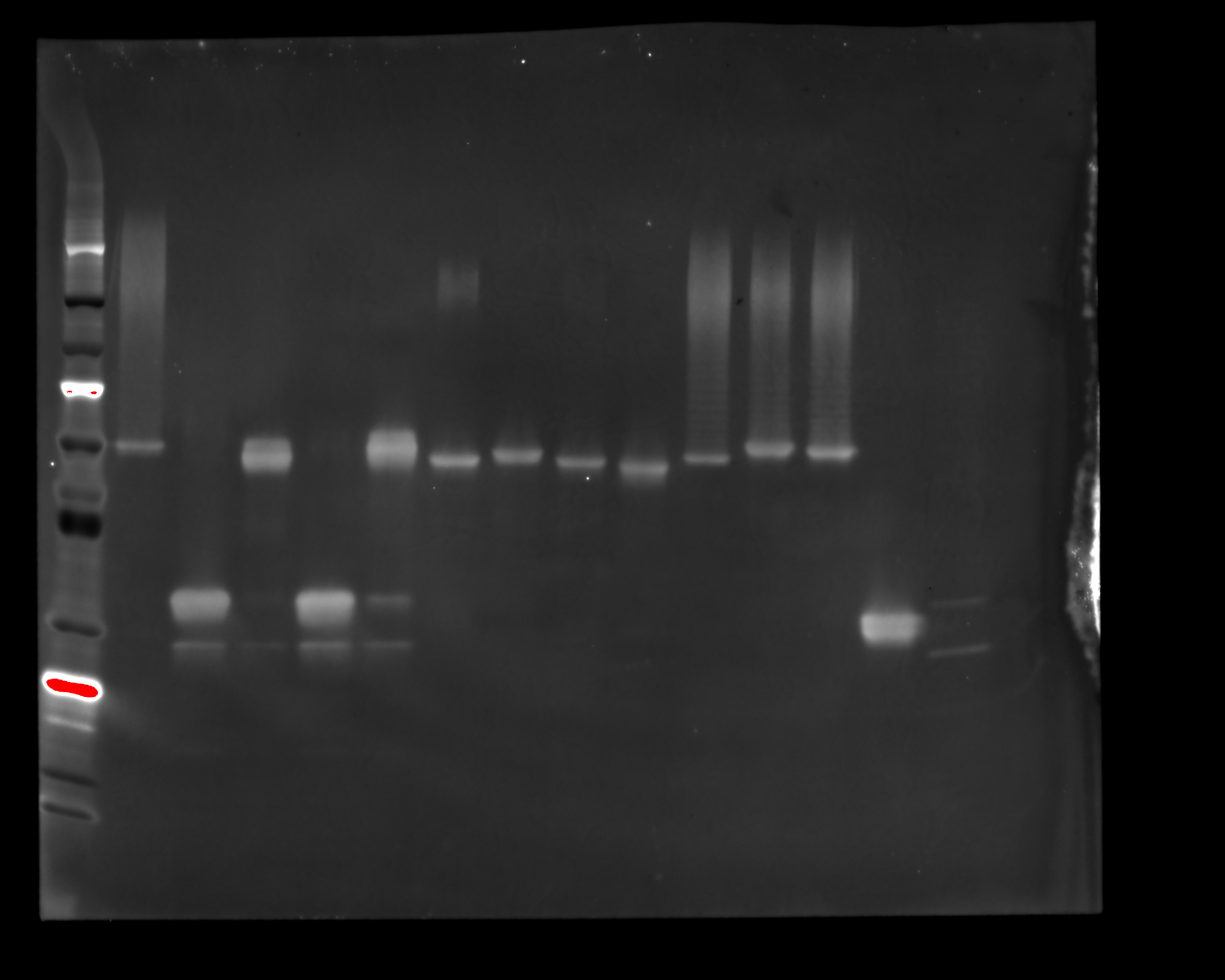

### 20241106_gel1_rerun_reprobe_his488_4 (Multichannel)_edit3.tif

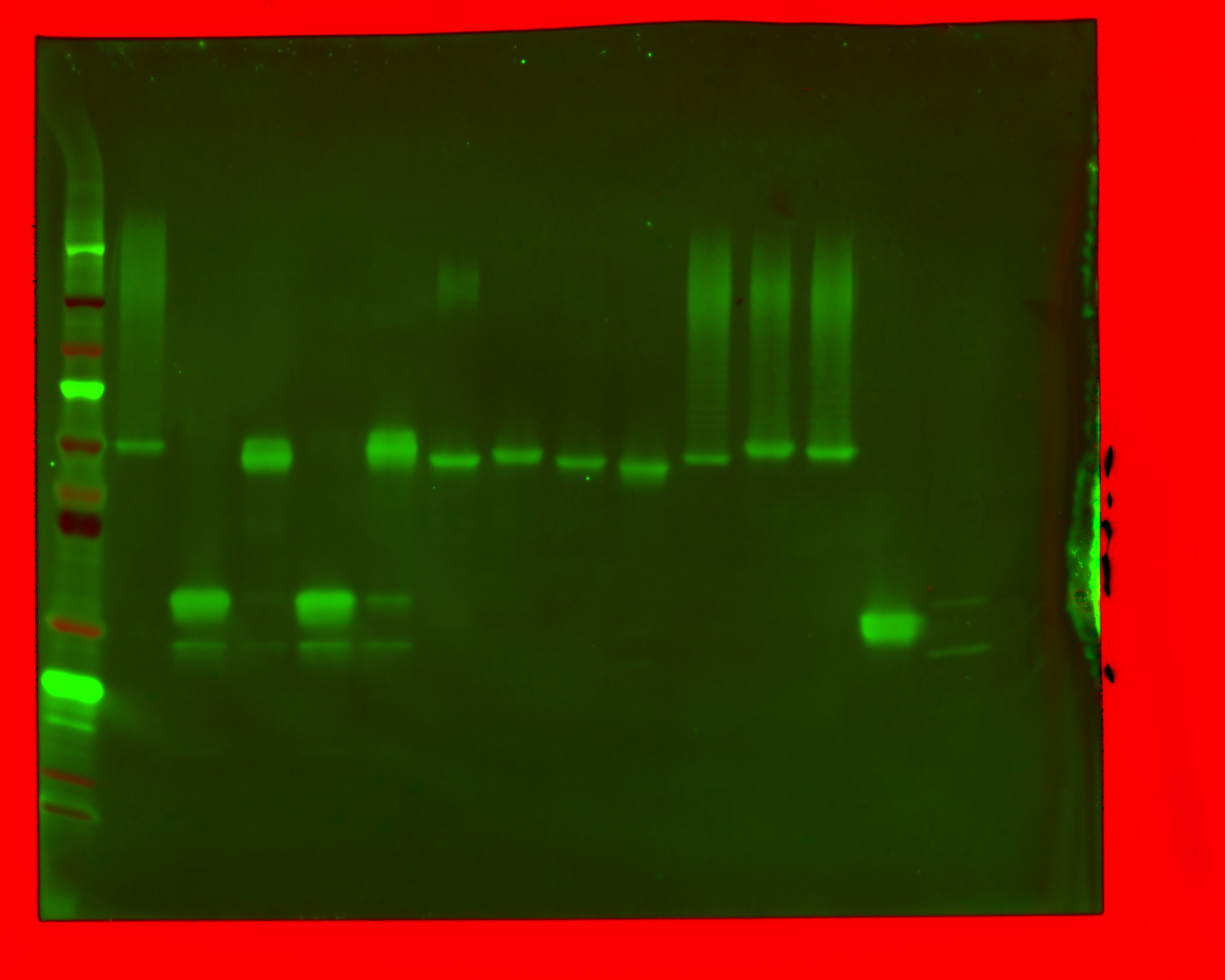

### 20241106_gel1_rerun_reprobe_his488_4 (Multichannel)_edit4.tif

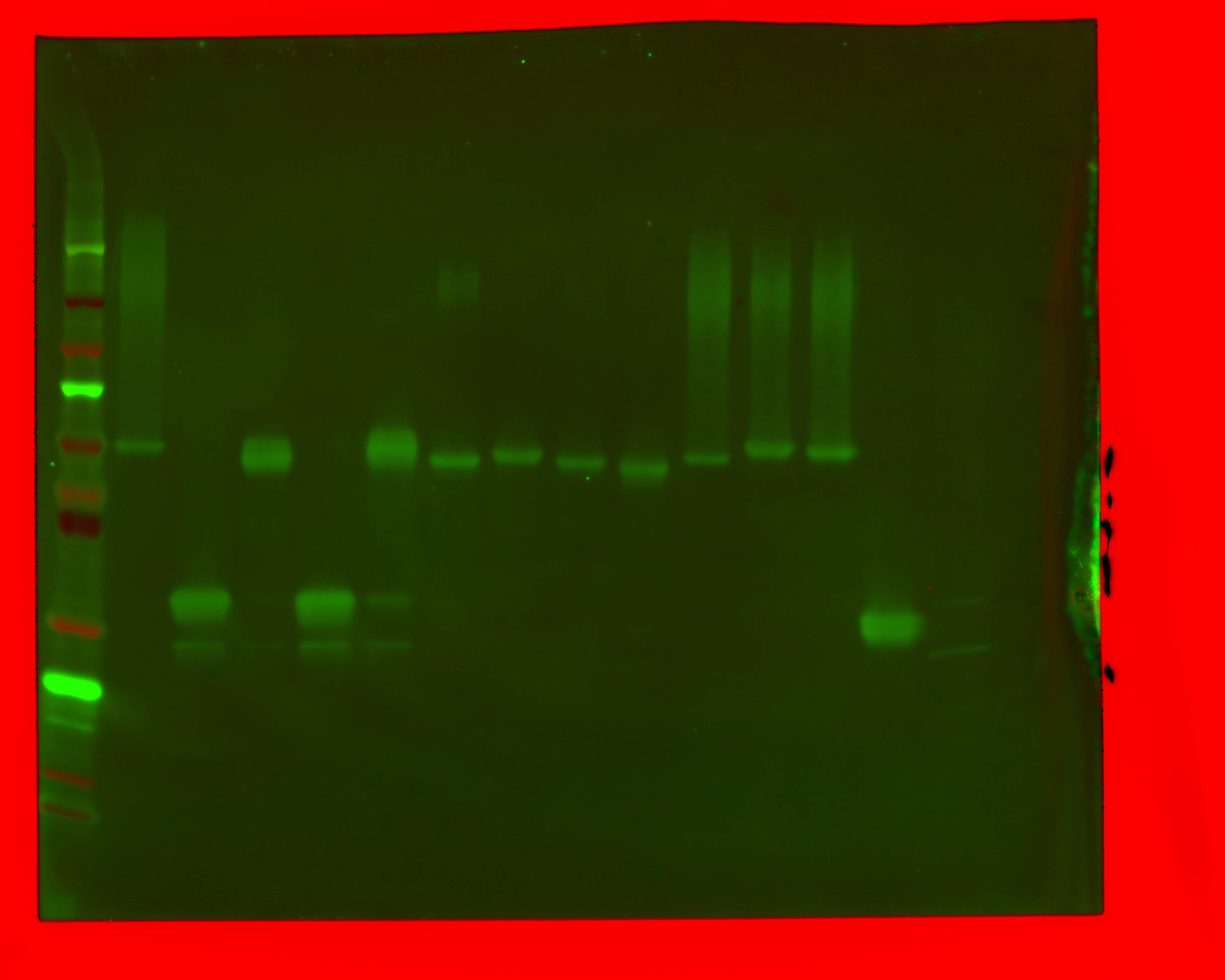

### 20241106_gel1_rerun_reprobe_his488_4 (Multichannel)_edit.tif

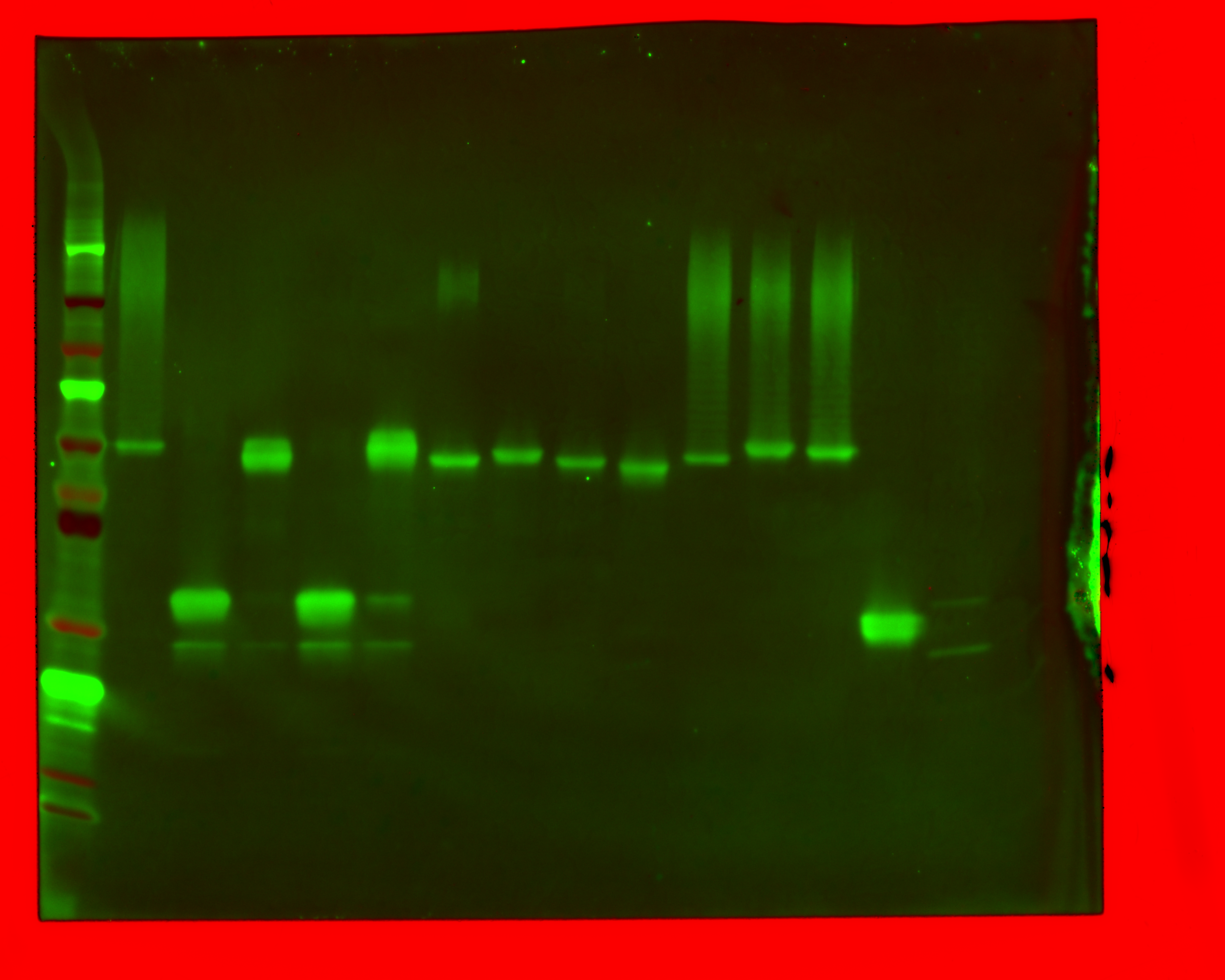

### 20241106_gel1_rerun_reprobe_his488_4(Composite).tif

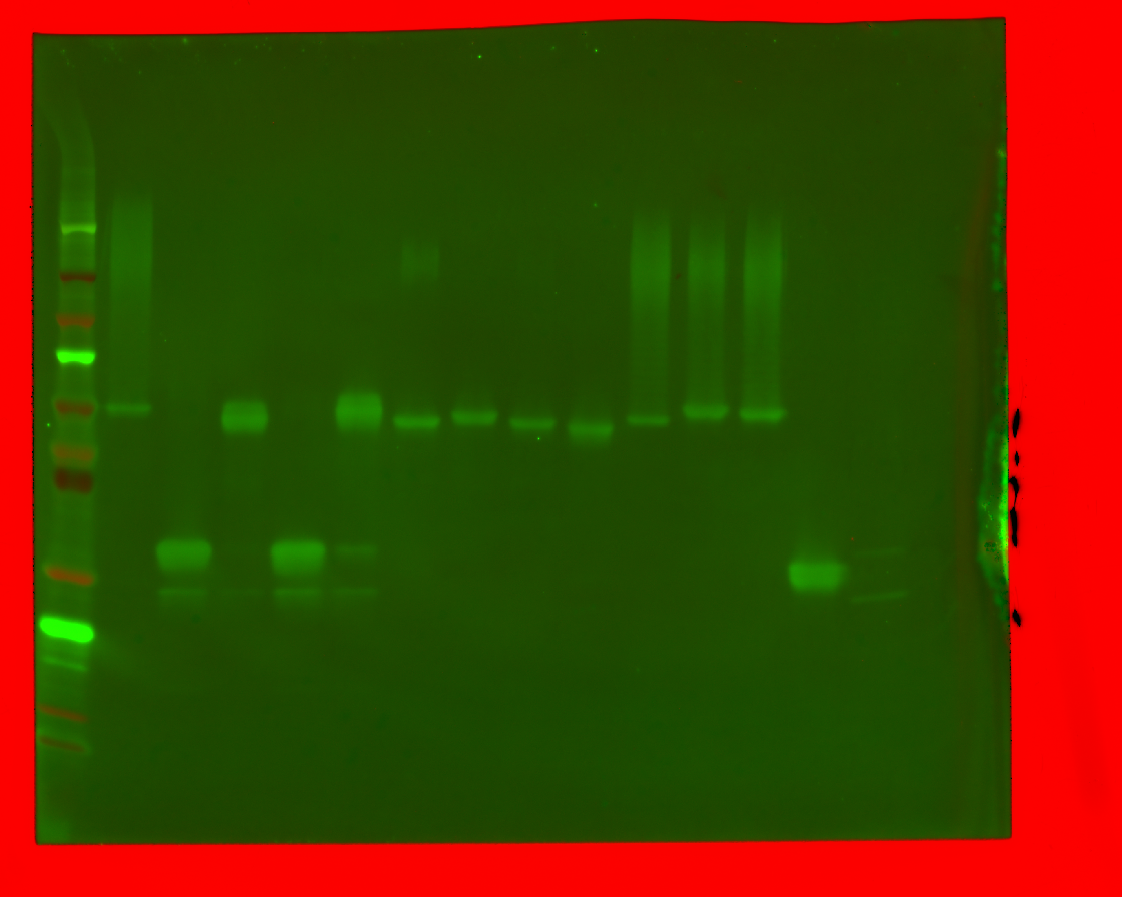

### 20241106_gel1_rerun_reprobe_his488_6(DyLight 488).tif

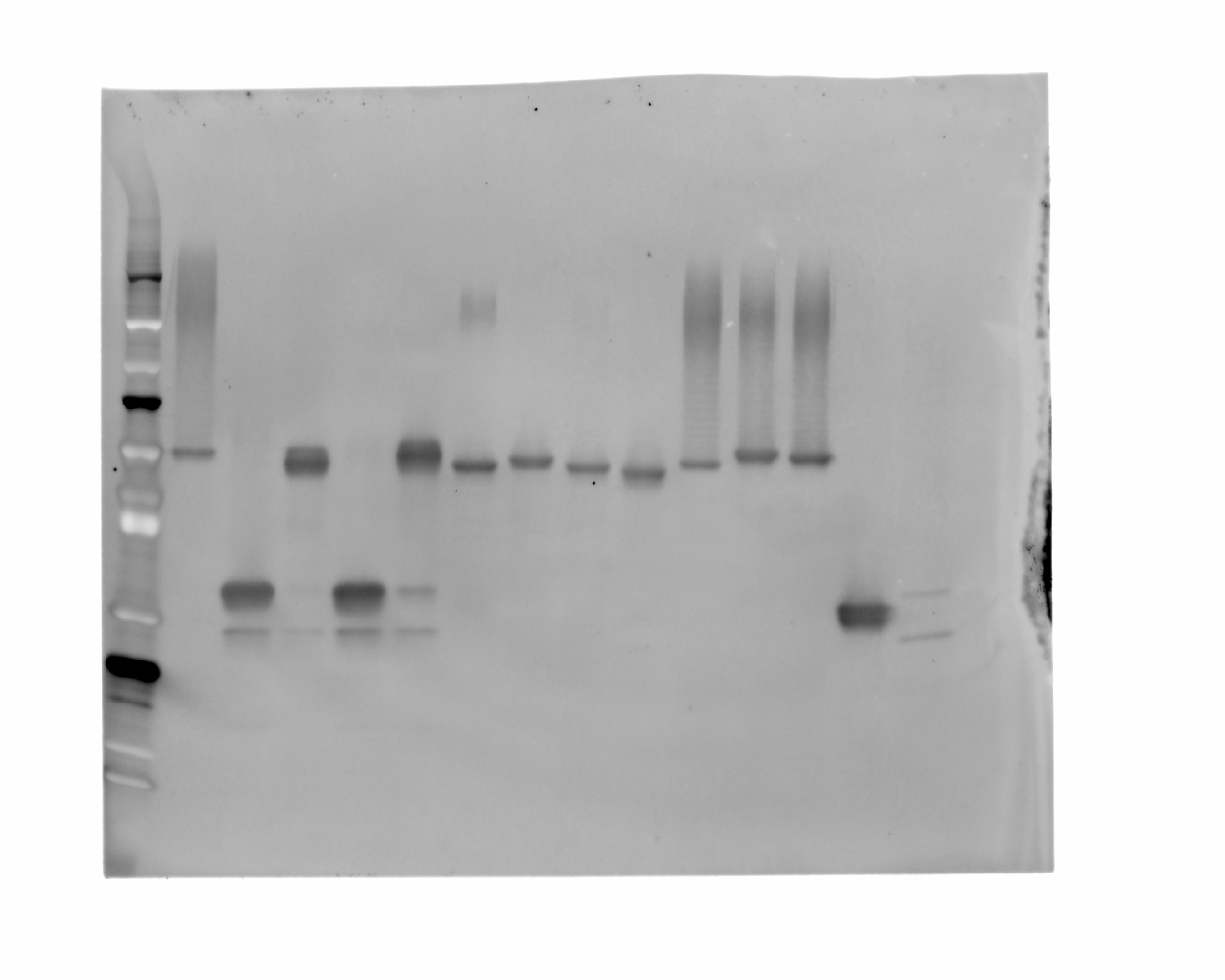

### 20241114_purity_gel4_27high_flaghrp_02(Composite).tif

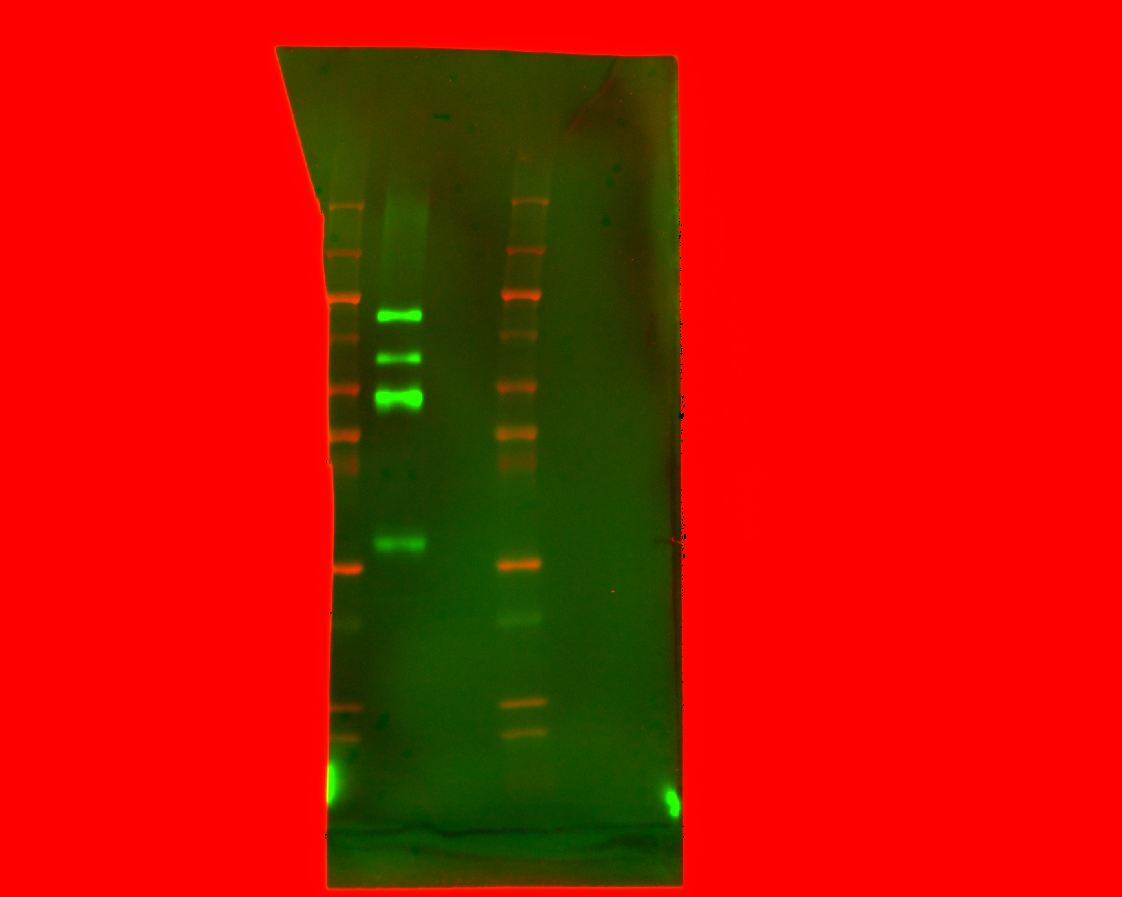

### 20241114_purity_gel4_27high_flaghrp_03(Composite).tif

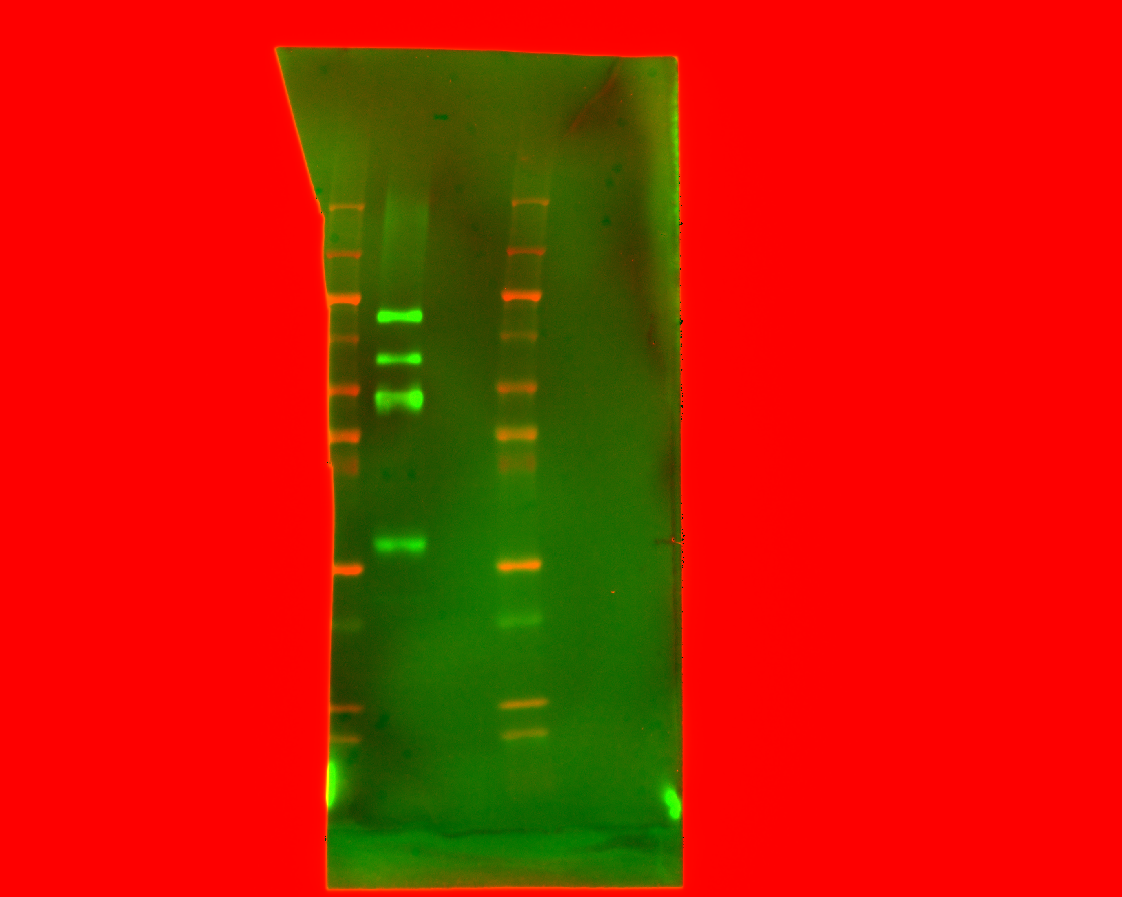

### 20241114_purity_gel4_26272921_his488_2(Composite).tif

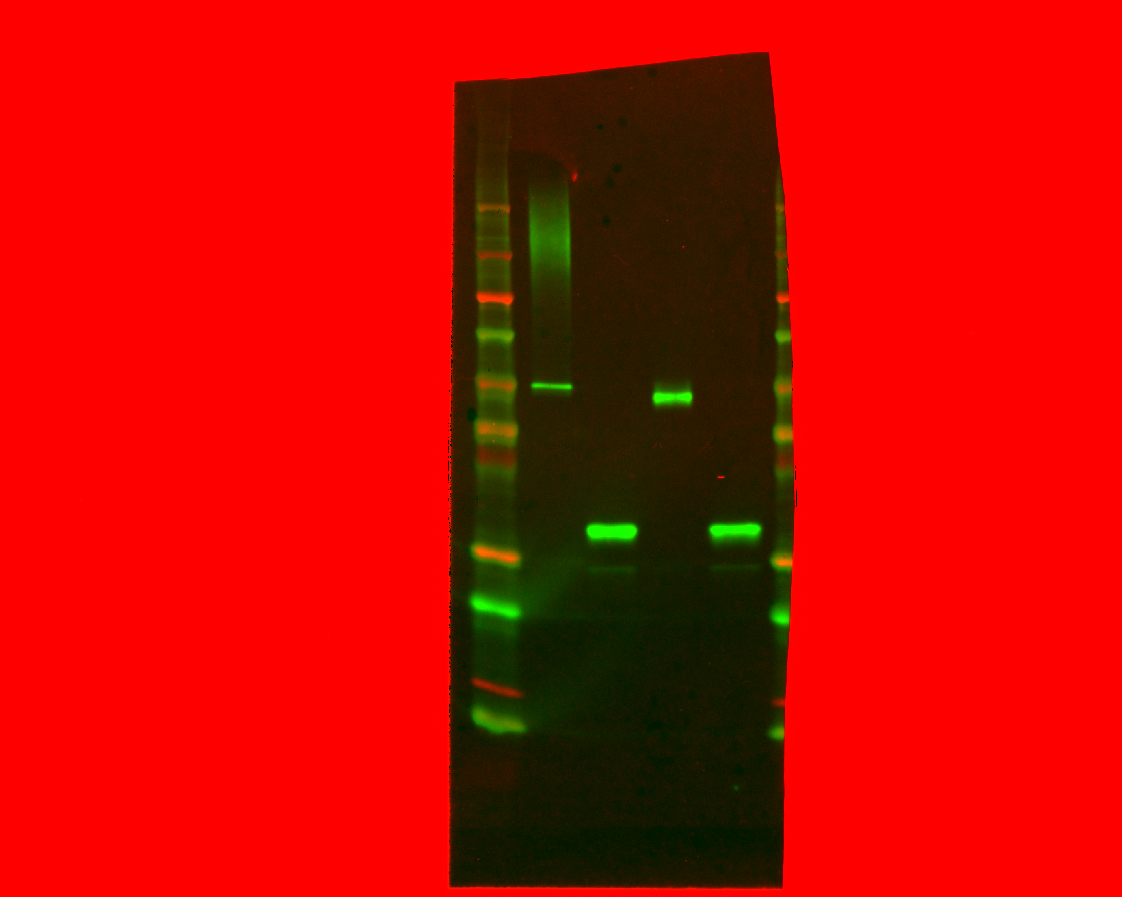

### 20241114_purity_gel4_26272921_his488_3(Composite).tif

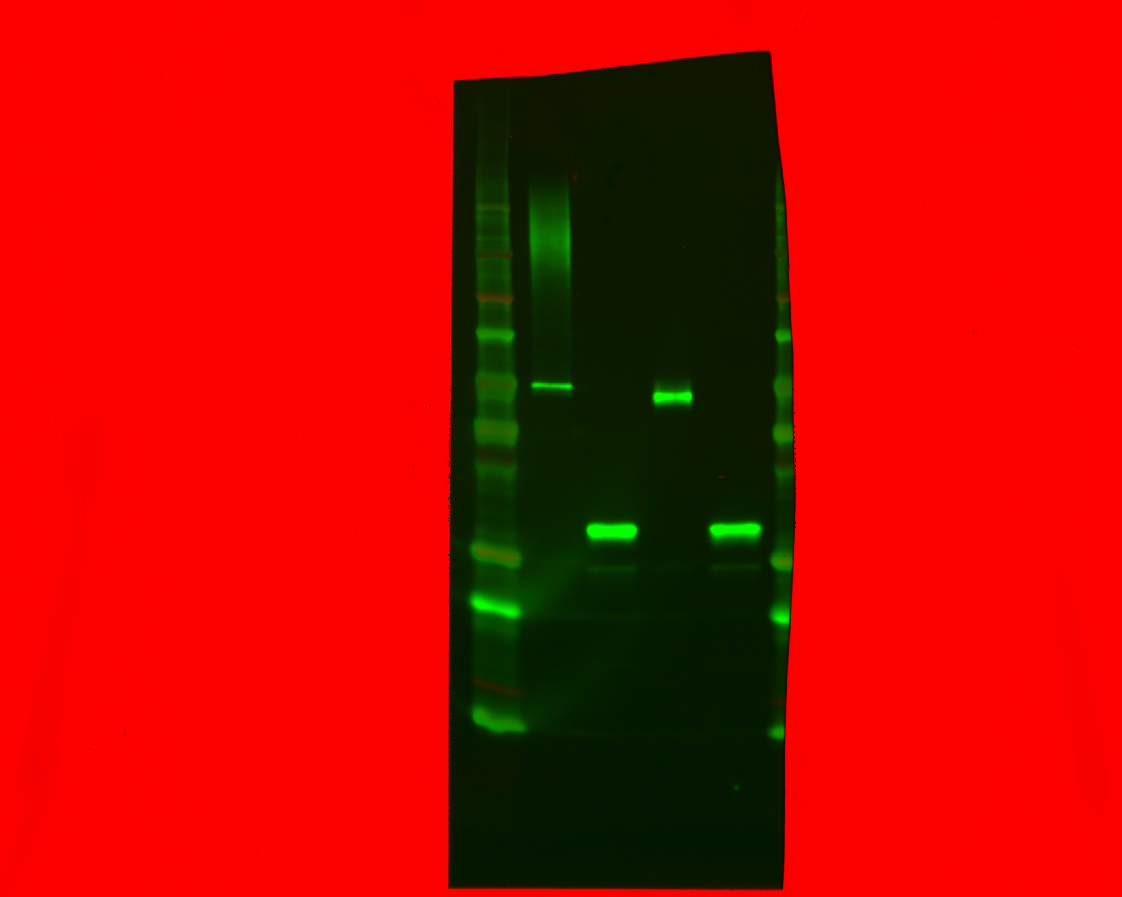

### 20241114_purity_gel4_26272921_his488_6(Alexa 488).tif

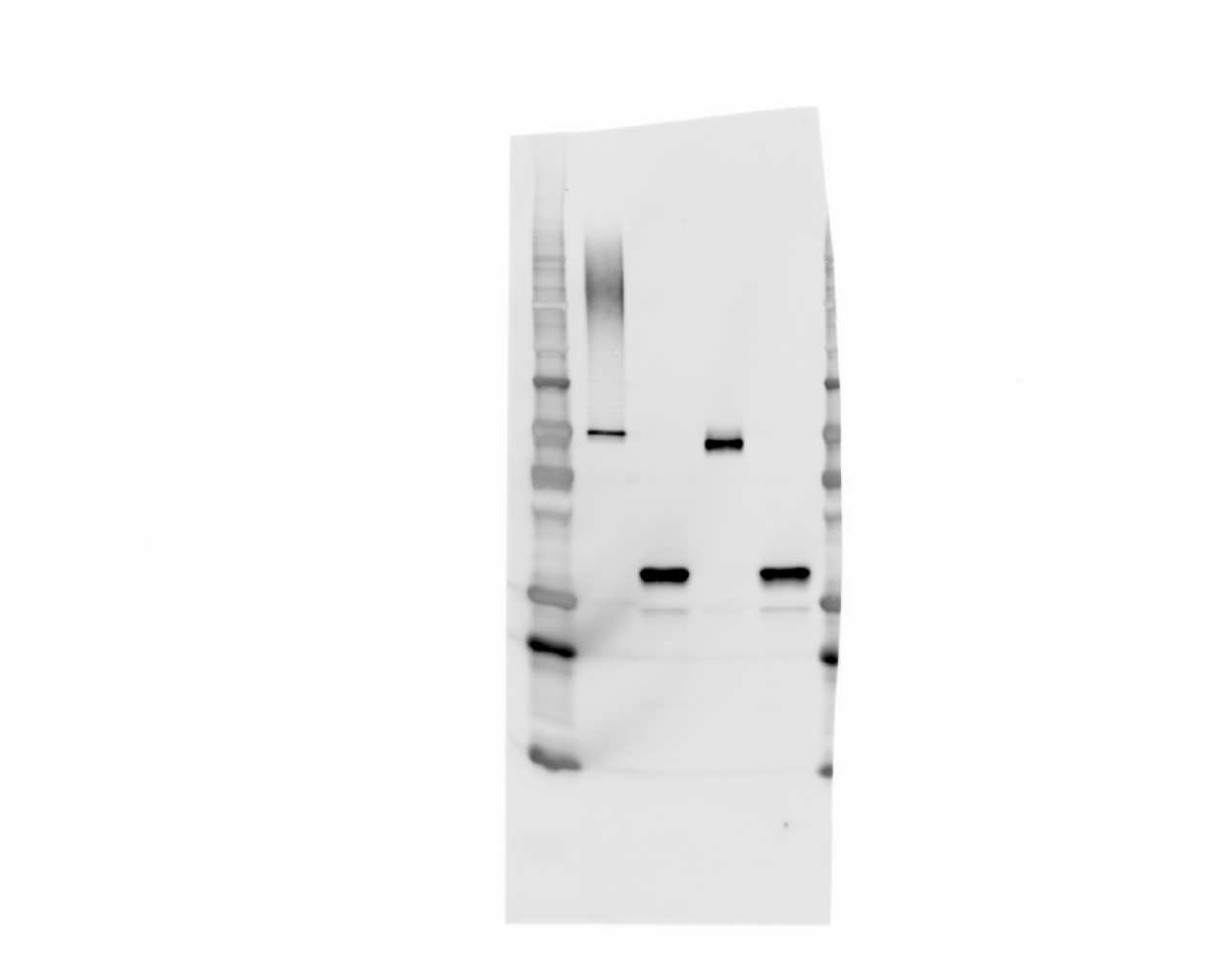

### 20241114_purity_gel4_26272921_his488_6(Composite).tif

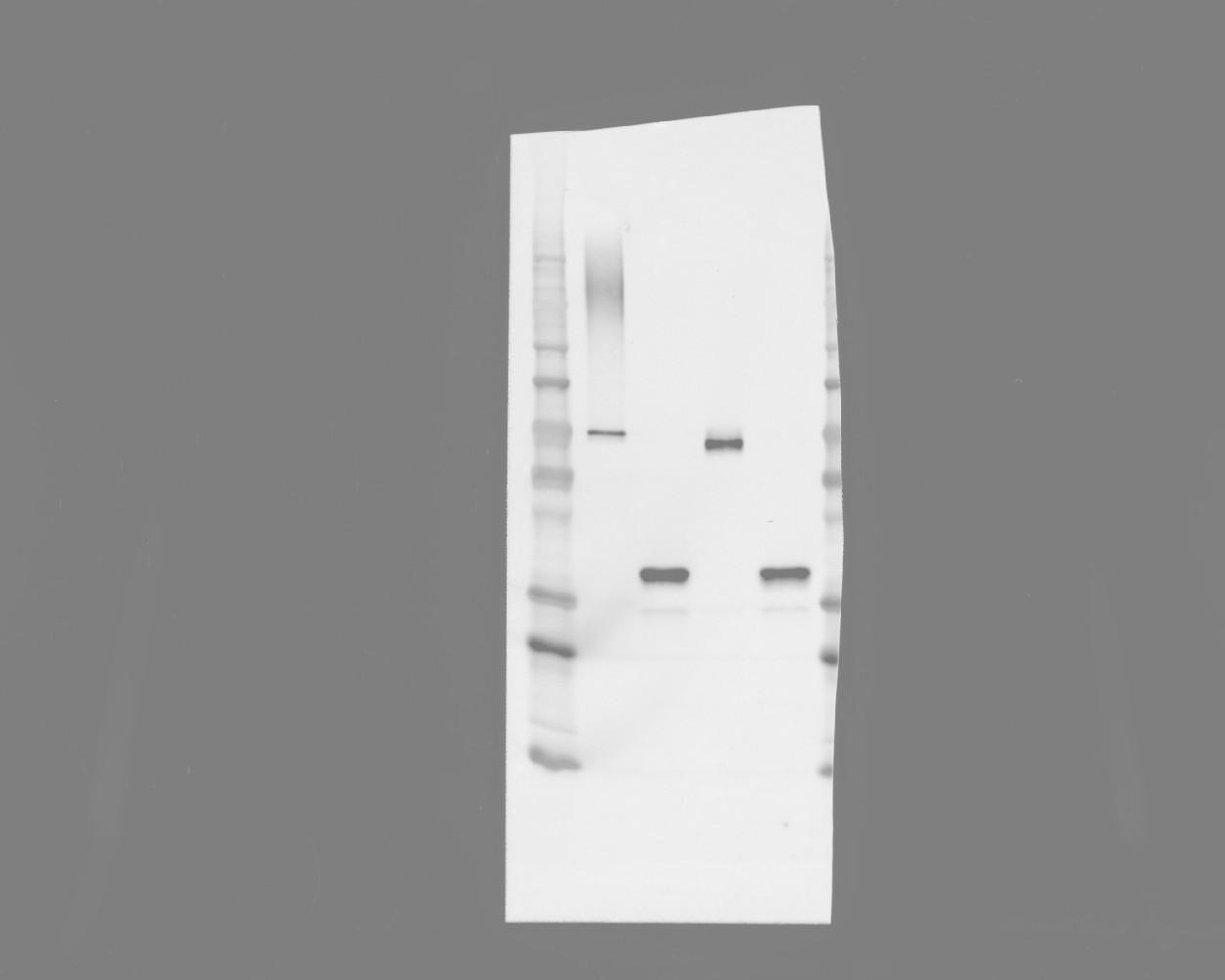

### 20241114_purity_gel4_26272921_his488_8(Alexa 488).tif

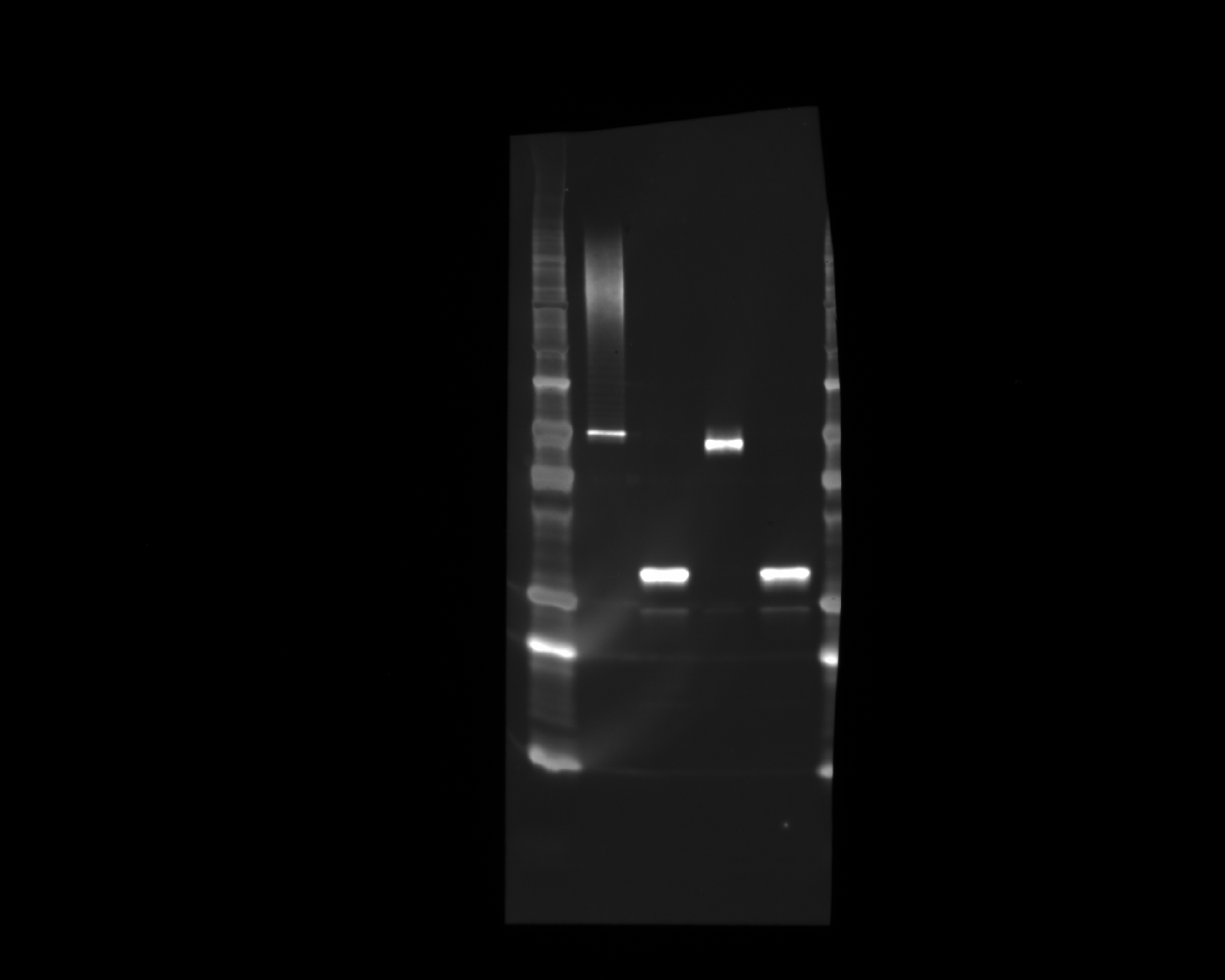

### 20241114_purity_gel4_26272921_his488_8(Composite).tif

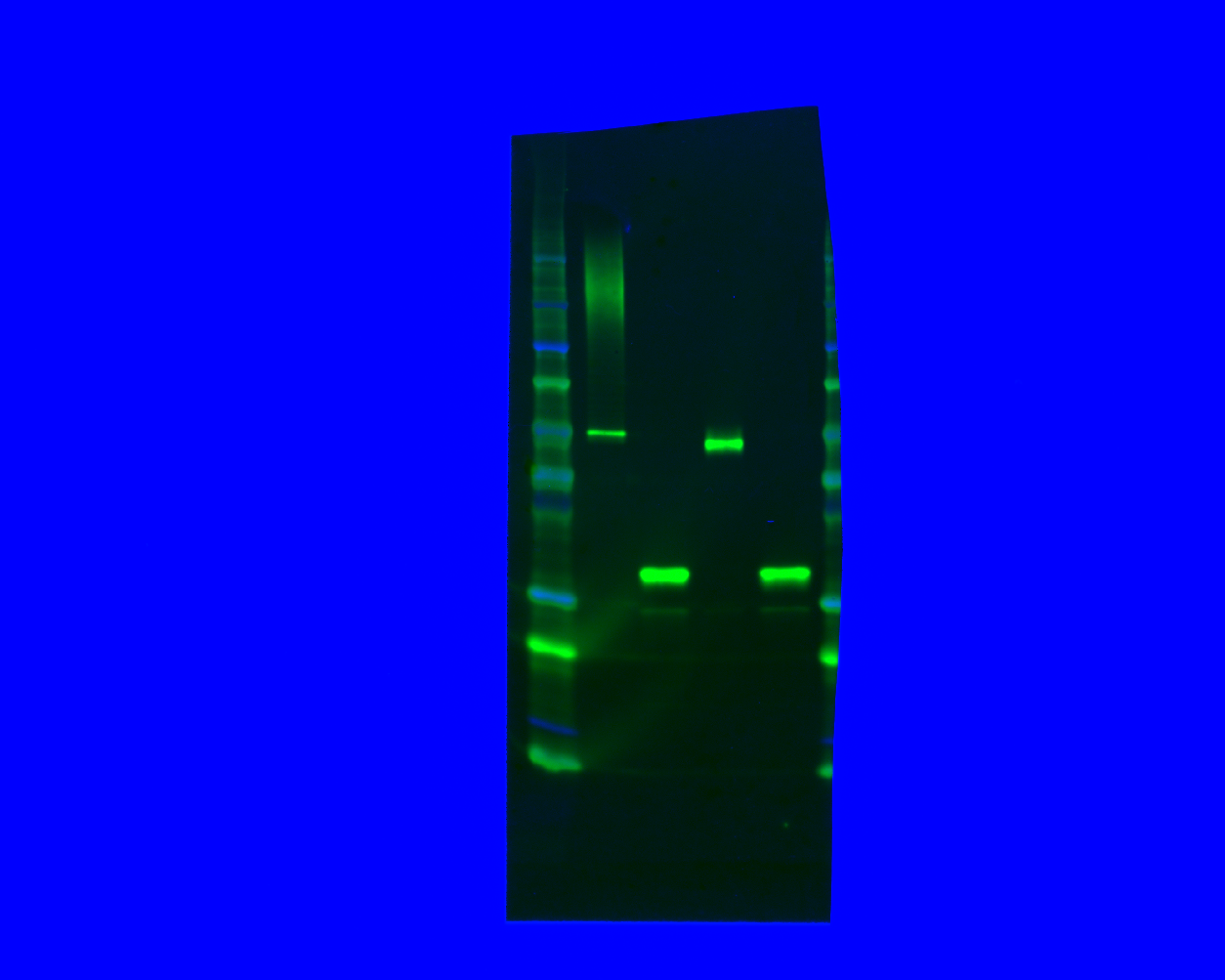

### 20241114_purity_gel4_27292124_flaghrp_04(Composite).tif

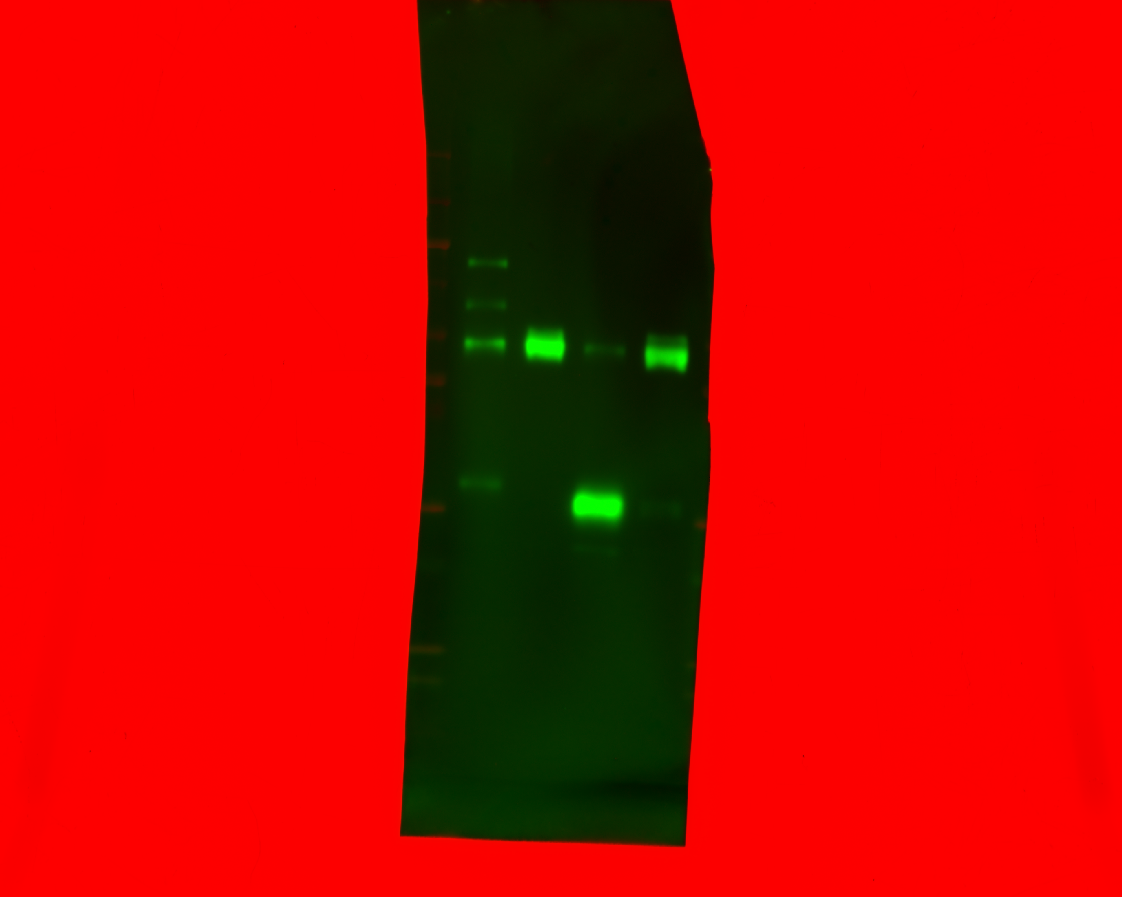

### 20241114_purity_gel4_27292124_flaghrp_05(Composite).tif

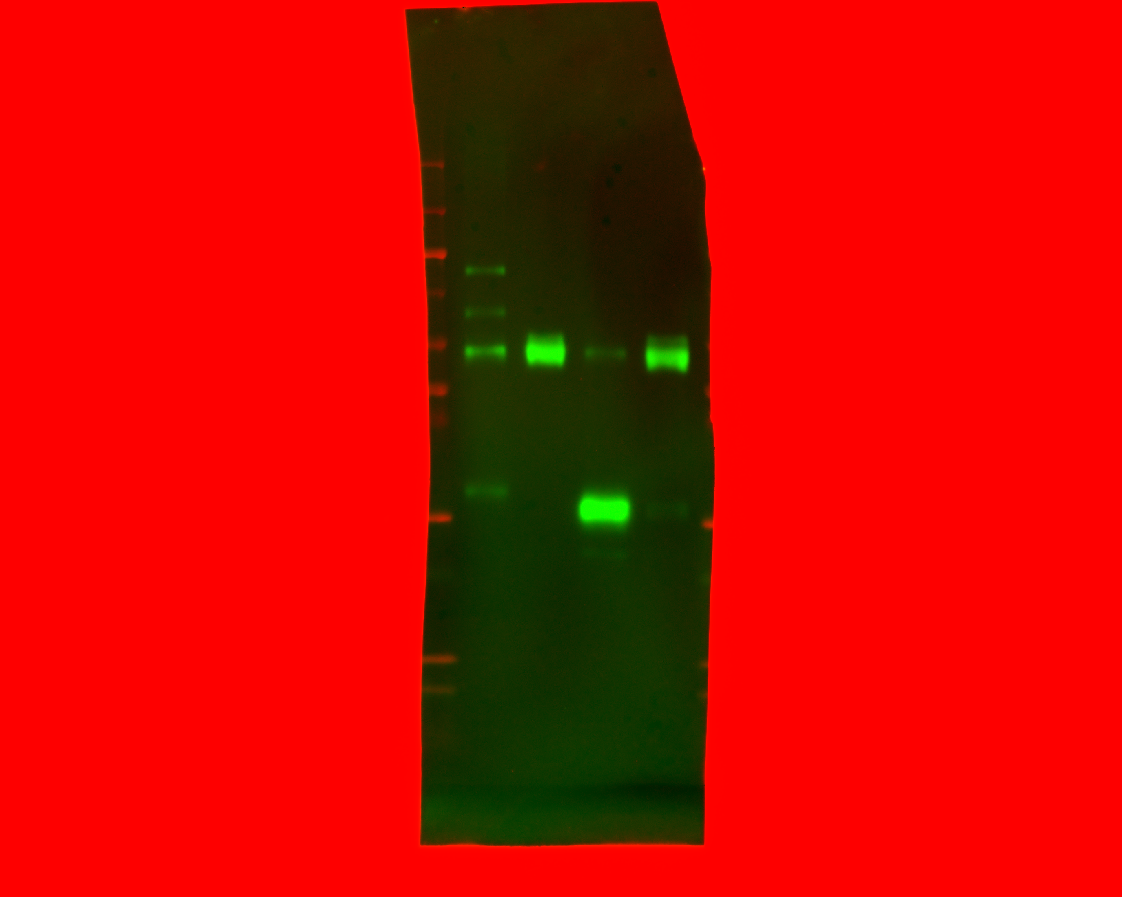

### 20241114_purity_gel4_27292124_flaghrp_06(Composite).tif

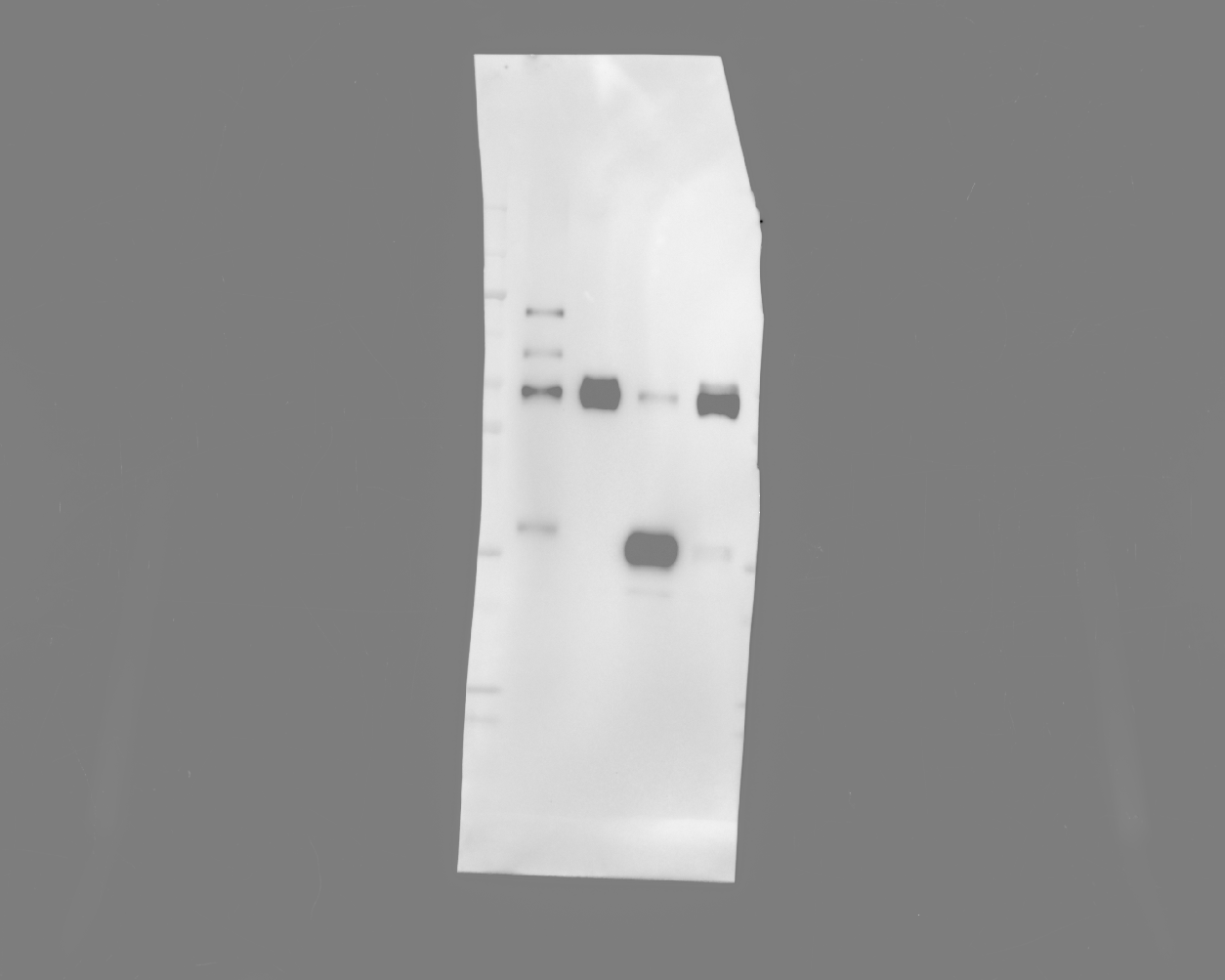

### 20241114_purity_gel4_27292124_flaghrp_07(Composite).tif

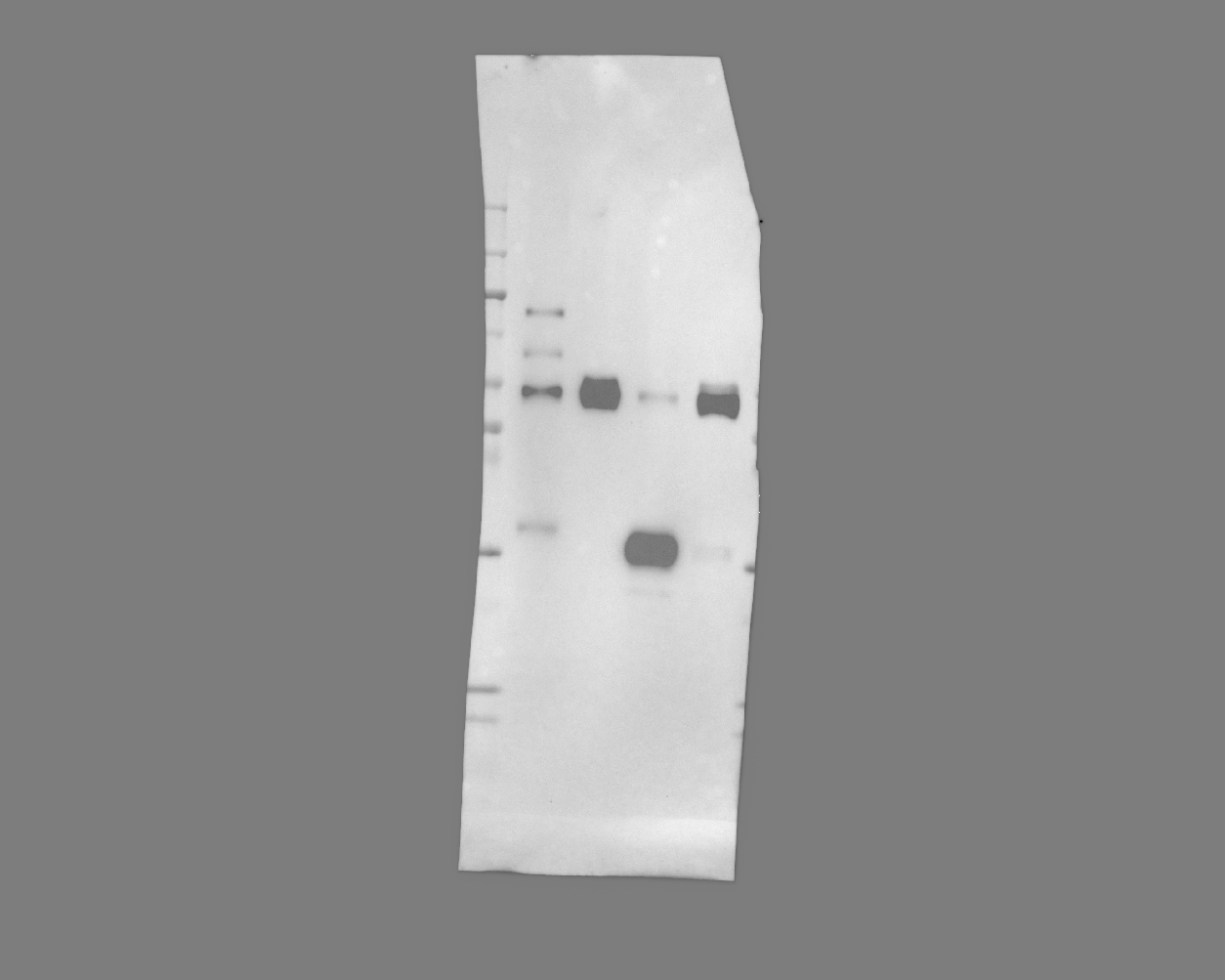

### 20241114_purity_gel4_27292124_flaghrp_08(Composite).tif

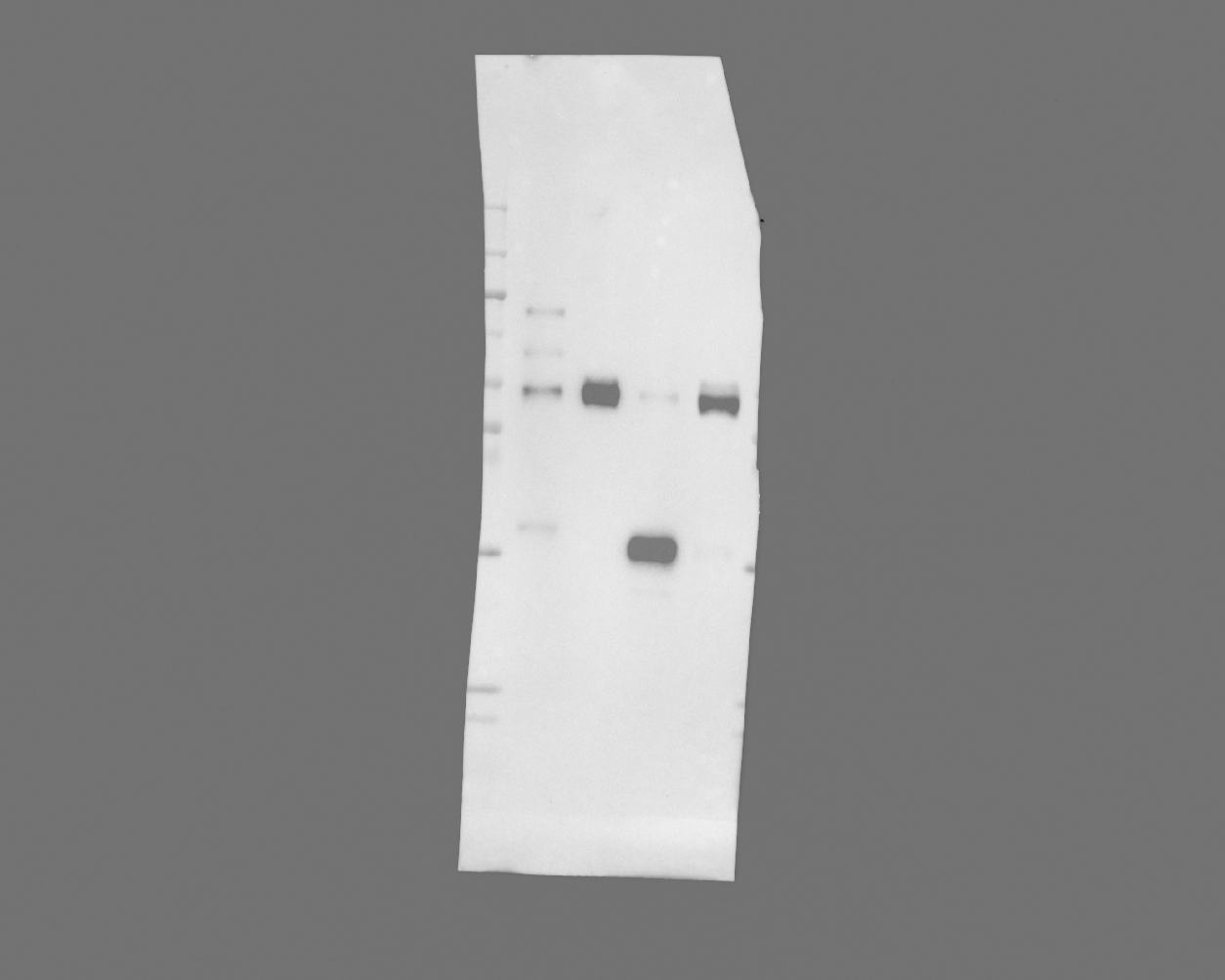

### 20241114_purity_gel4_27292124_flaghrp_09(Composite).tif

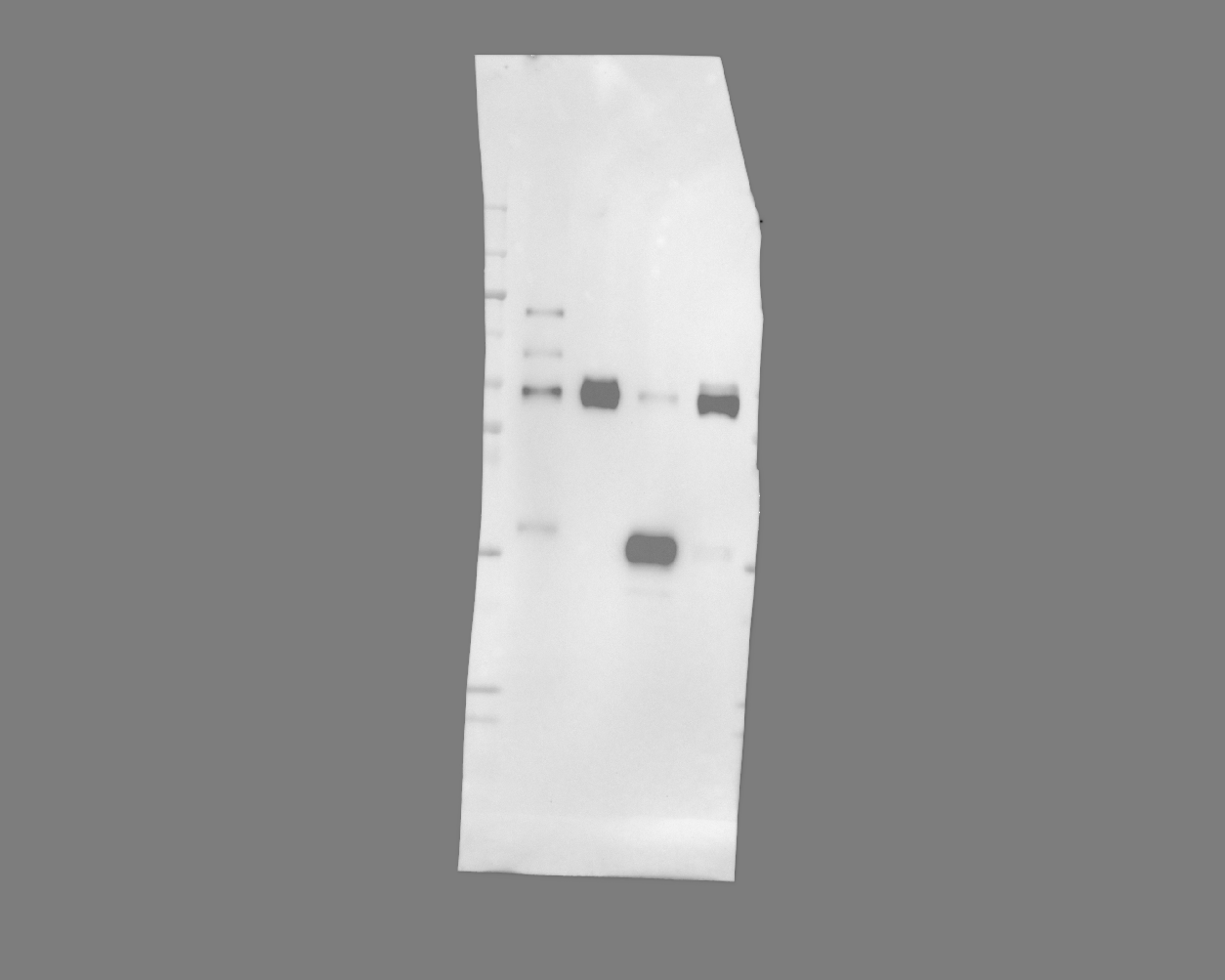

### 20241114_purity_gel4_27292124_flaghrp_10(Composite).tif

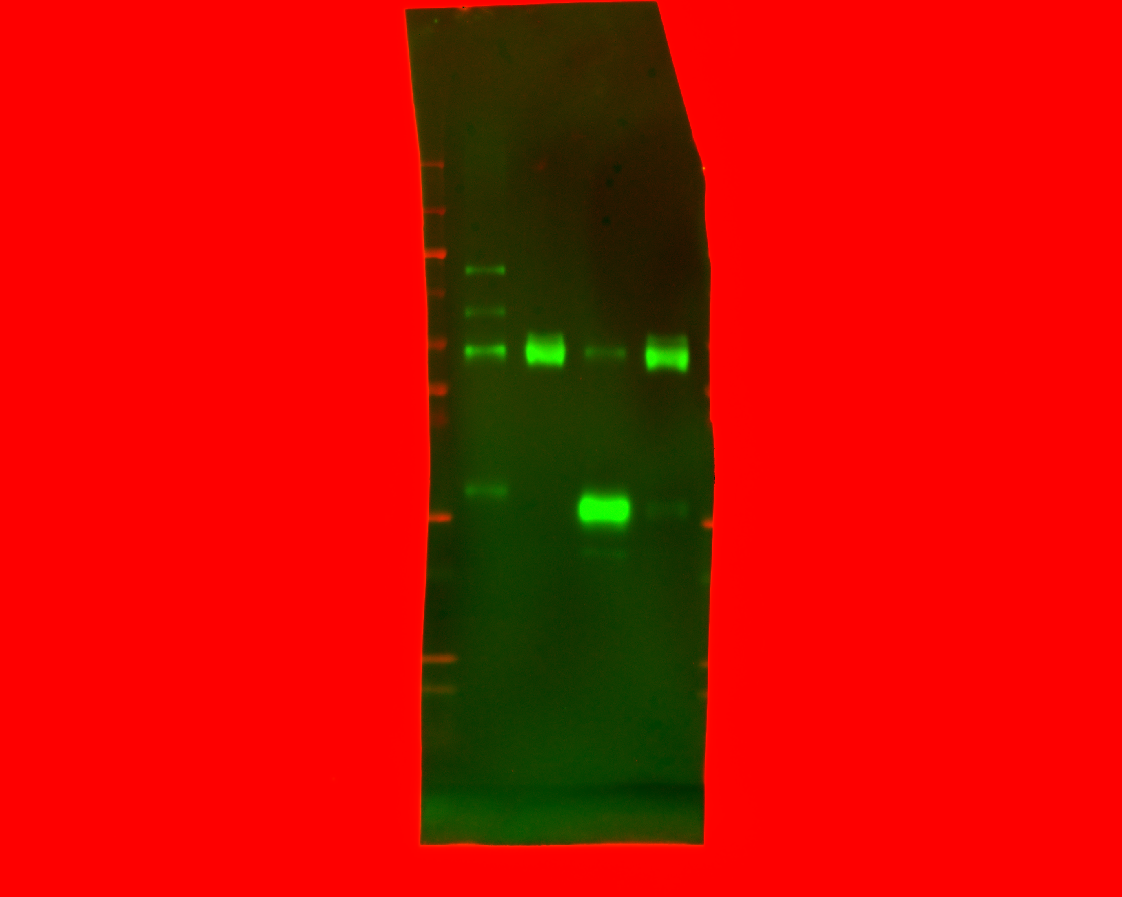

### 20241114_purity_gel4_27292124_flaghrp_11(Composite).tif

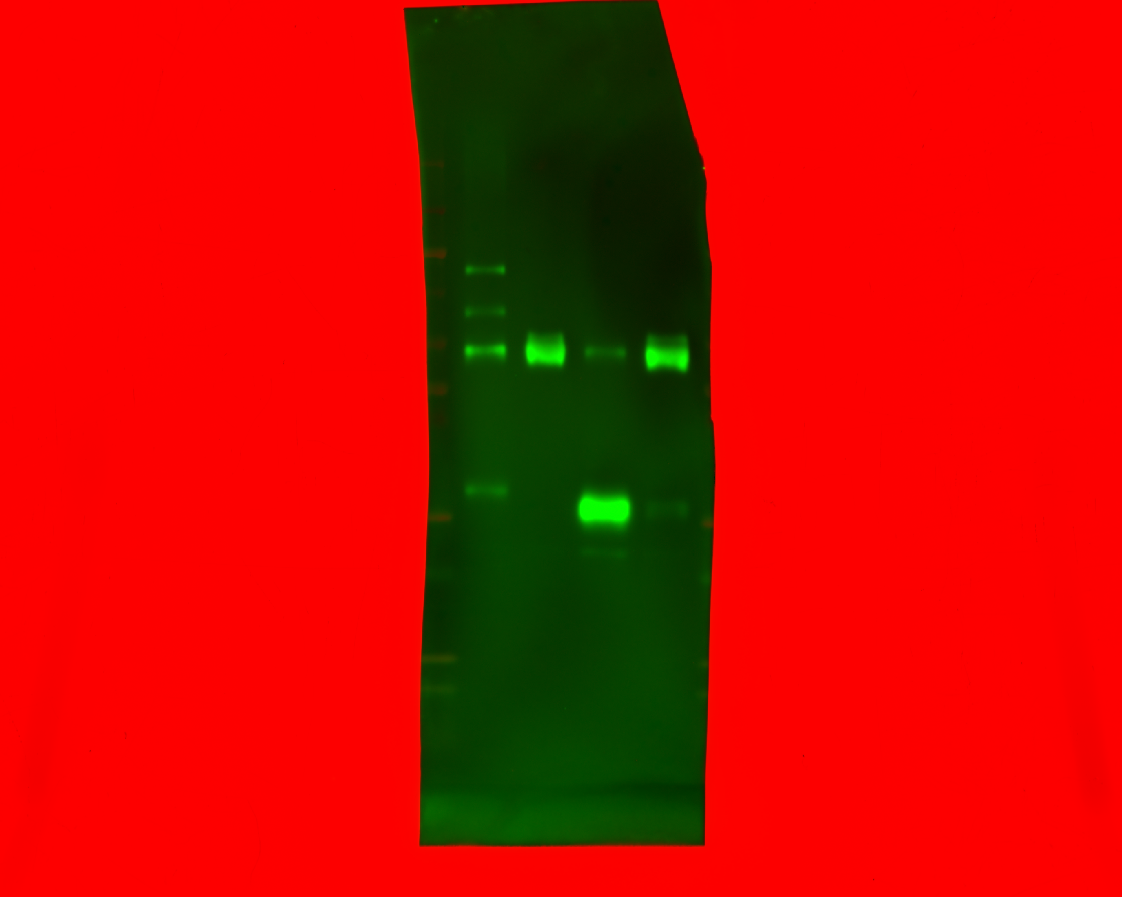

### 20241119_ut7variantgel_gel2_hh3_7(Colorimetric).tif

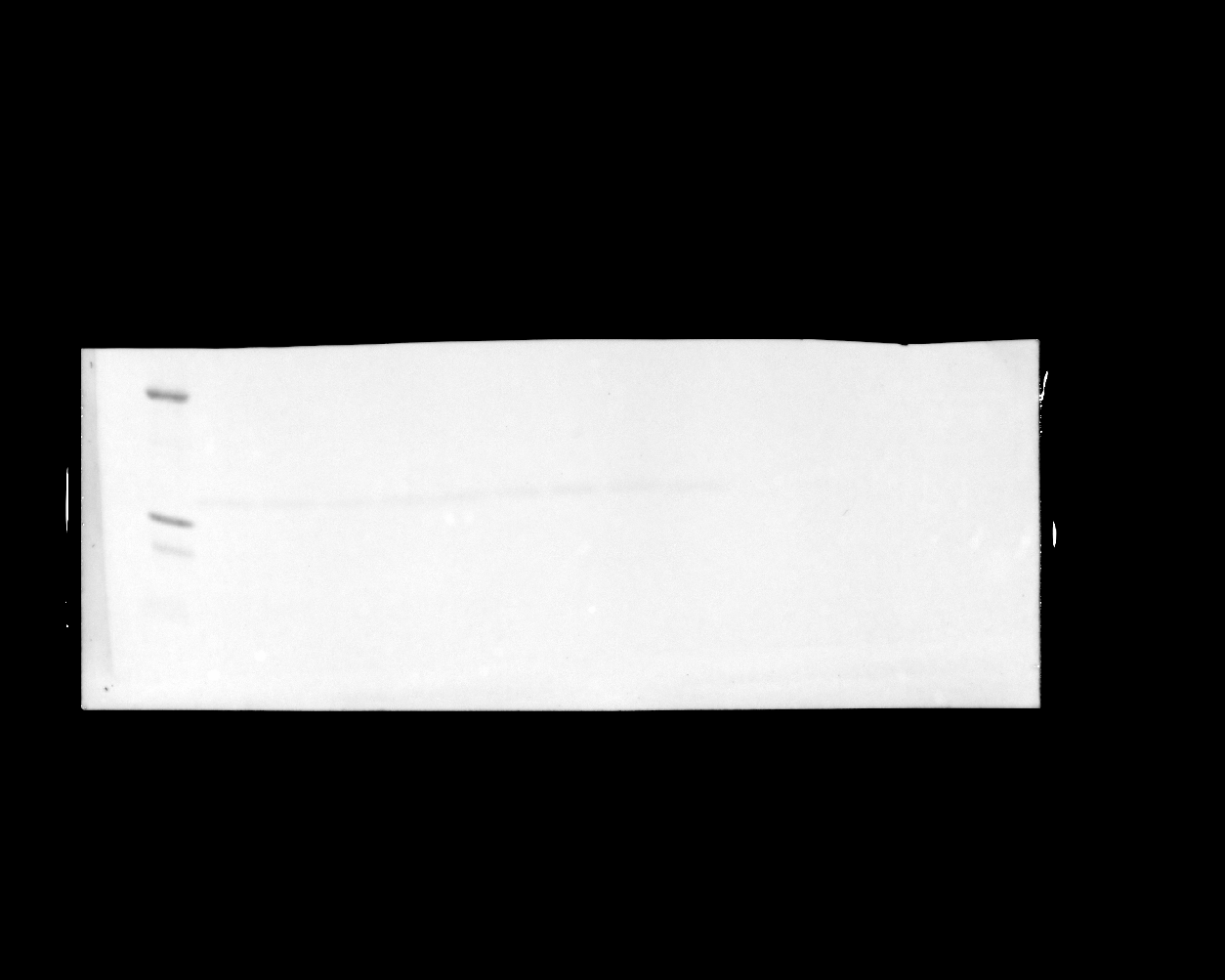

### 20241119_ut7variantgel_gel2_hh3_7(Composite).tif

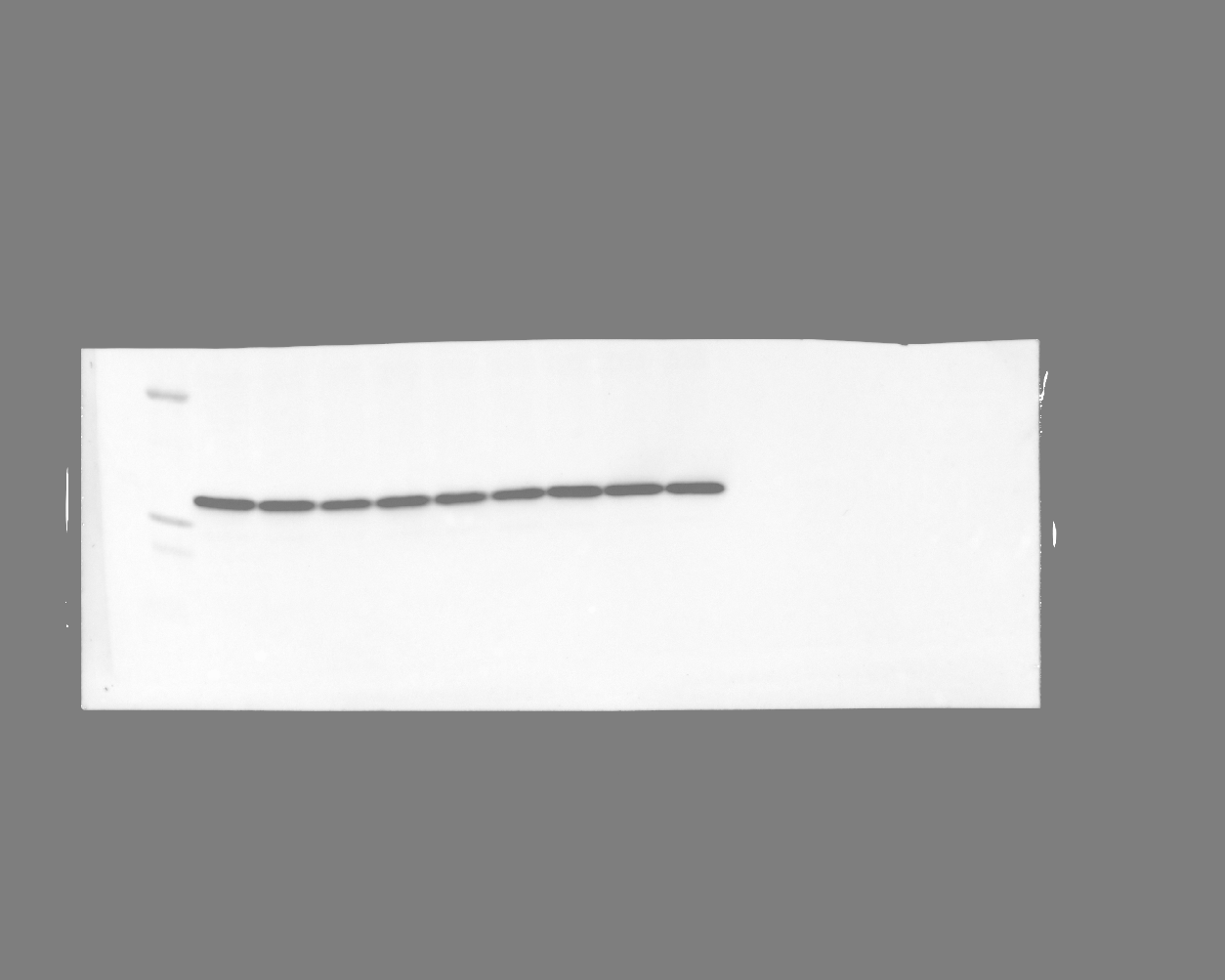

### 20241119_ut7variantgel_gel2_stat5_phospho_28(Chemiluminescence).tif

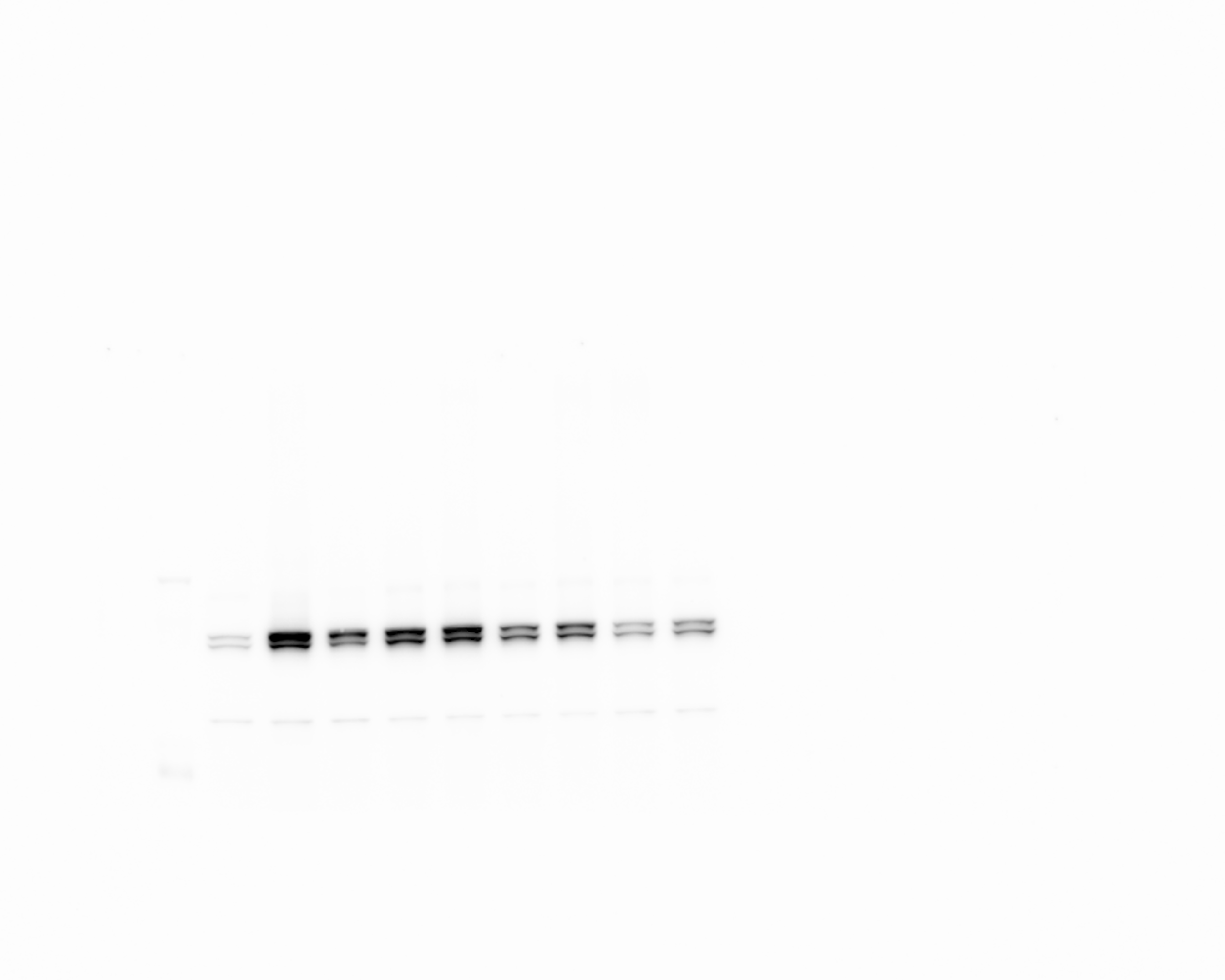

### 20241119_ut7variantgel_gel2_stat5_phospho_28(Colorimetric).tif

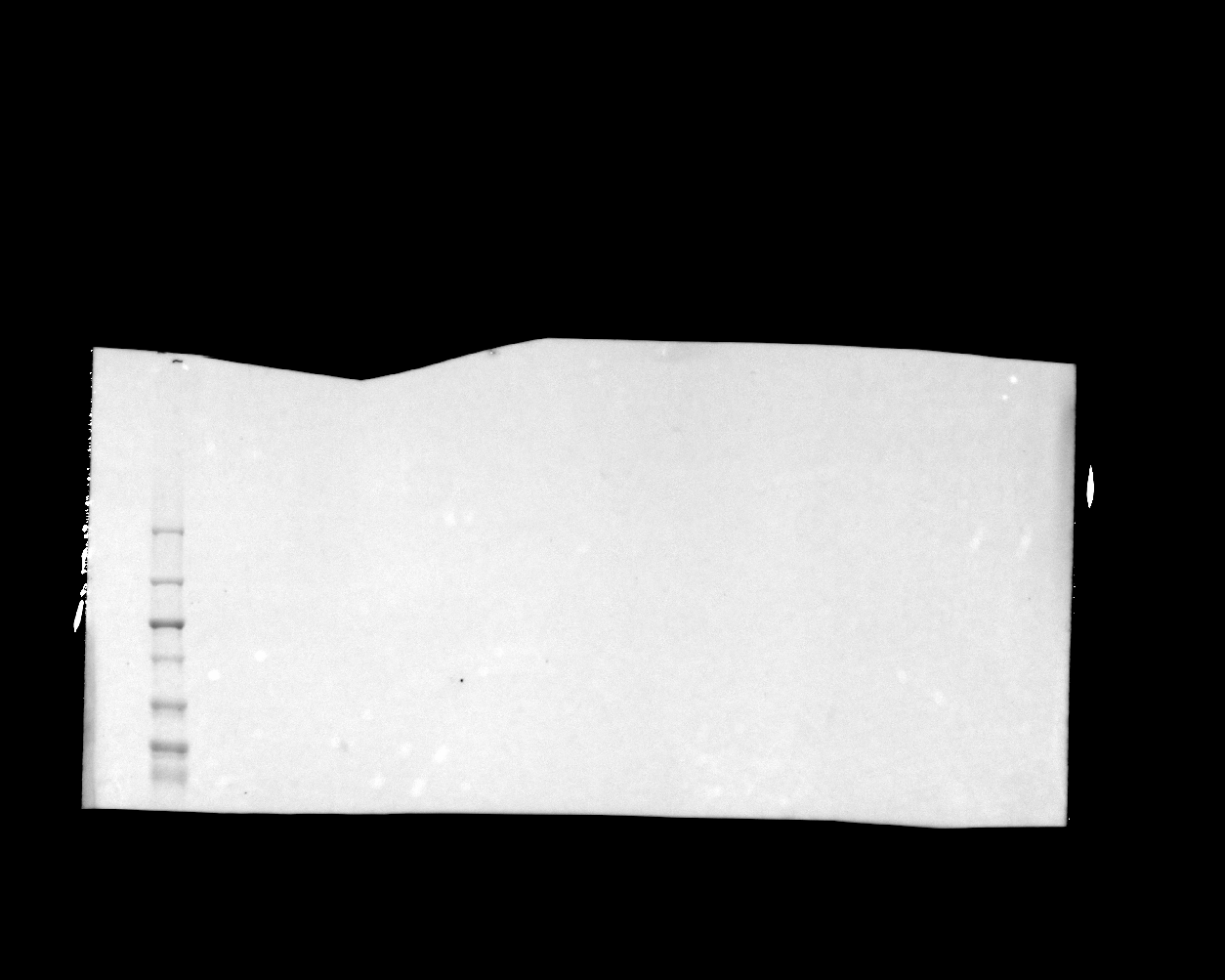

### 20241119_ut7variantgel_gel2_stat5_phospho_28(Composite).tif

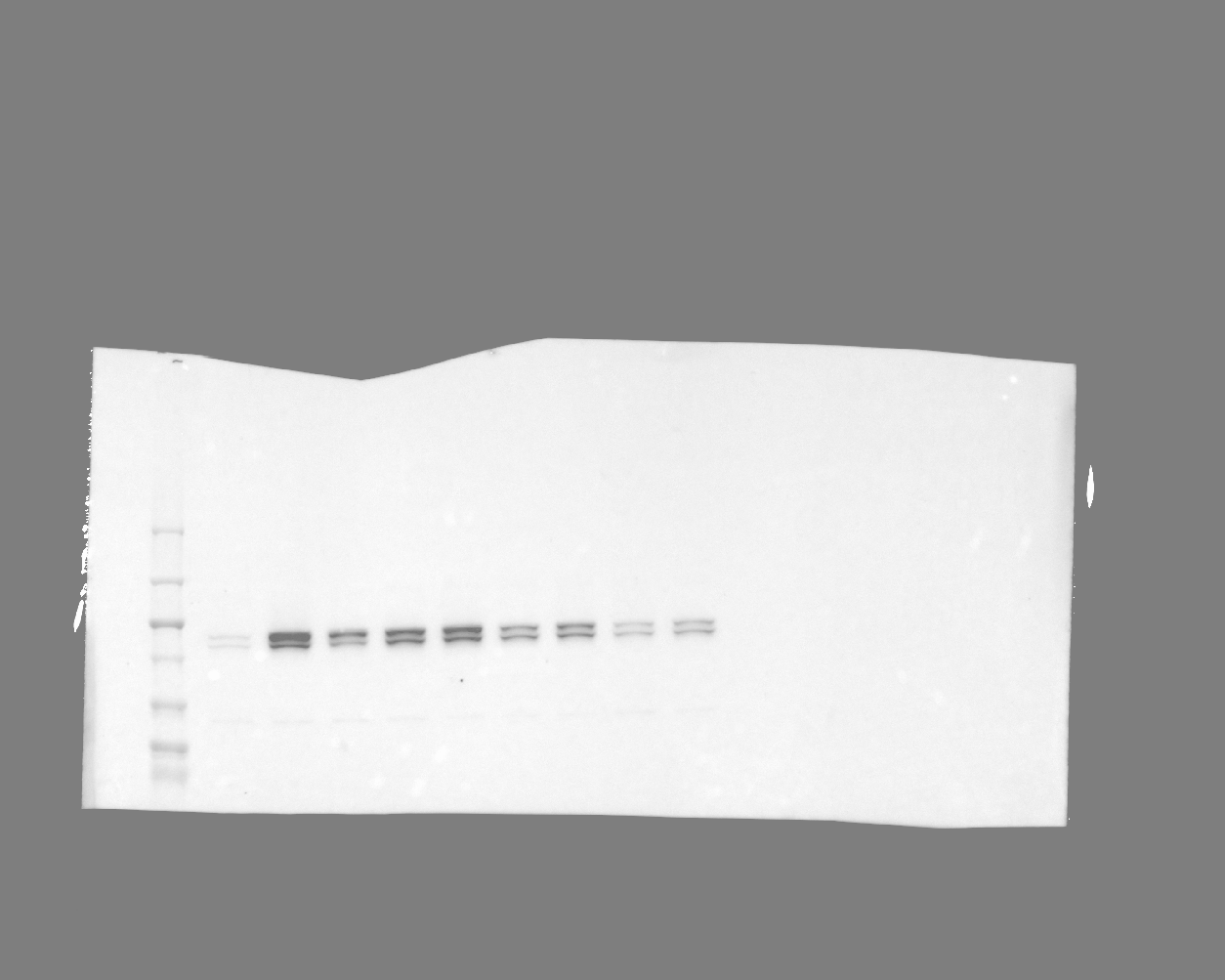

### 20241119_ut7variantgel_gel4_hh3_7(Chemiluminescence).tif

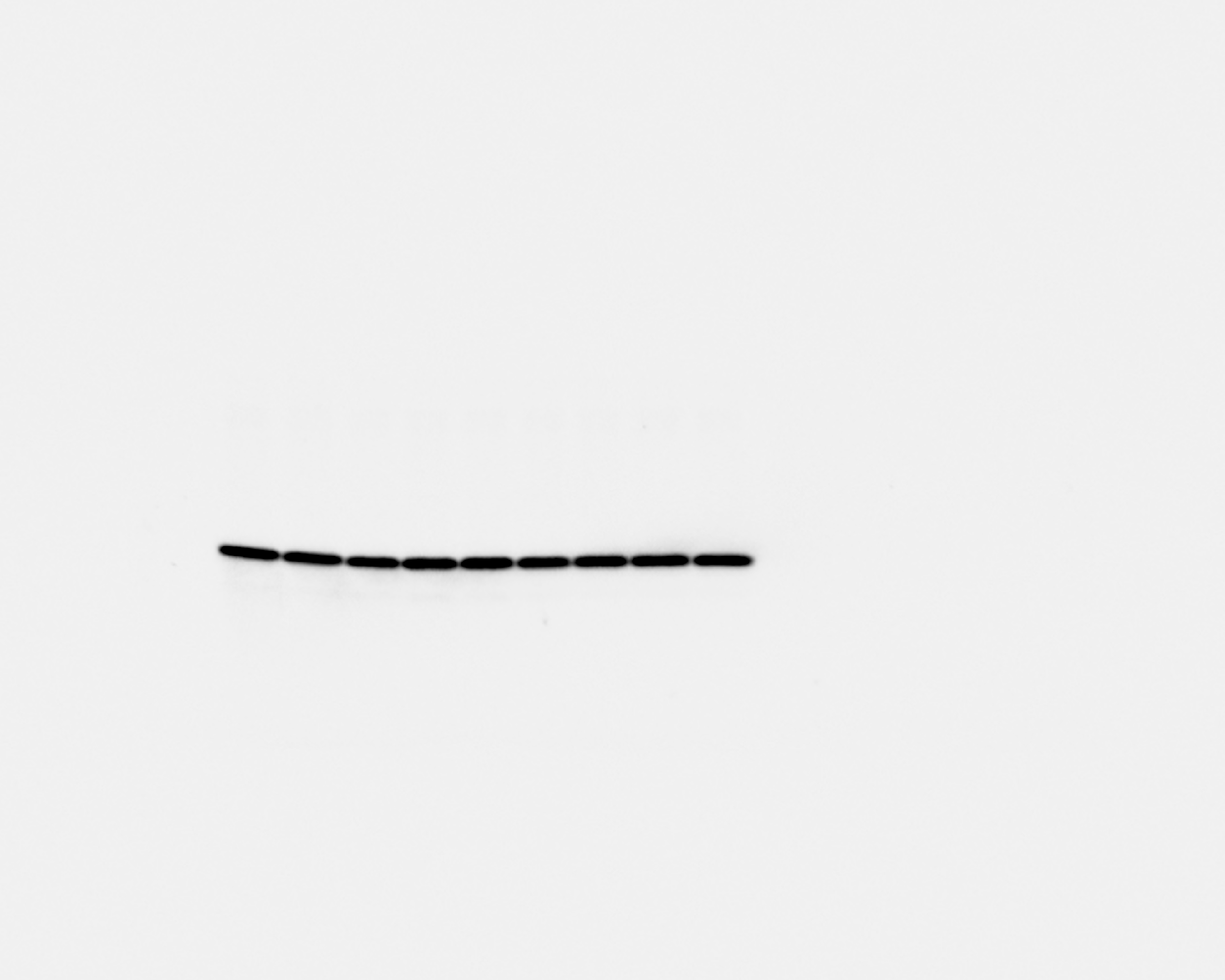

### 20241119_ut7variantgel_gel4_hh3_7(Colorimetric).tif

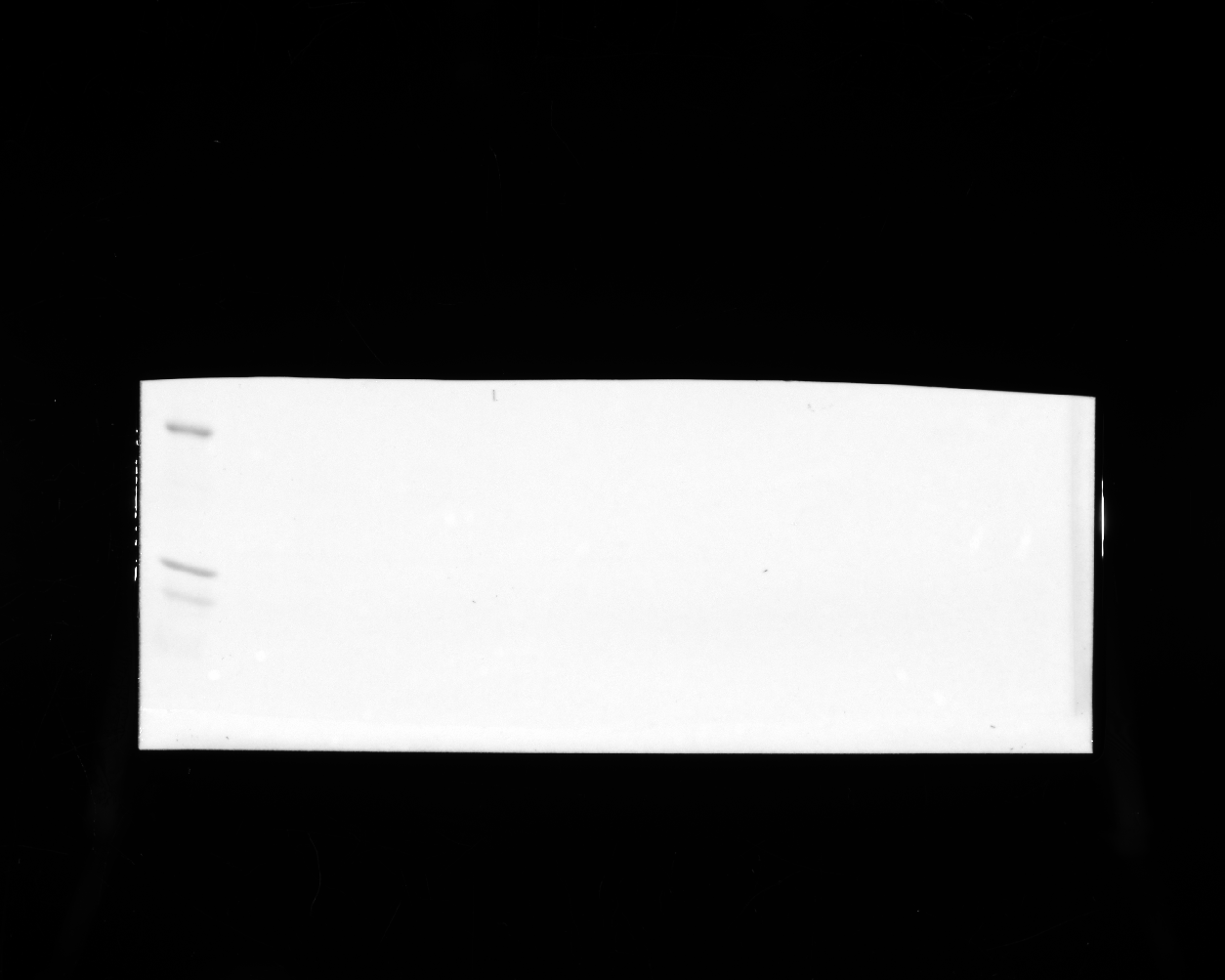

### 20241119_ut7variantgel_gel4_hh3_7(Composite).tif

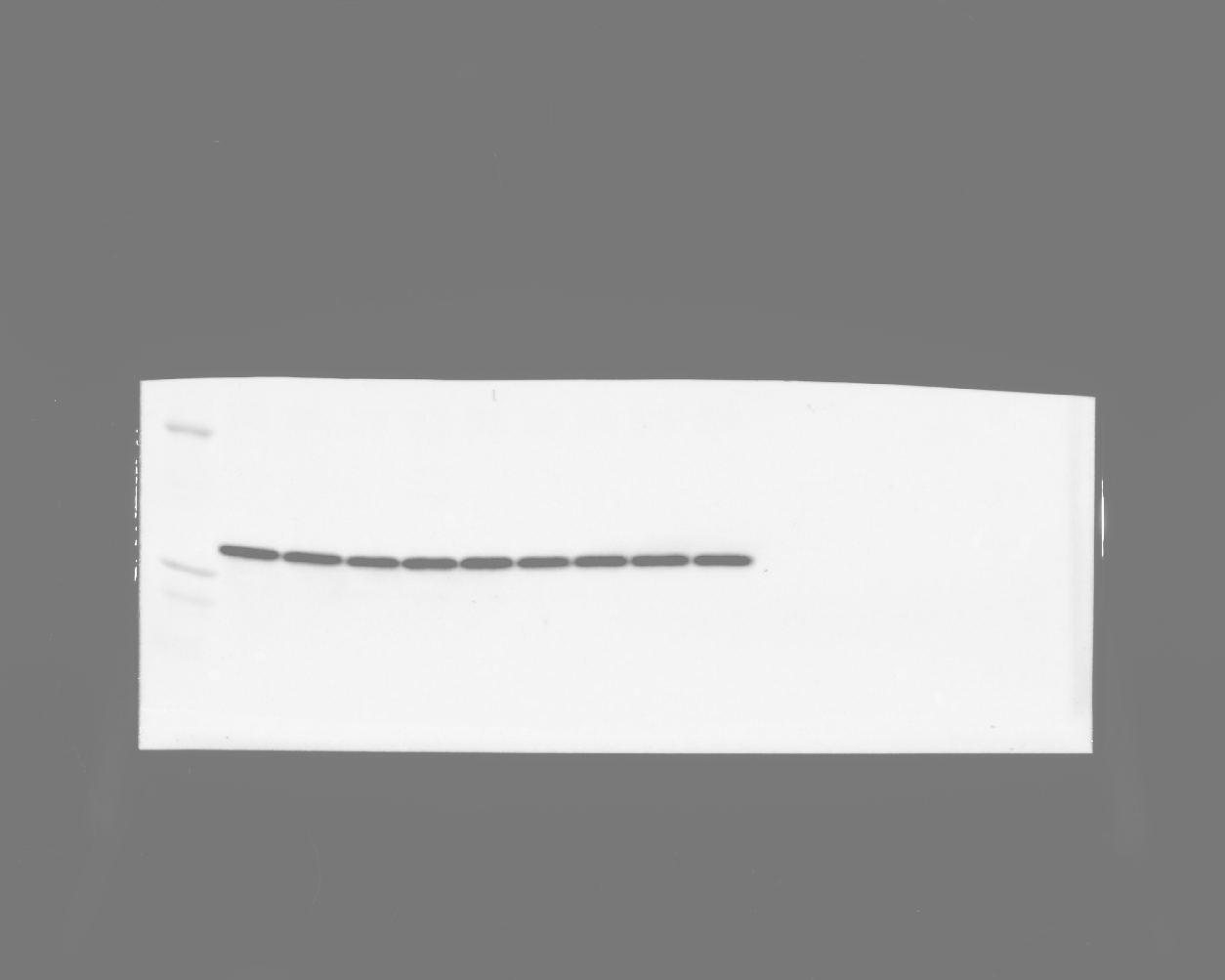

### 20241119_ut7variantgel_gel4_stat5_7(Chemiluminescence).tif

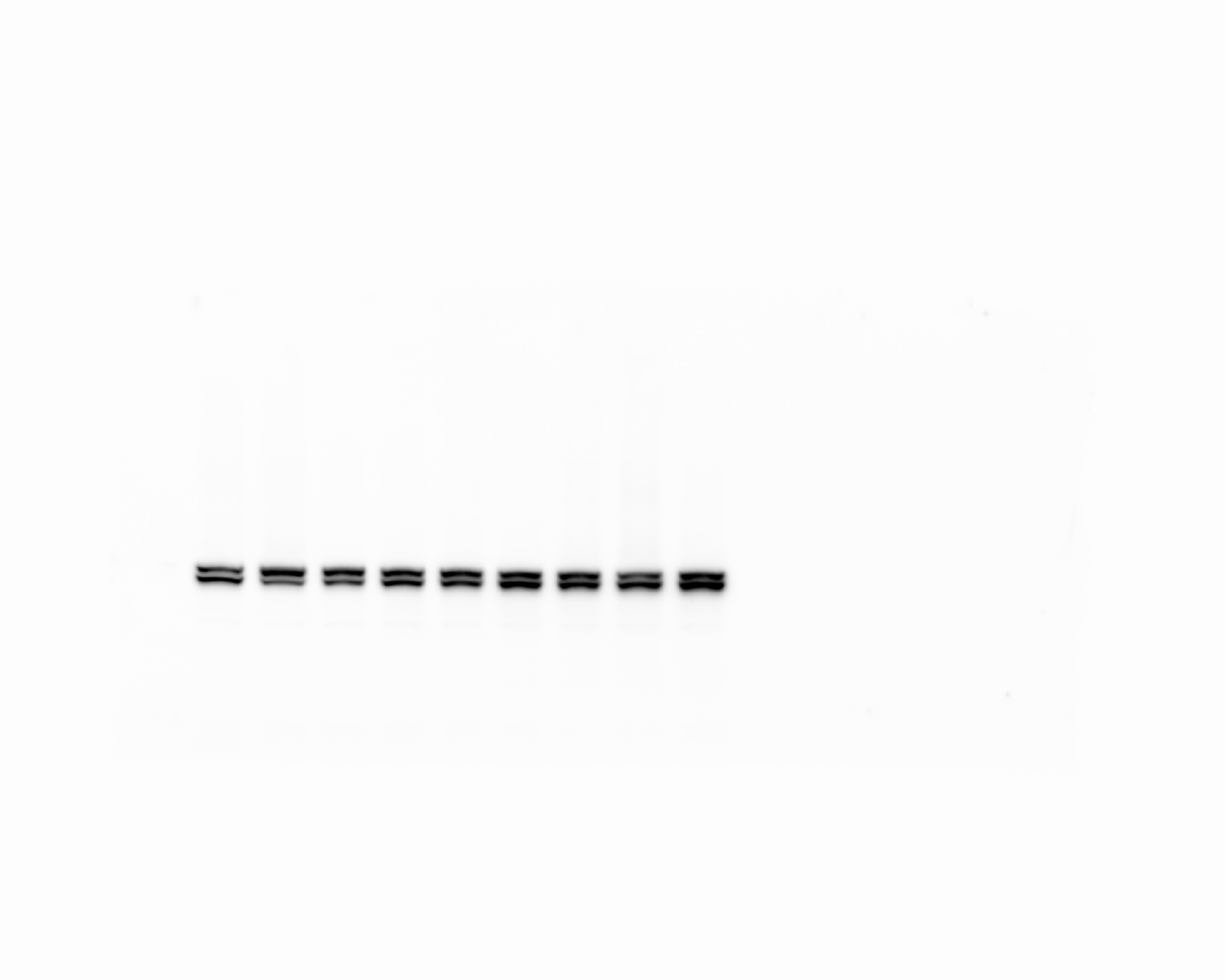

### 20241119_ut7variantgel_gel4_stat5_7(Colorimetric).tif

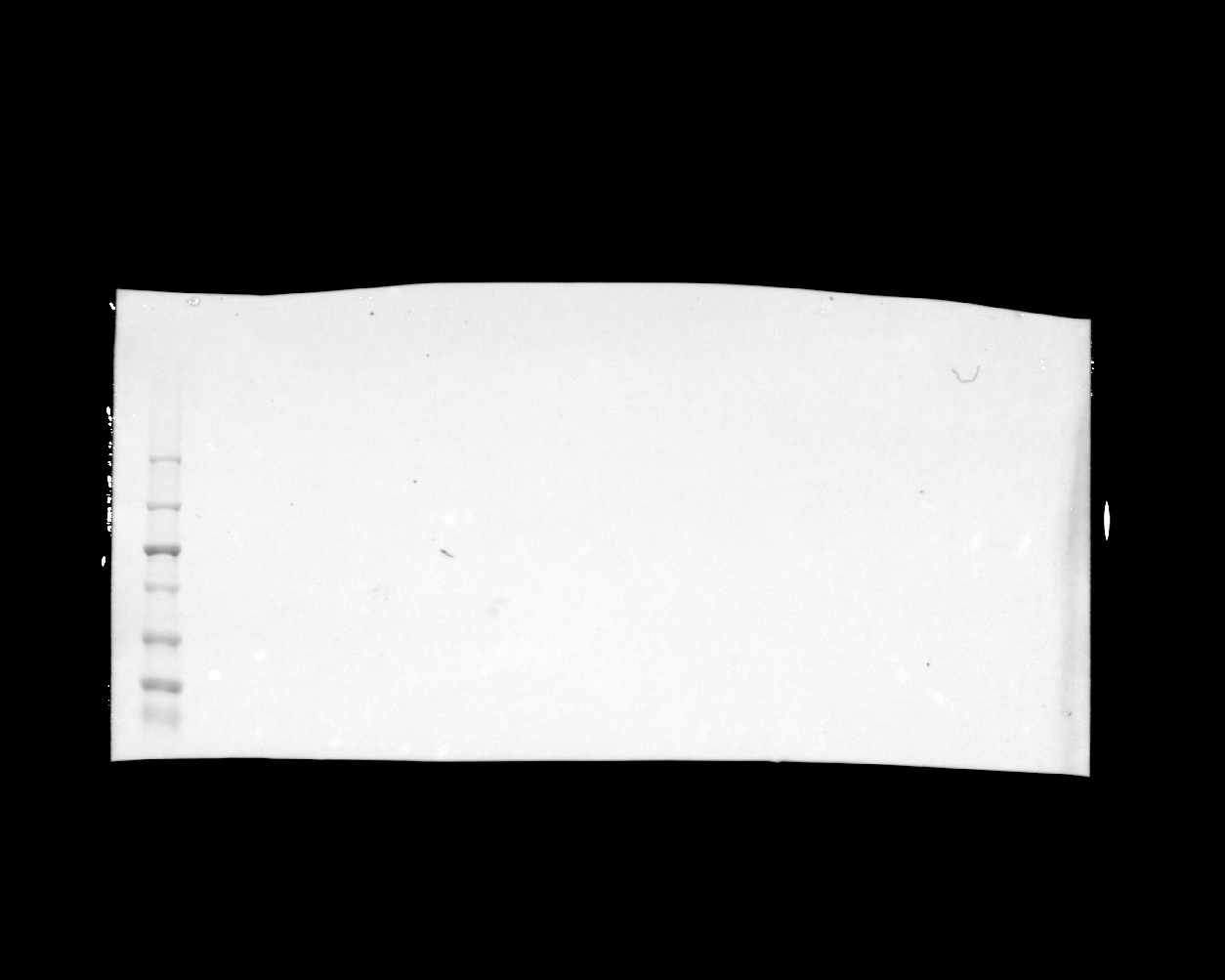

### 20241119_ut7variantgel_gel4_stat5_7(Composite).tif

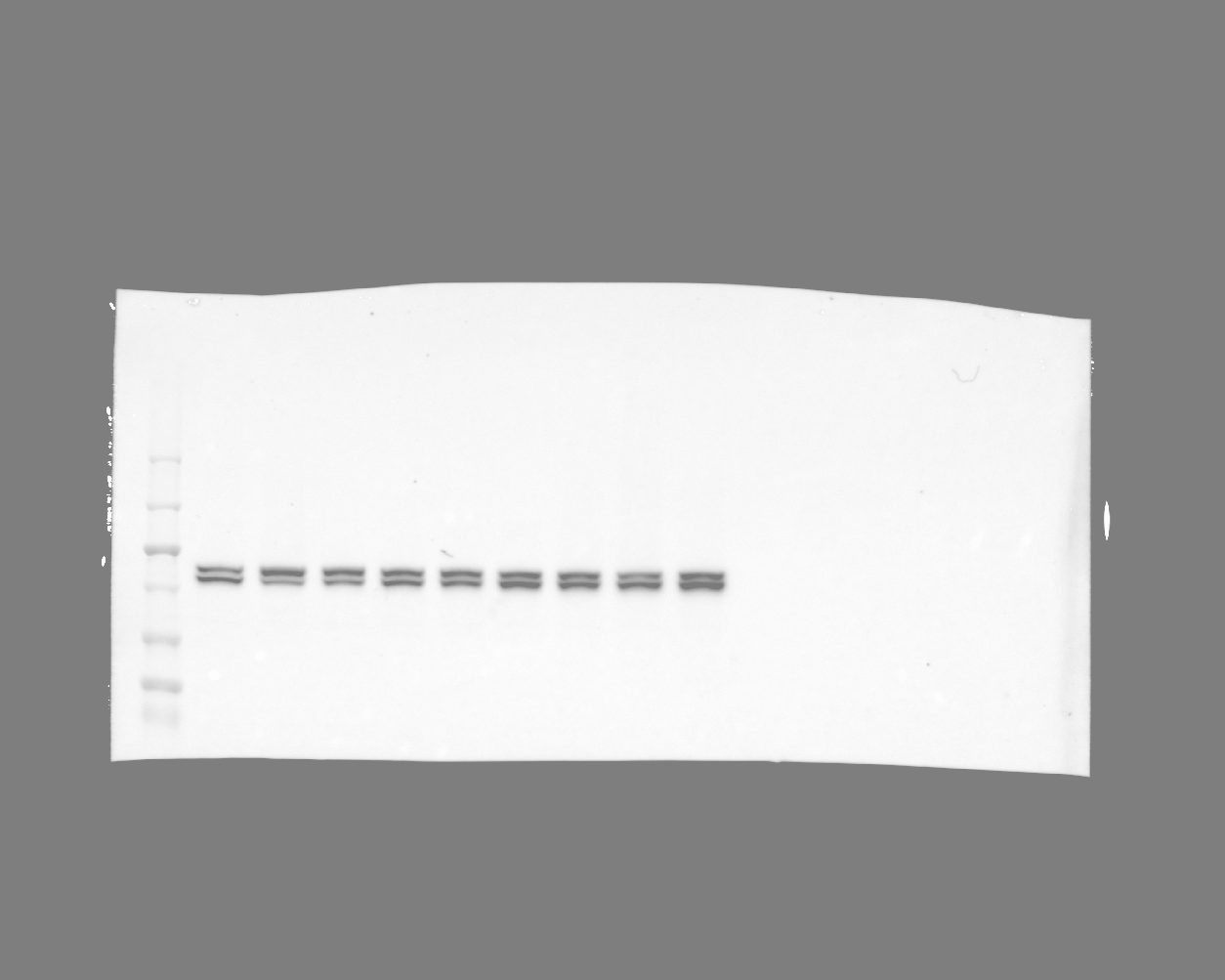

### 20241119_ut7variantgel_gel5_hh3_13(Chemiluminescence).tif

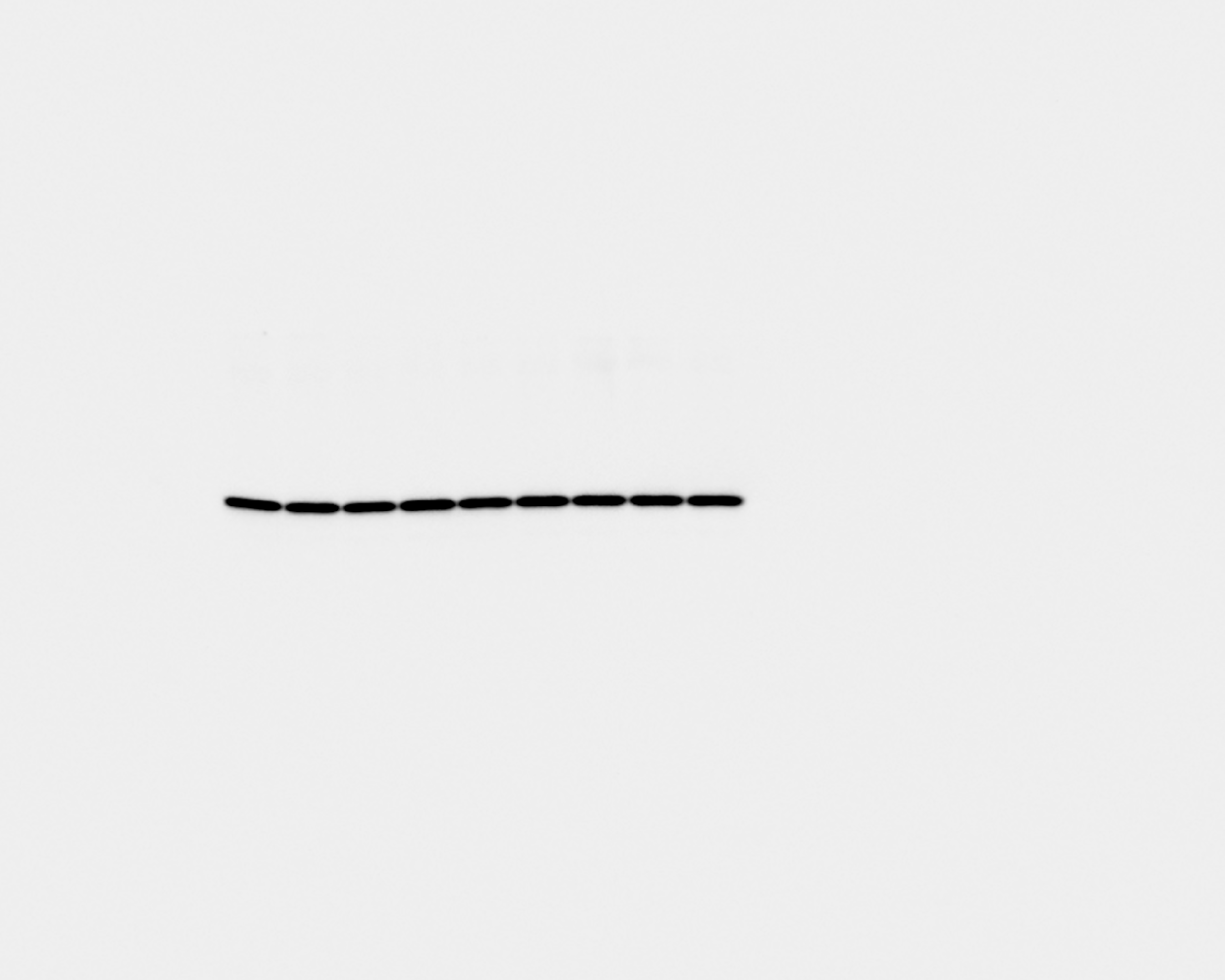

### 20241119_ut7variantgel_gel5_hh3_13(Colorimetric).tif

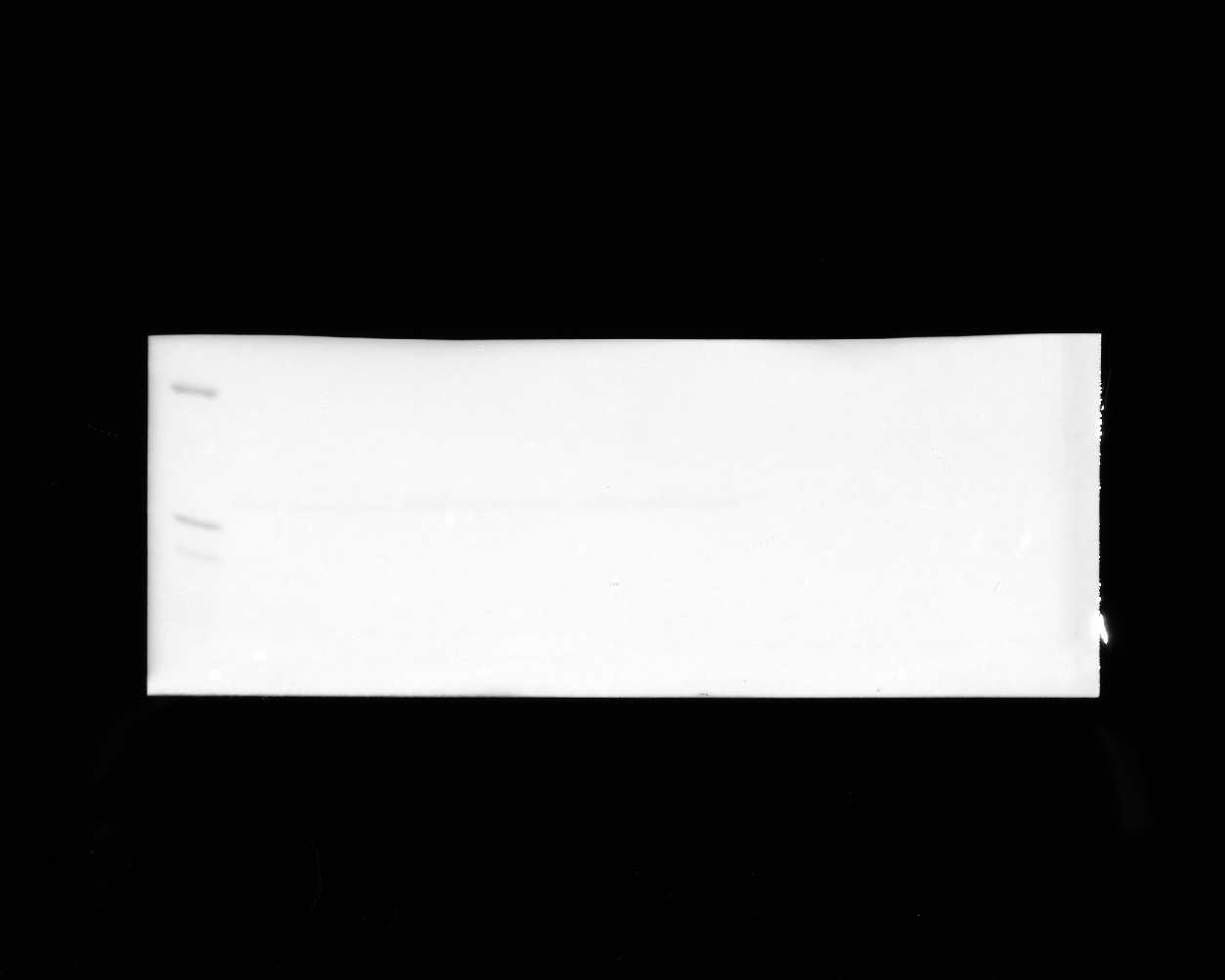

### 20241119_ut7variantgel_gel5_hh3_13(Composite).tif

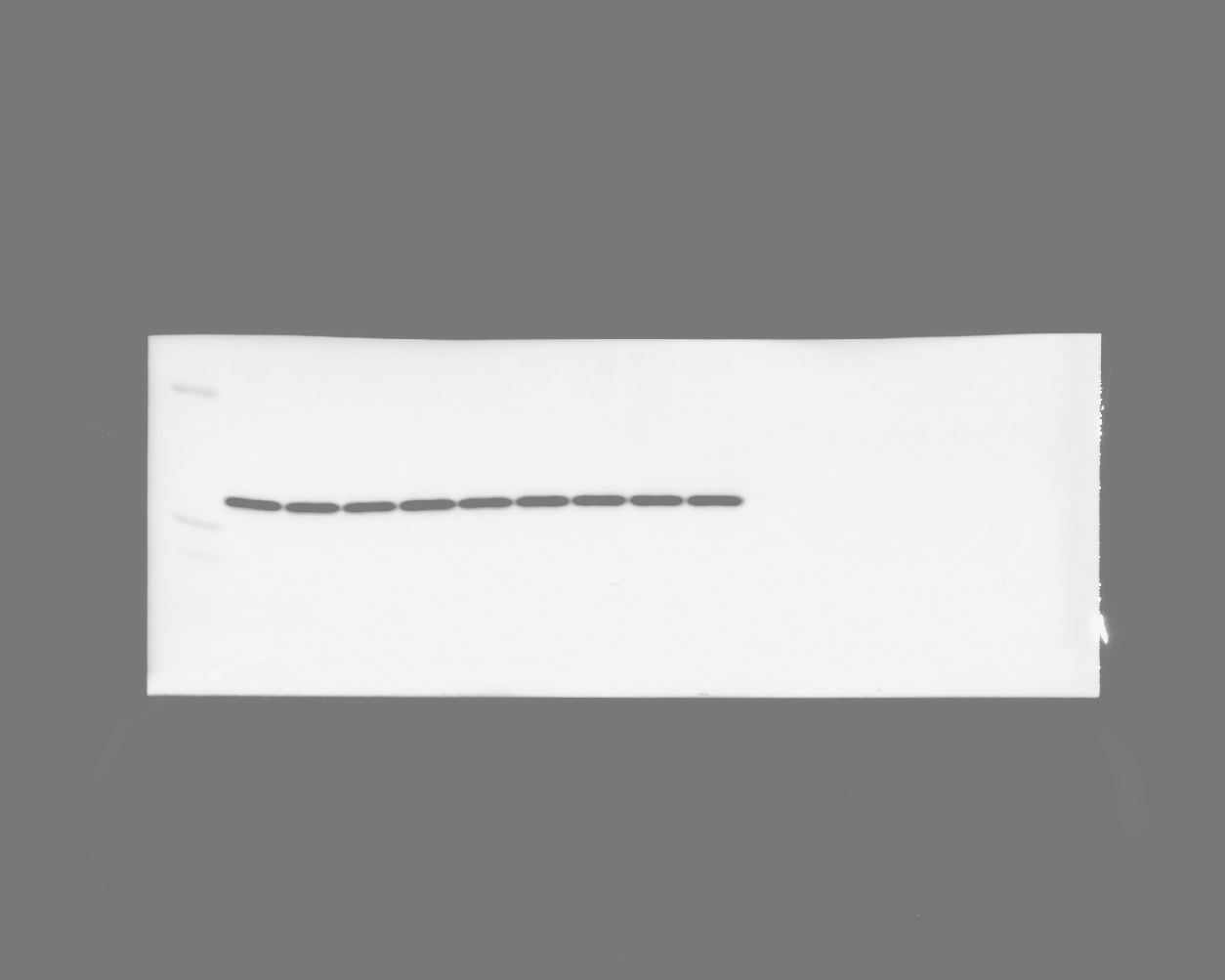

### 20241119_ut7variantgel_gel5_stat5_phospho_16 (Multichannel).tif

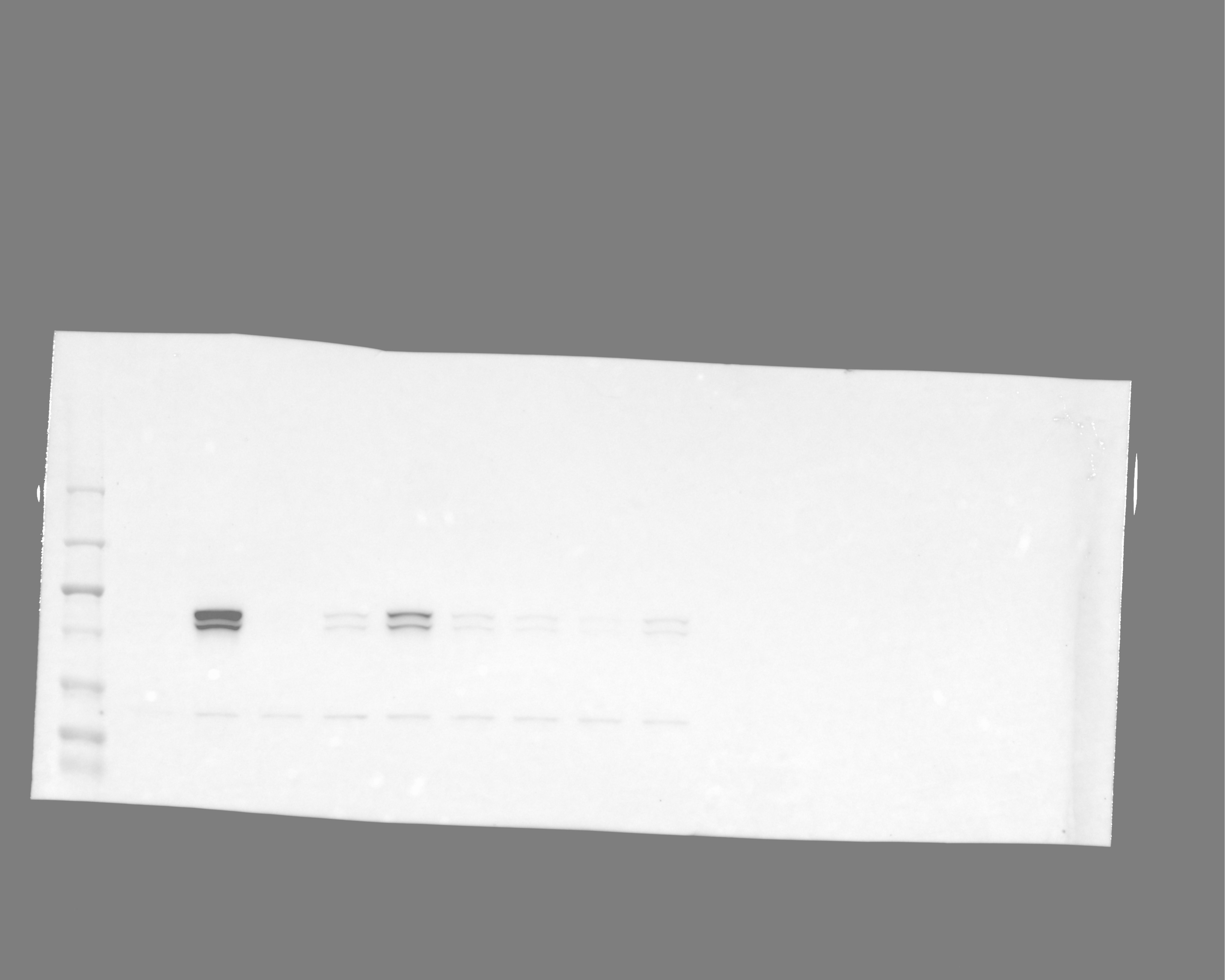

### 20241119_ut7variantgel_gel5_stat5_phospho_16(Chemiluminescence).tif

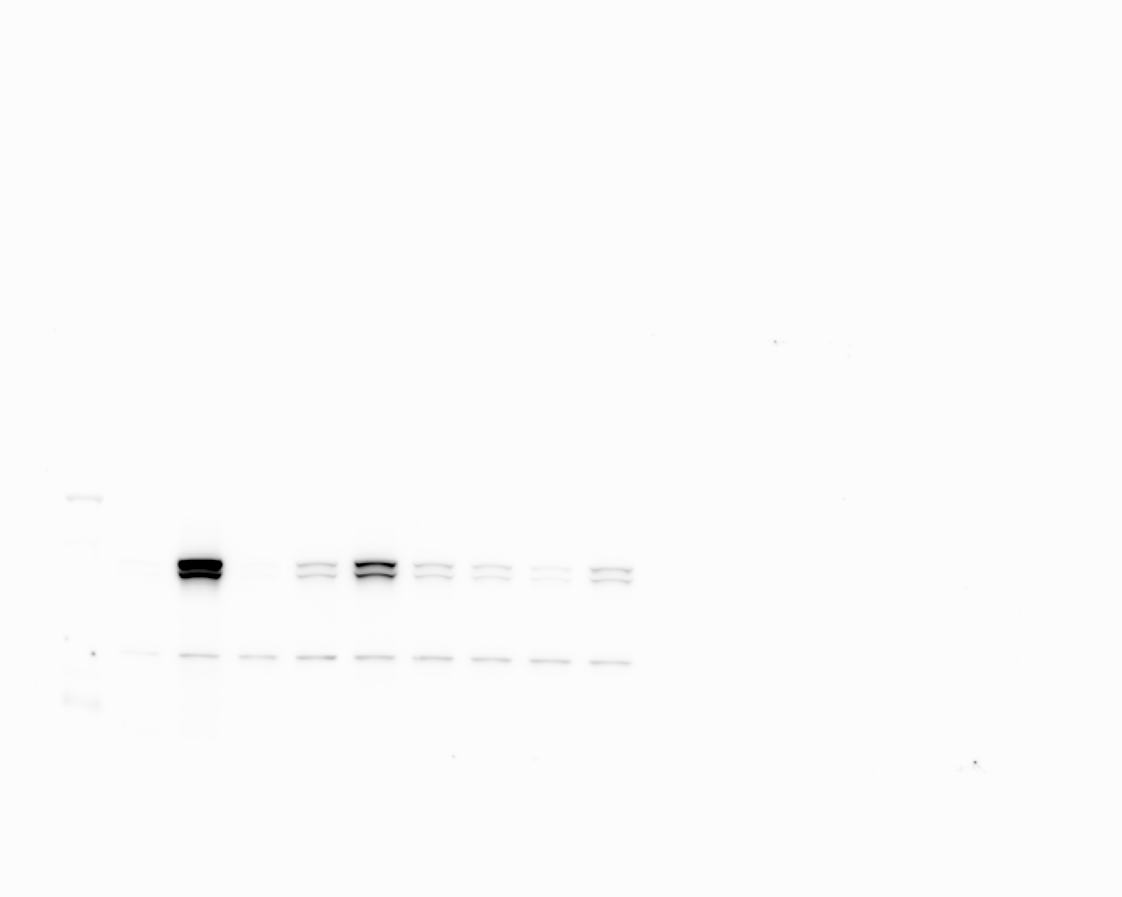

### 20241119_ut7variantgel_gel5_stat5_phospho_16(Colorimetric).tif

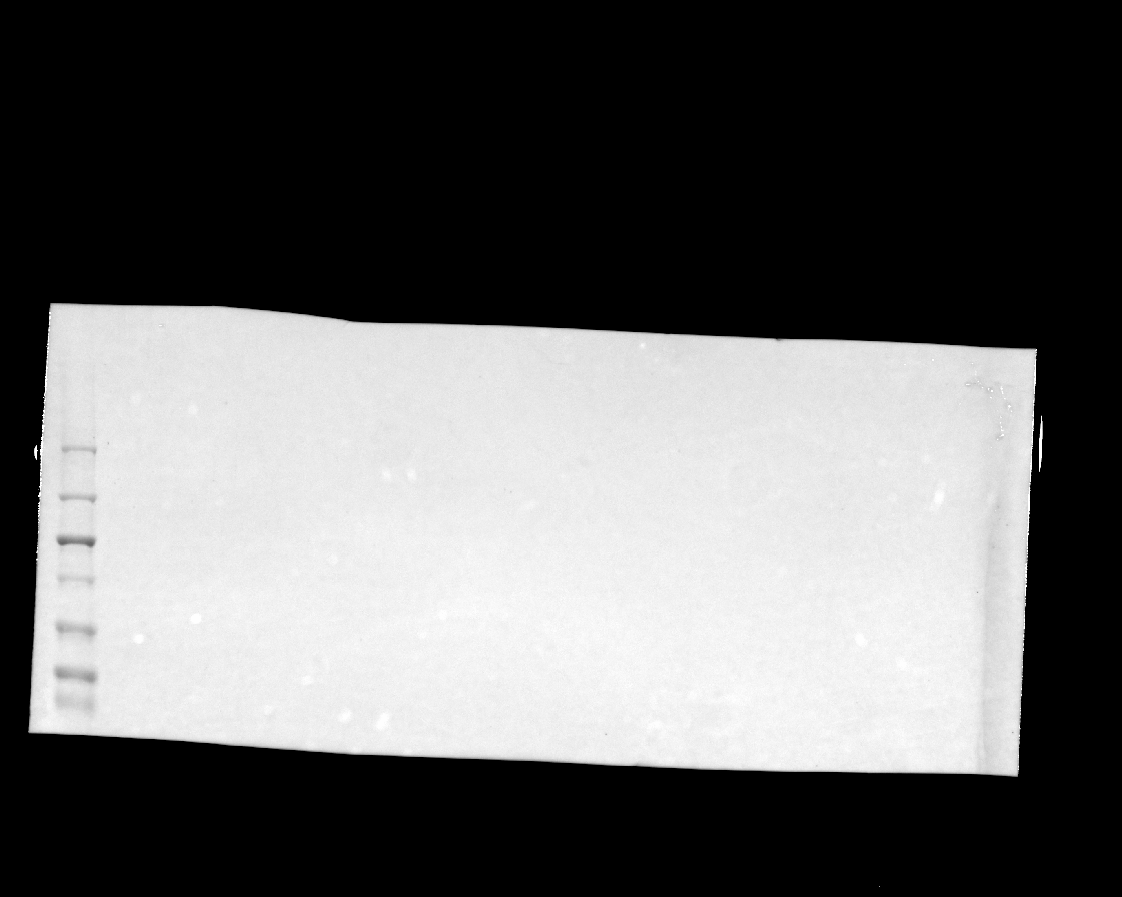

### 20241119_ut7variantgel_gel6_hh3_7(Chemiluminescence).tif

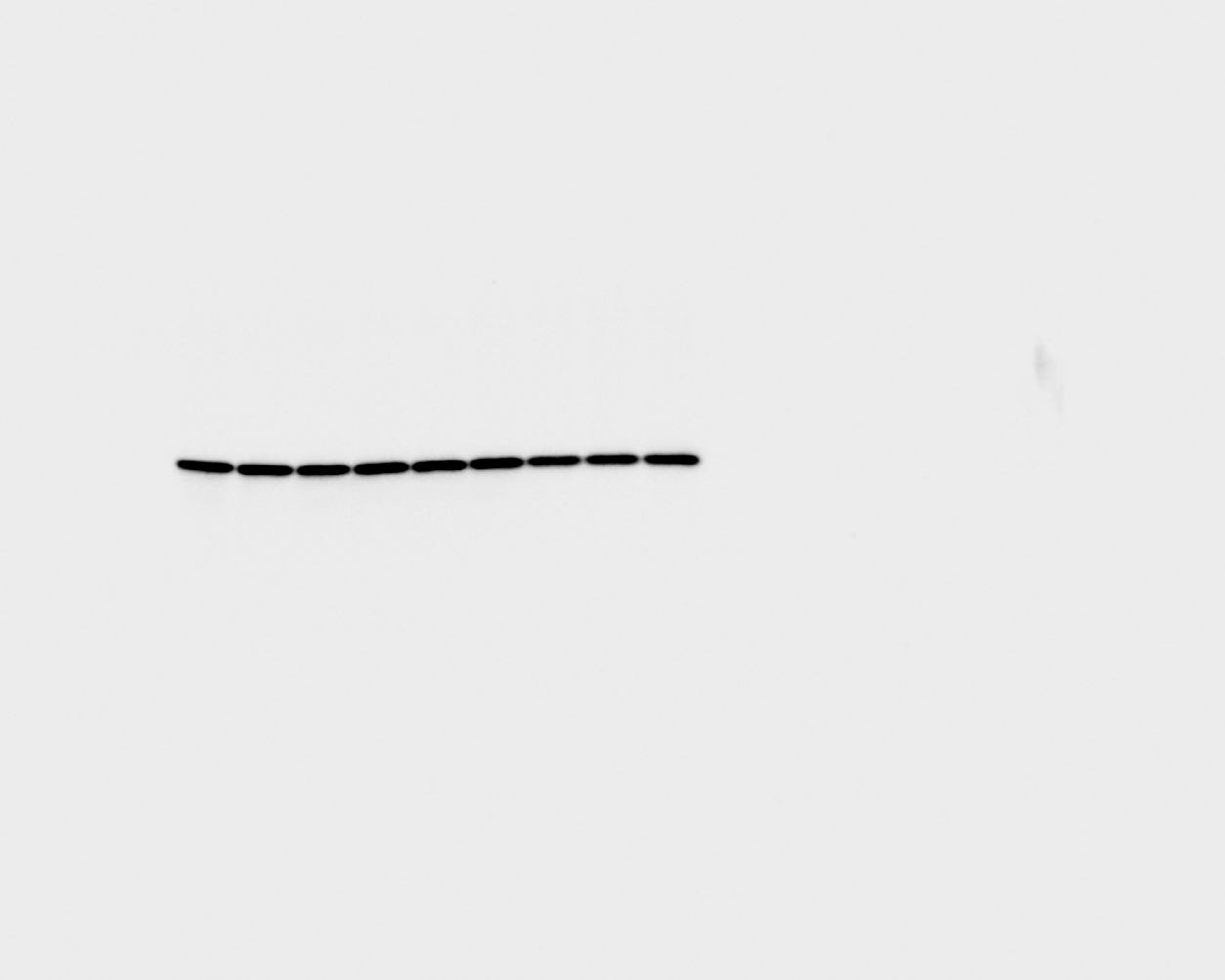

### 20241119_ut7variantgel_gel6_hh3_7(Colorimetric).tif

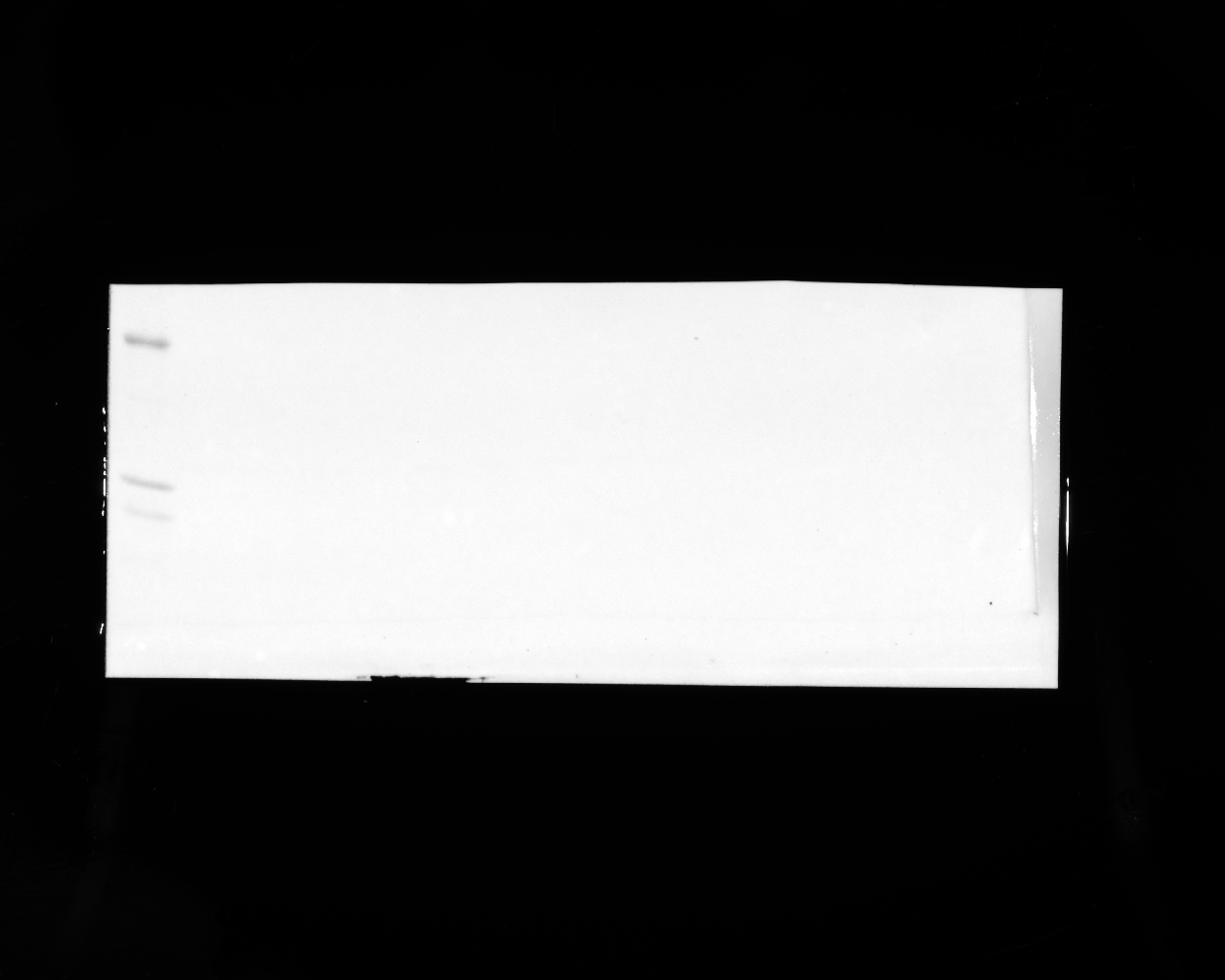

### 20241119_ut7variantgel_gel6_hh3_7(Composite).tif

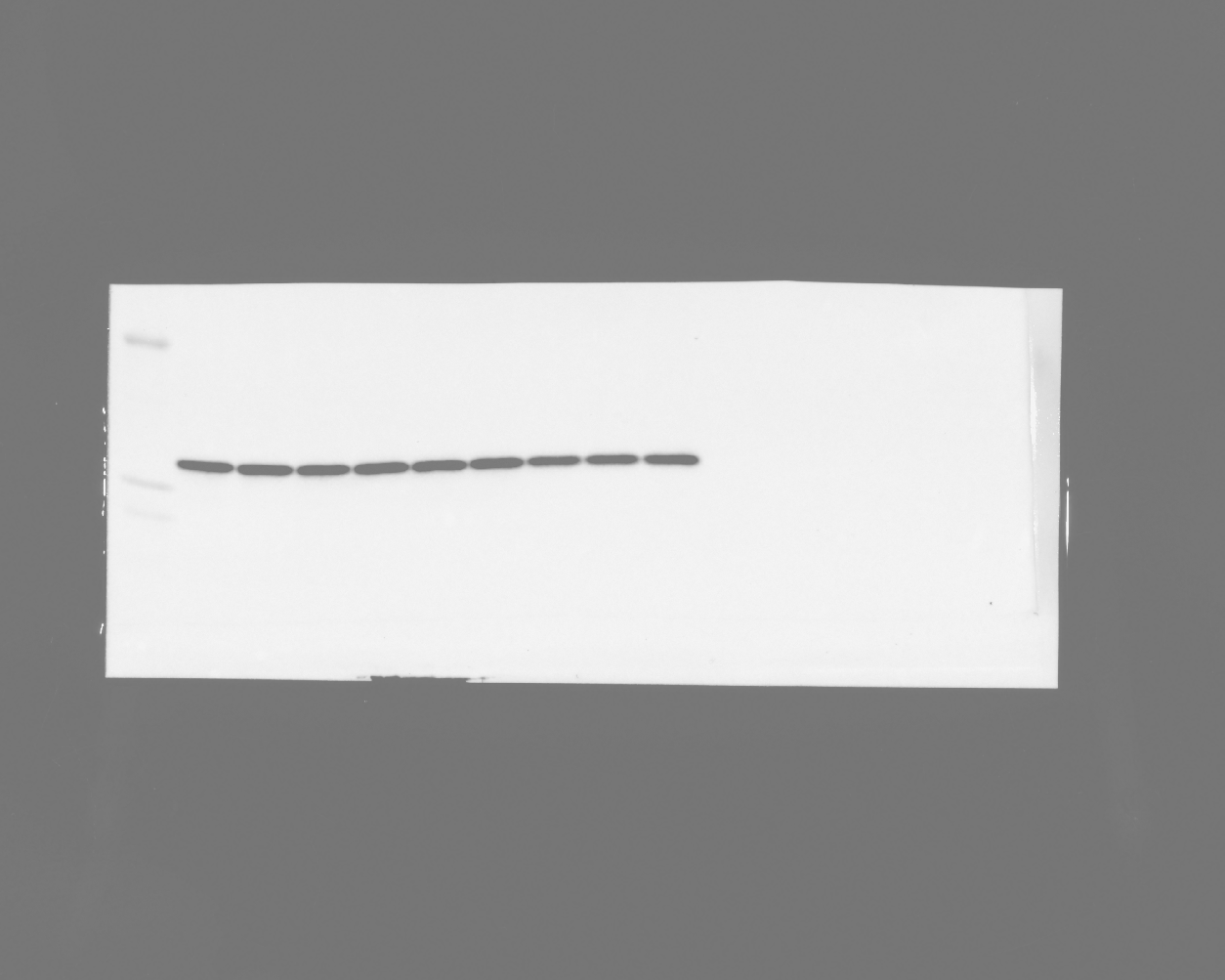

### 20241119_ut7variantgel_gel6_stat5_8 (Multichannel)_usedinpaper.tif

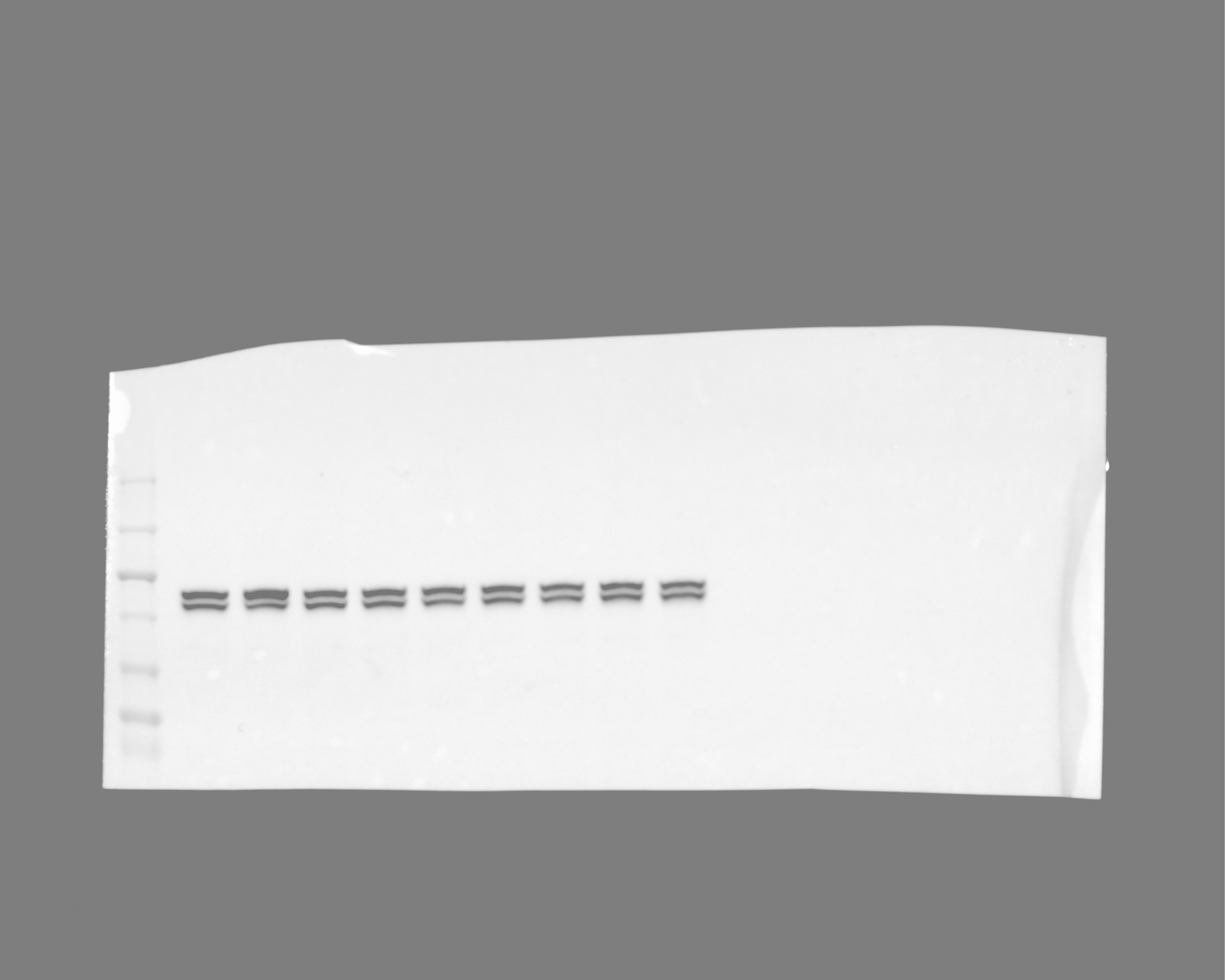

### 20241119_ut7variantgel_gel6_stat5_8(Chemiluminescence).tif

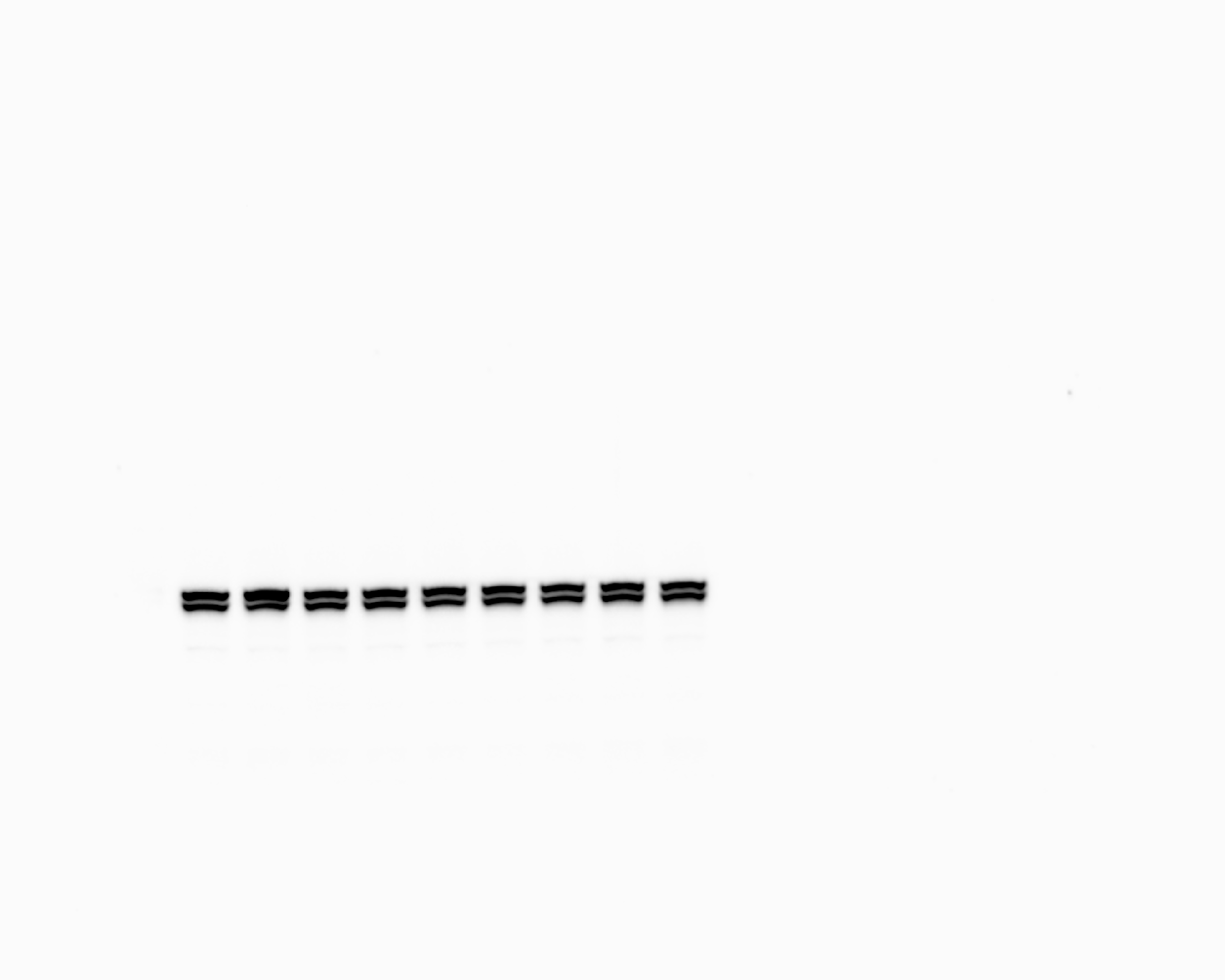

### 20241119_ut7variantgel_gel6_stat5_8(Colorimetric).tif

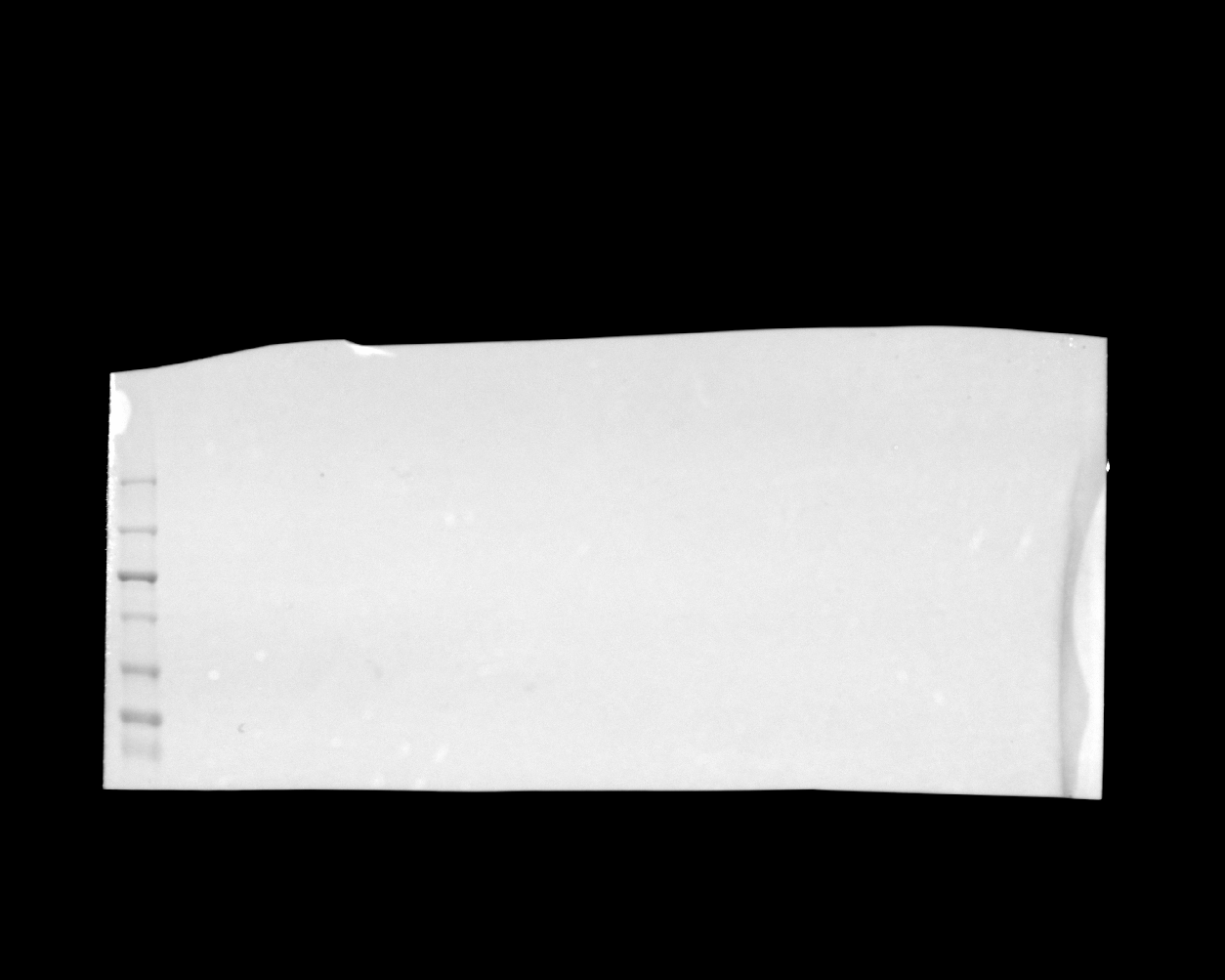
