## Supplementary figures and images for "Rational design of selective bispecific EPO-R/CD131 agonists"

### 20240216_dualscfv_purif_26_his_2 (Multichannel)_final.tif

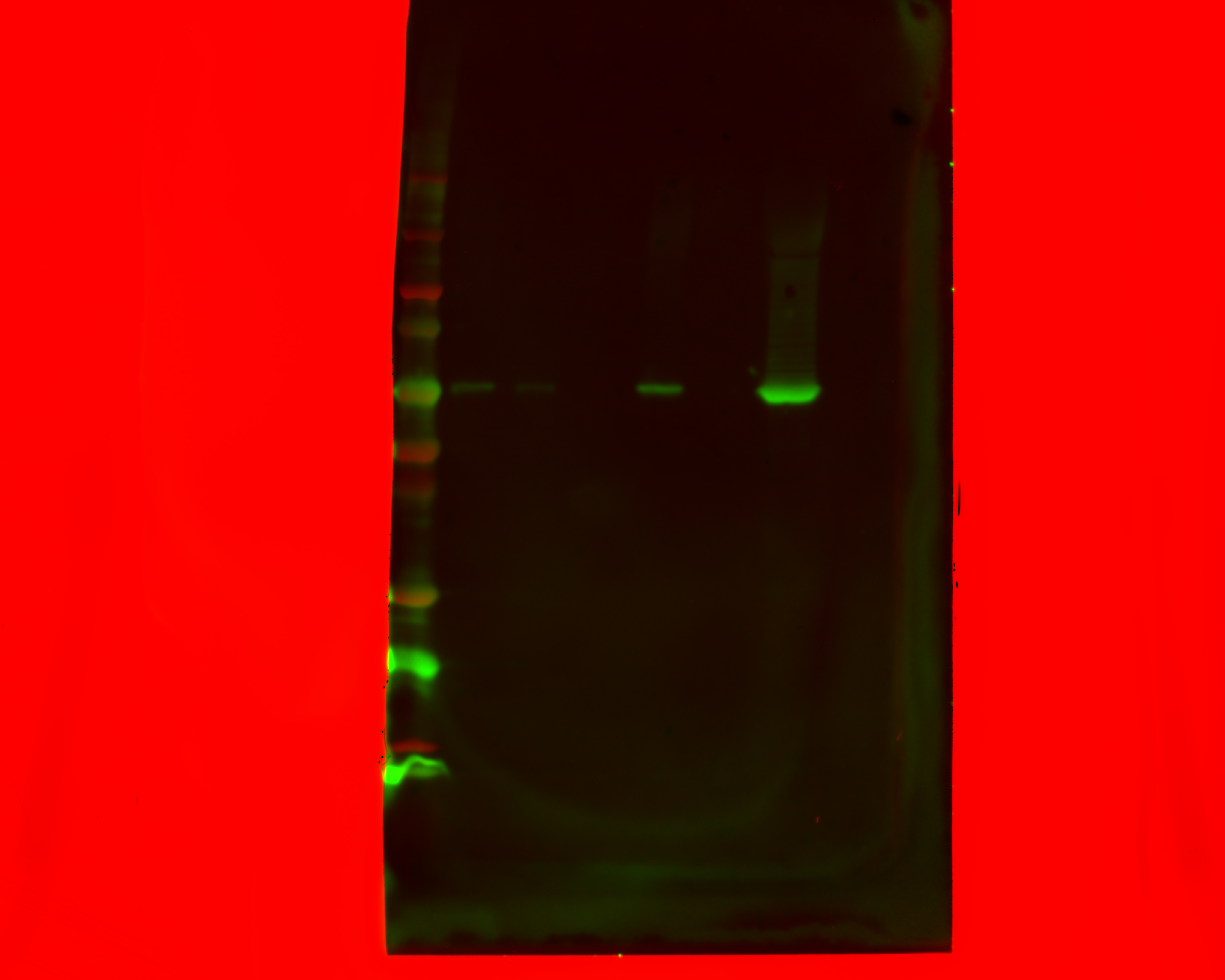

### 20240216_dualscfv_purif_26_his_2(Chemiluminescence).tif

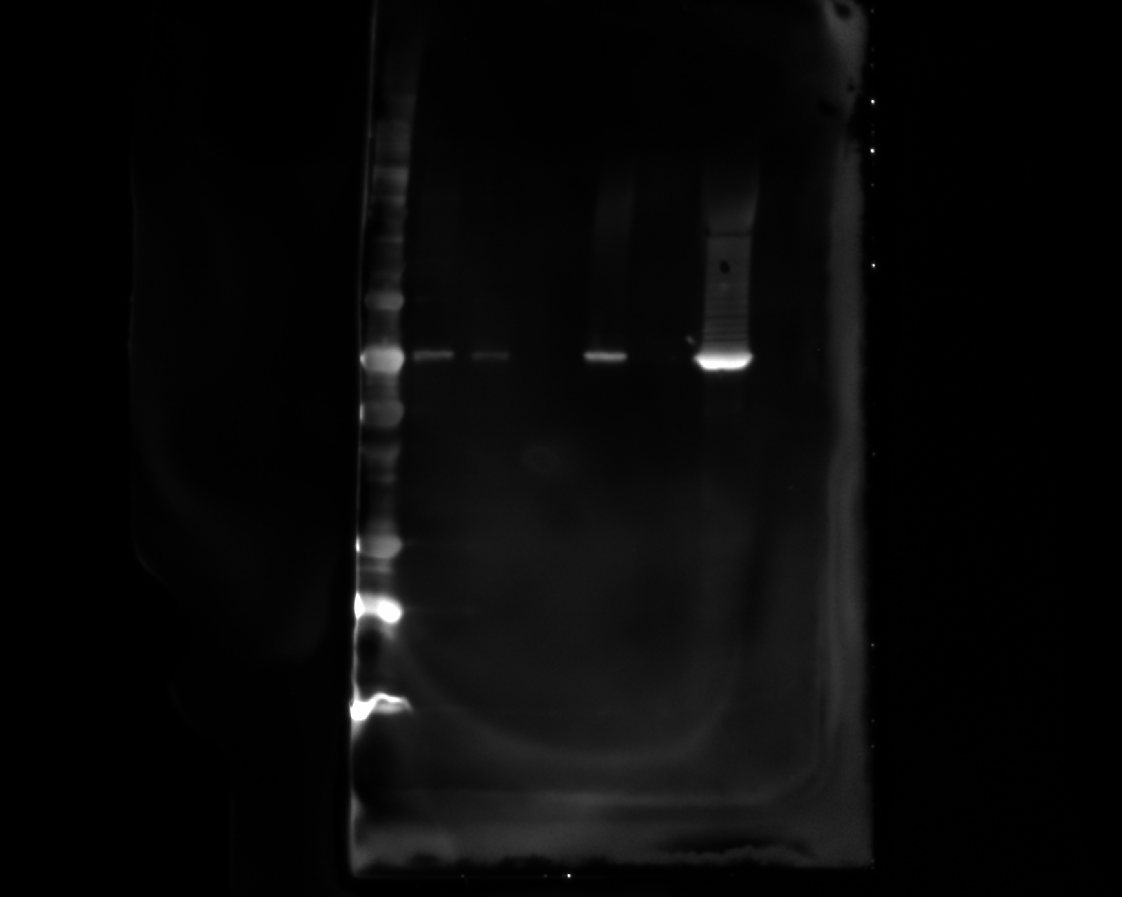

### 20240216_dualscfv_purif_26_his_2(Colorimetric).tif

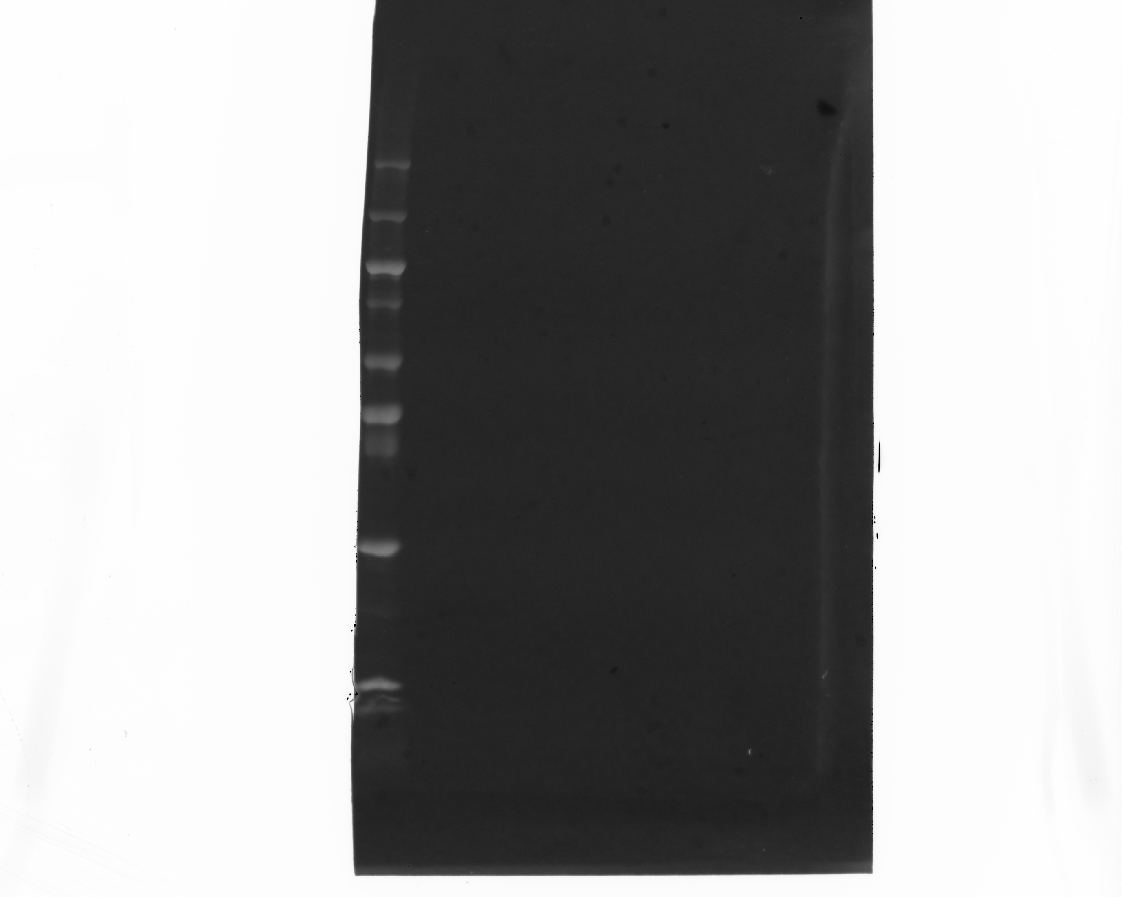

### 20240216_dualscfv_purif_26_his_2(Composite).tif

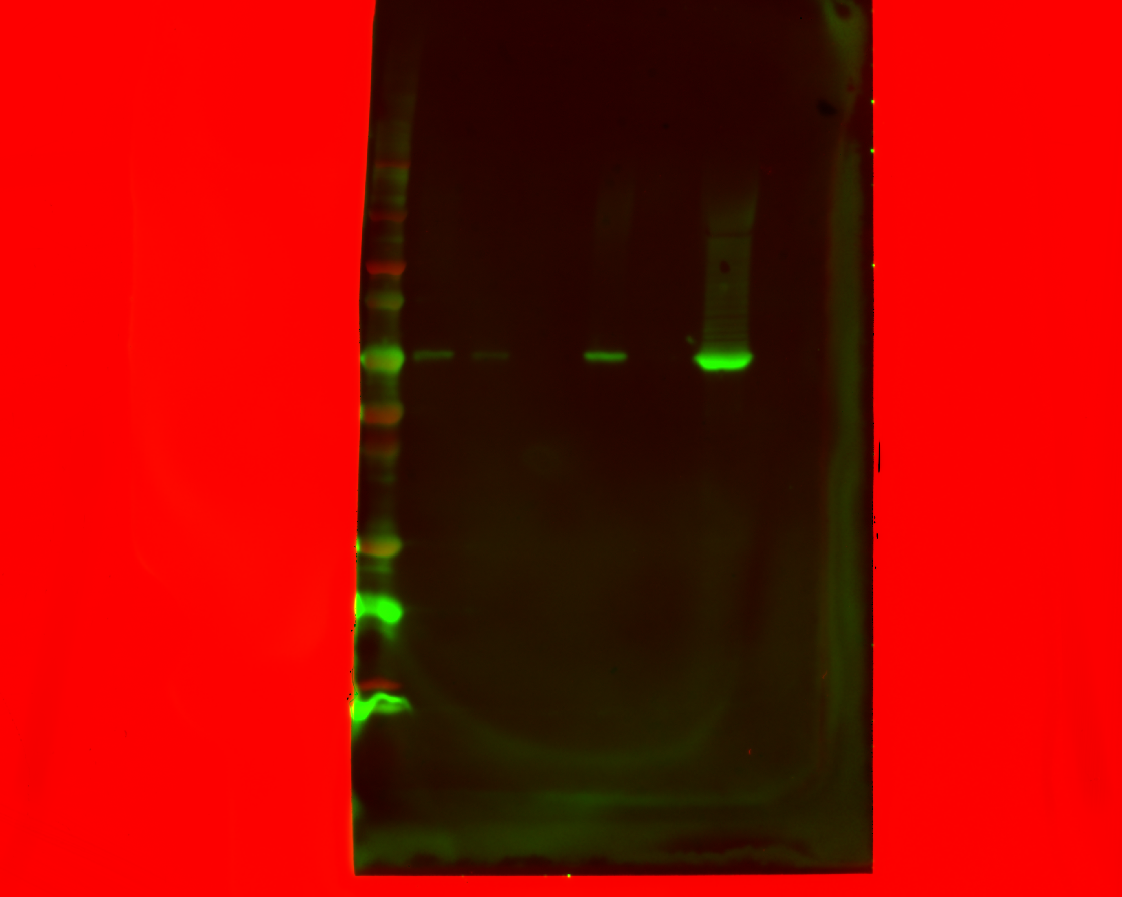

### 20240328_hek_phosphogel_2_gel1_flag_10(Chemiluminescence).tif

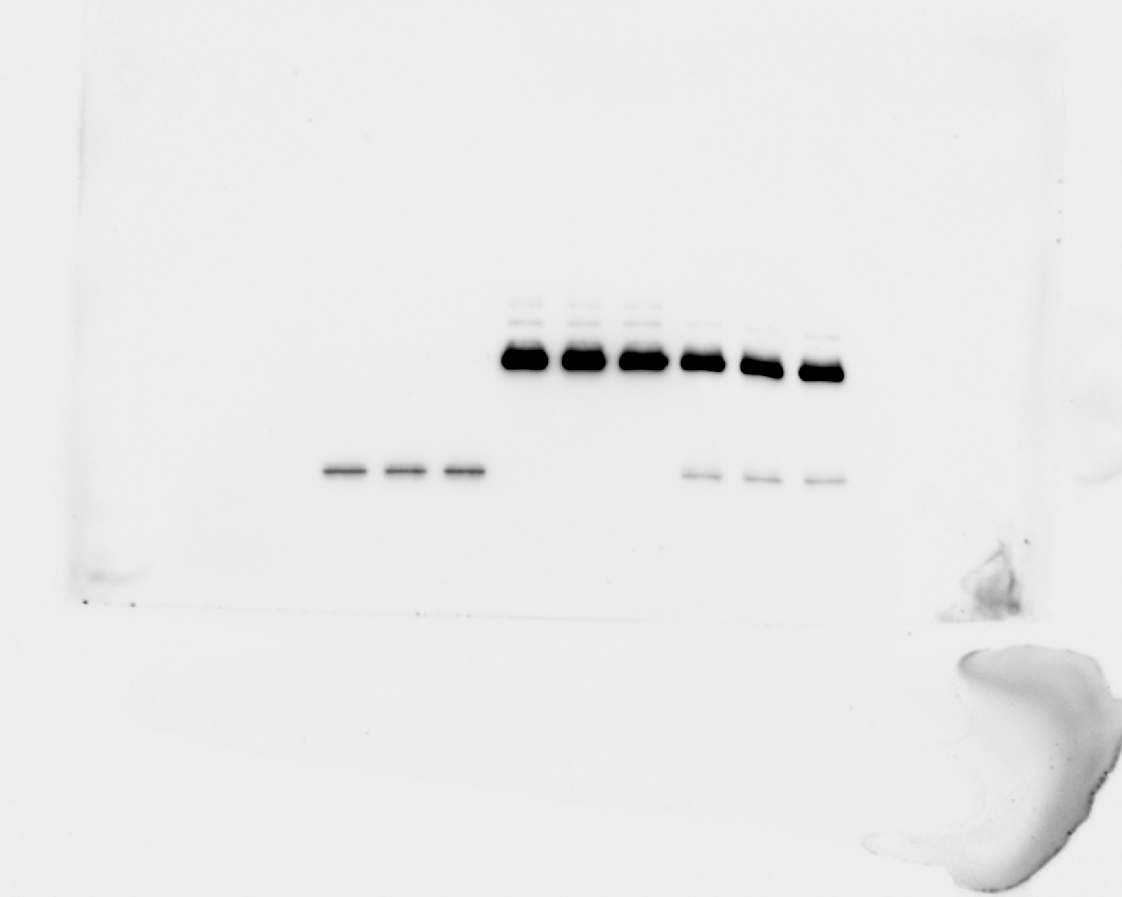

### 20240328_hek_phosphogel_2_gel1_flag_10(Colorimetric).tif

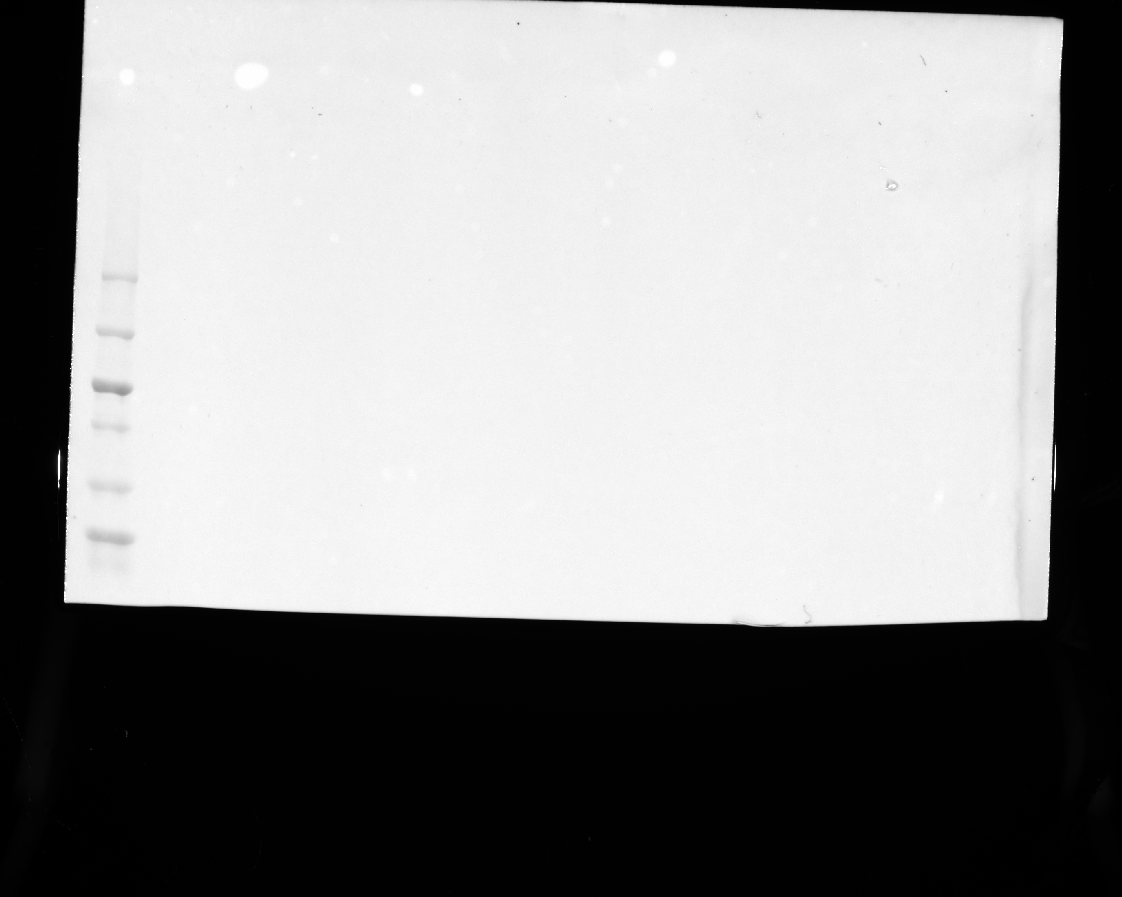

### 20240328_hek_phosphogel_2_gel1_flag_10(Composite).tif

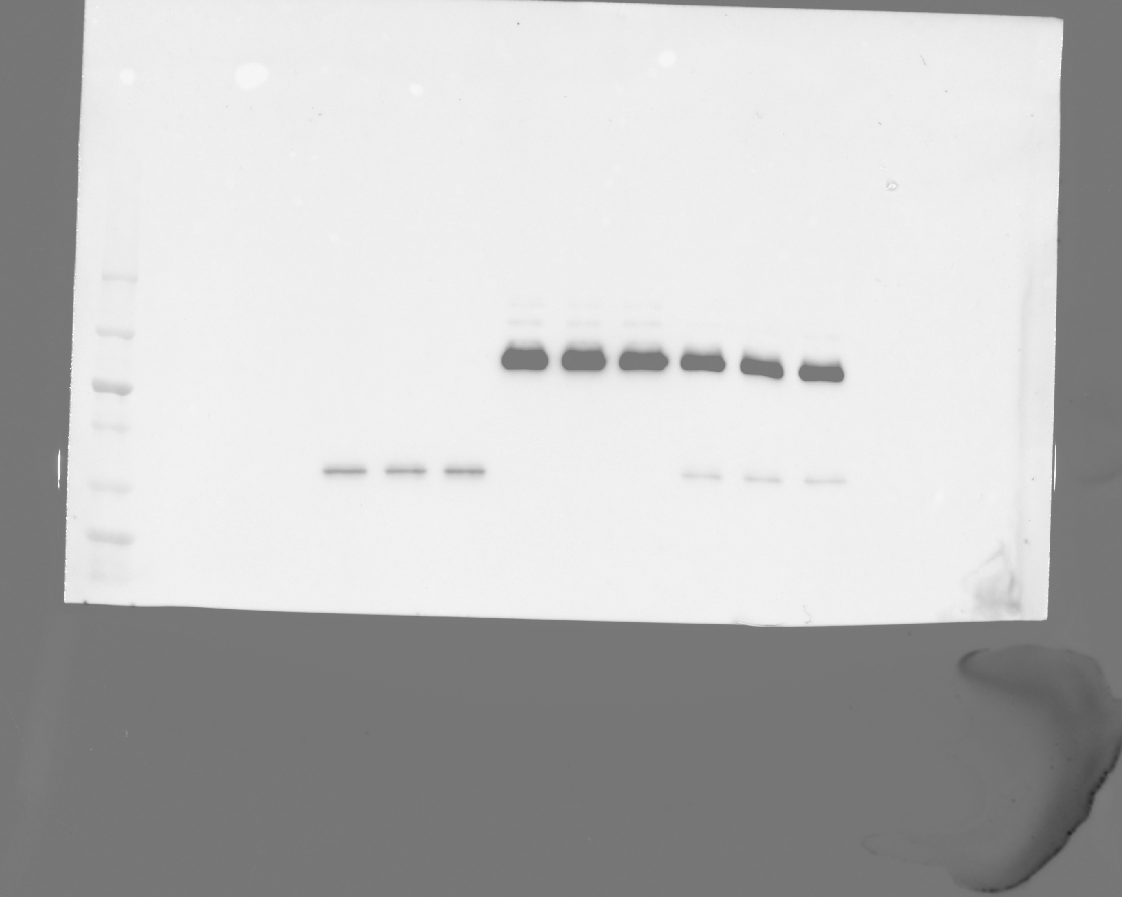

### 20240328_hek_phosphogel_2_gel1_flag_12(Chemiluminescence).tif

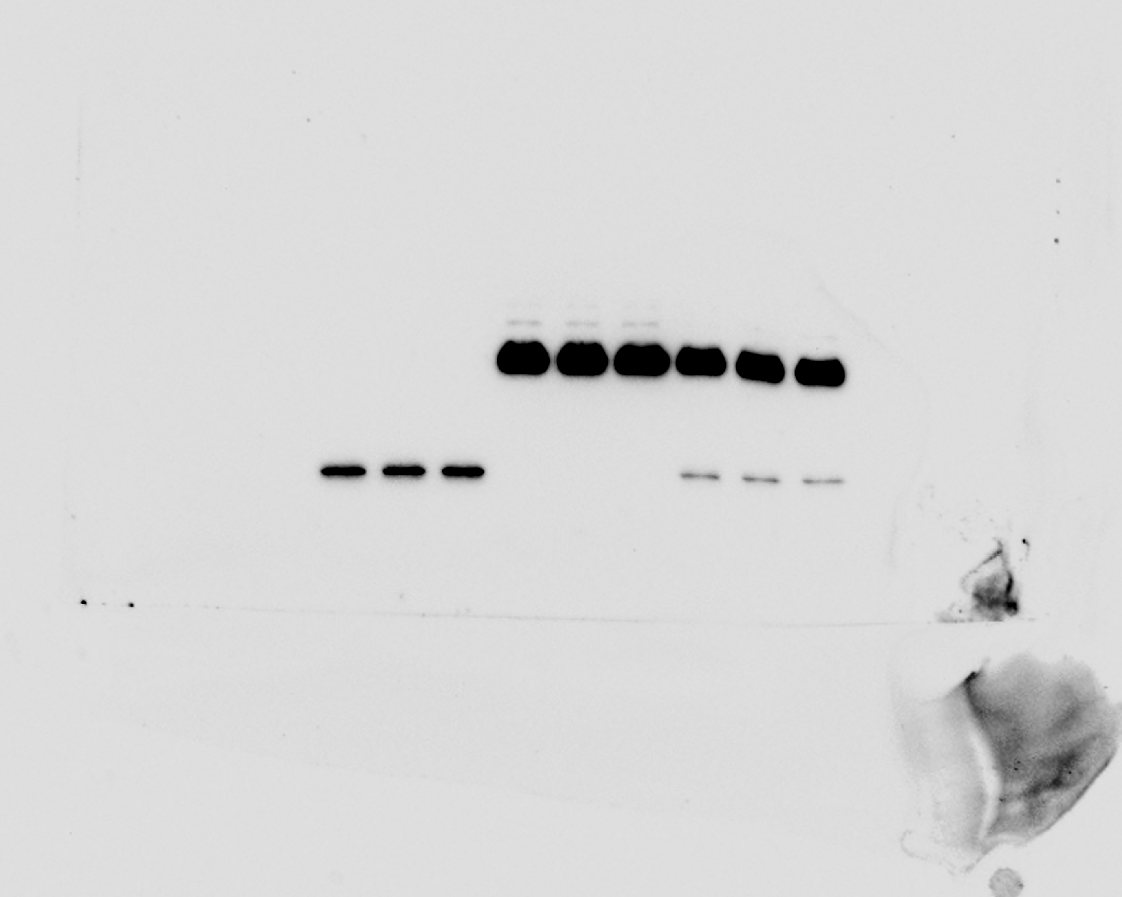

### 20240328_hek_phosphogel_2_gel1_flag_12(Colorimetric).tif

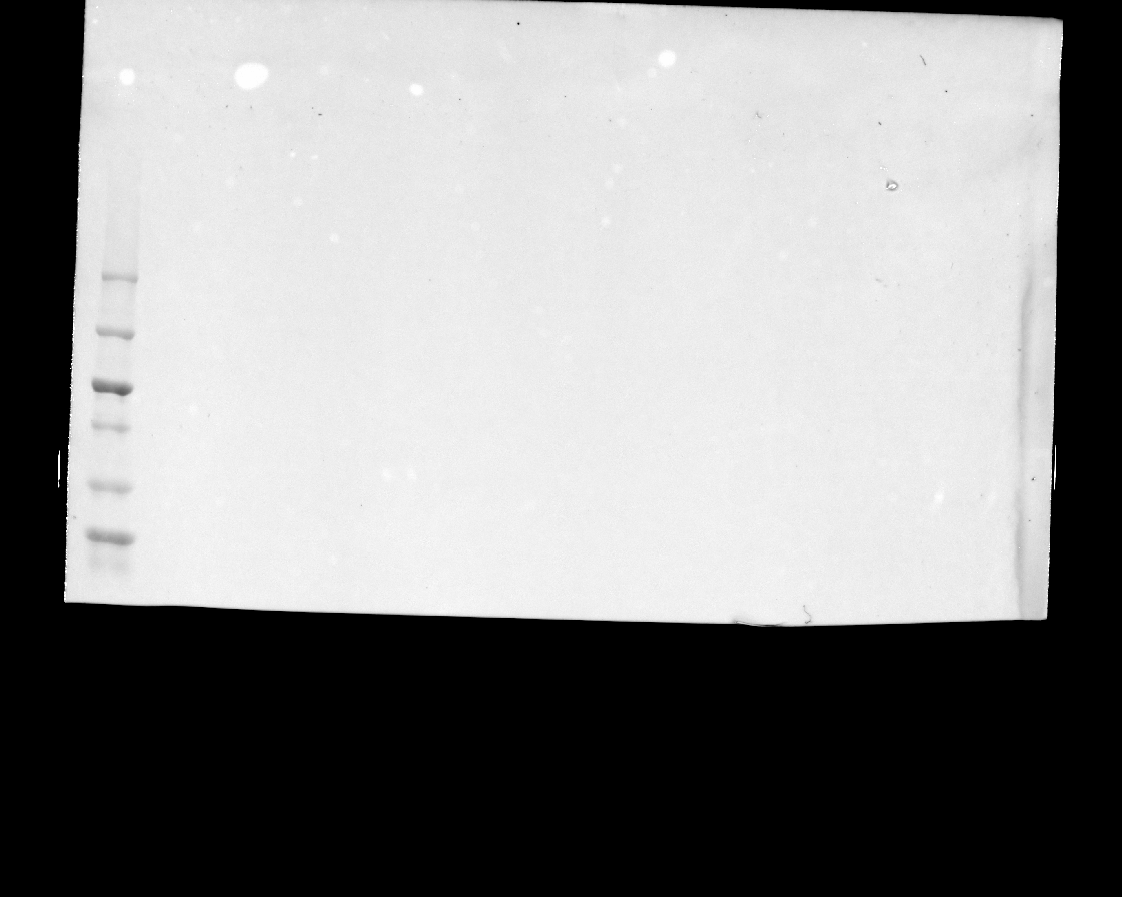

### 20240328_hek_phosphogel_2_gel1_flag_12(Composite).tif

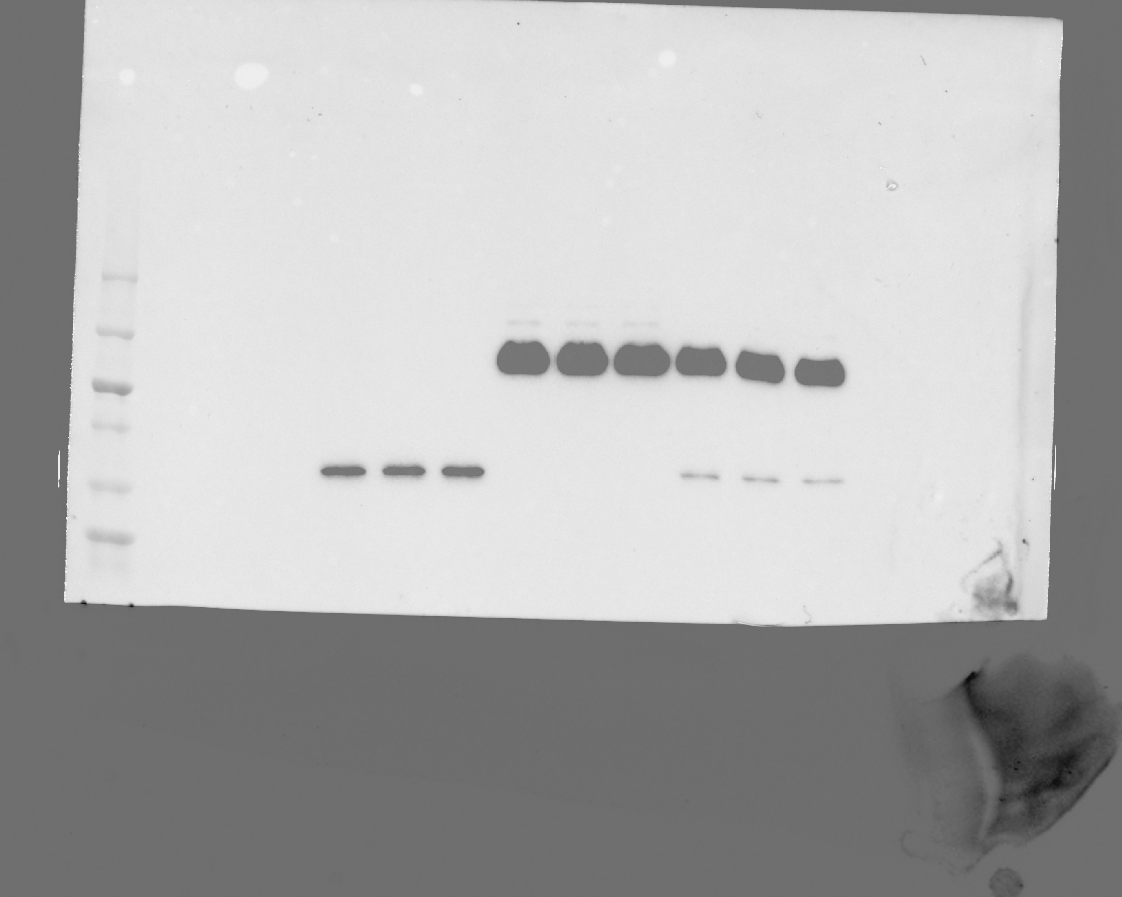

### 20240328_hek_phosphogel_2_gel1_hh3_stat5_10(Chemiluminescence).tif

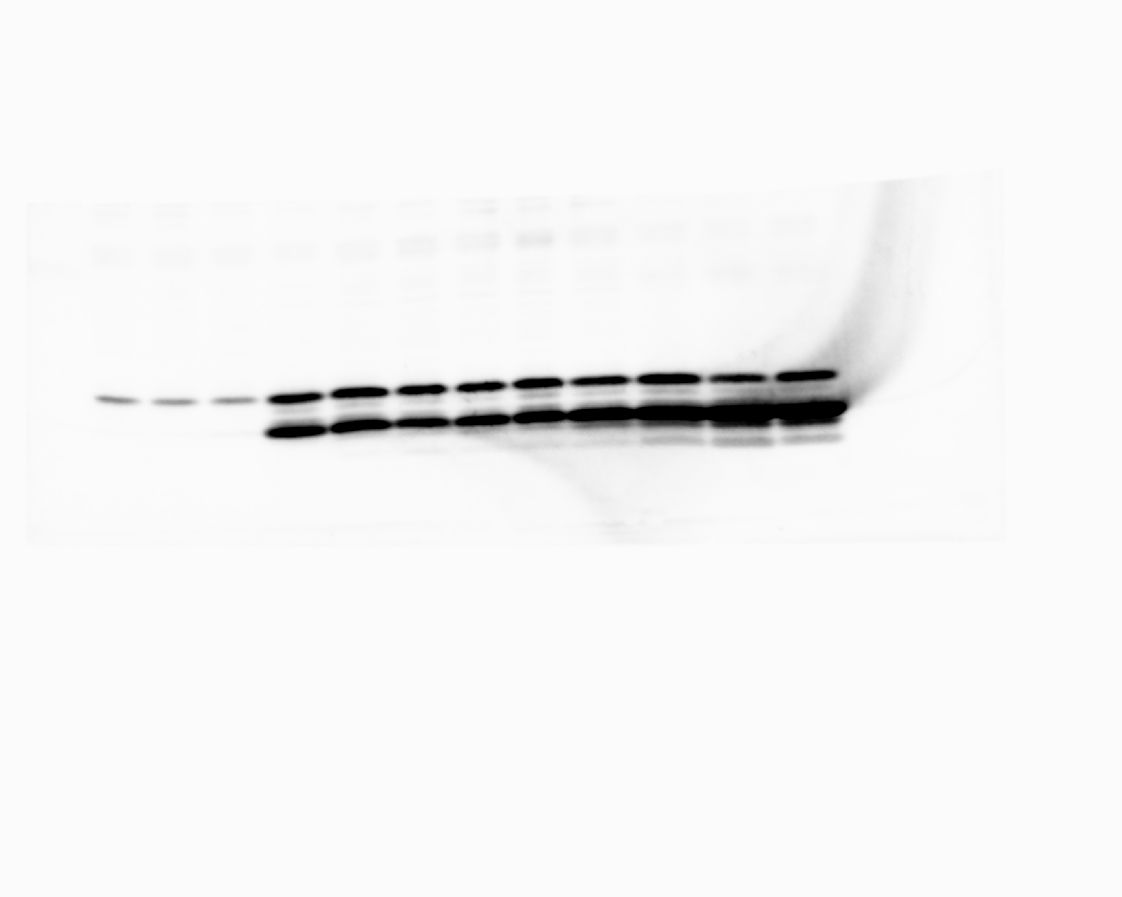

### 20240328_hek_phosphogel_2_gel1_hh3_stat5_10(Colorimetric).tif

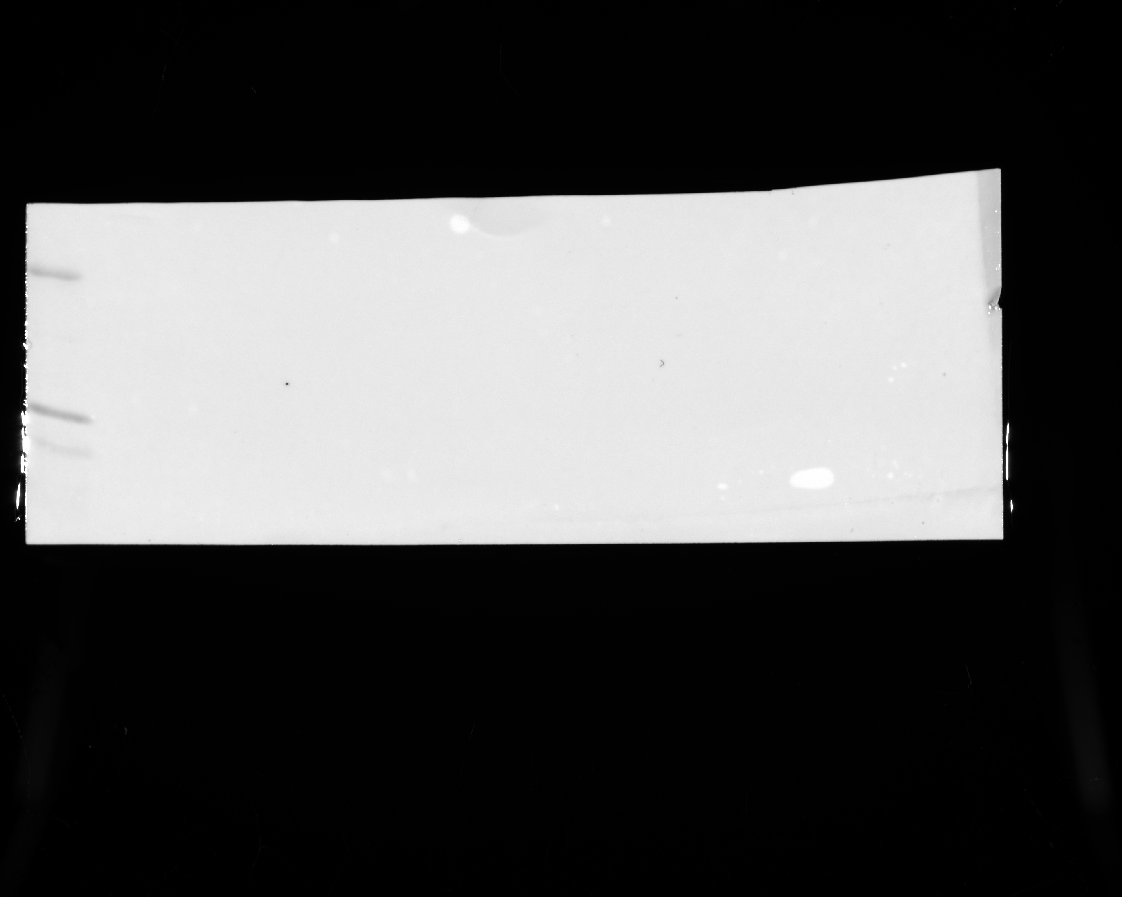

### 20240328_hek_phosphogel_2_gel1_hh3_stat5_10(Composite).tif

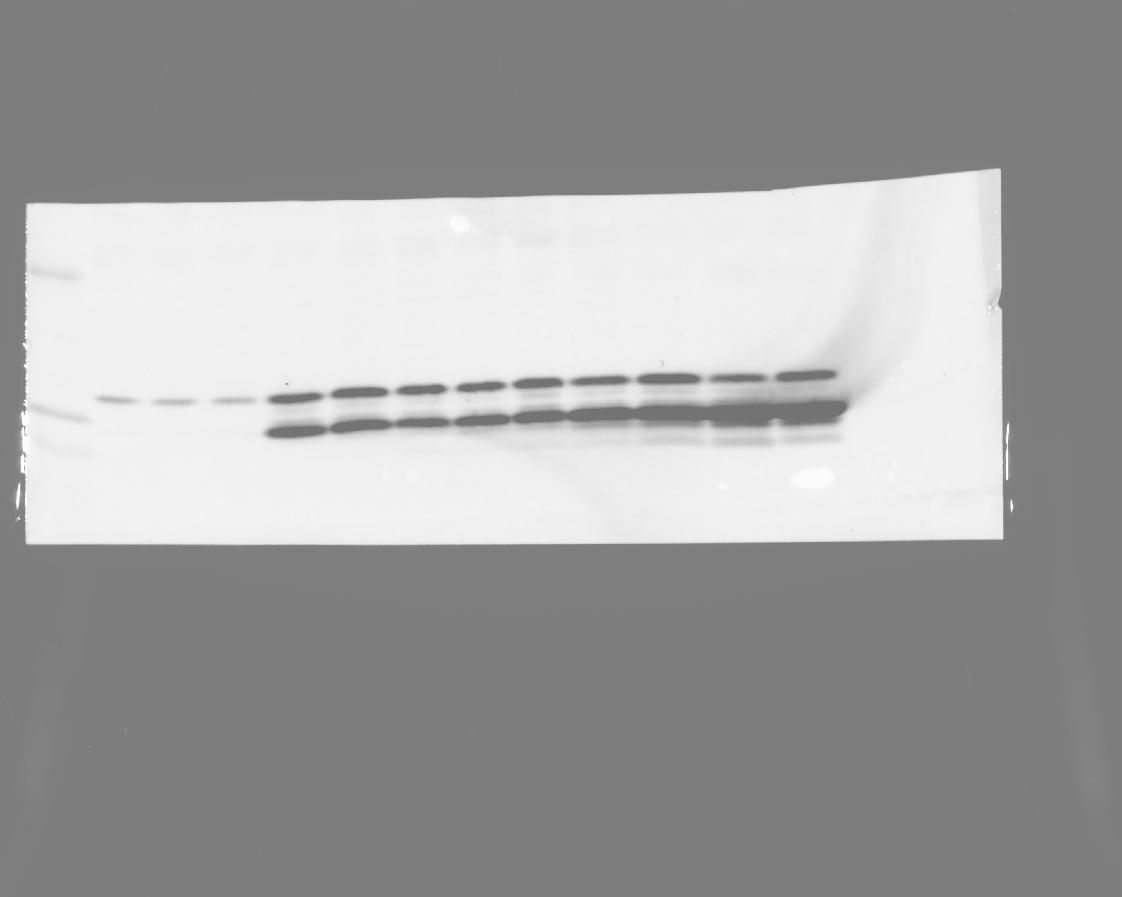

### 20240328_hek_phosphogel_2_gel1_hh3_stat5_phospho_6(Chemiluminescence).tif

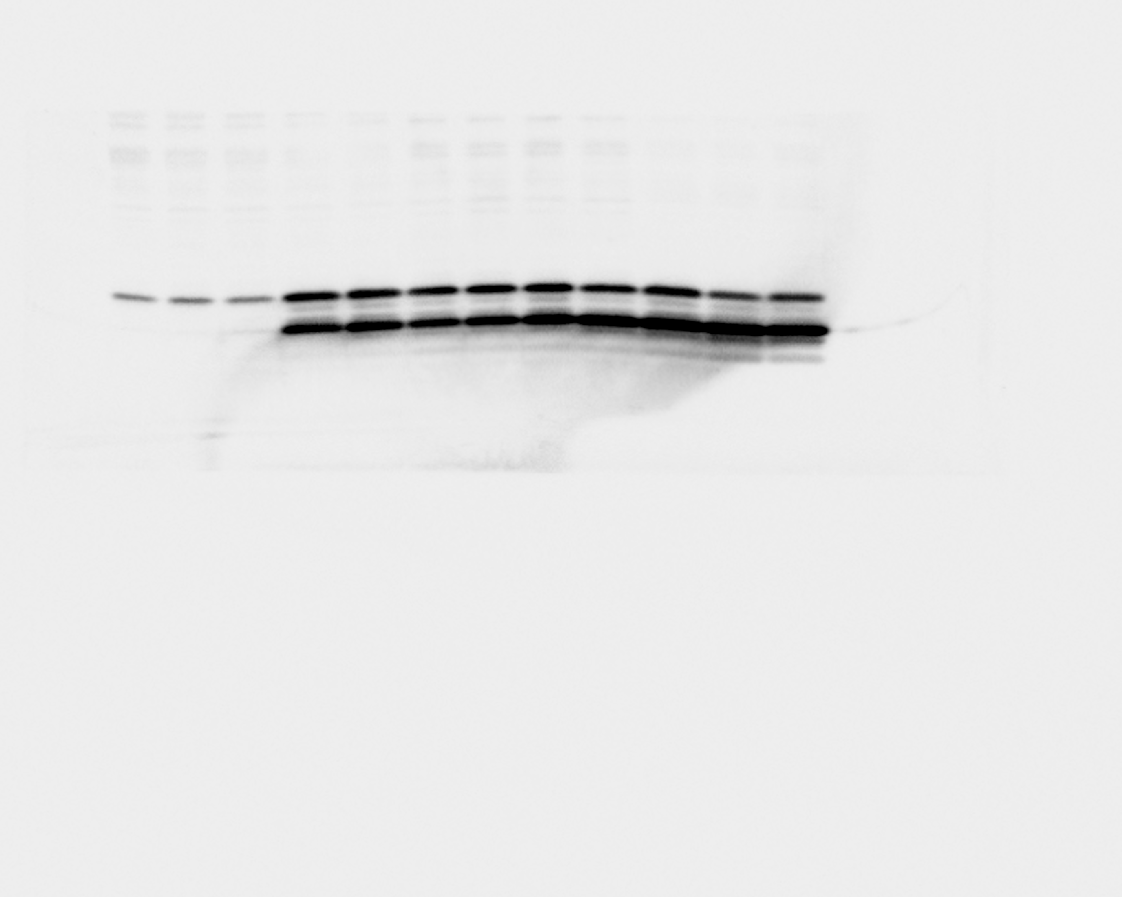

### 20240328_hek_phosphogel_2_gel1_hh3_stat5_phospho_6(Colorimetric).tif

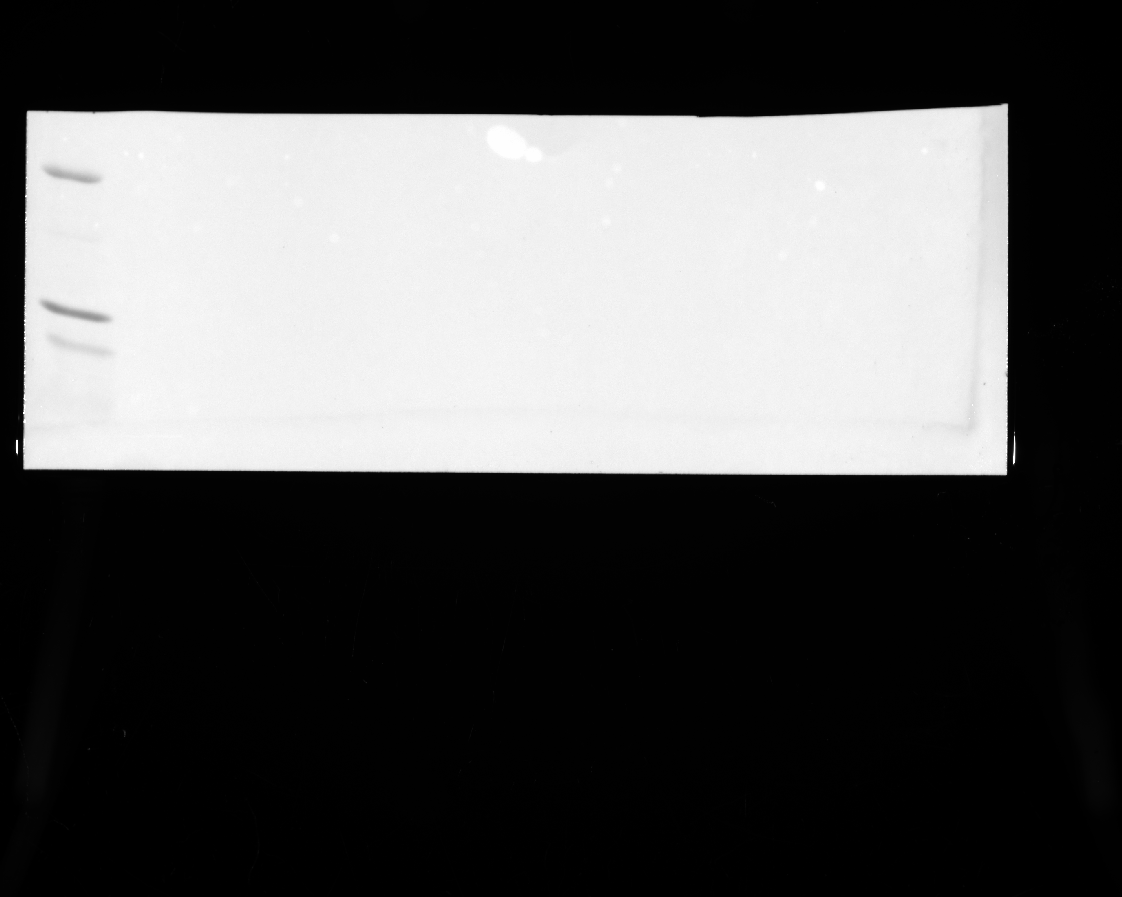

### 20240328_hek_phosphogel_2_gel1_hh3_stat5_phospho_6(Composite).tif

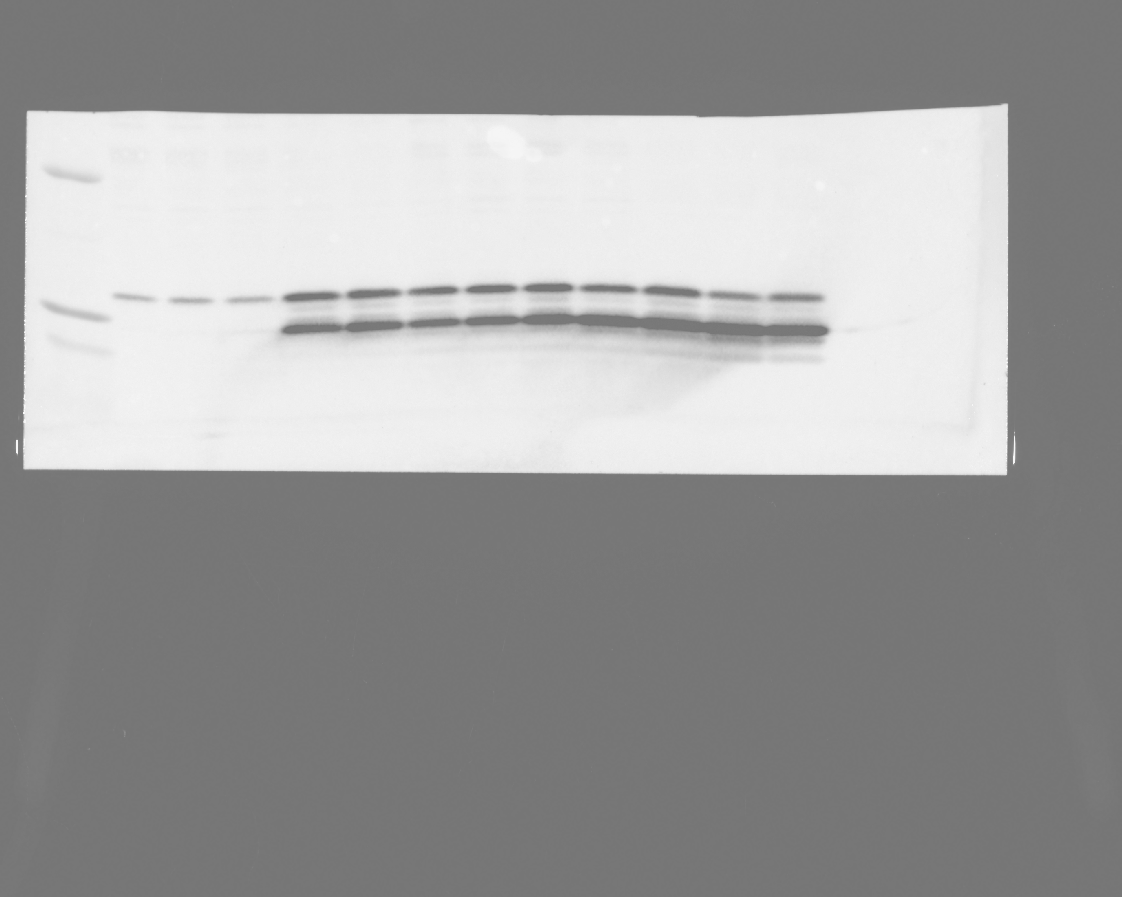

### 20240328_hek_phosphogel_2_gel1_stat5_12(Chemiluminescence).tif

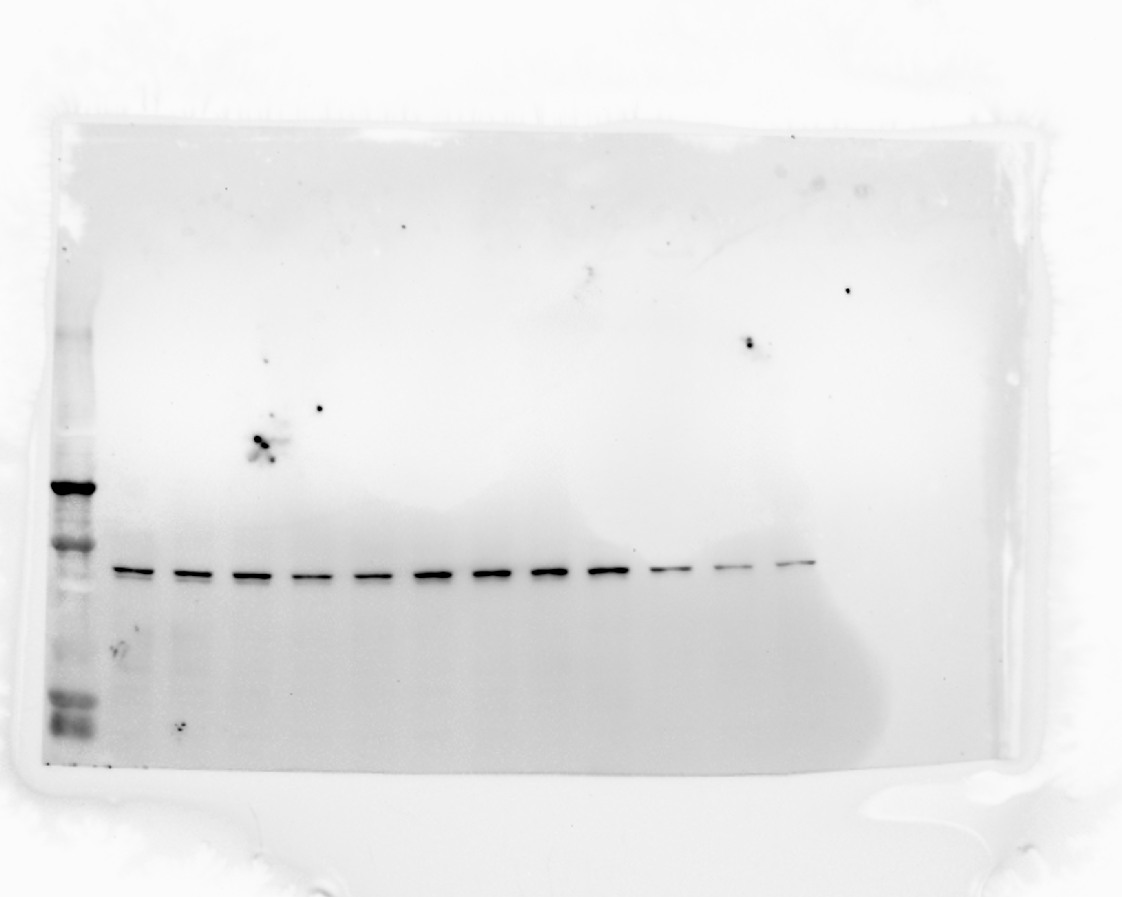

### 20240328_hek_phosphogel_2_gel1_stat5_12(Colorimetric).tif

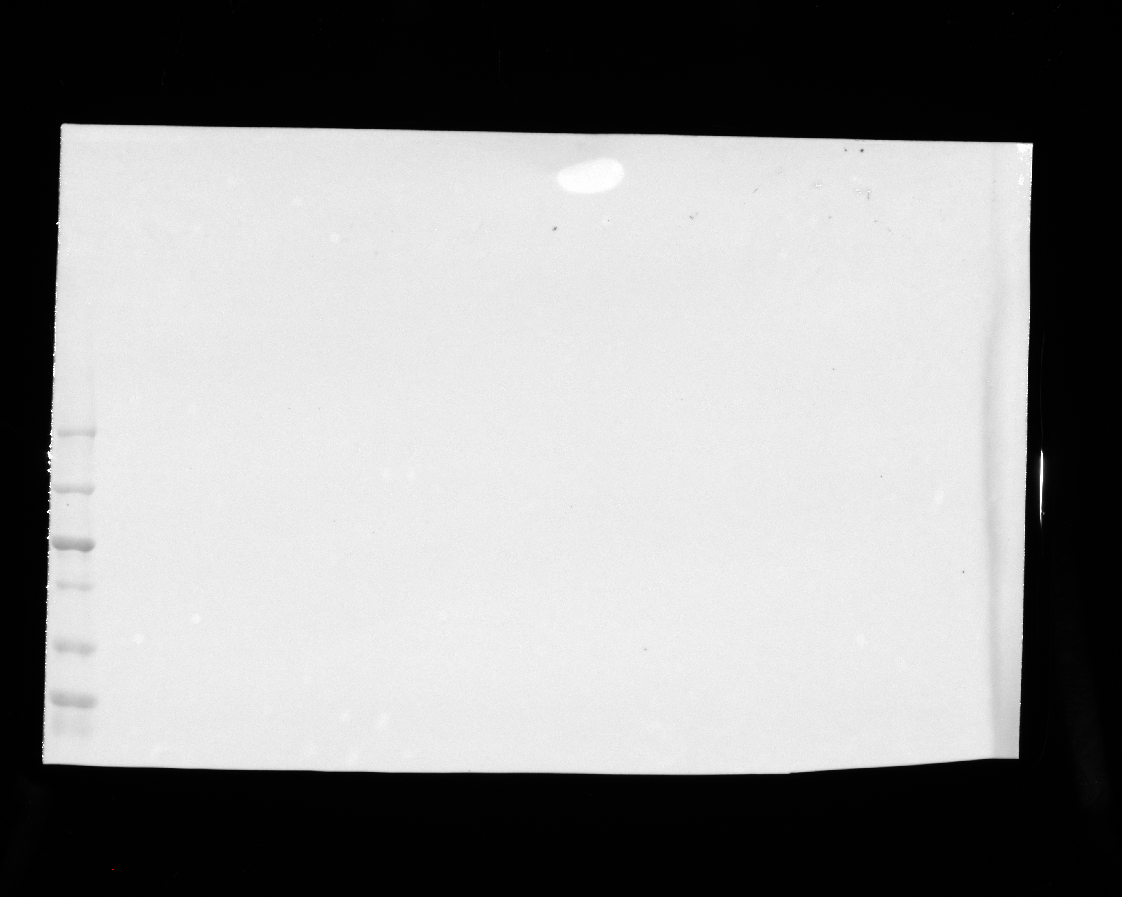

### 20240328_hek_phosphogel_2_gel1_stat5_12(Composite).tif

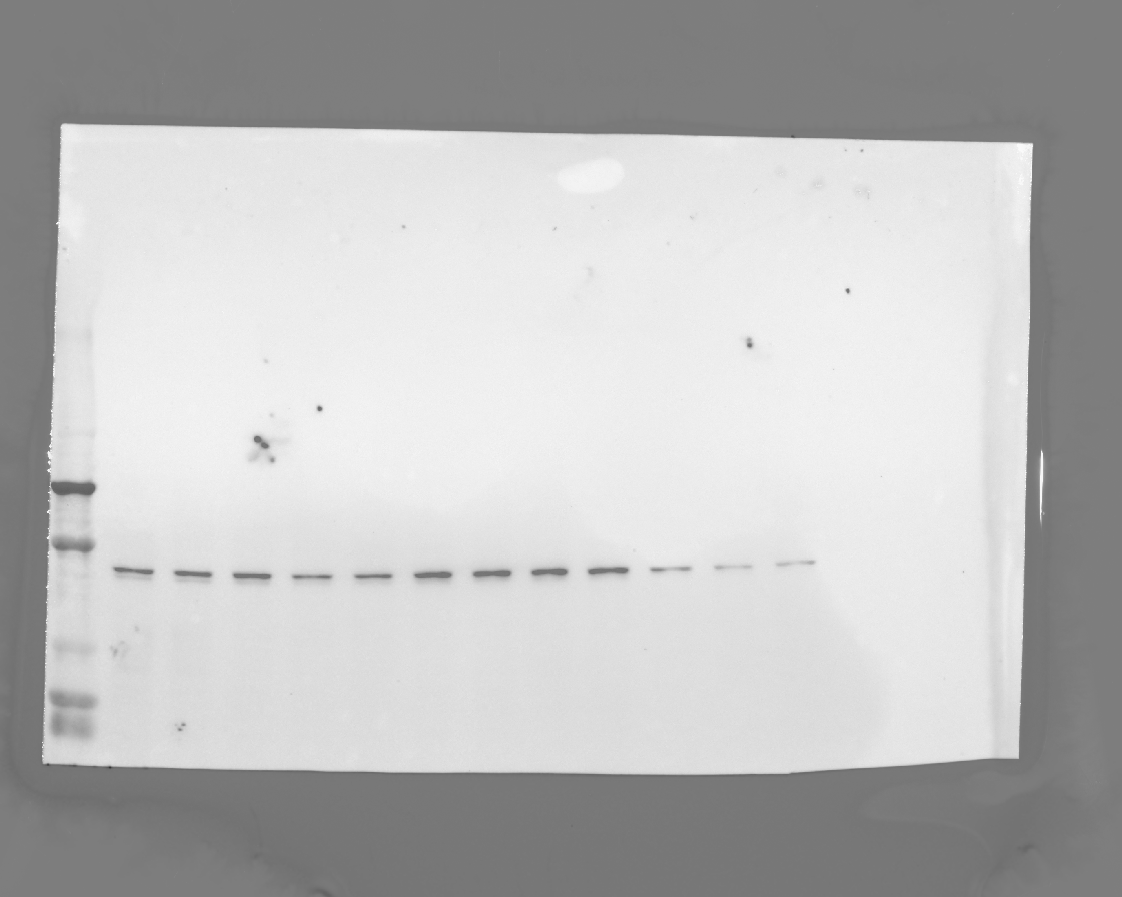

### 20240328_hek_phosphogel_2_gel1_stat5_phospho_7(Chemiluminescence).tif

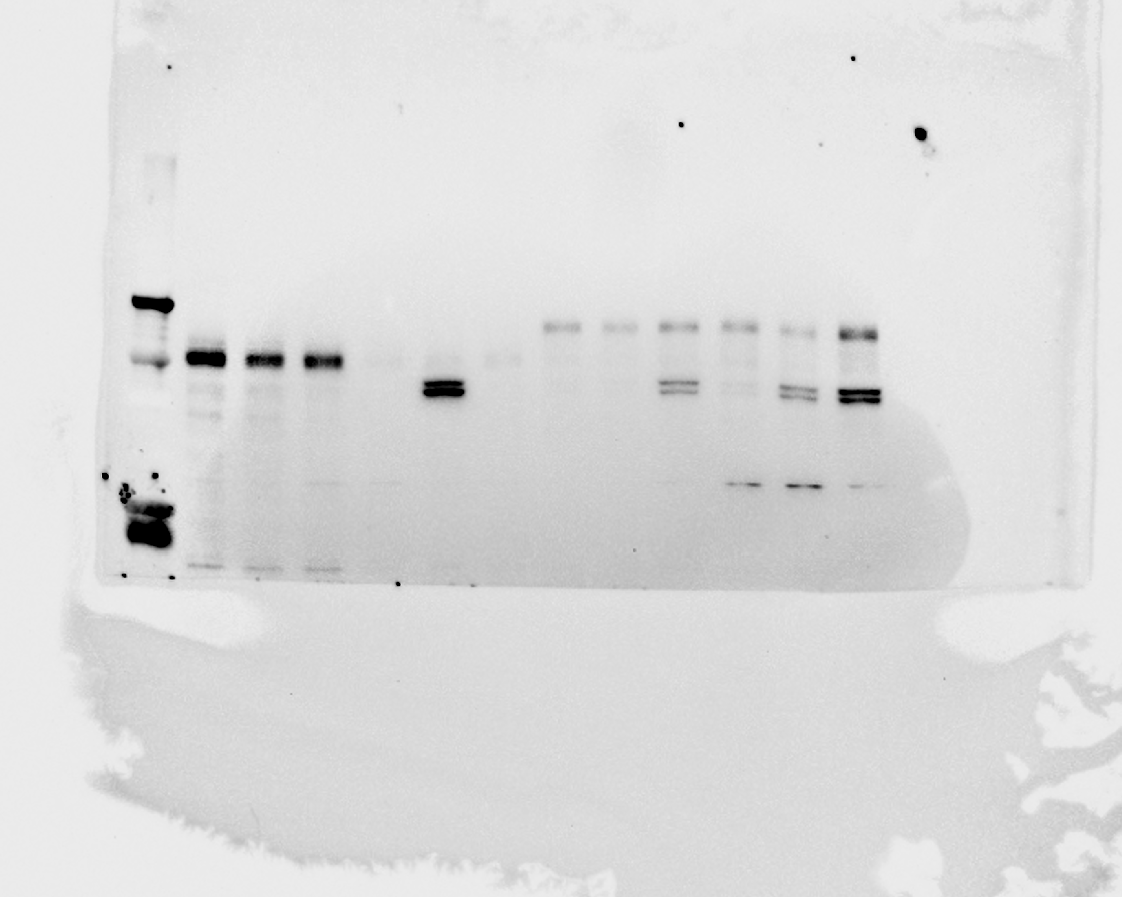

### 20240328_hek_phosphogel_2_gel1_stat5_phospho_7(Colorimetric).tif

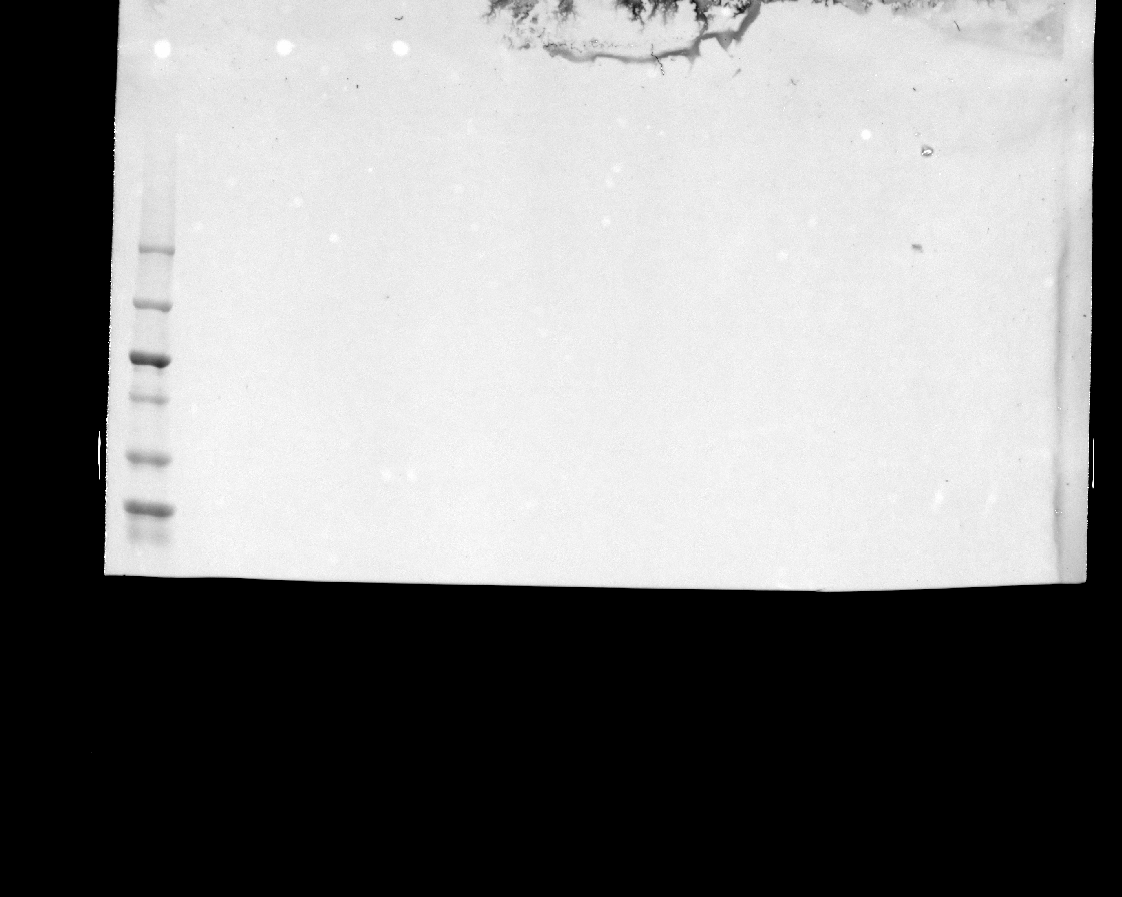

### 20240328_hek_phosphogel_2_gel1_stat5_phospho_7(Composite).tif

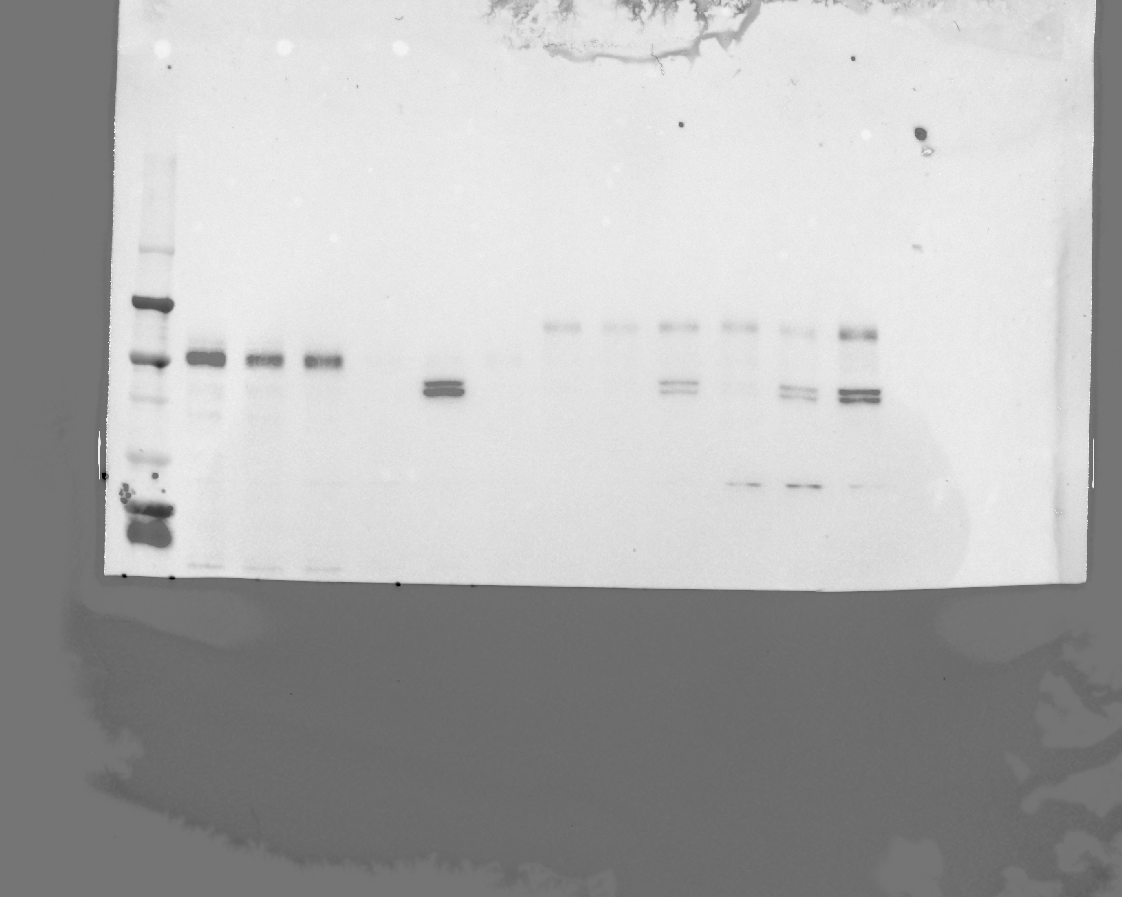

### 20240328_hek_phosphogel_2_gel2_flag_10(Chemiluminescence).tif

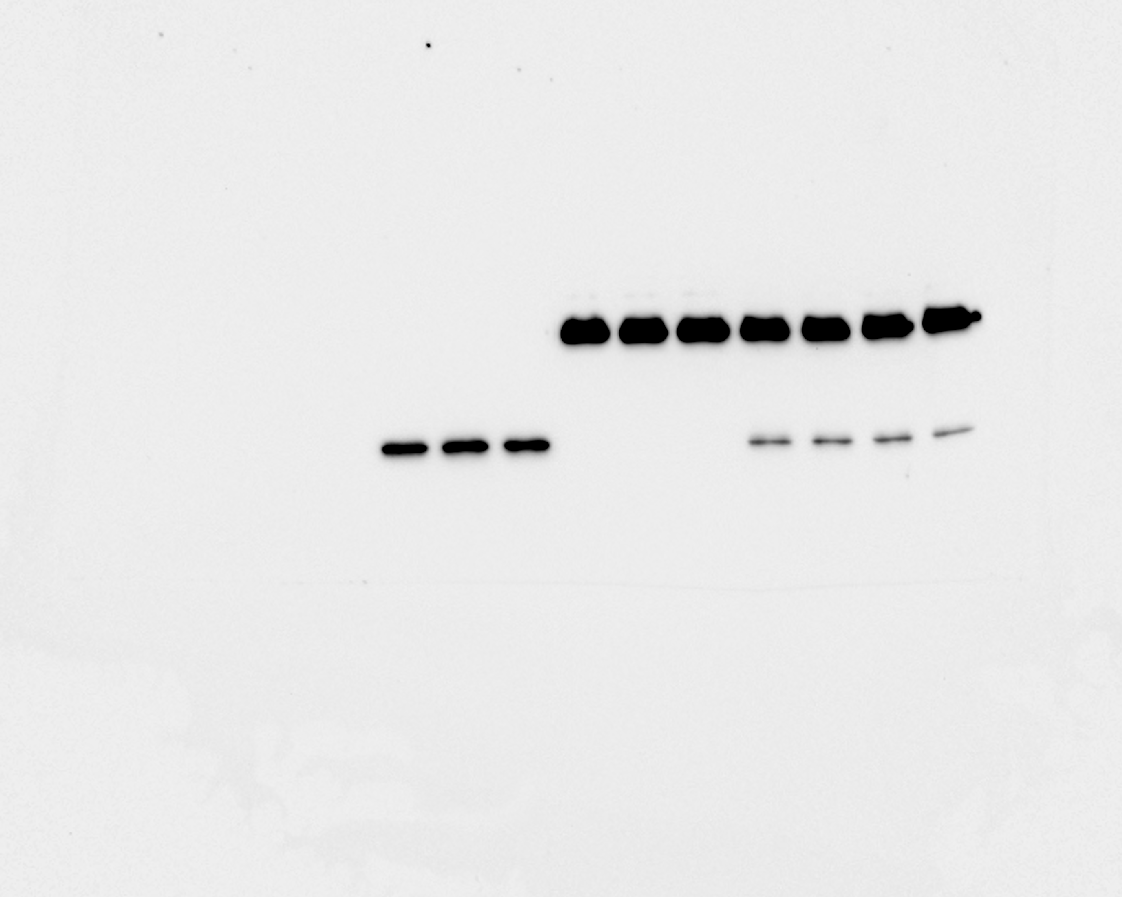

### 20240328_hek_phosphogel_2_gel2_flag_10(Colorimetric).tif

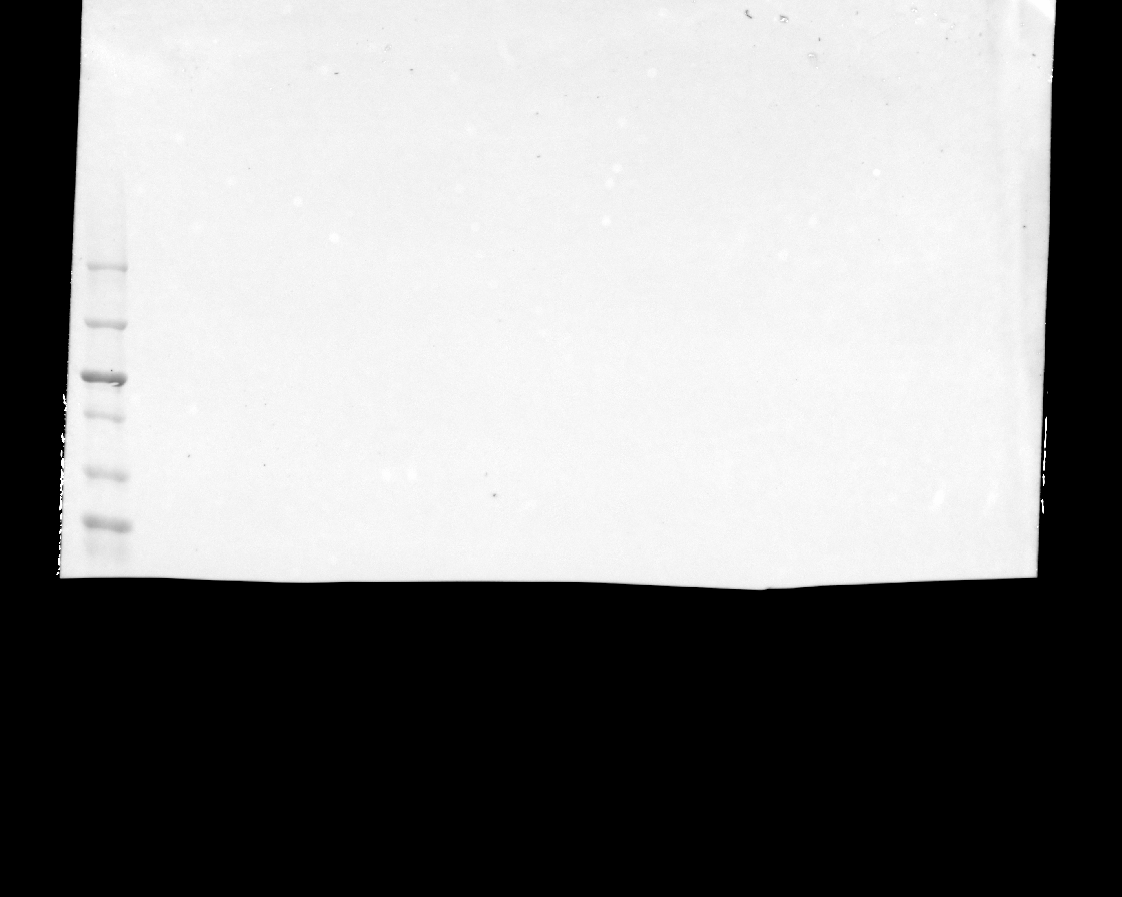

### 20240328_hek_phosphogel_2_gel2_flag_10(Composite).tif

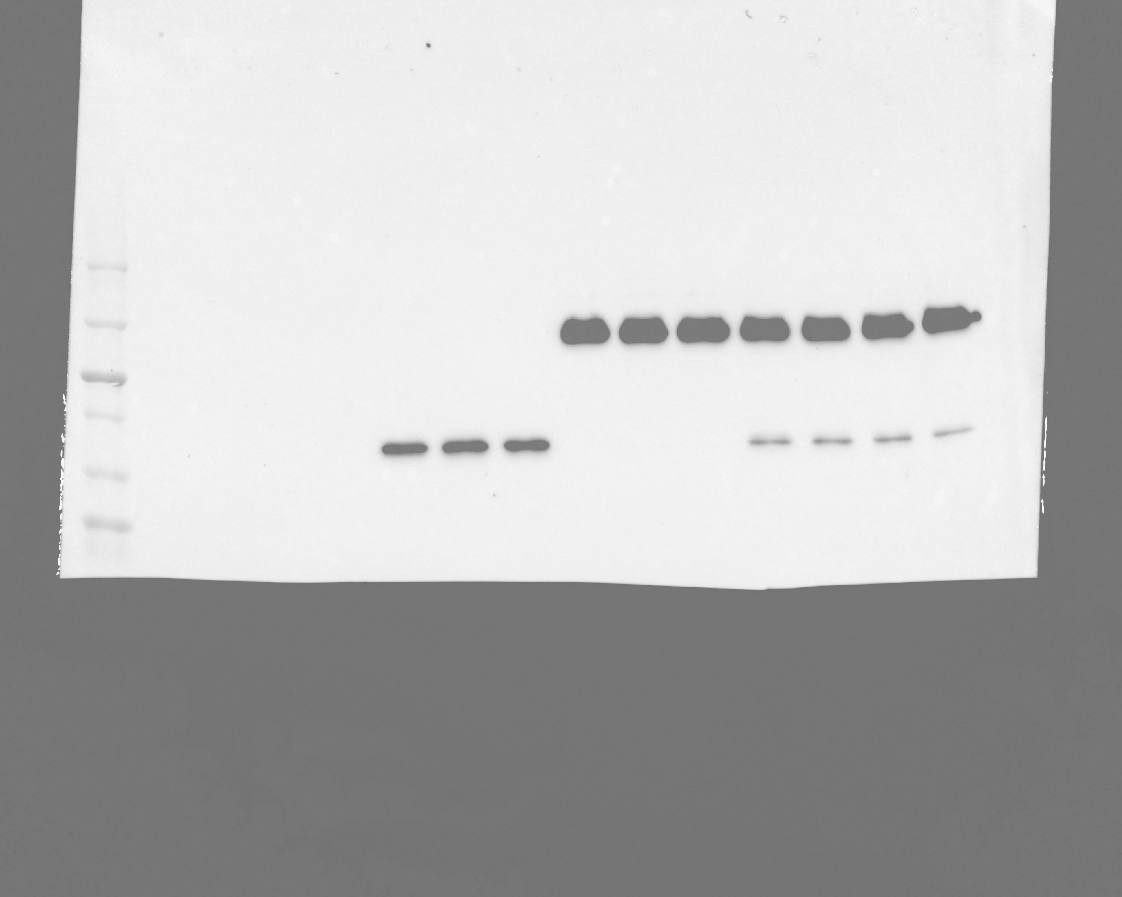

### 20240328_hek_phosphogel_2_gel2_hh3_stat5_5(Chemiluminescence).tif

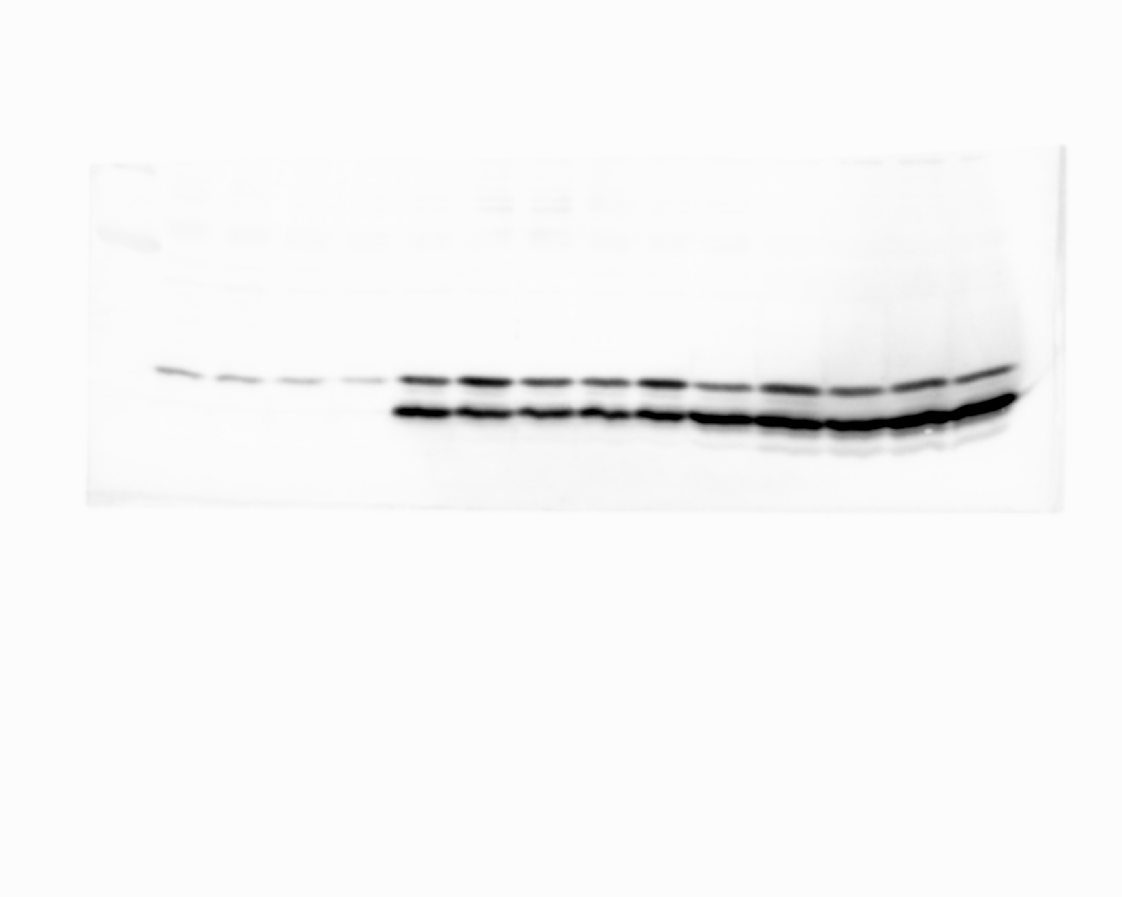

### 20240328_hek_phosphogel_2_gel2_hh3_stat5_5(Colorimetric).tif

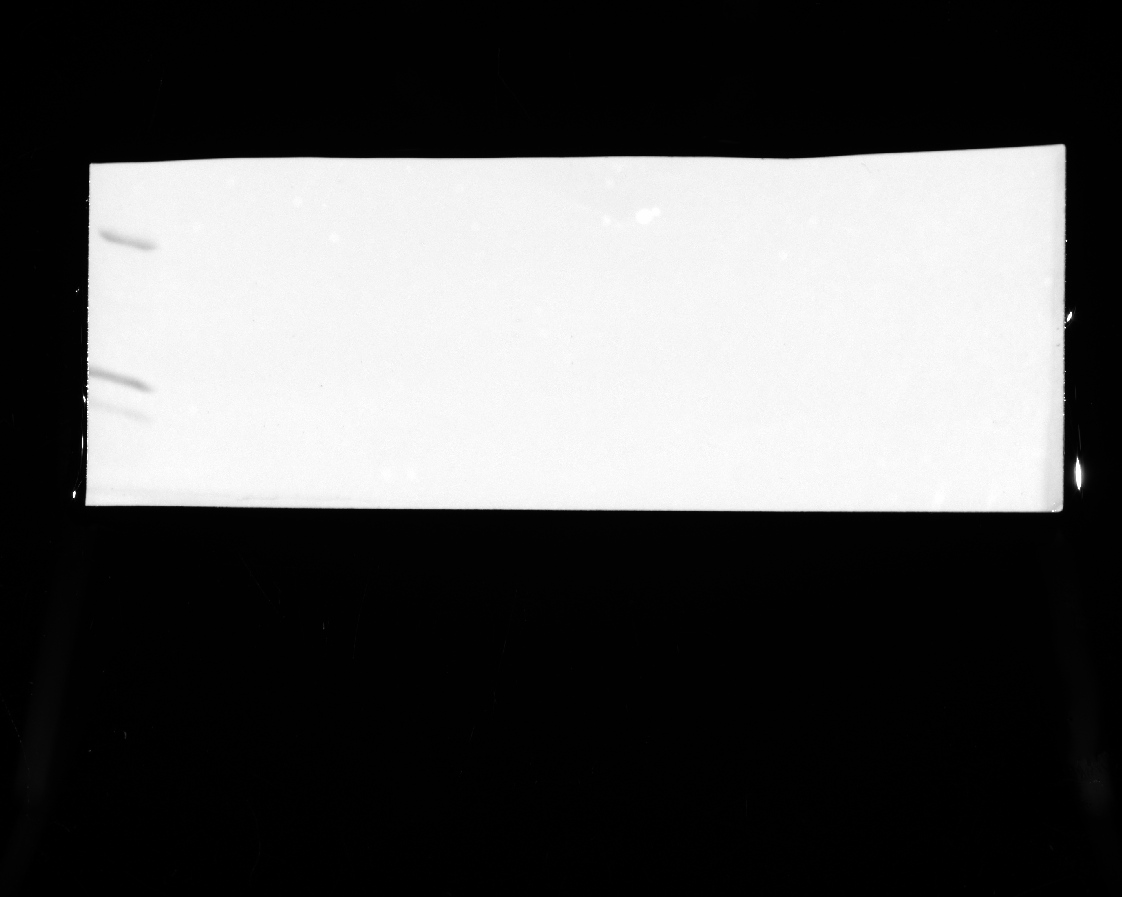

### 20240328_hek_phosphogel_2_gel2_hh3_stat5_5(Composite).tif

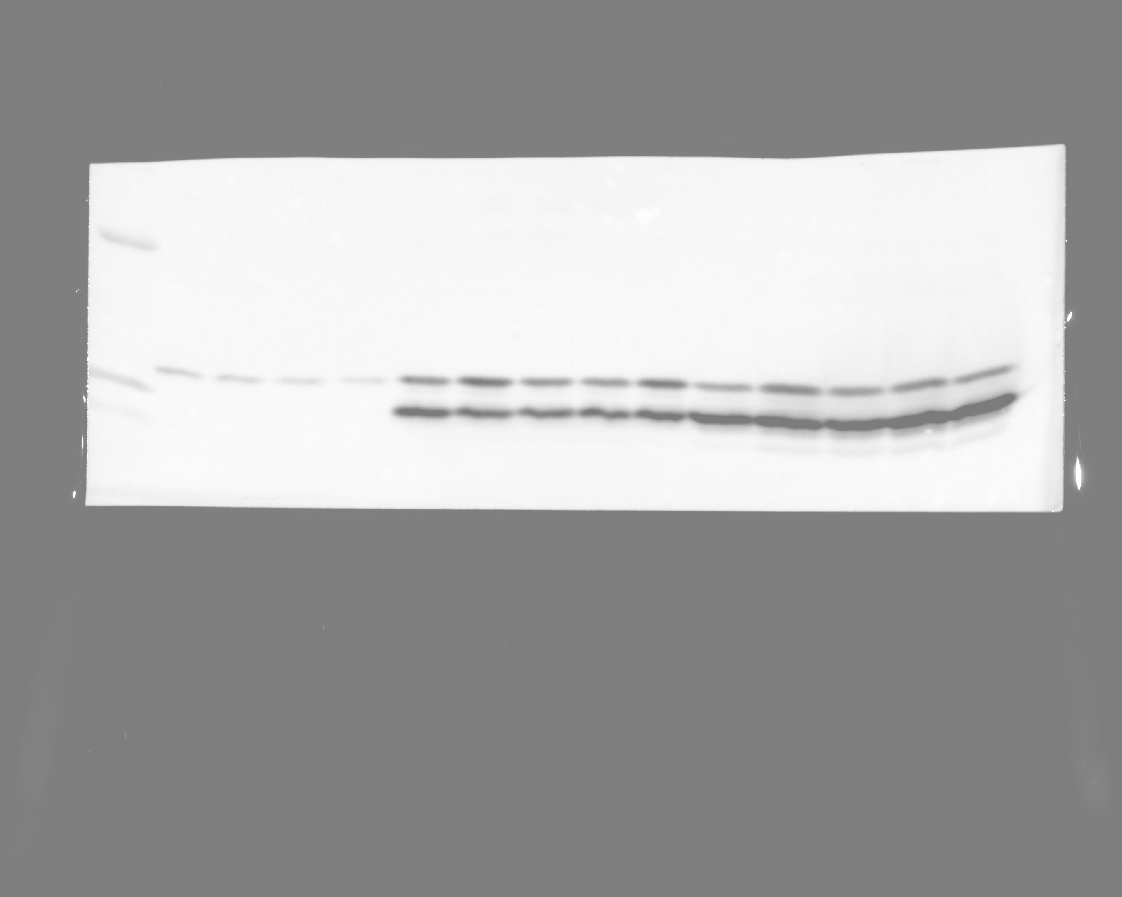

### 20240328_hek_phosphogel_2_gel2_hh3_stat5_phospho_4(Chemiluminescence).tif

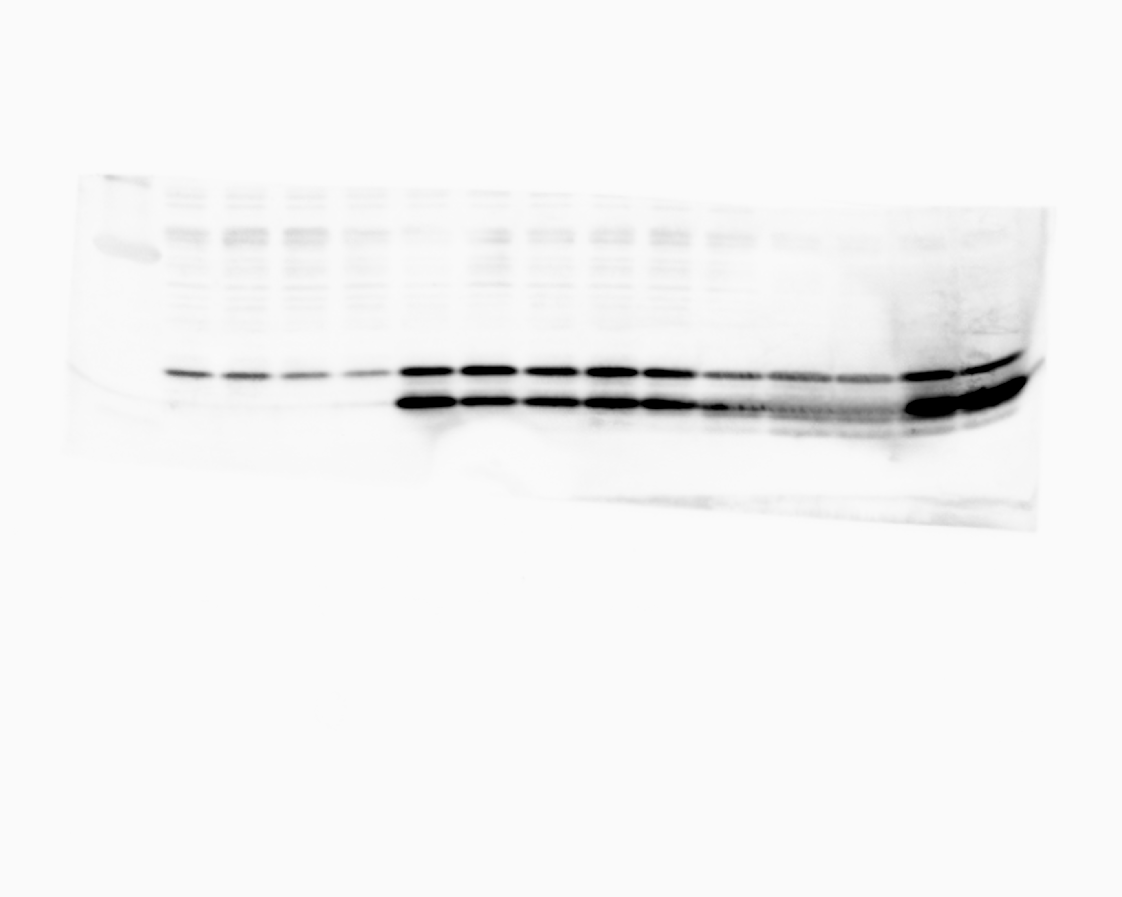

### 20240328_hek_phosphogel_2_gel2_hh3_stat5_phospho_4(Colorimetric).tif

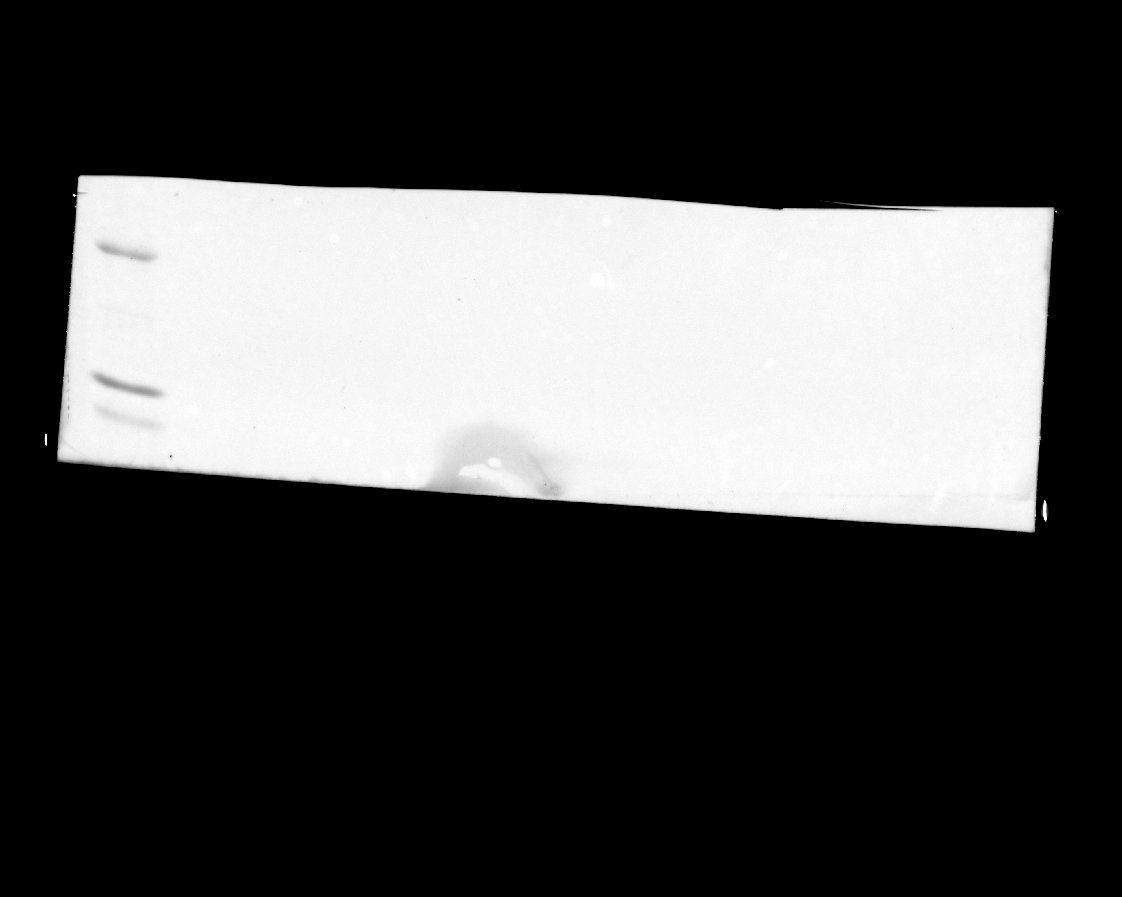

### 20240328_hek_phosphogel_2_gel2_hh3_stat5_phospho_4(Composite).tif

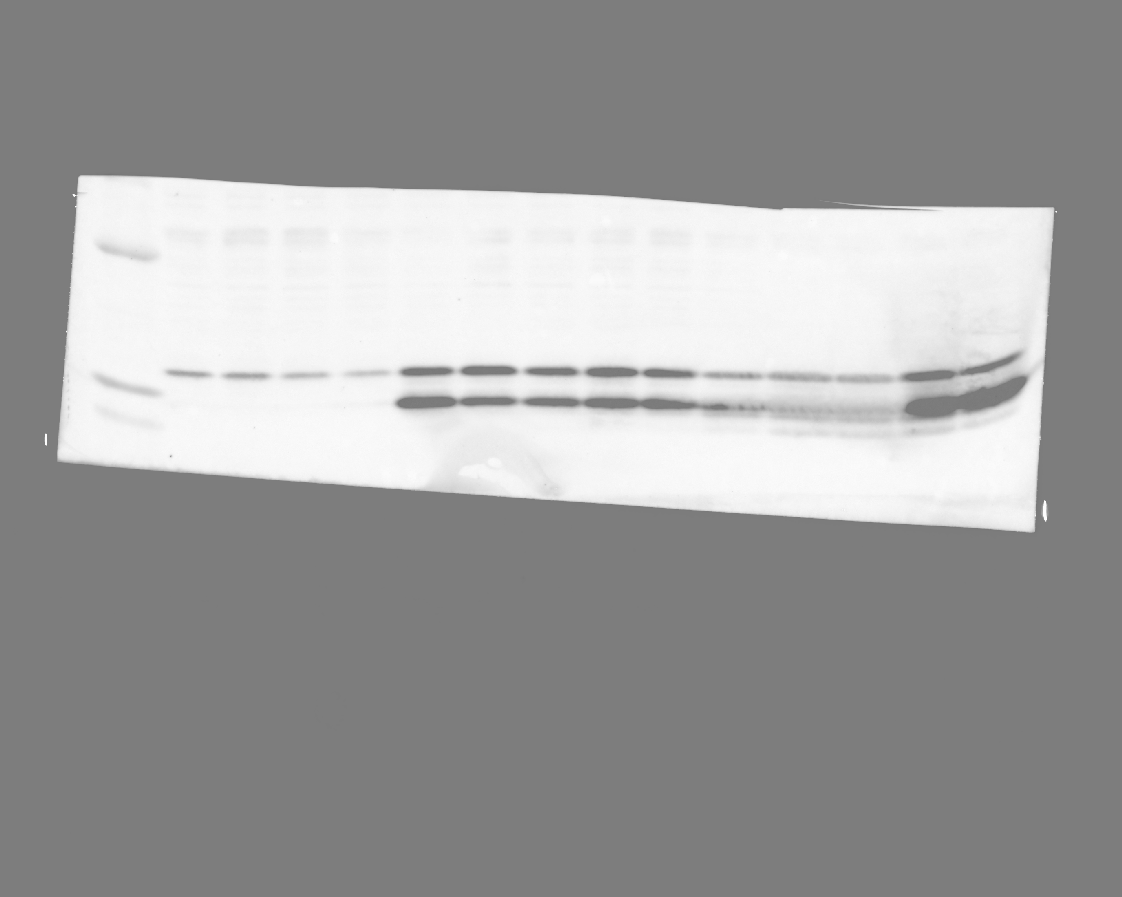

### 20240328_hek_phosphogel_2_gel2_stat5_10(Chemiluminescence).tif

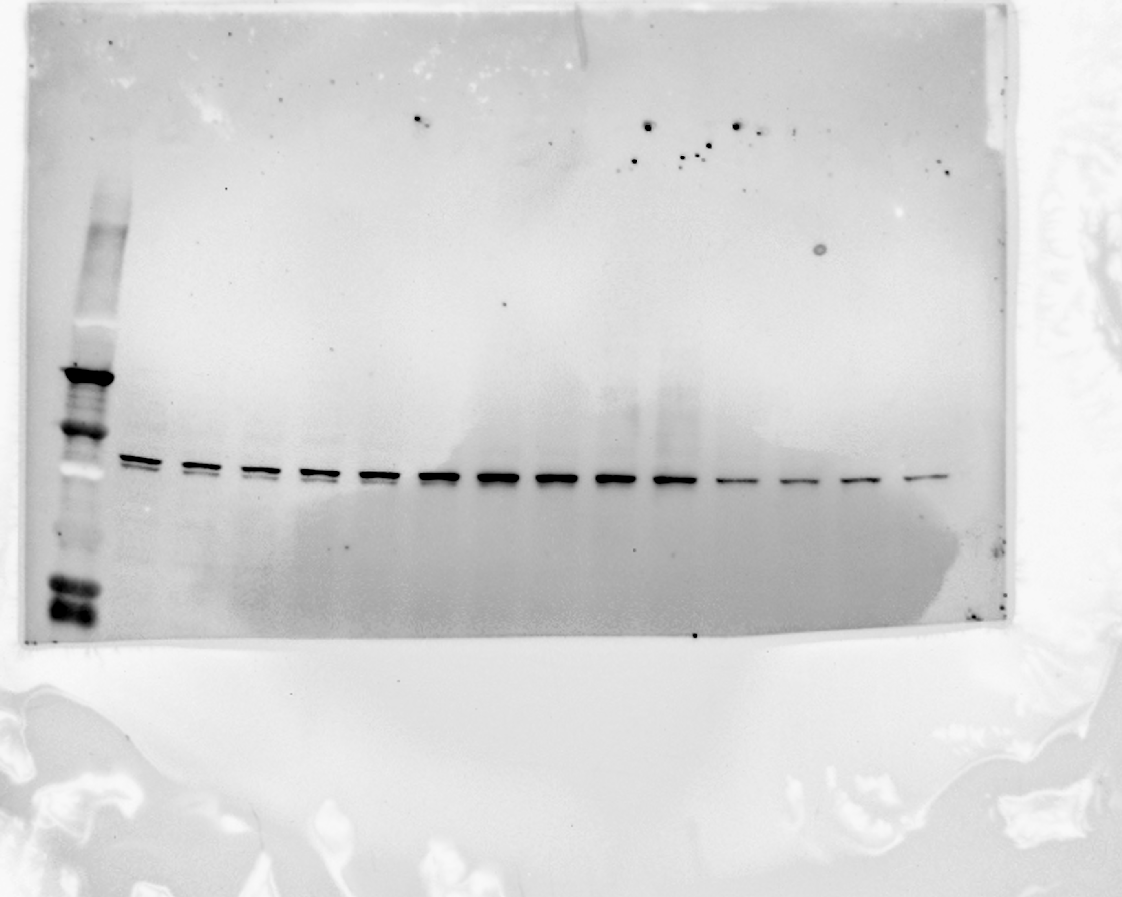

### 20240328_hek_phosphogel_2_gel2_stat5_10(Colorimetric).tif

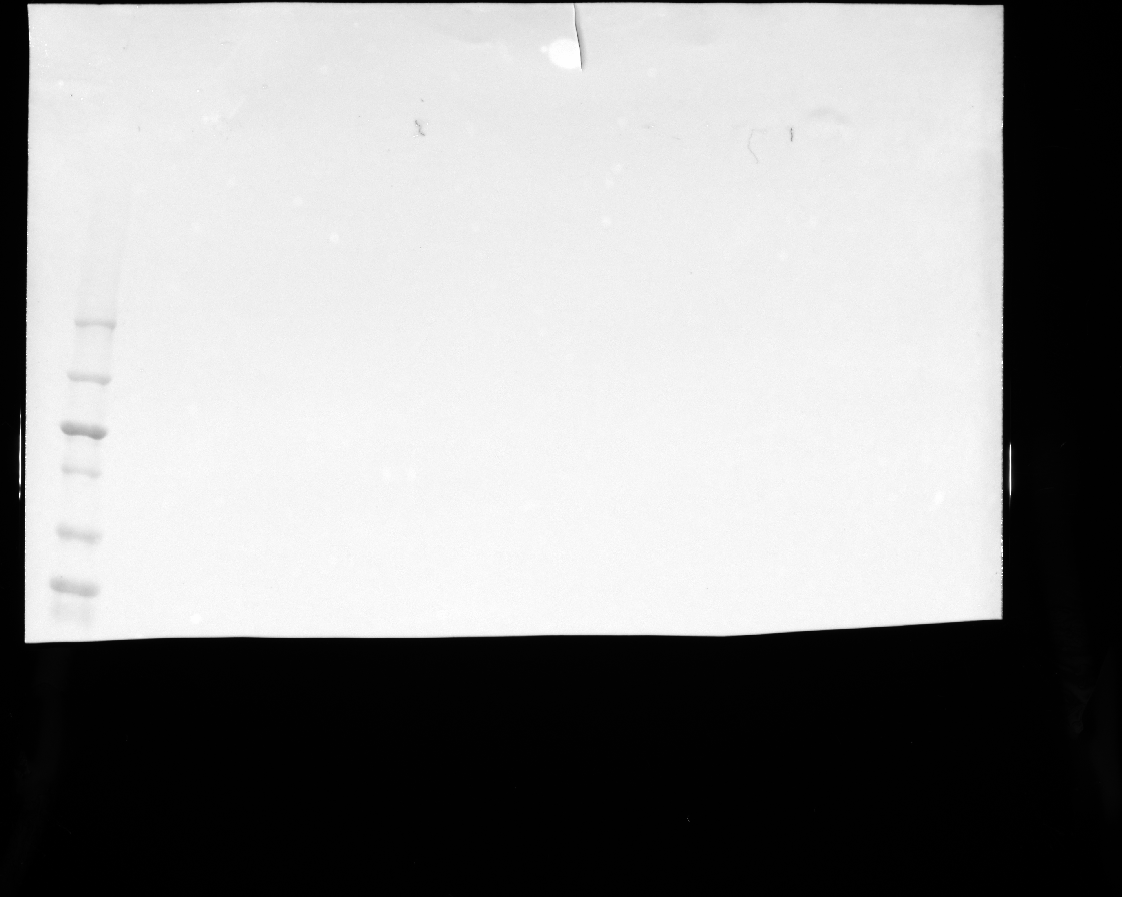

### 20240328_hek_phosphogel_2_gel2_stat5_10(Composite).tif

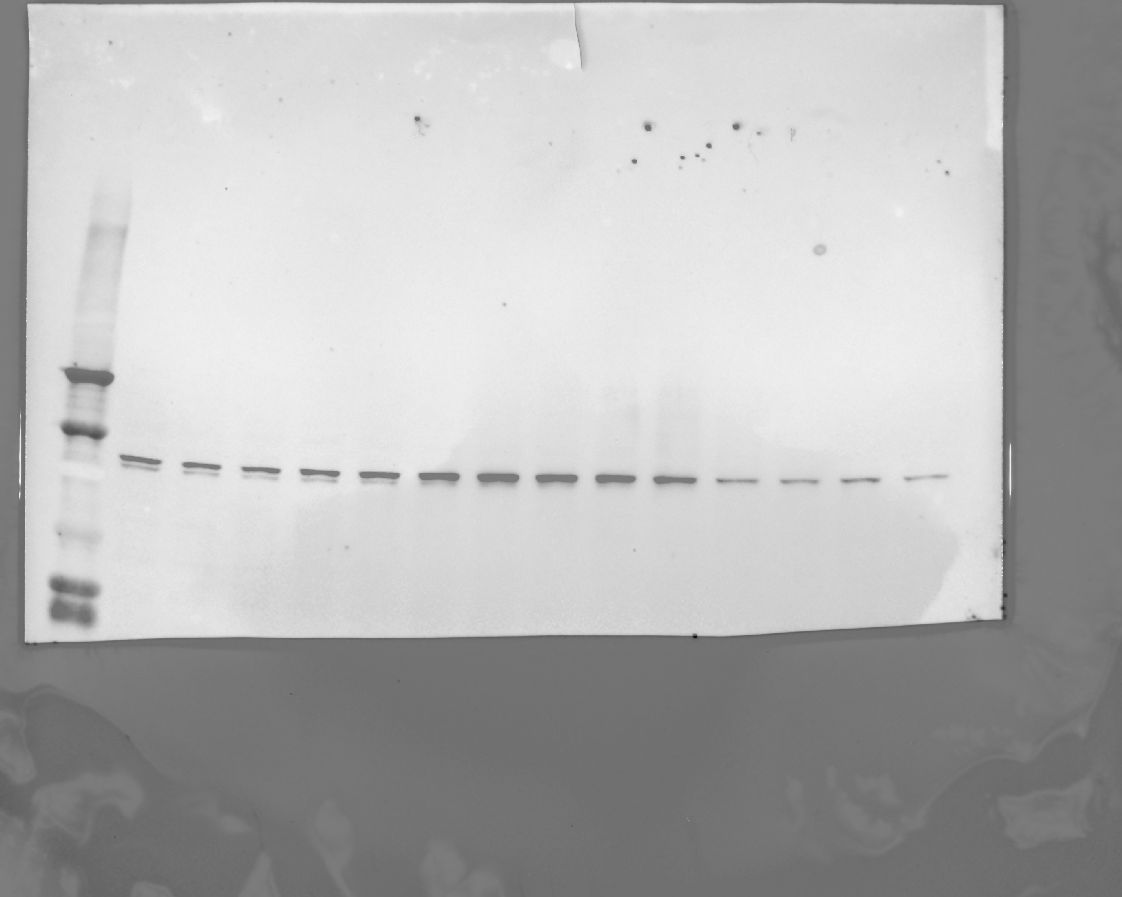

### 20240328_hek_phosphogel_2_gel2_stat5_10(Composite)_2.tif

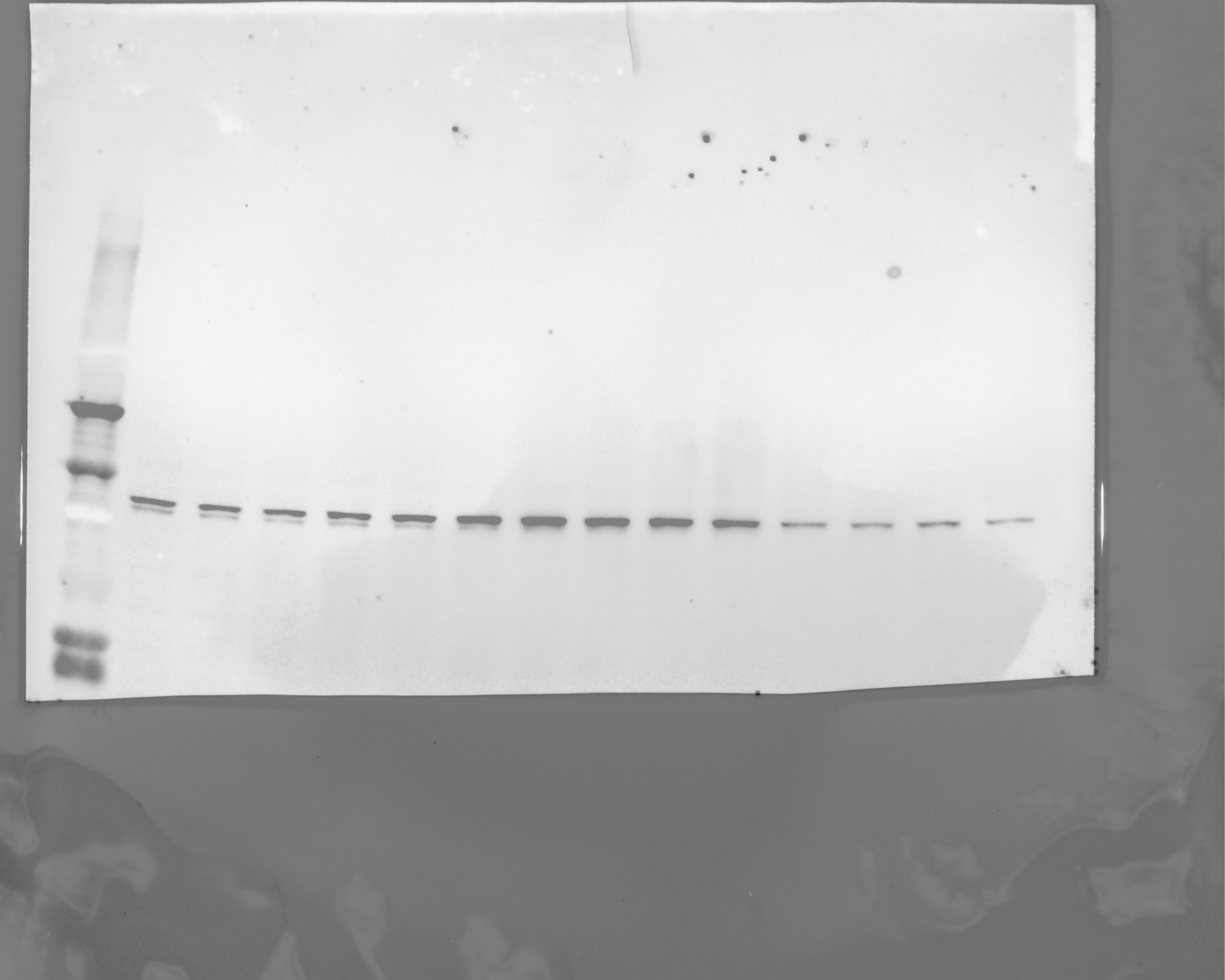

### 20240328_hek_phosphogel_2_gel2_stat5_phospho_6(Chemiluminescence).tif

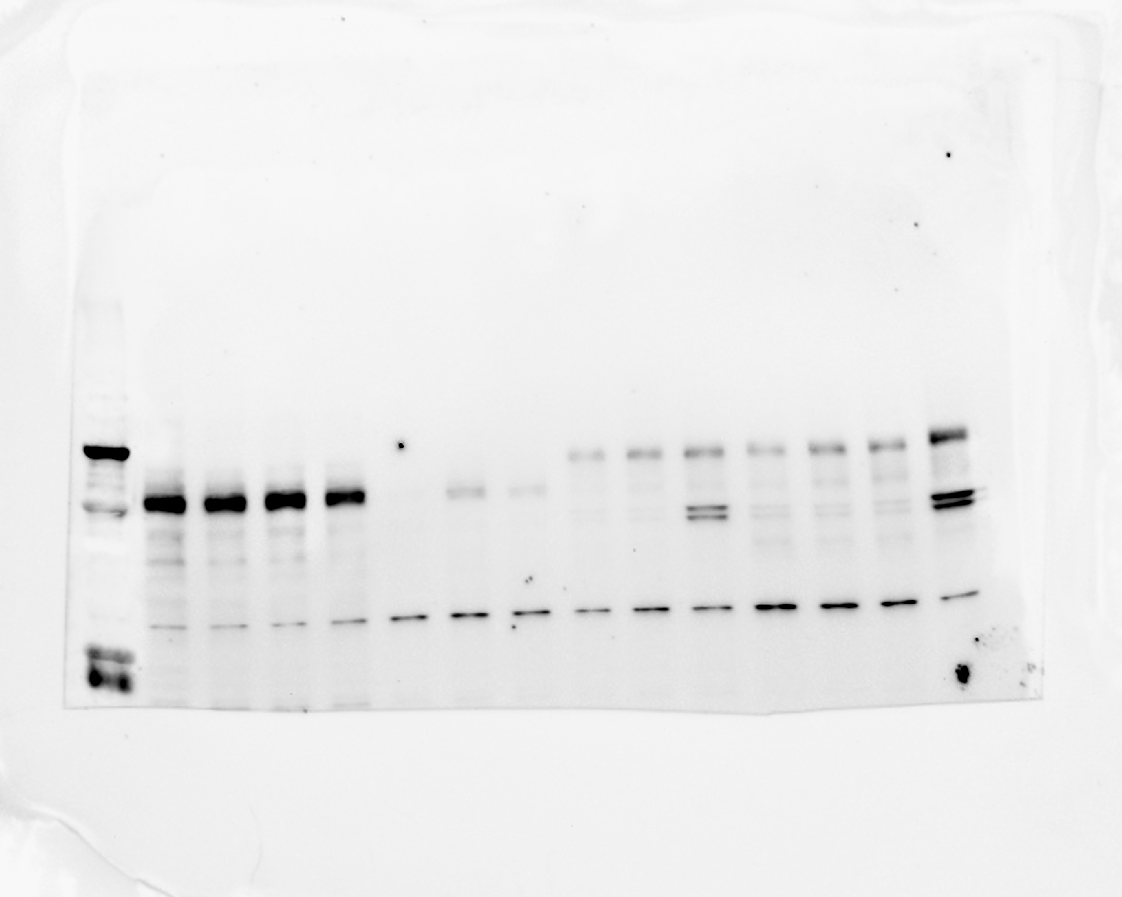

### 20240328_hek_phosphogel_2_gel2_stat5_phospho_6(Colorimetric).tif

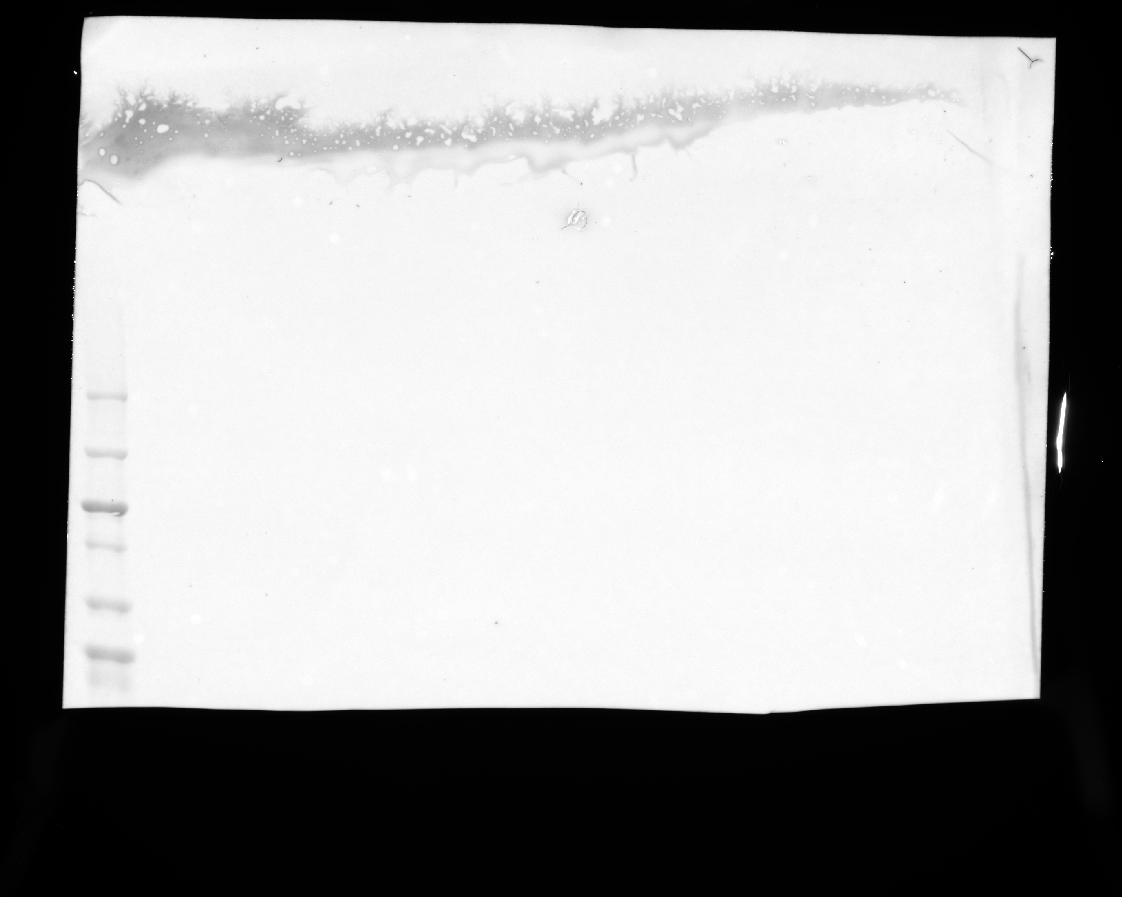

### 20240328_hek_phosphogel_2_gel2_stat5_phospho_6(Composite).tif

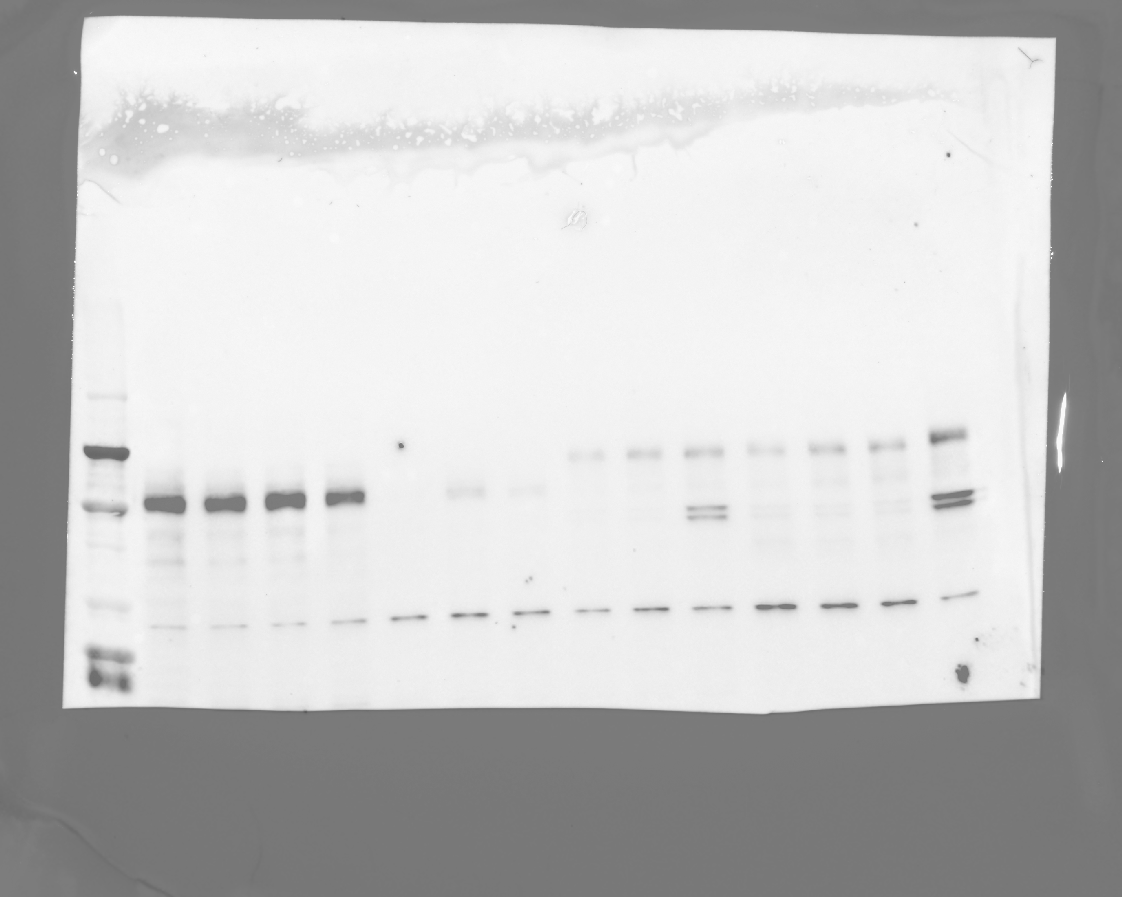

### 20240618_p26_3s_p29_purif_3_coomassie(Coomassie Blue).tif

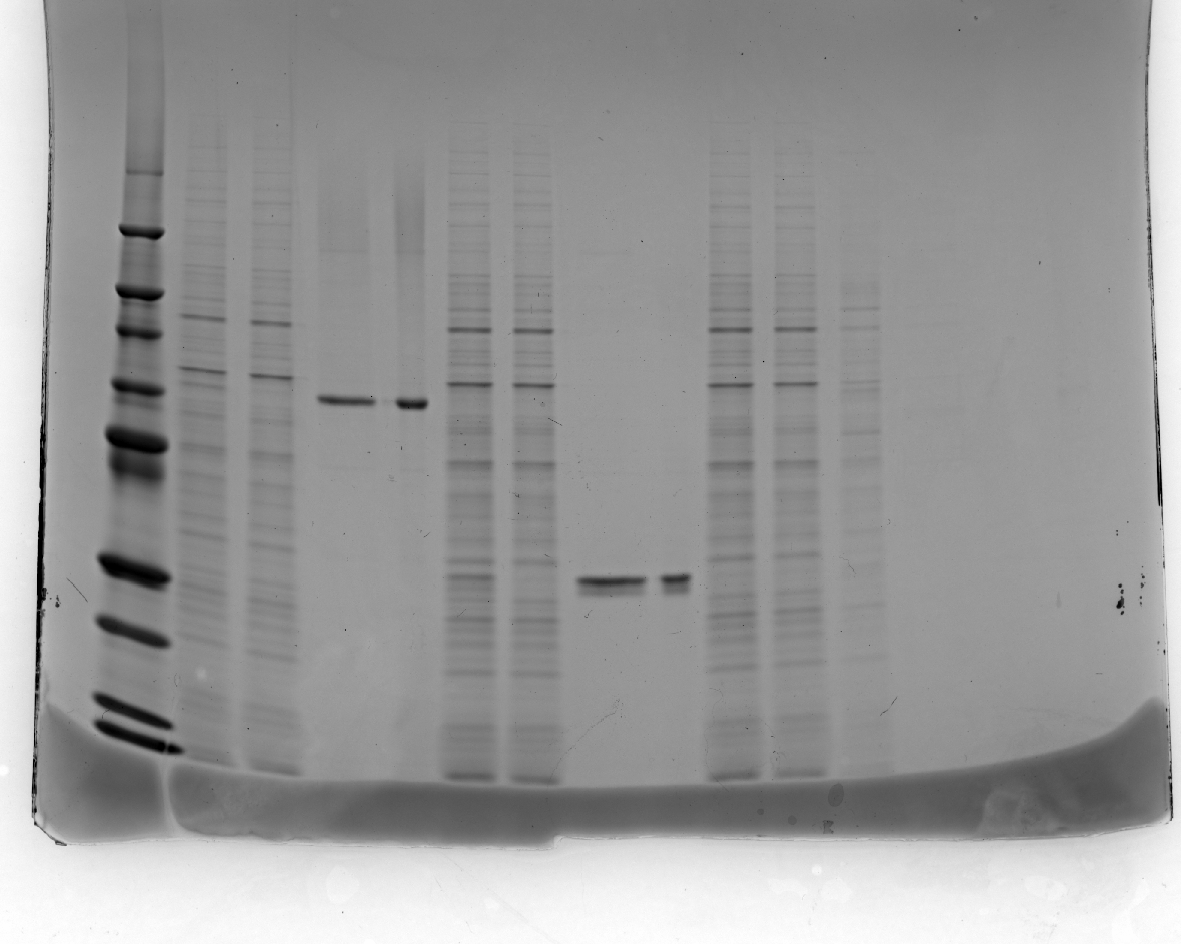

### 20240829_hek_phosphogel_5_linkervars_flaghrp_10(Chemiluminescence).tif

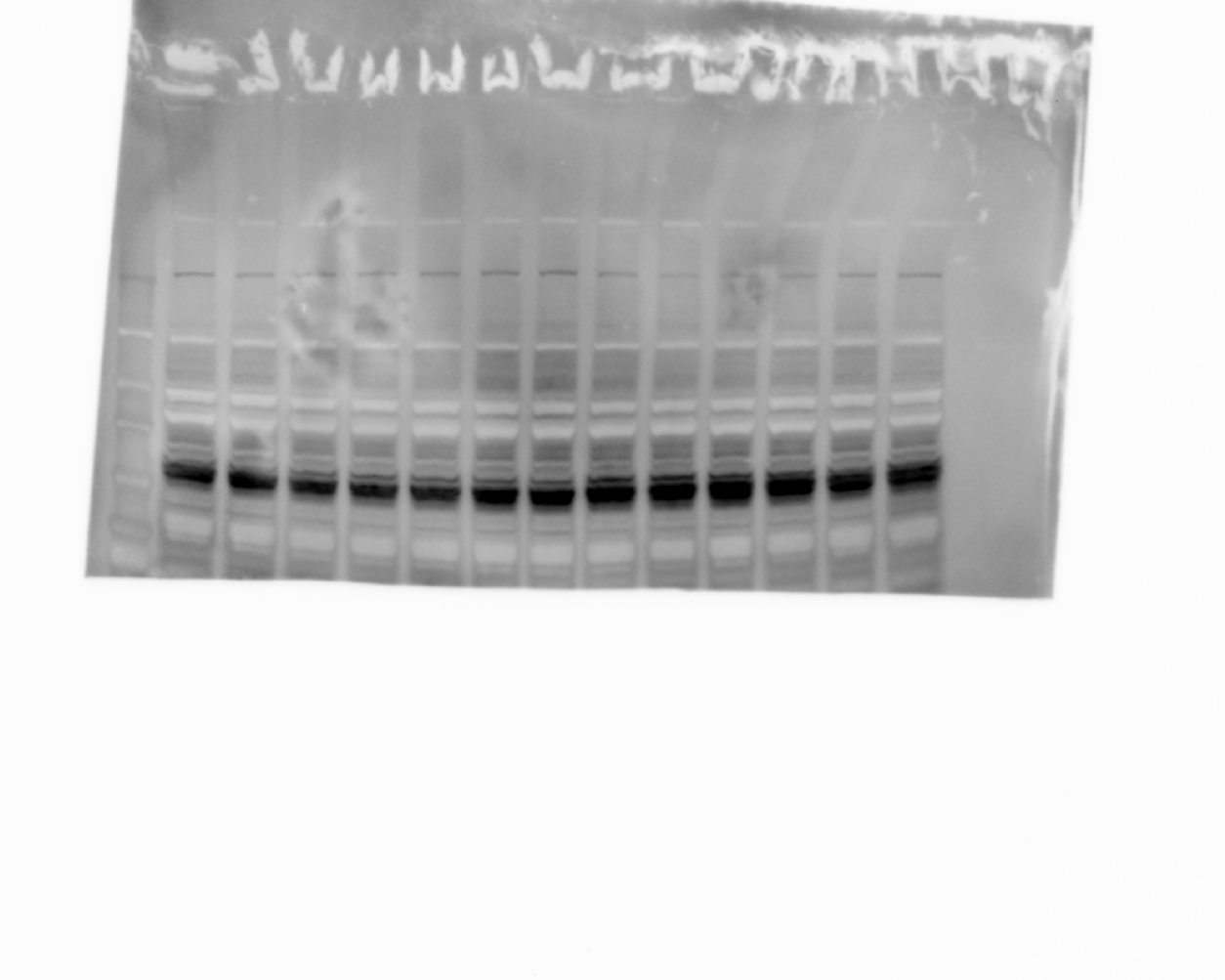

### 20240829_hek_phosphogel_5_linkervars_flaghrp_10(Colorimetric).tif

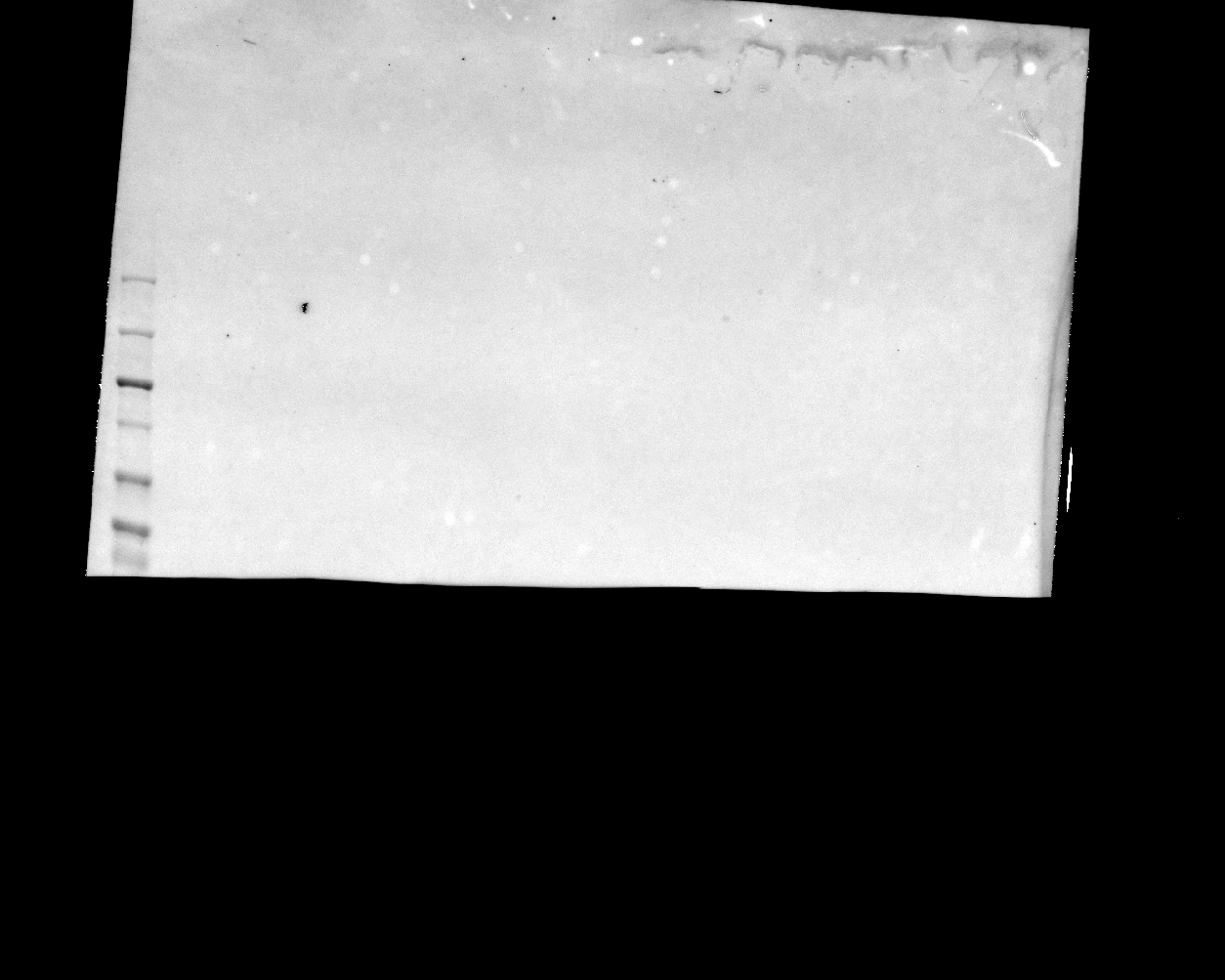

### 20240829_hek_phosphogel_5_linkervars_flaghrp_10(Composite).tif

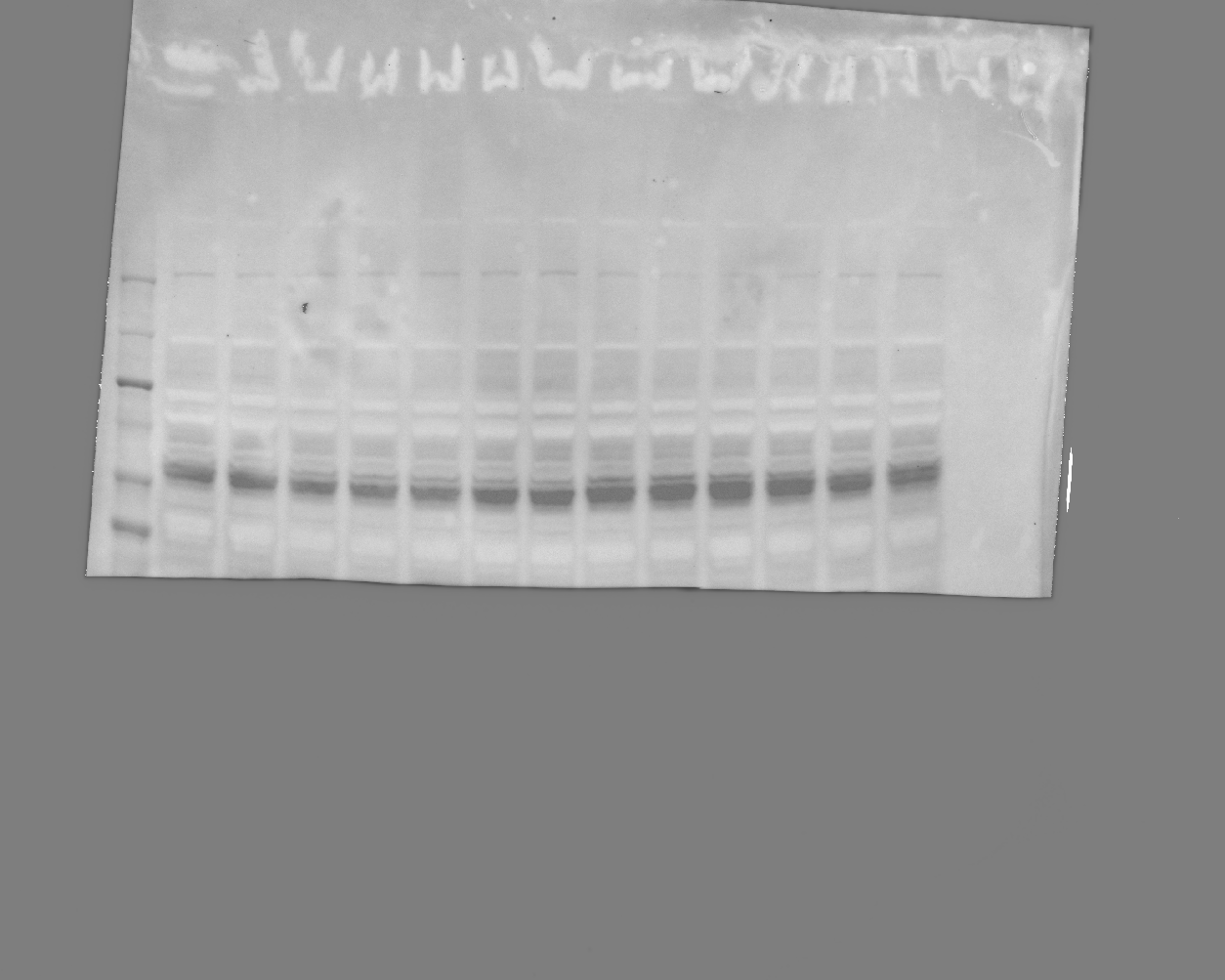

### 20240829_hek_phosphogel_5_linkervars_gel1_hh3_10(Chemiluminescence).tif

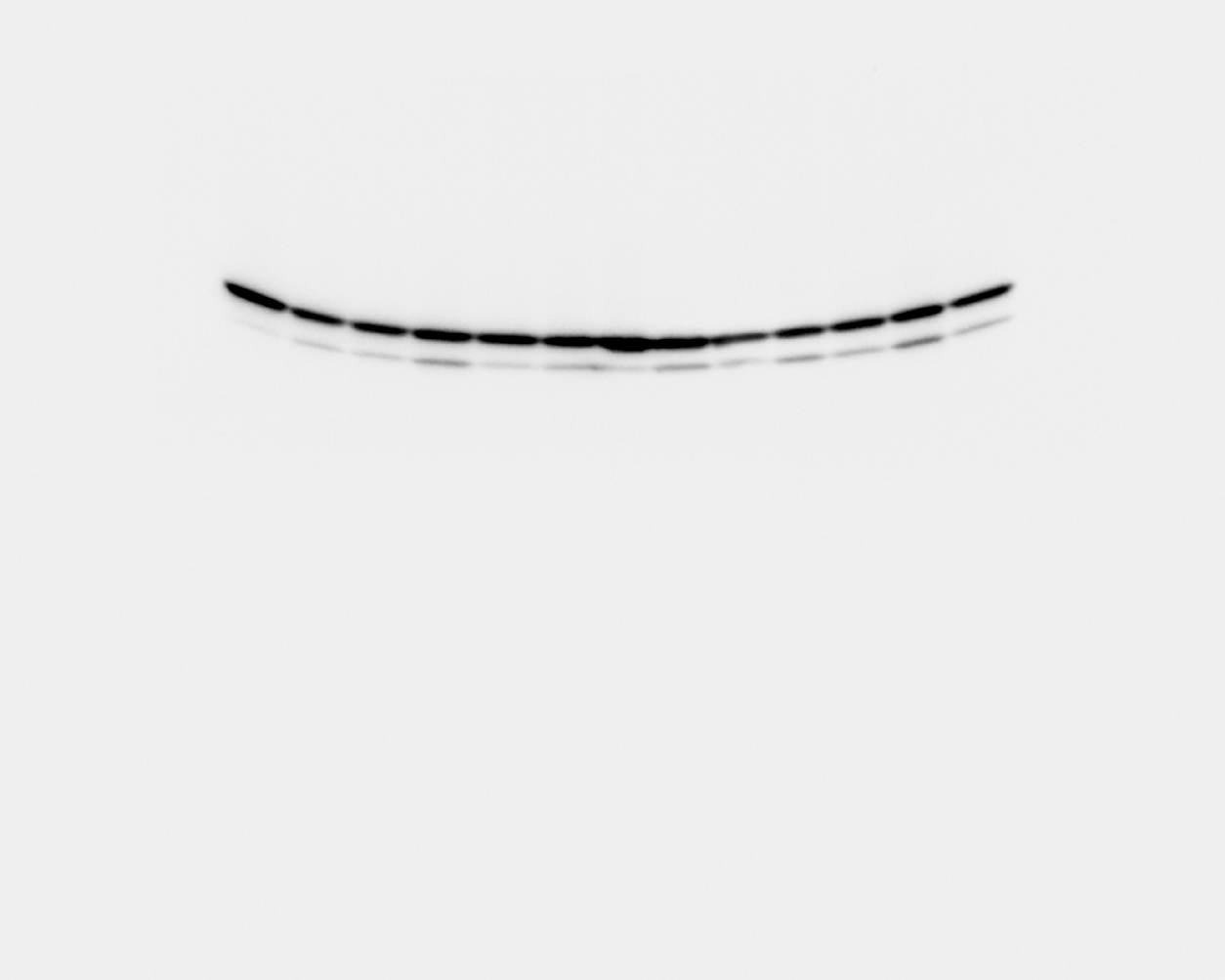

### 20240829_hek_phosphogel_5_linkervars_gel1_hh3_10(Colorimetric).tif

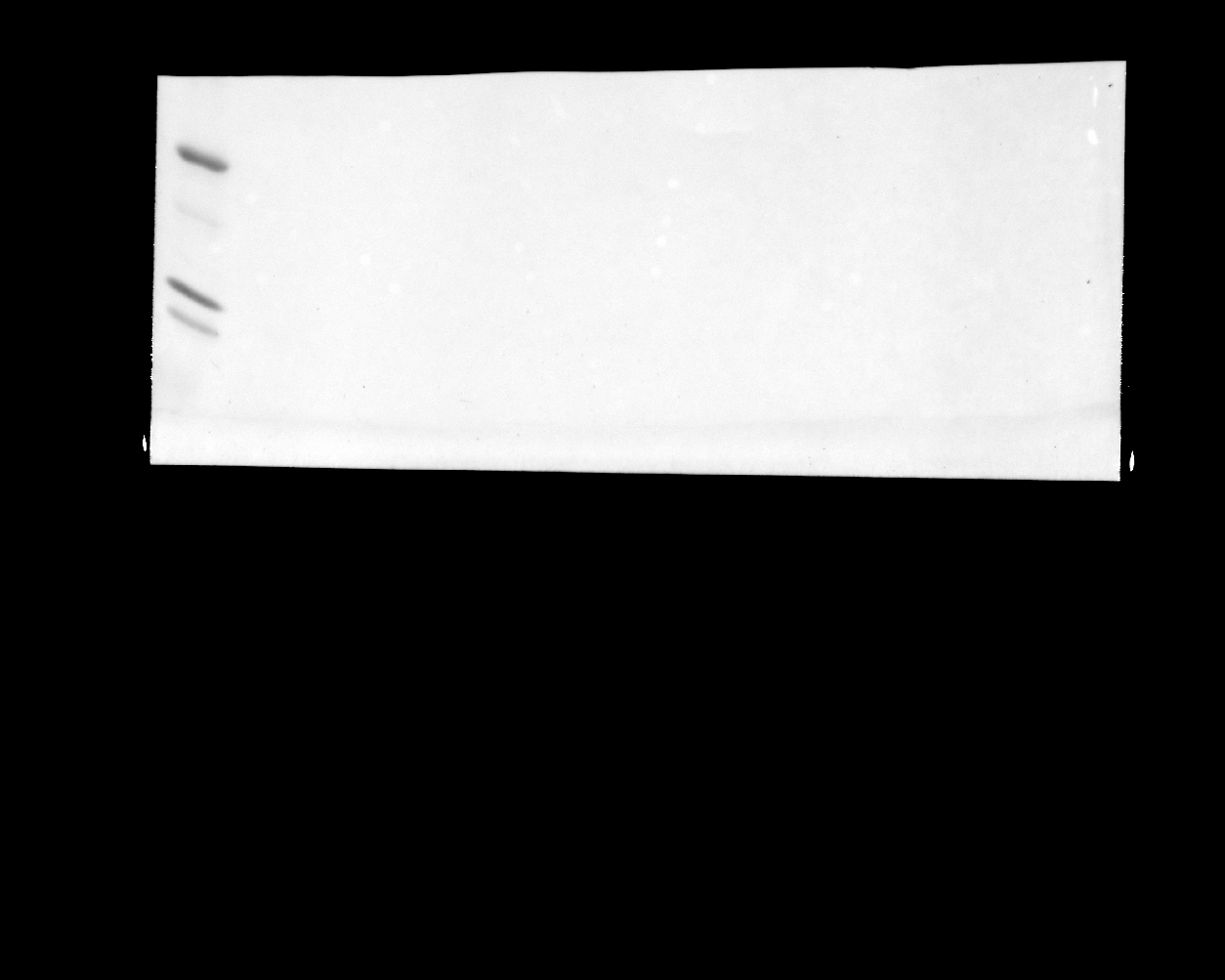

### 20240829_hek_phosphogel_5_linkervars_gel1_hh3_10(Composite).tif

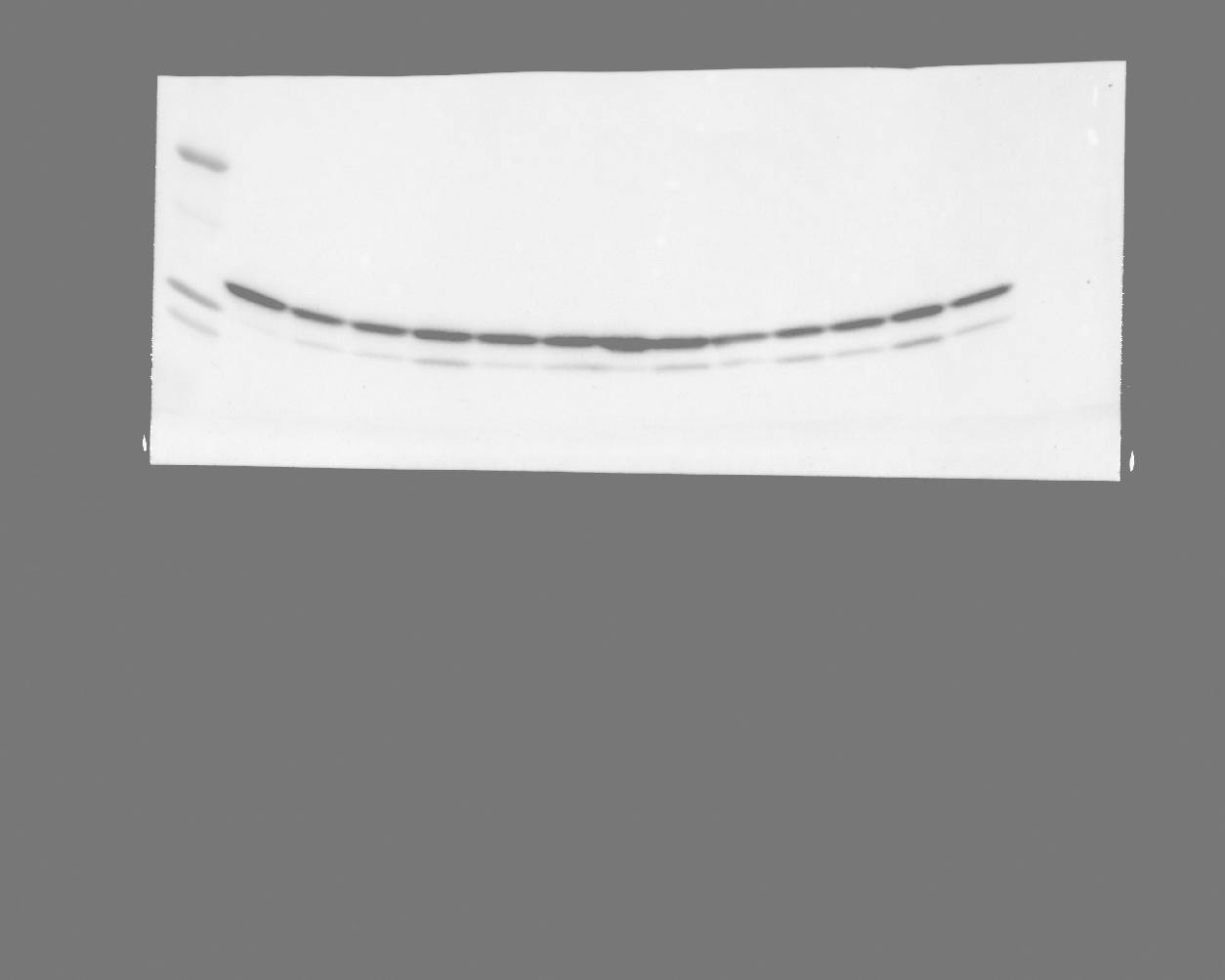

### 20240829_hek_phosphogel_5_linkervars_gel1_stat5_phospho_05(Chemiluminescence).tif

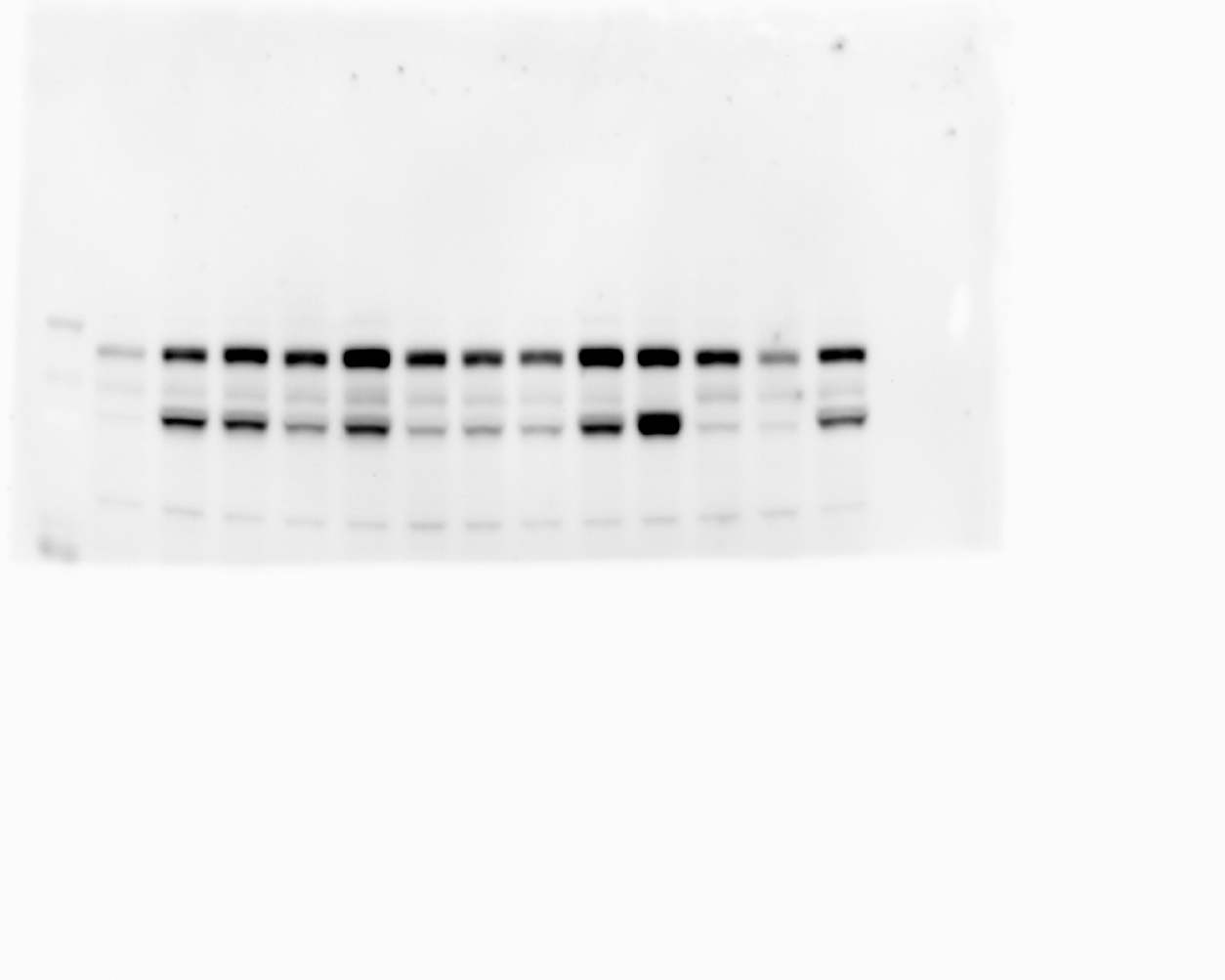

### 20240829_hek_phosphogel_5_linkervars_gel1_stat5_phospho_05(Colorimetric).tif

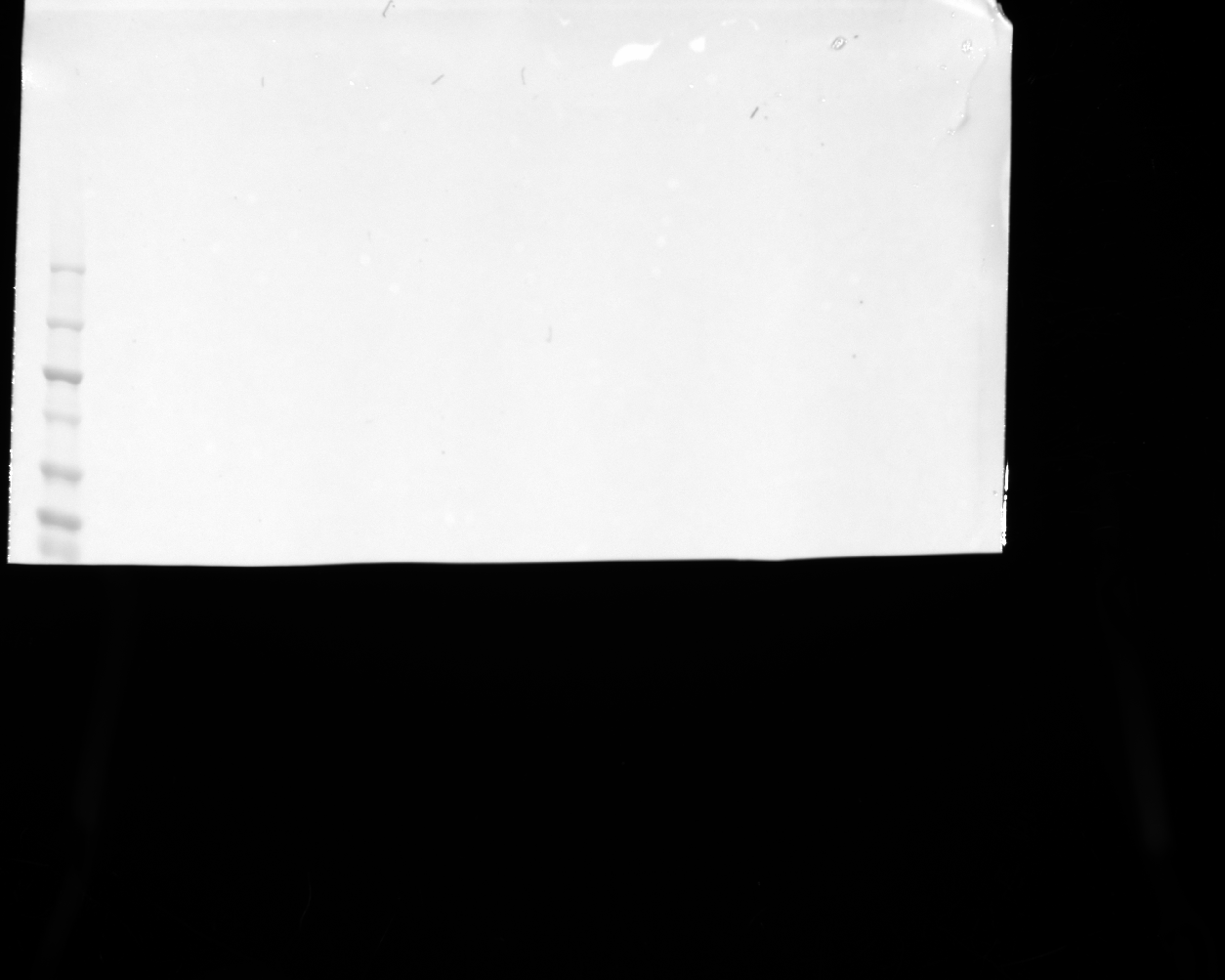

### 20240829_hek_phosphogel_5_linkervars_gel1_stat5_phospho_05(Composite).tif

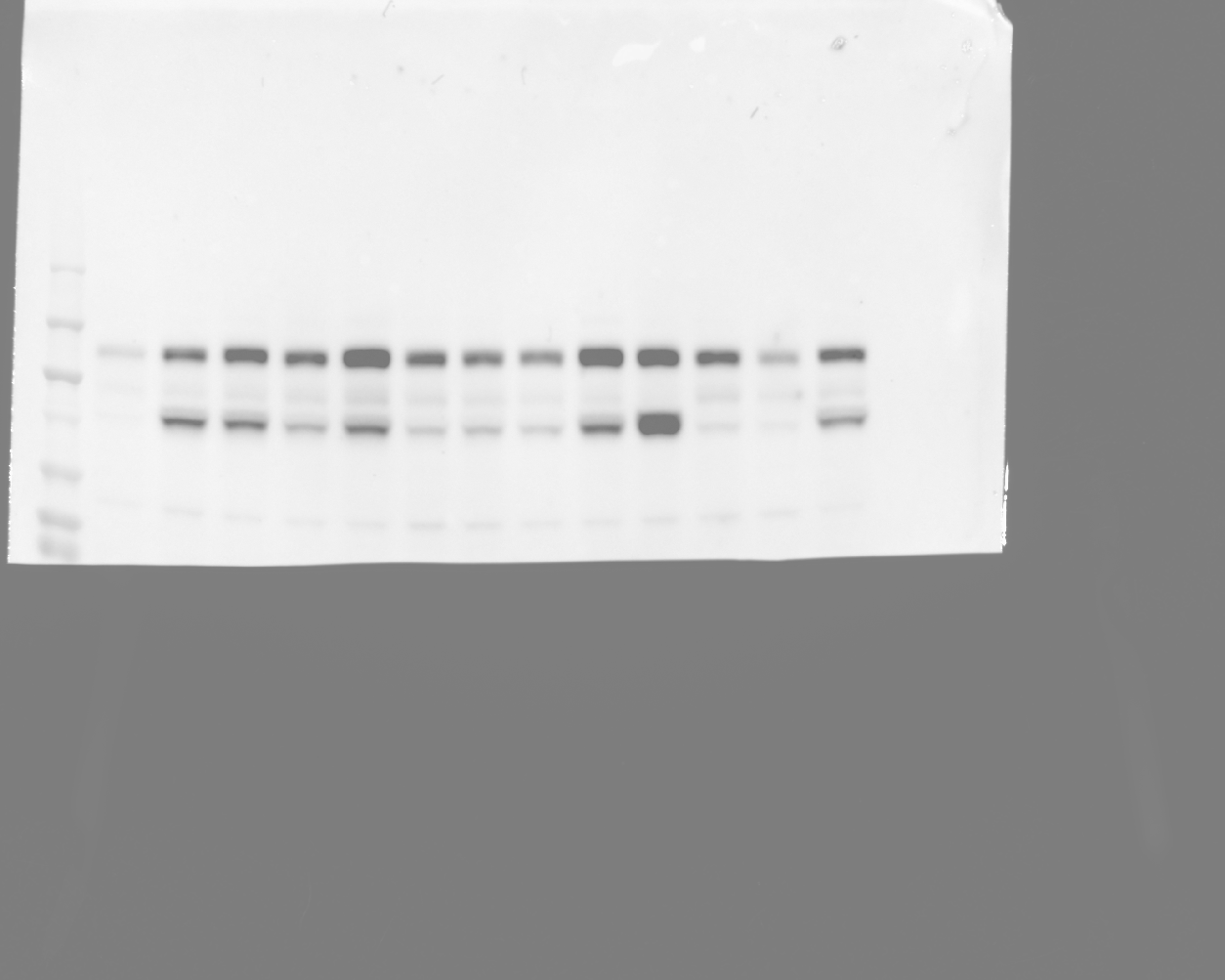

### 20240829_hek_phosphogel_5_linkervars_gel1_stat5_phospho_11(Chemiluminescence).tif

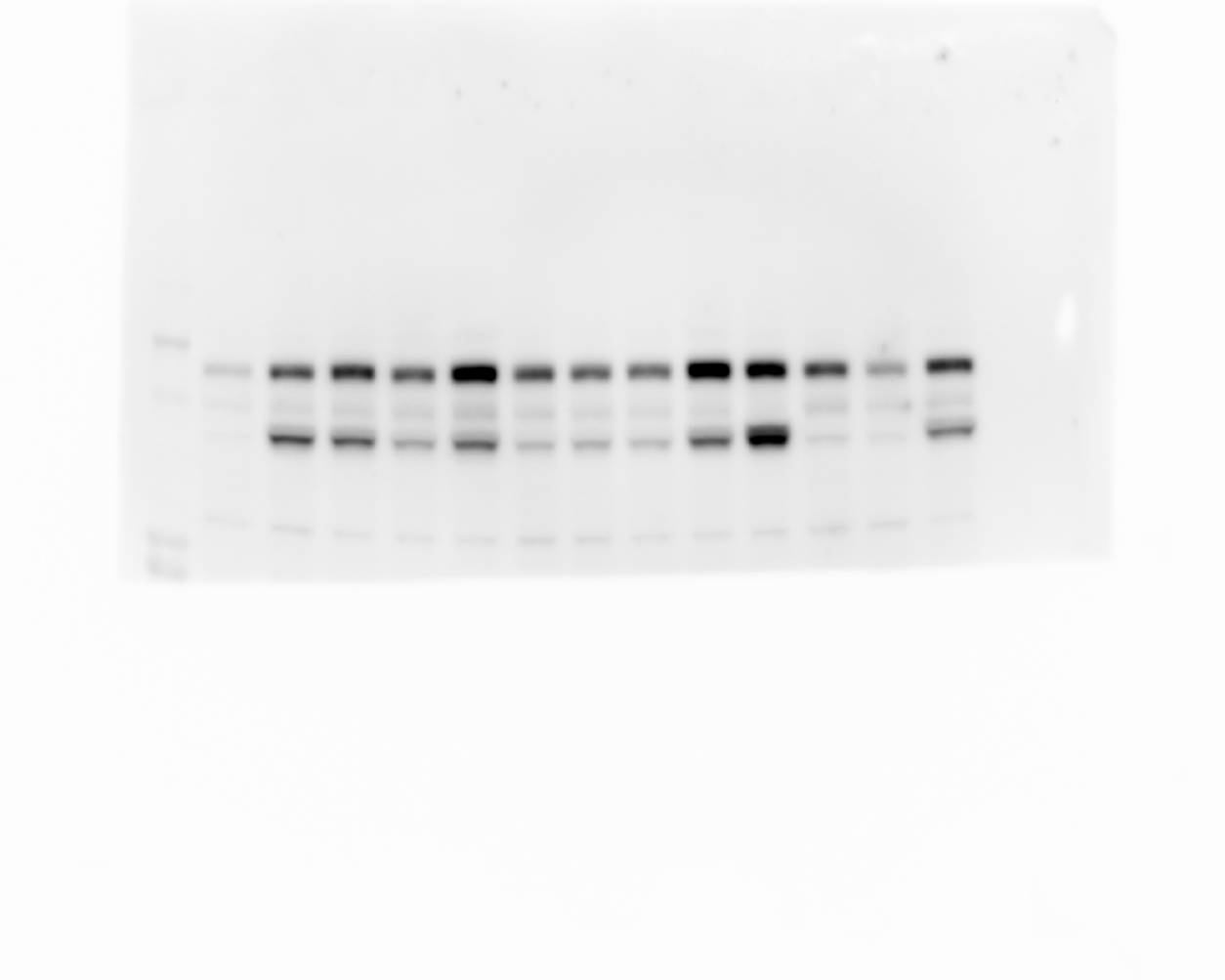

### 20240829_hek_phosphogel_5_linkervars_gel1_stat5_phospho_11(Colorimetric).tif

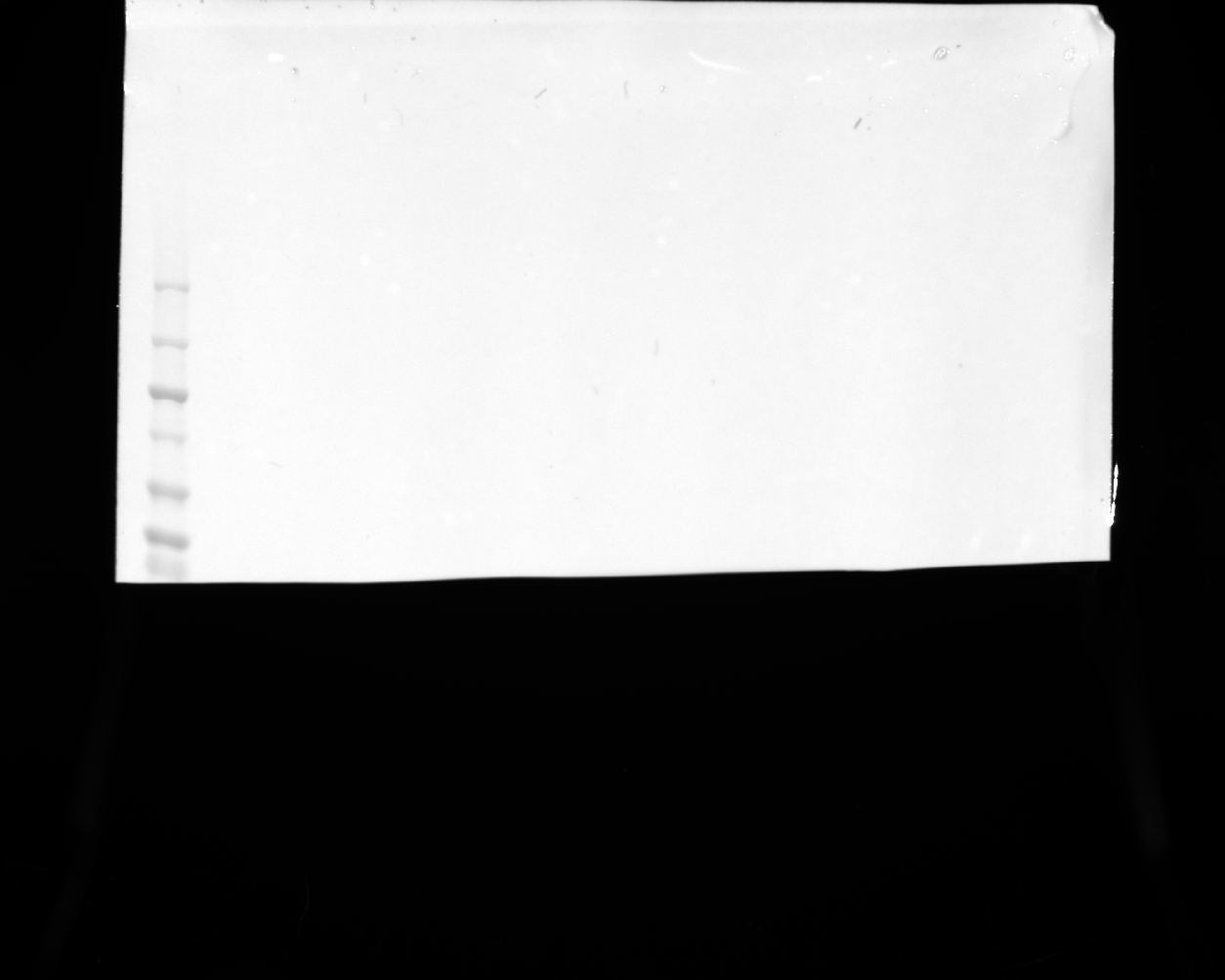

### 20240829_hek_phosphogel_5_linkervars_gel1_stat5_phospho_11(Composite)_lowexp.tif

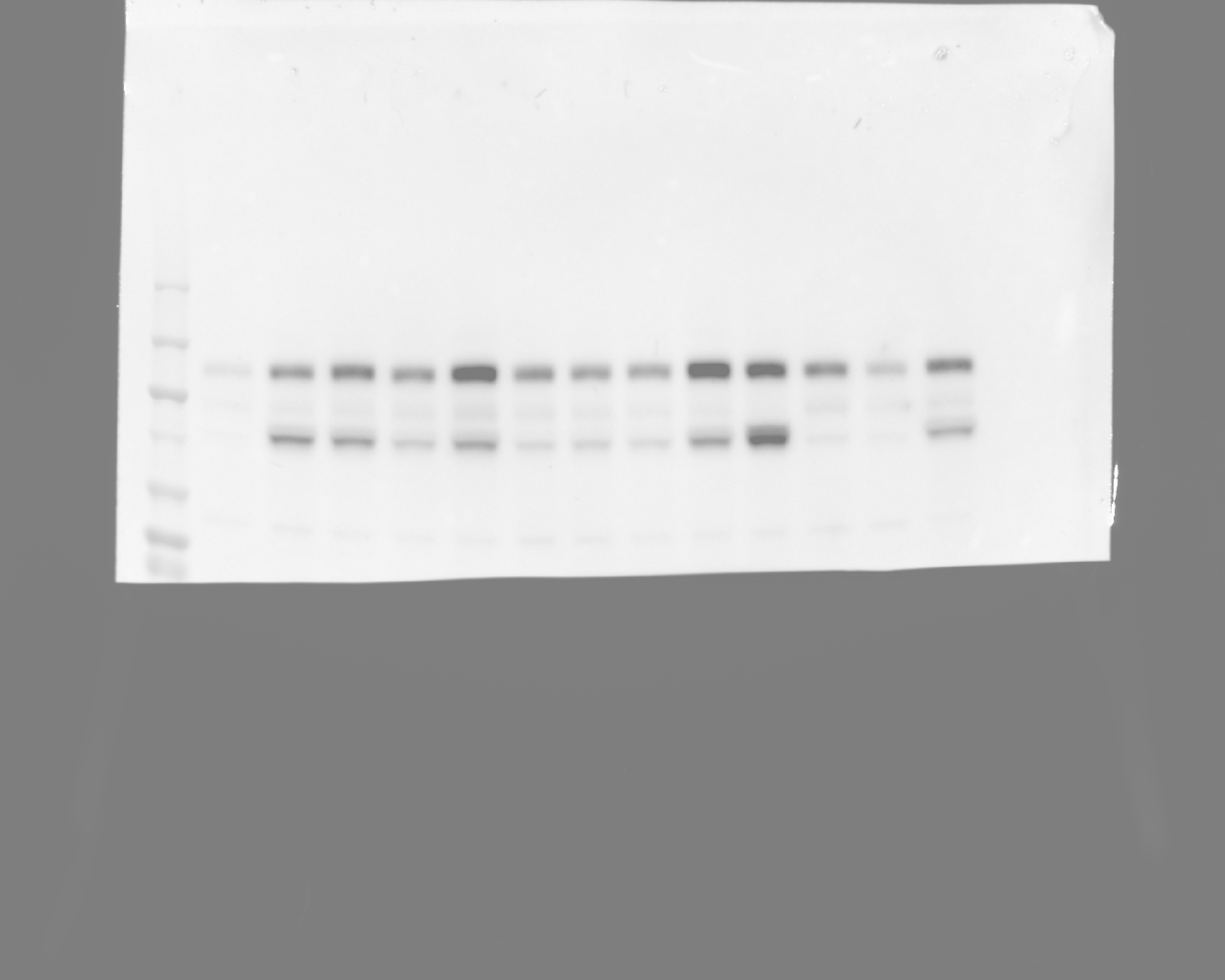

### 20240829_hek_phosphogel_5_linkervars_gel2_hh3_5(Chemiluminescence).tif

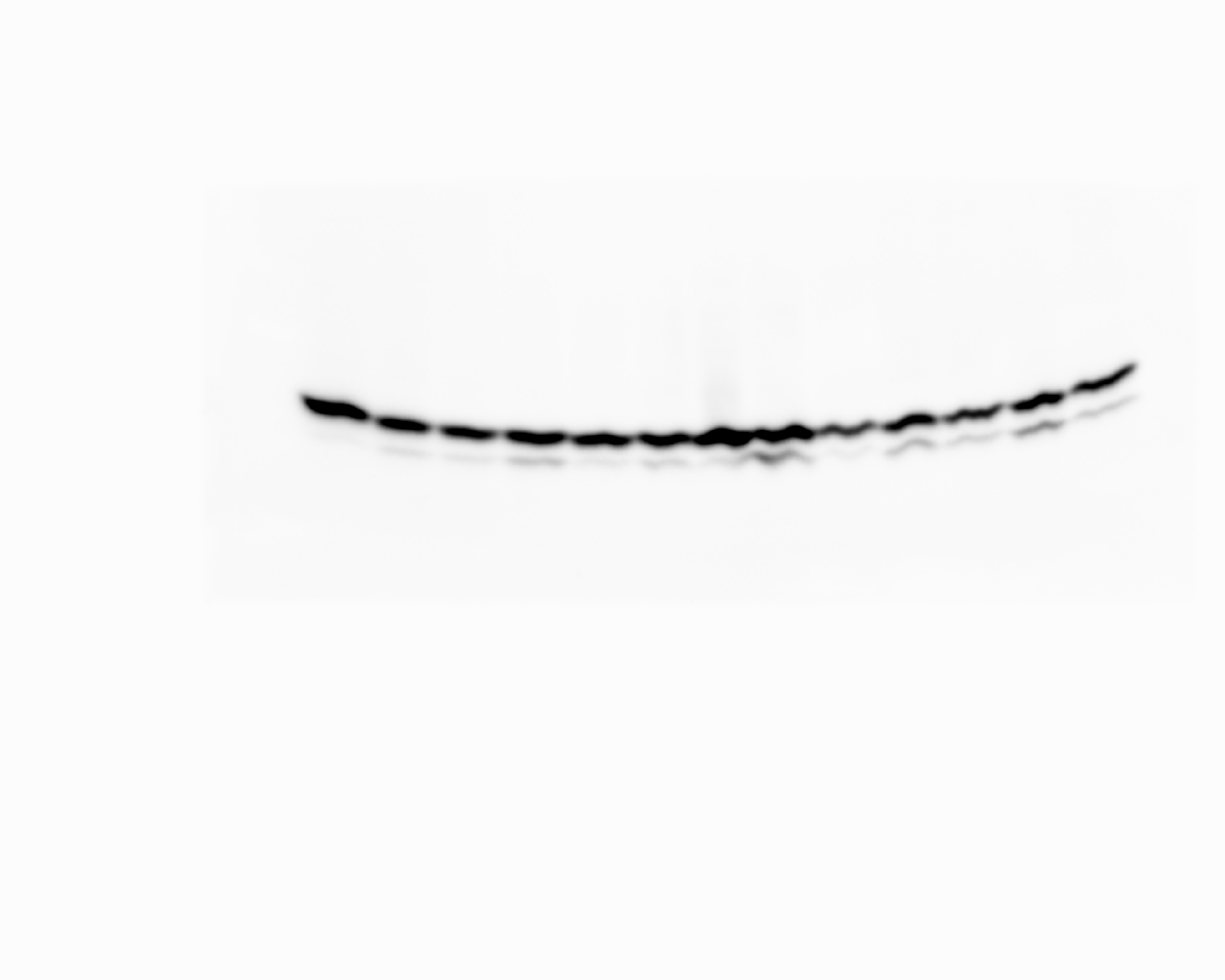

### 20240829_hek_phosphogel_5_linkervars_gel2_hh3_5(Colorimetric).tif

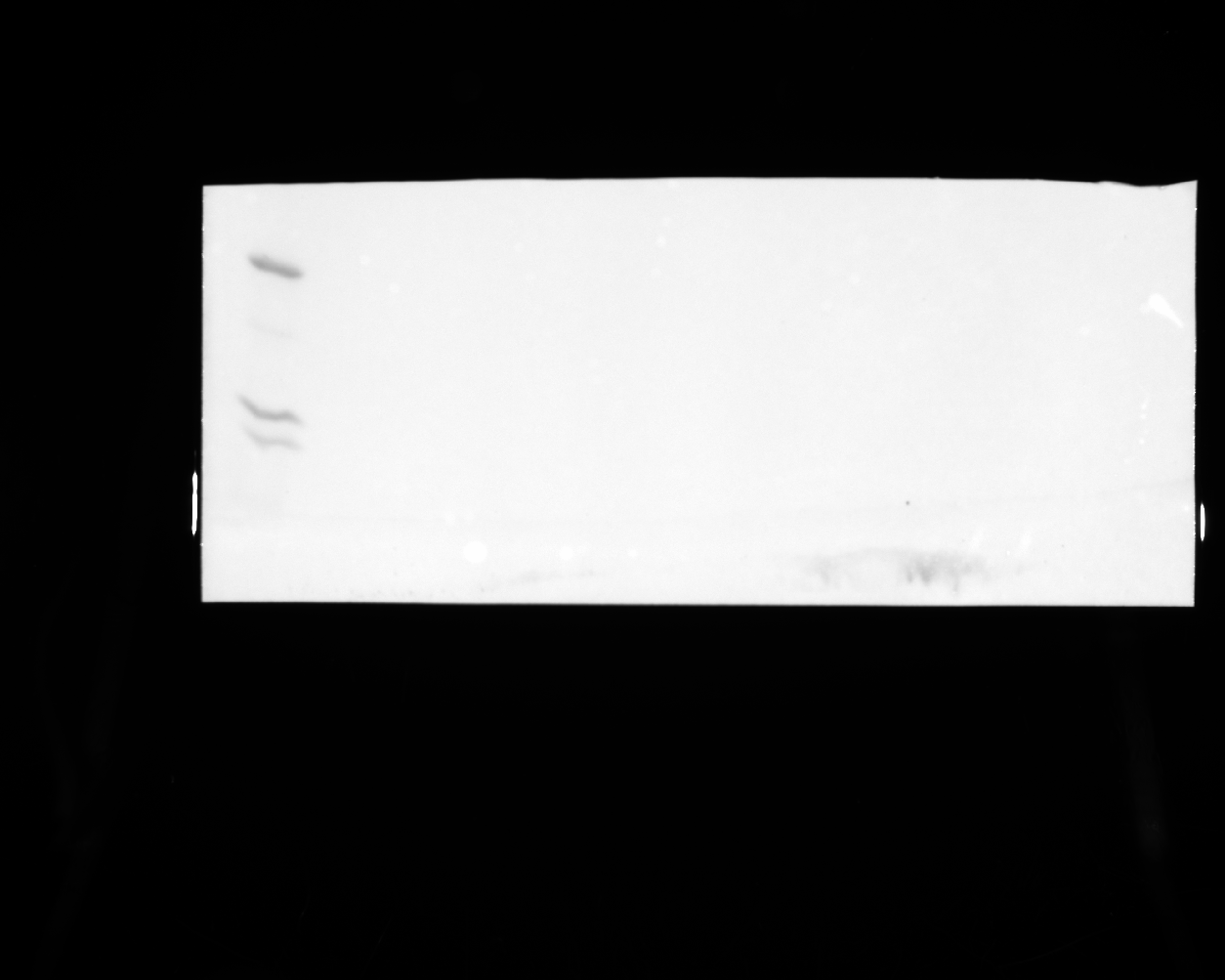

### 20240829_hek_phosphogel_5_linkervars_gel2_hh3_5(Composite).tif

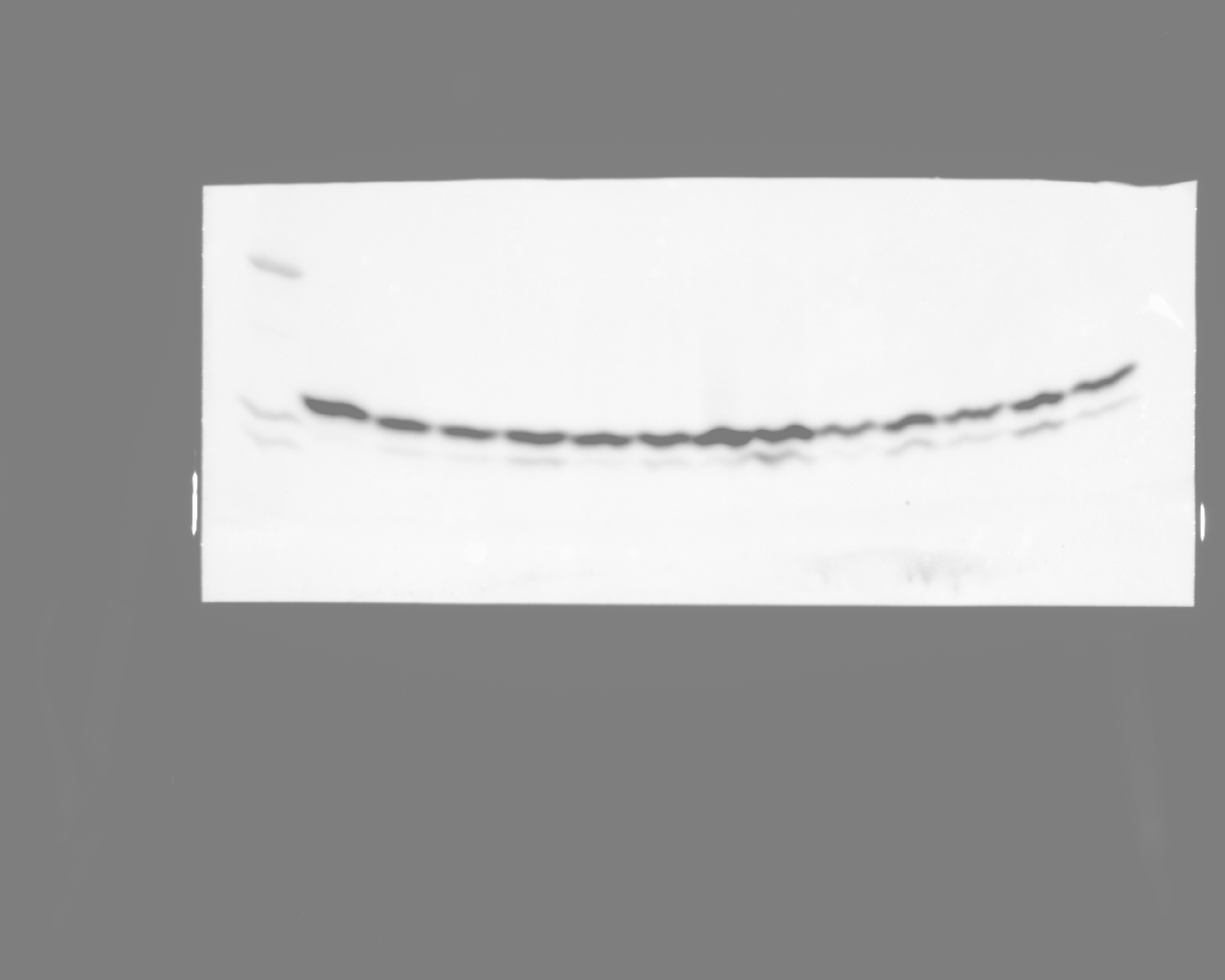

### 20240829_hek_phosphogel_5_linkervars_gel2_stat5_8(Chemiluminescence).tif

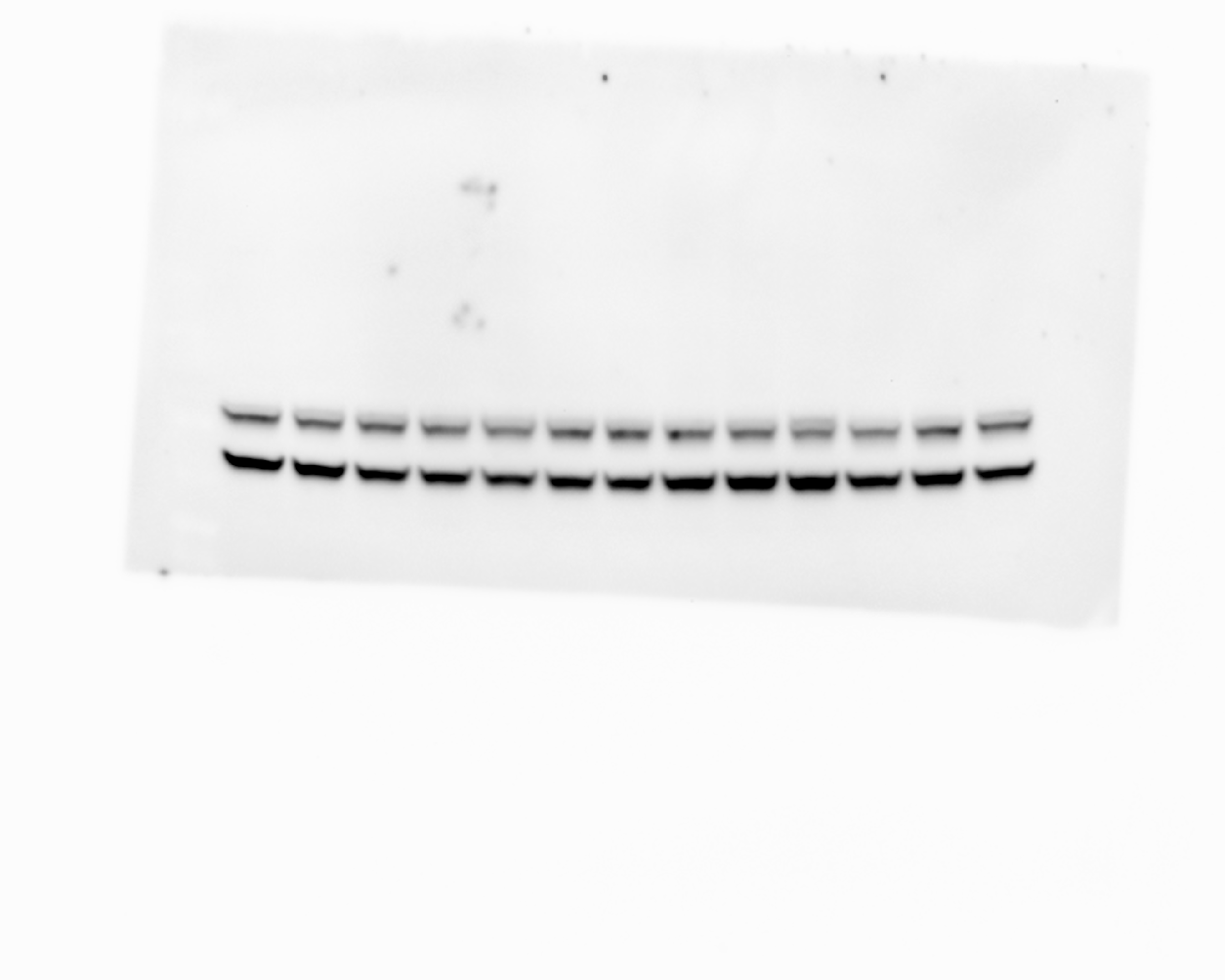

### 20240829_hek_phosphogel_5_linkervars_gel2_stat5_8(Colorimetric).tif

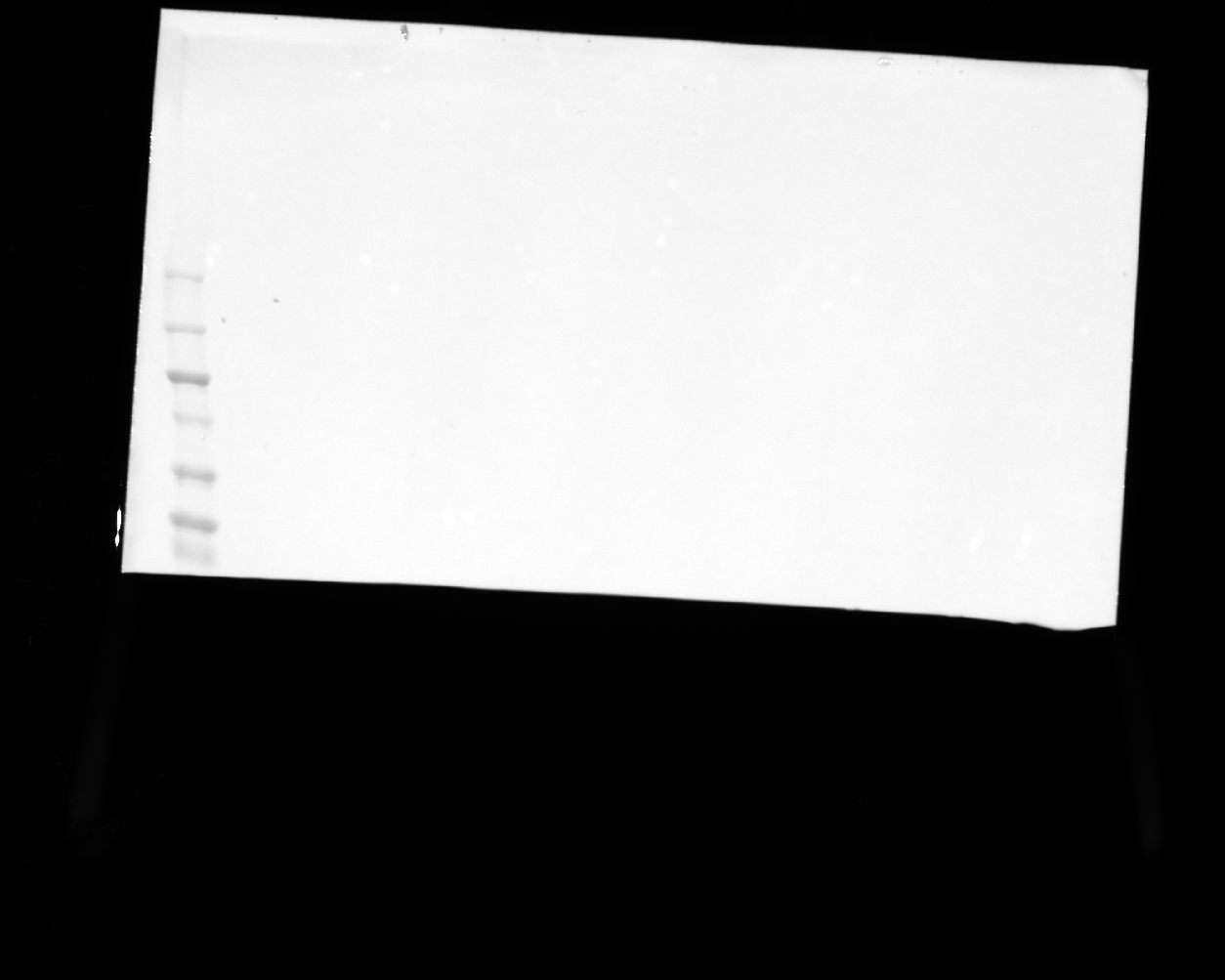

### 20240829_hek_phosphogel_5_linkervars_gel2_stat5_8(Composite).tif

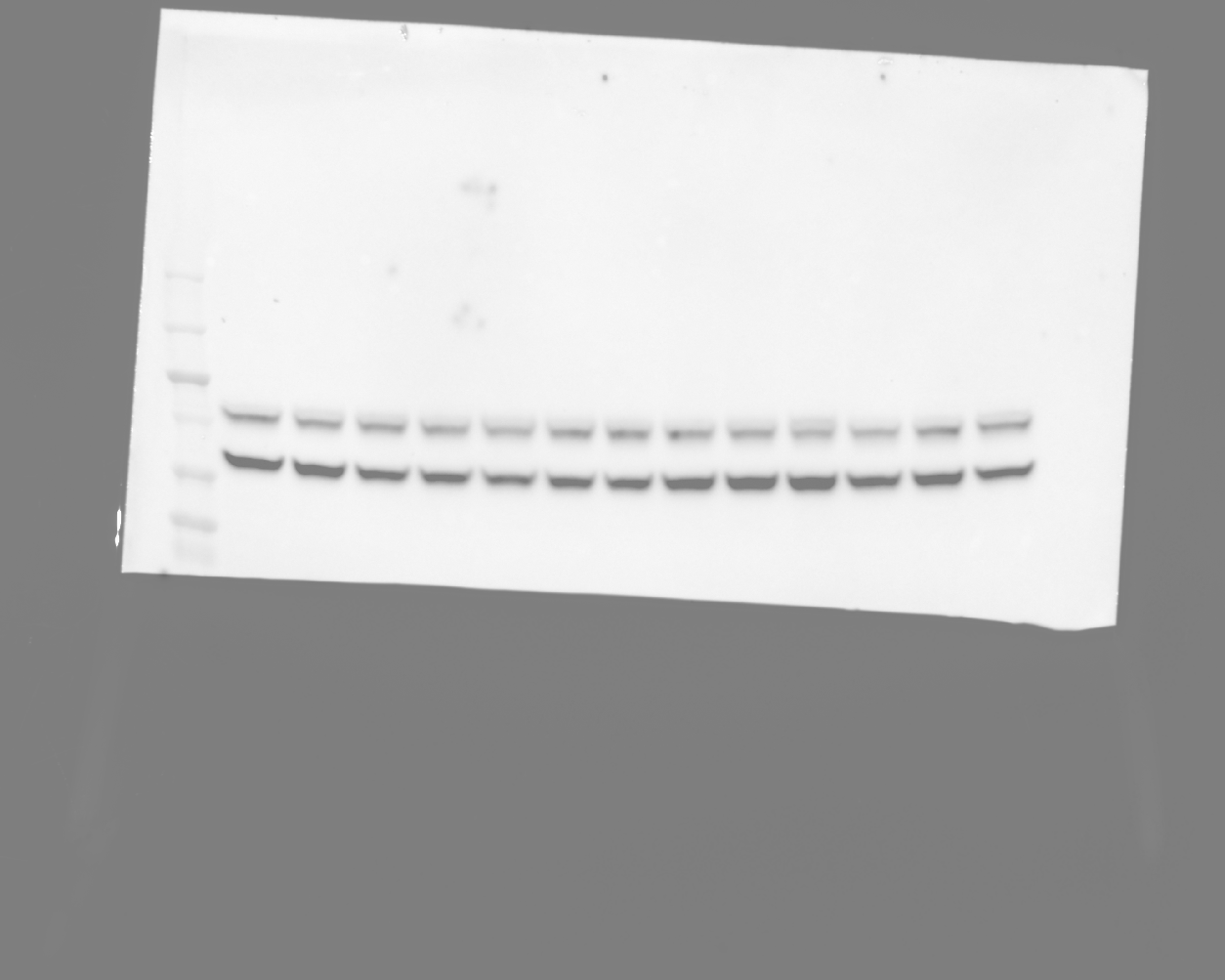

### 20240829_hek_phosphogel_5_linkervars_gel4_hh3_12(Chemiluminescence).tif

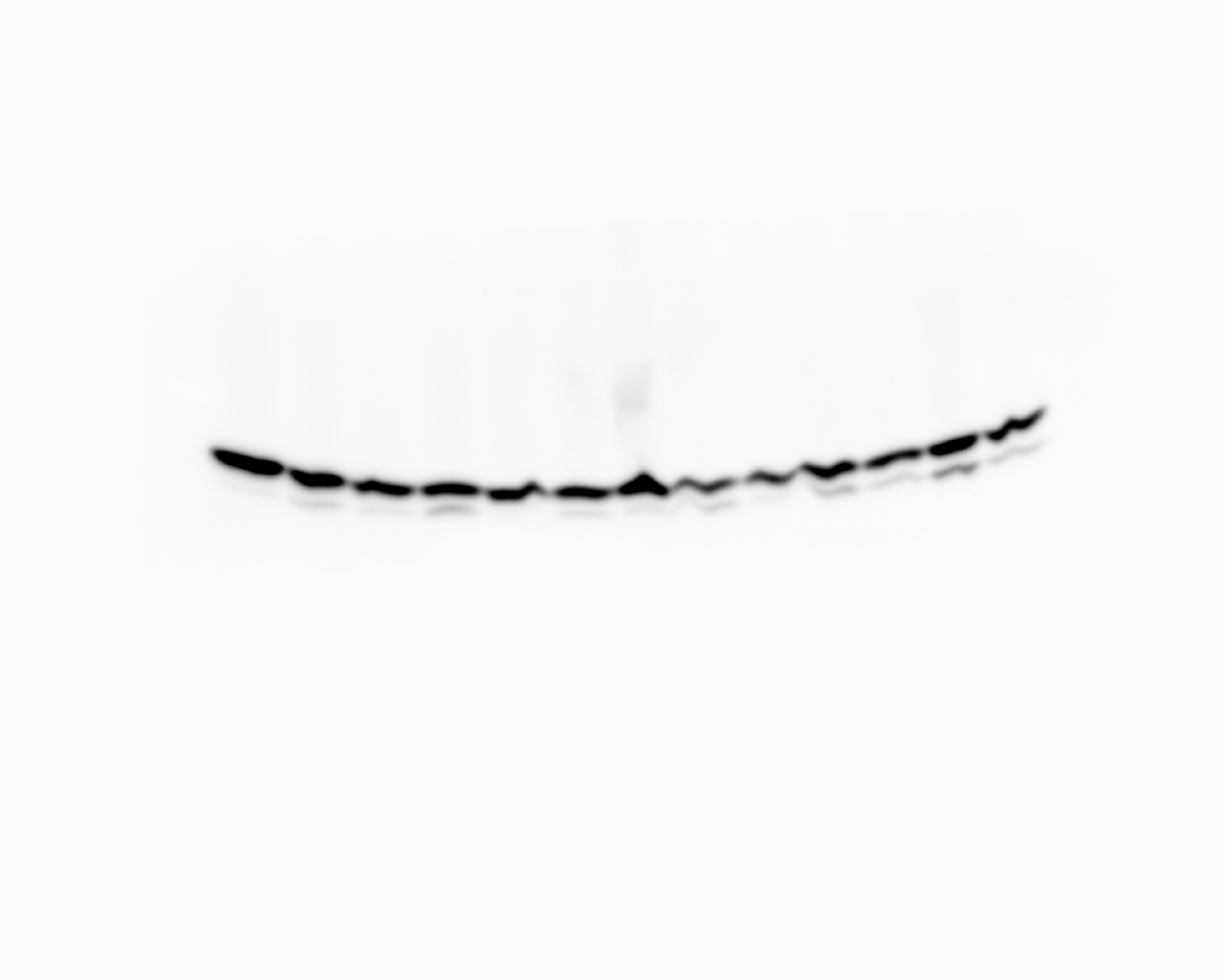

### 20240829_hek_phosphogel_5_linkervars_gel4_hh3_12(Colorimetric).tif

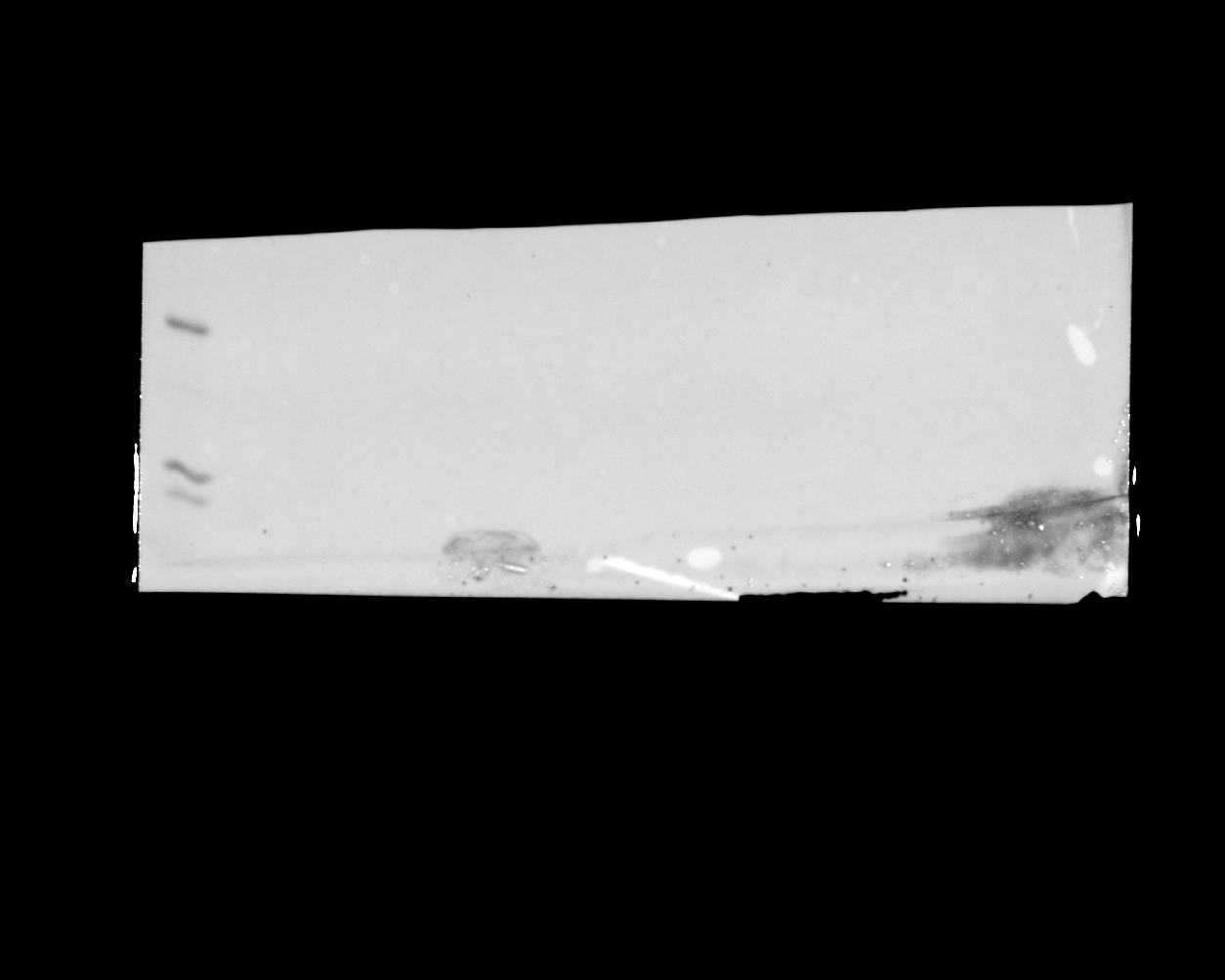

### 20240829_hek_phosphogel_5_linkervars_gel4_hh3_12(Composite).tif

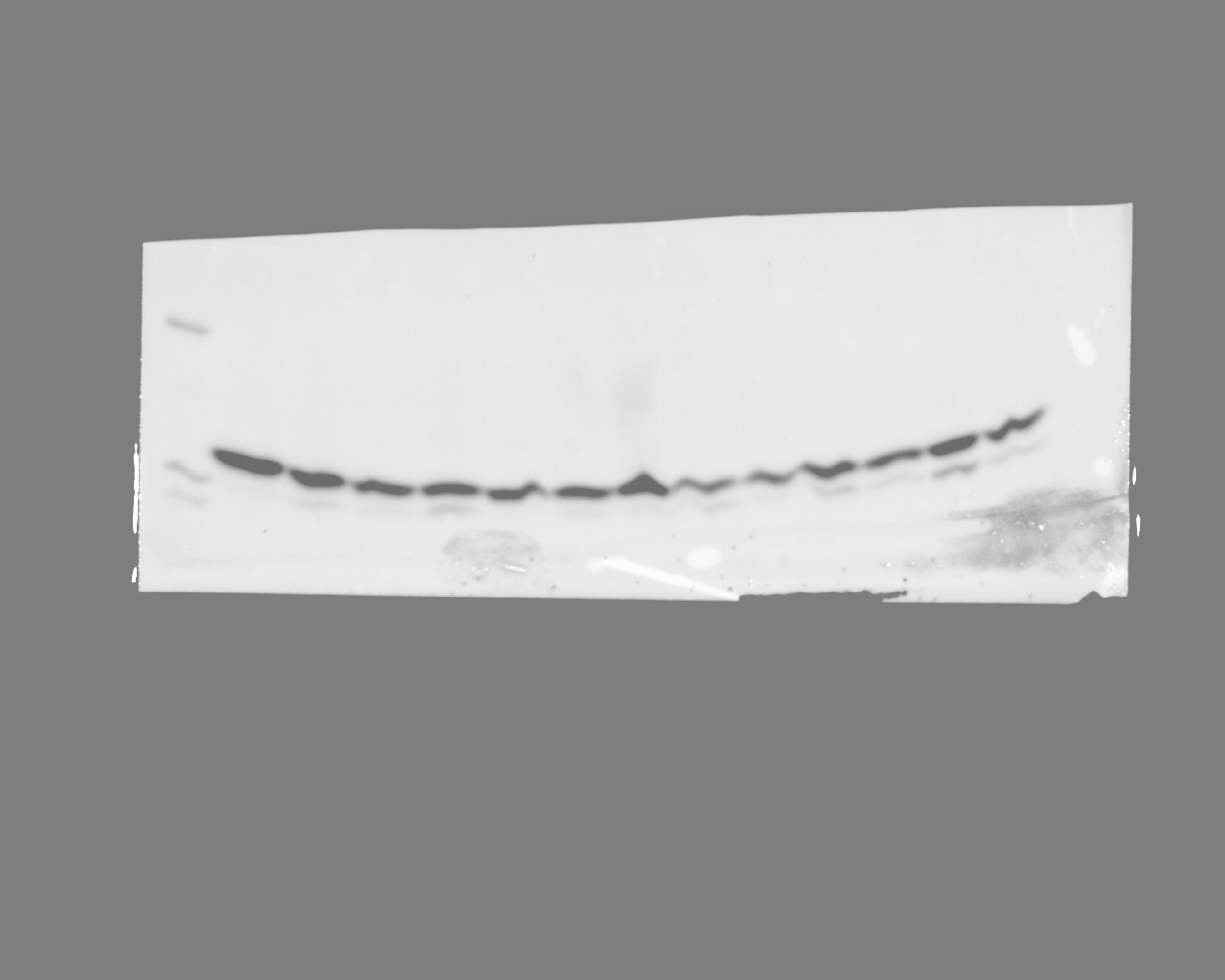

### 20240829_hek_phosphogel_5_linkervars_ha488_4(DyLight 488).tif

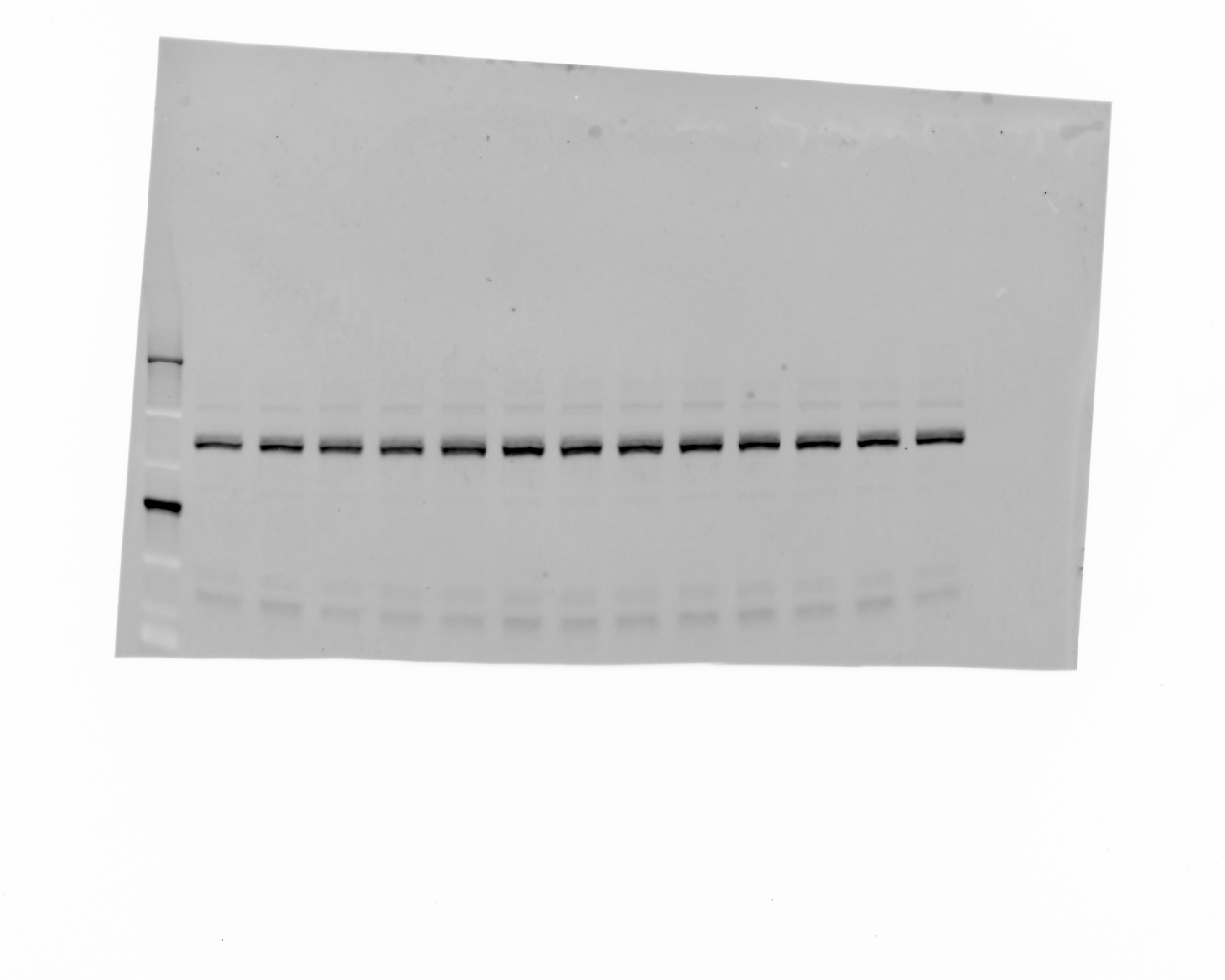

### 20240829_hek_phosphogel_5_linkervars_ha488_6(Colorimetric).tif

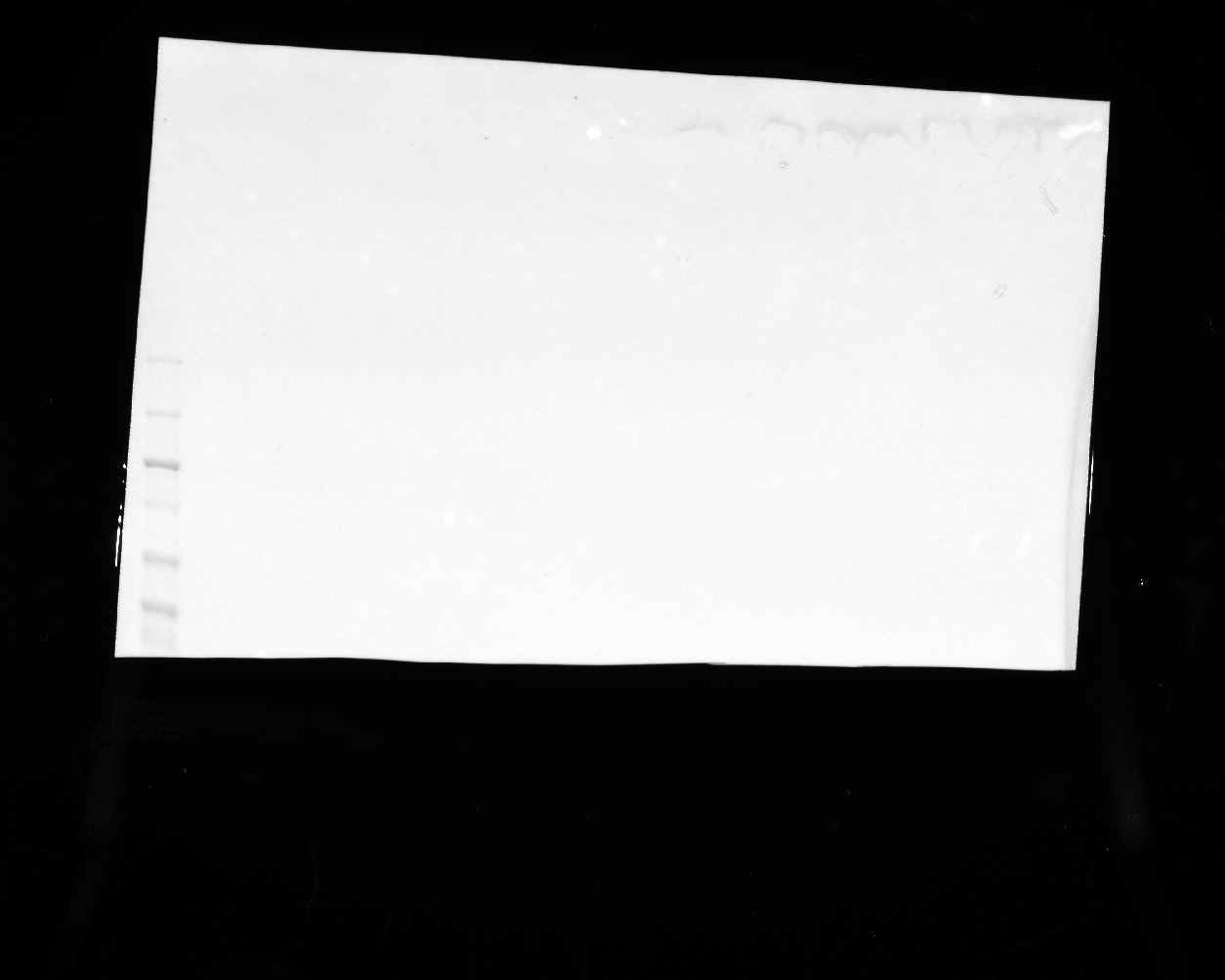

### 20241114_phosphogel2_phosphogel3_paperrerun_gel2_stat5_16(Chemiluminescence).tif

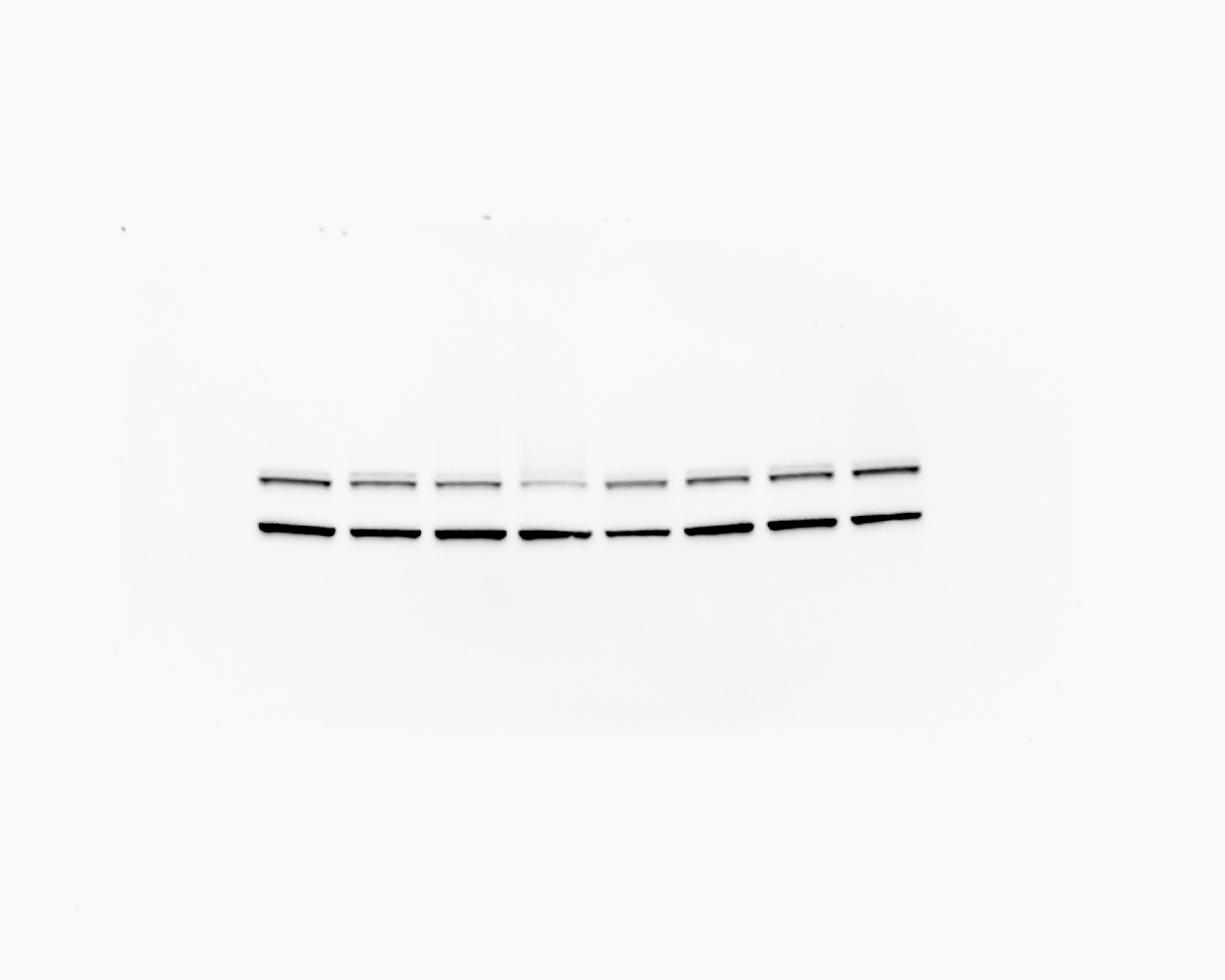

### 20241114_phosphogel2_phosphogel3_paperrerun_gel2_stat5_16(Colorimetric).tif

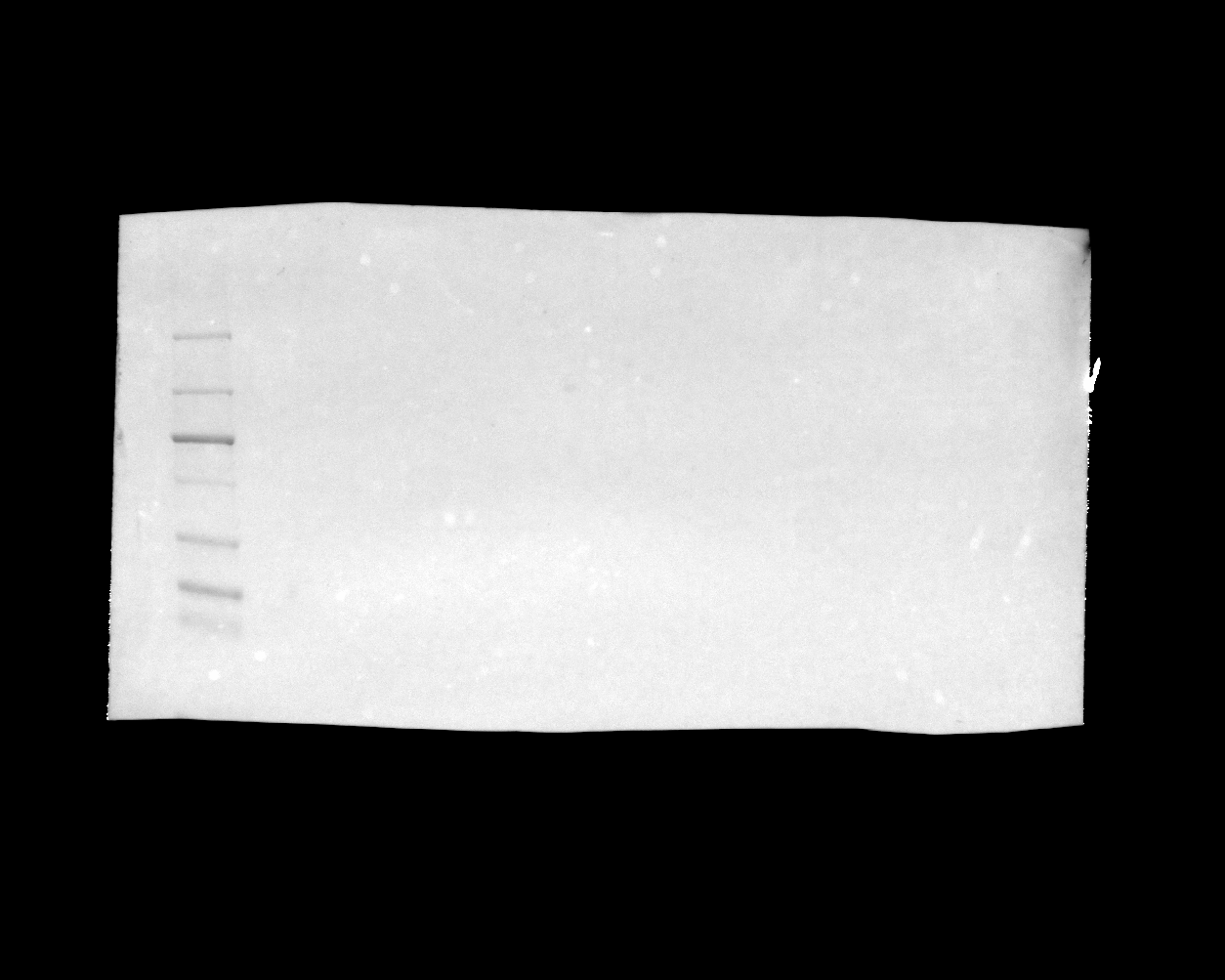

### 20241114_phosphogel2_phosphogel3_paperrerun_gel2_stat5_16(Composite).tif

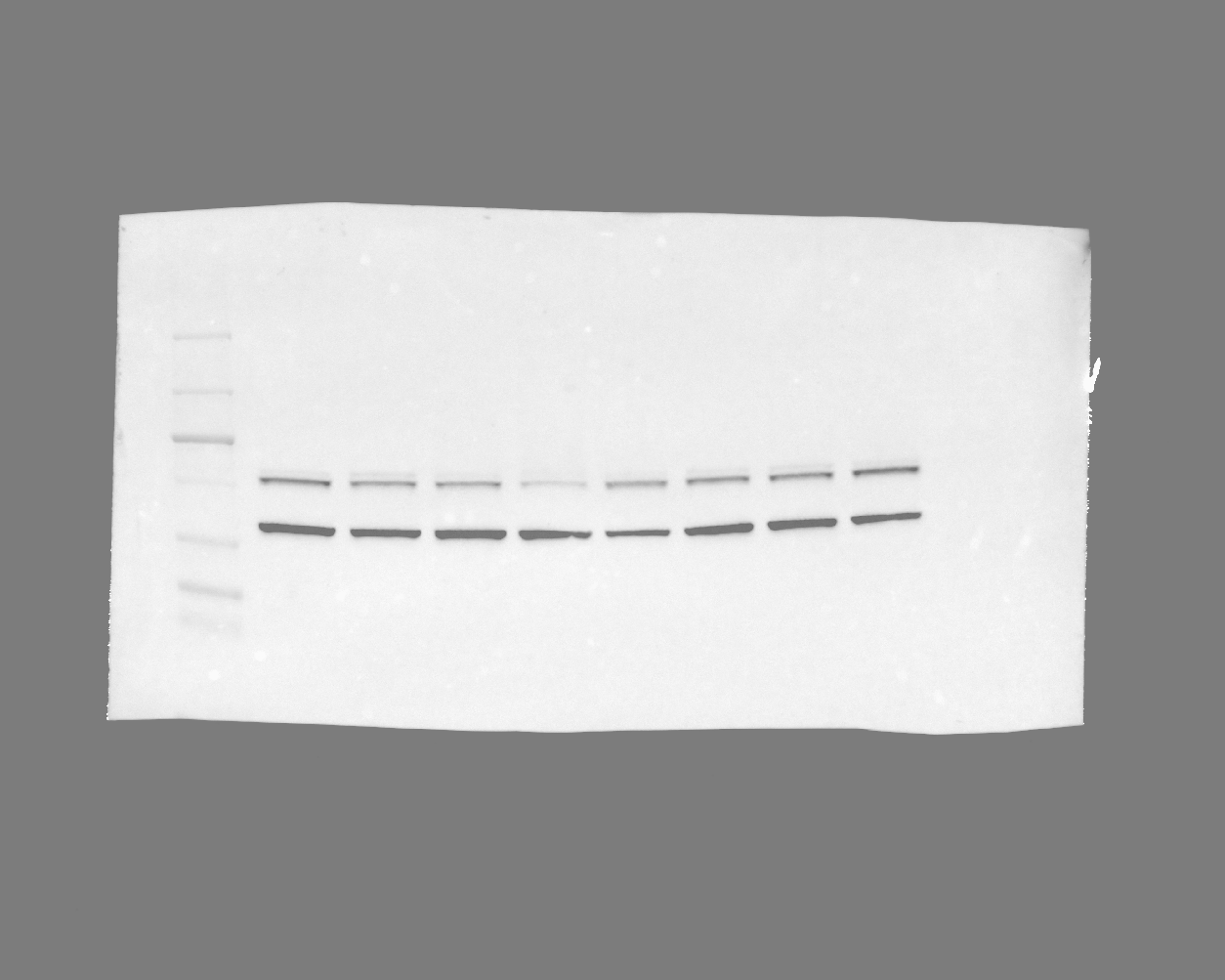

### 20241114_phosphogel2_phosphogel3_paperrerun_gel3_hh3_7(Chemiluminescence).tif

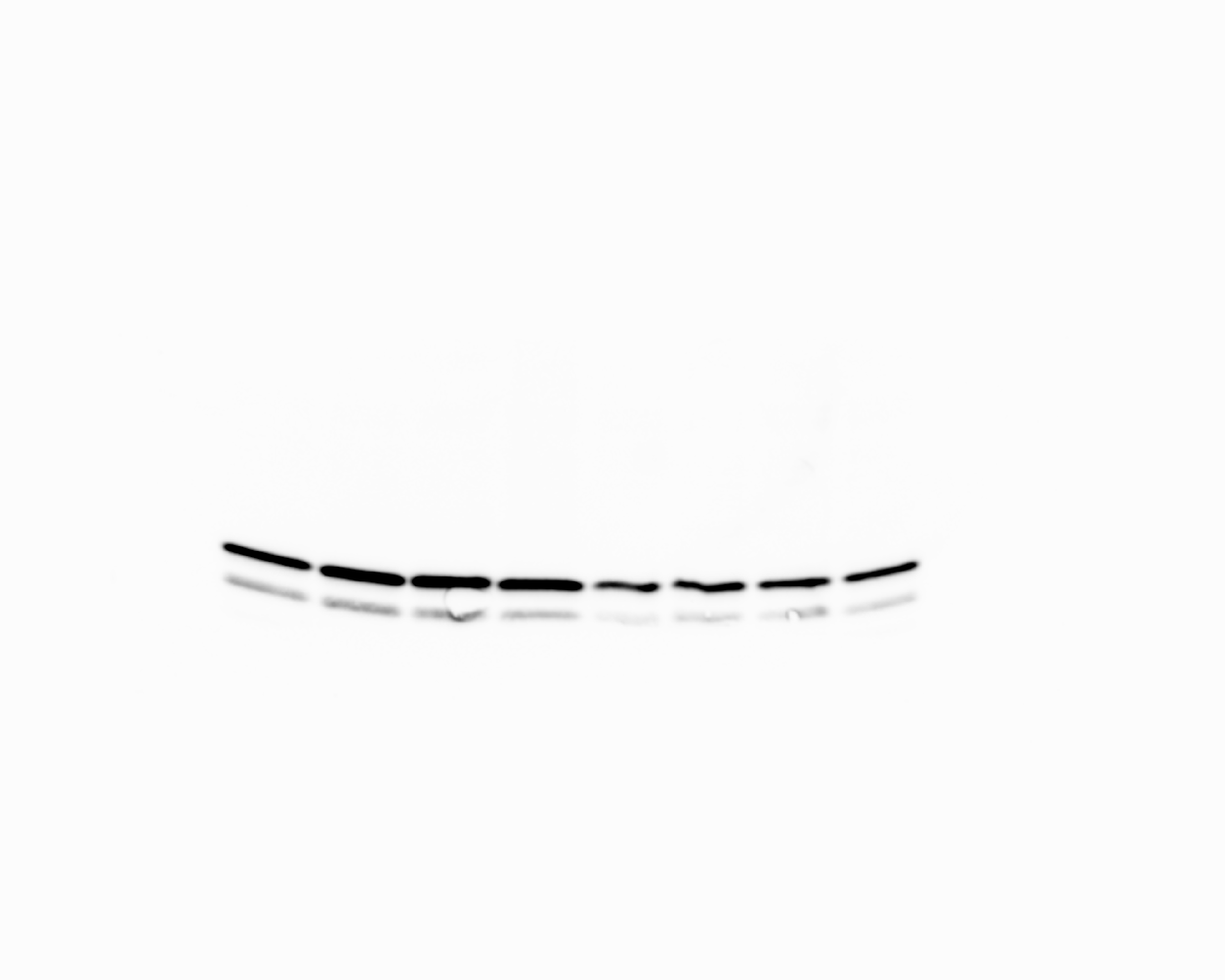

### 20241114_phosphogel2_phosphogel3_paperrerun_gel3_hh3_7(Colorimetric).tif

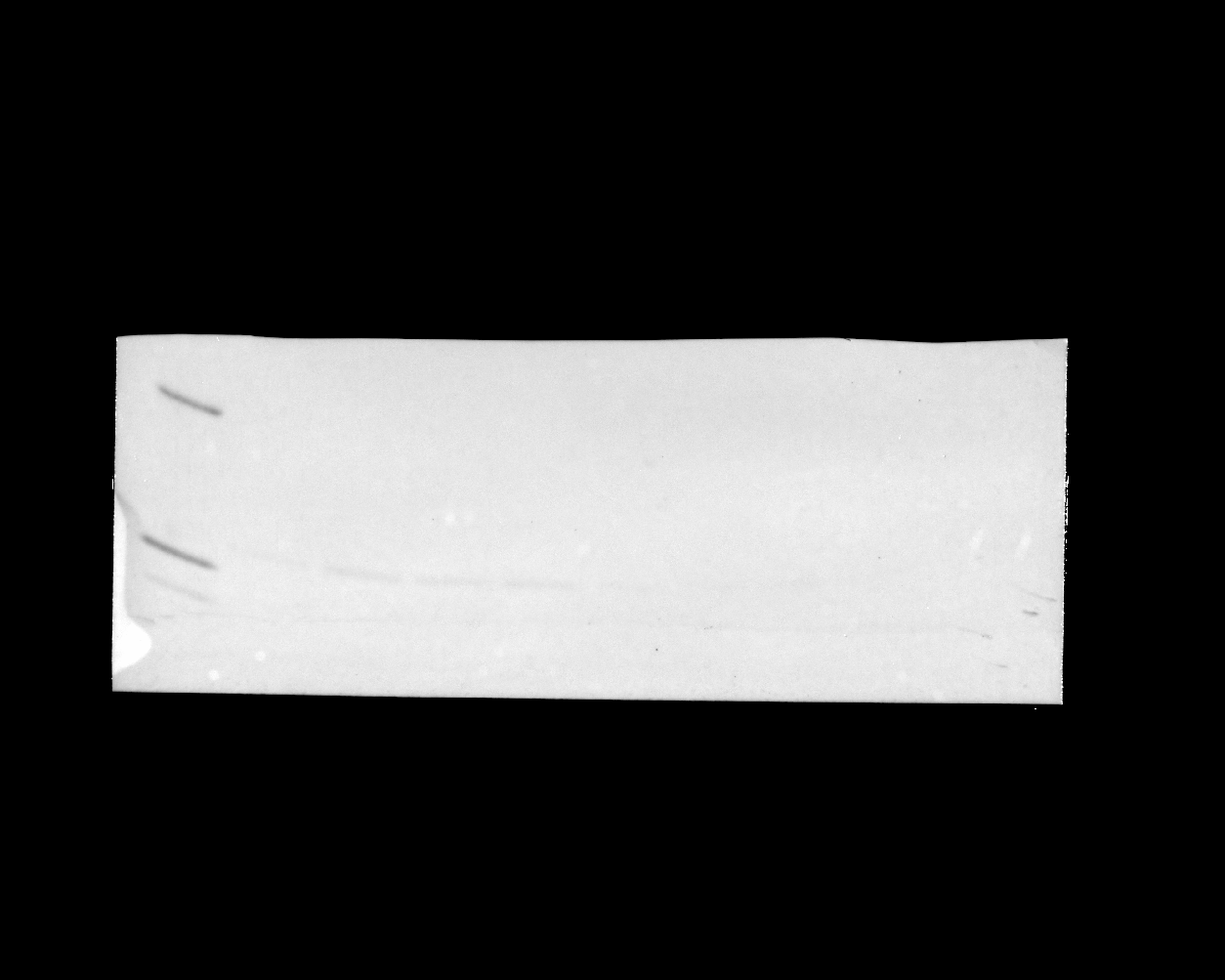

### 20241114_phosphogel2_phosphogel3_paperrerun_gel3_hh3_7(Composite).tif

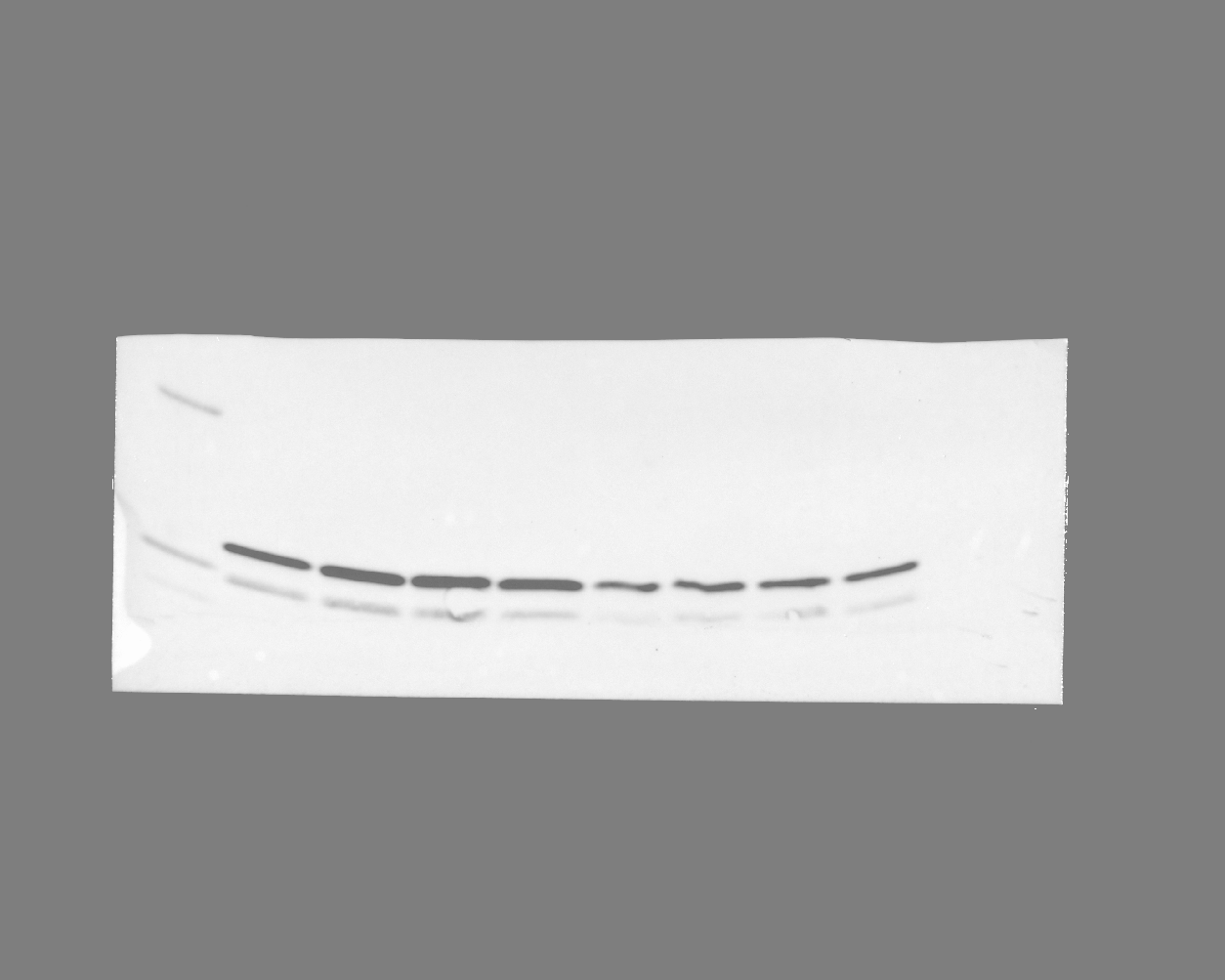

### 20241114_phosphogel2_phosphogel3_paperrerun_gel3_stat5_phospho_11 (Multichannel)_final2.tif

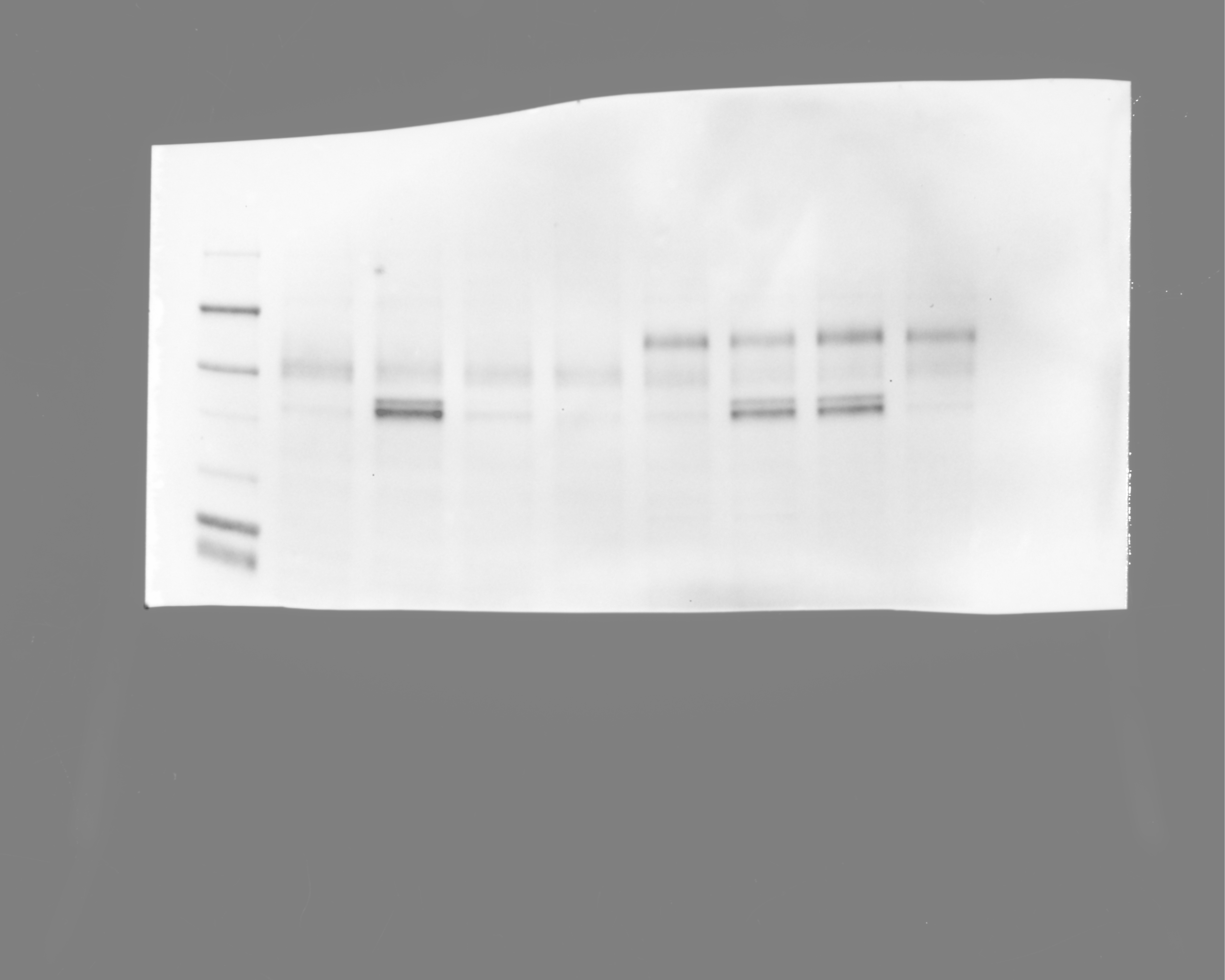

### 20241114_phosphogel2_phosphogel3_paperrerun_gel3_stat5_phospho_11 (Multichannel)_final.tif

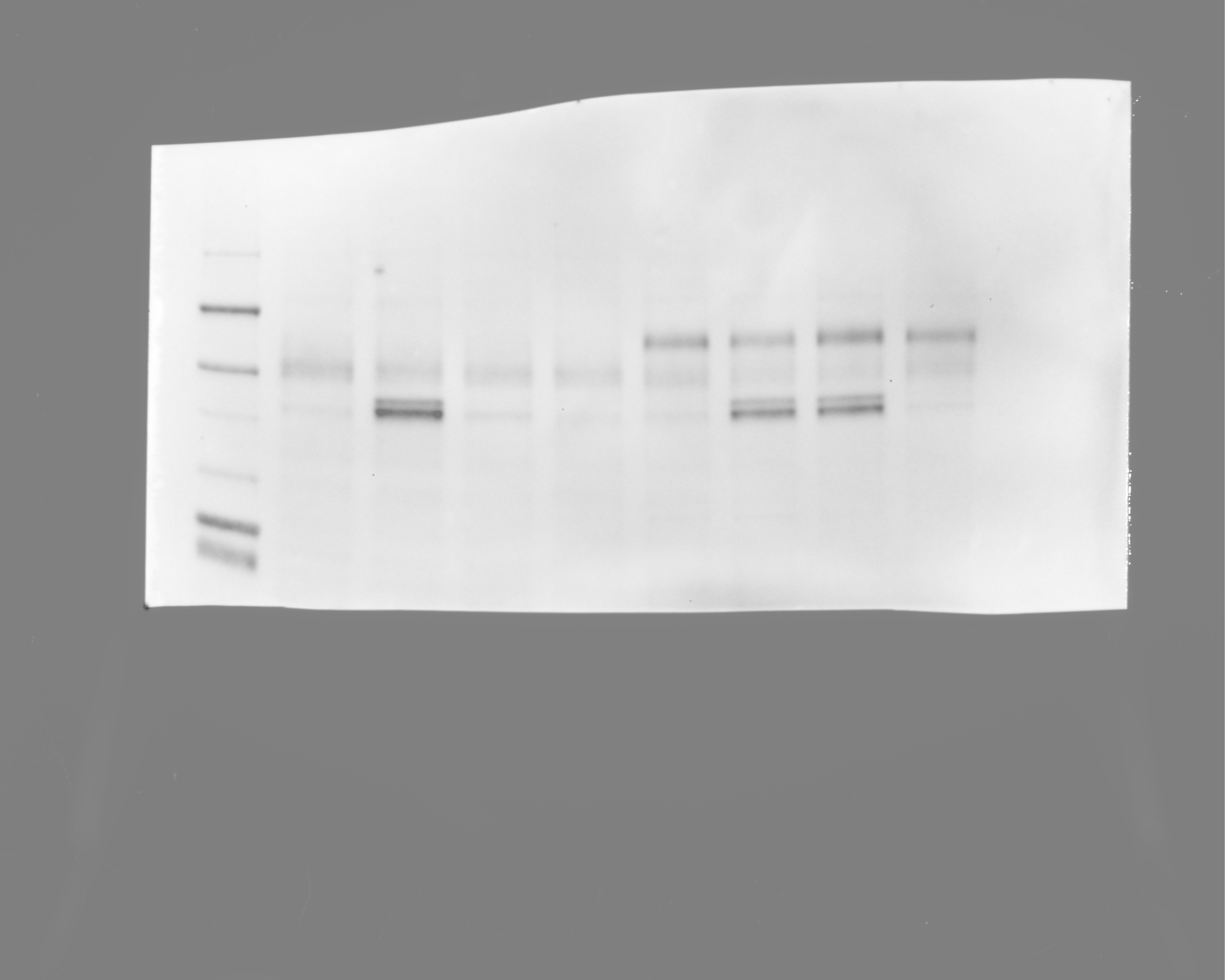

### 20241114_phosphogel2_phosphogel3_paperrerun_gel3_stat5_phospho_11(Chemiluminescence).tif

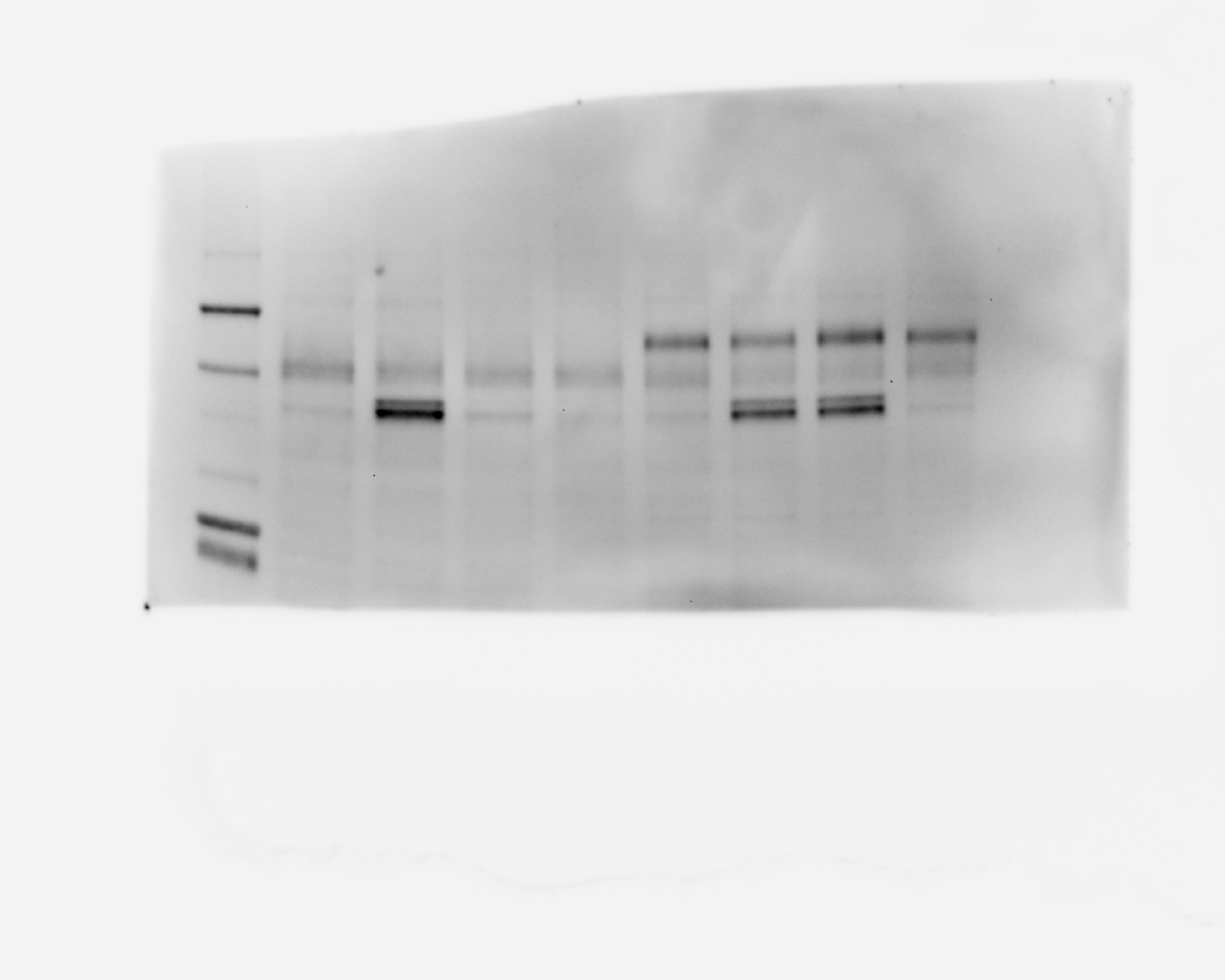

### 20241114_phosphogel2_phosphogel3_paperrerun_gel3_stat5_phospho_11(Colorimetric).tif

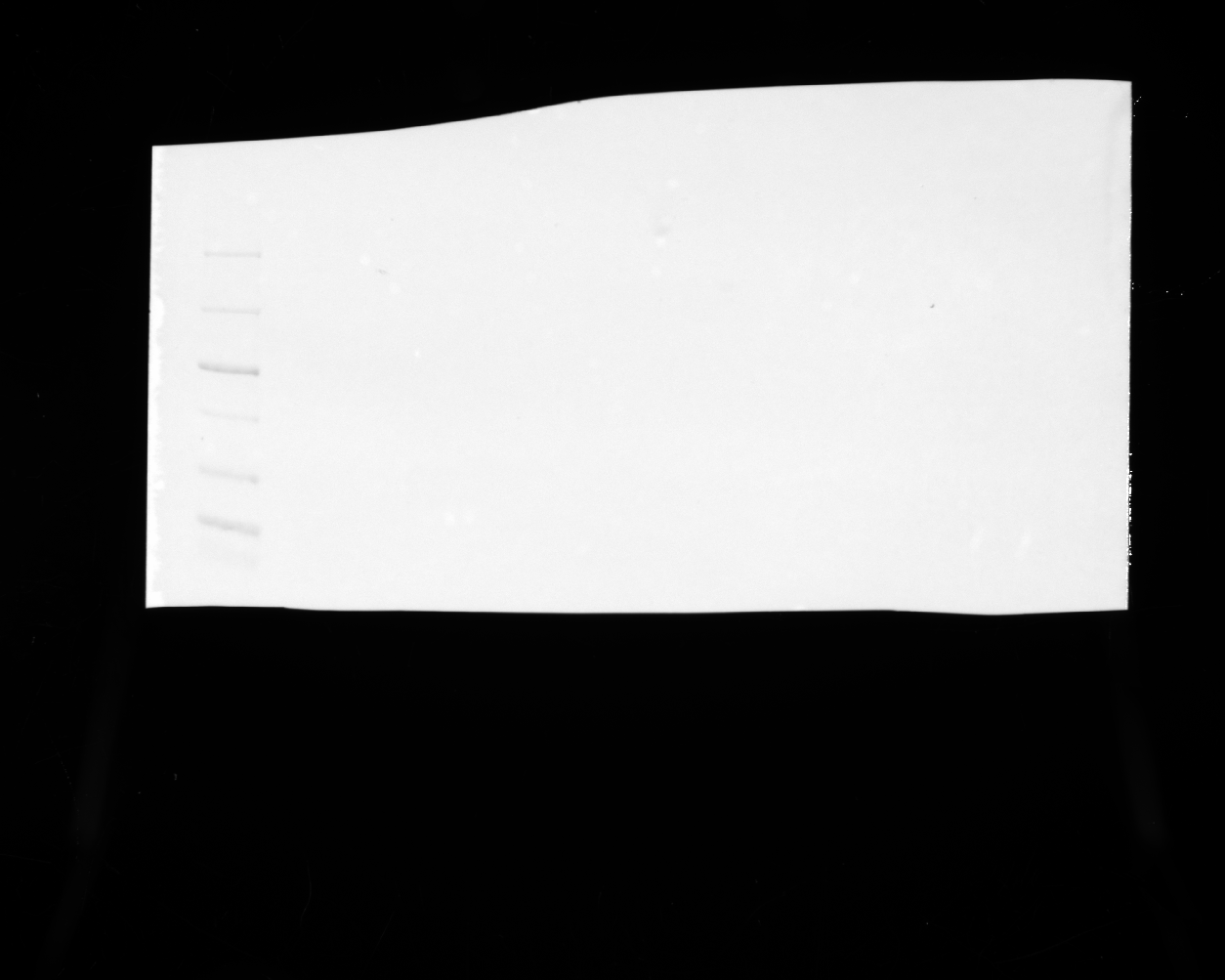

### 20241114_phosphogel2_phosphogel3_paperrerun_gel3_stat5_phospho_11(Composite).tif

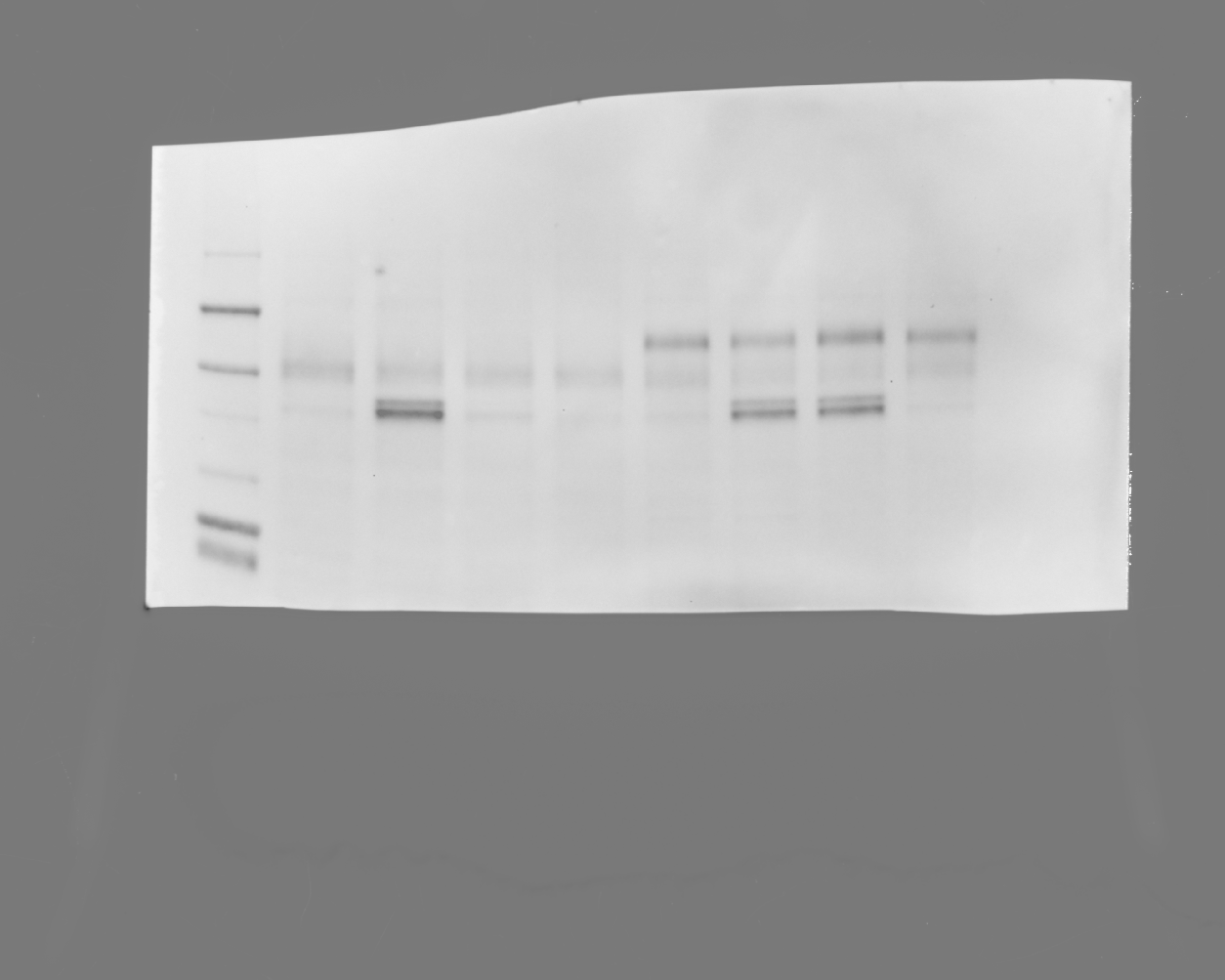

### 20241114_phosphogel2_phosphogel3_paperrerun_gel4_hh3_5(Chemiluminescence).tif

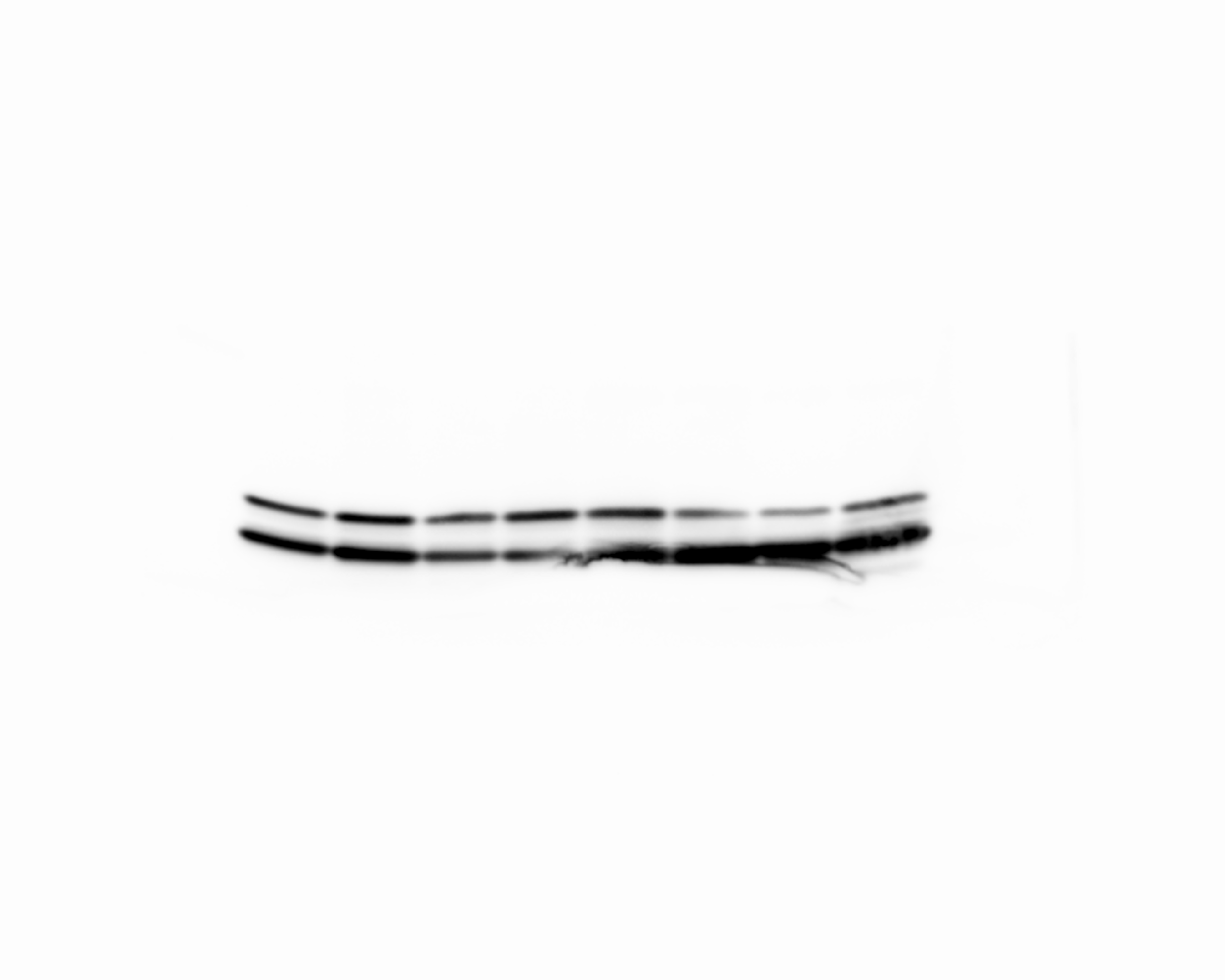

### 20241114_phosphogel2_phosphogel3_paperrerun_gel4_hh3_5(Colorimetric).tif

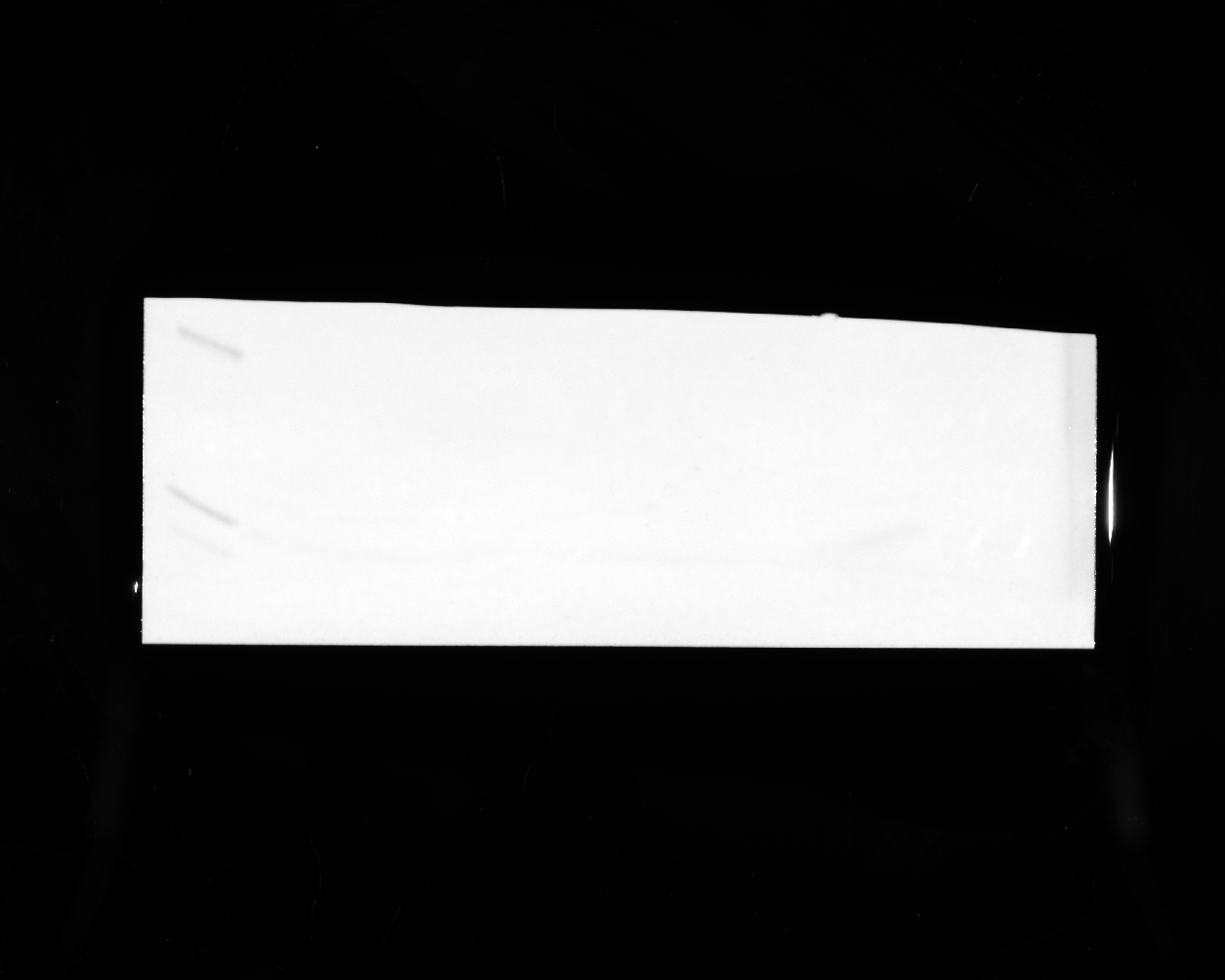

### 20241114_phosphogel2_phosphogel3_paperrerun_gel4_hh3_5(Composite).tif

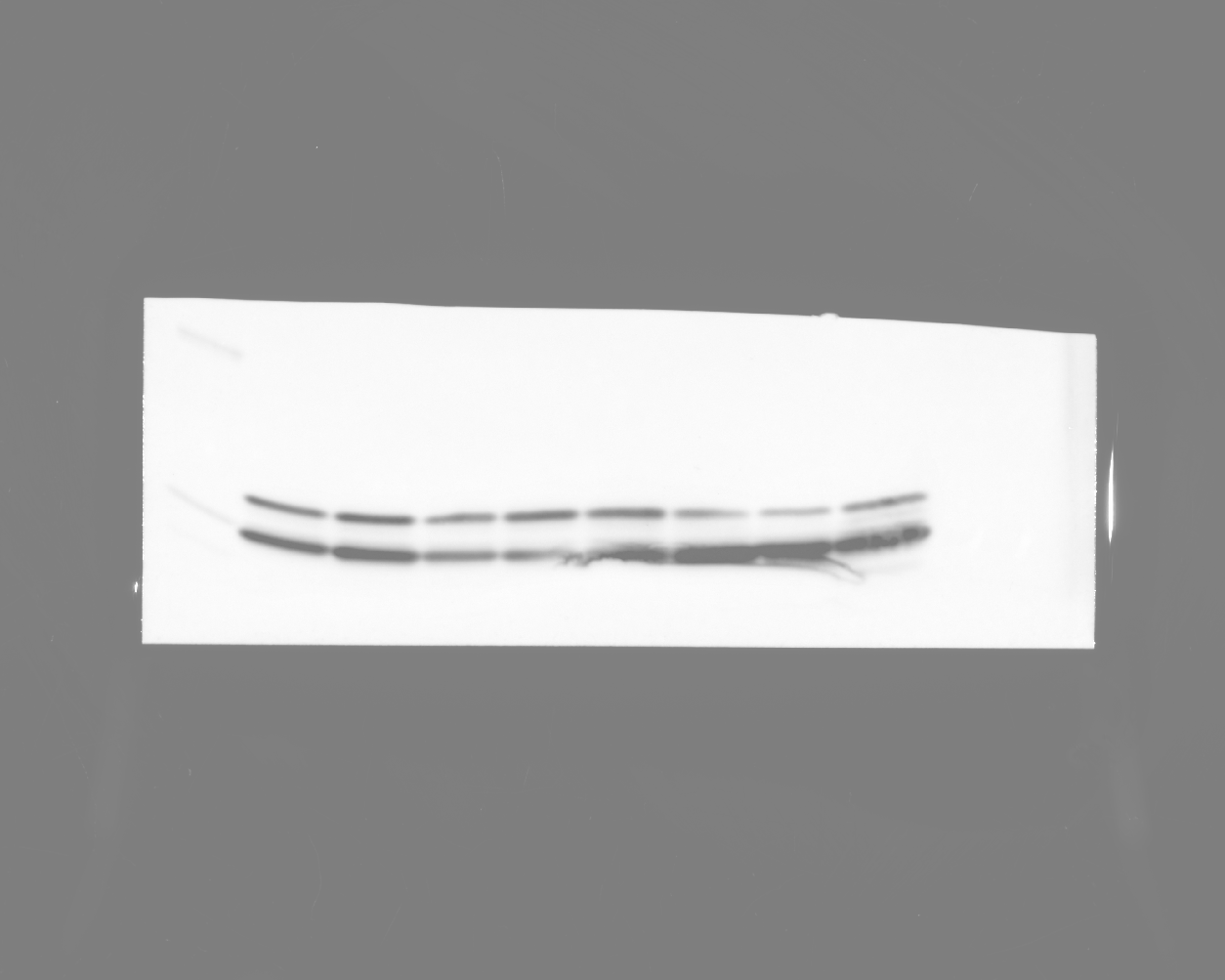

### 20241114_phosphogel2_phosphogel3_paperrerun_gel4_stat5_7(Chemiluminescence).tif

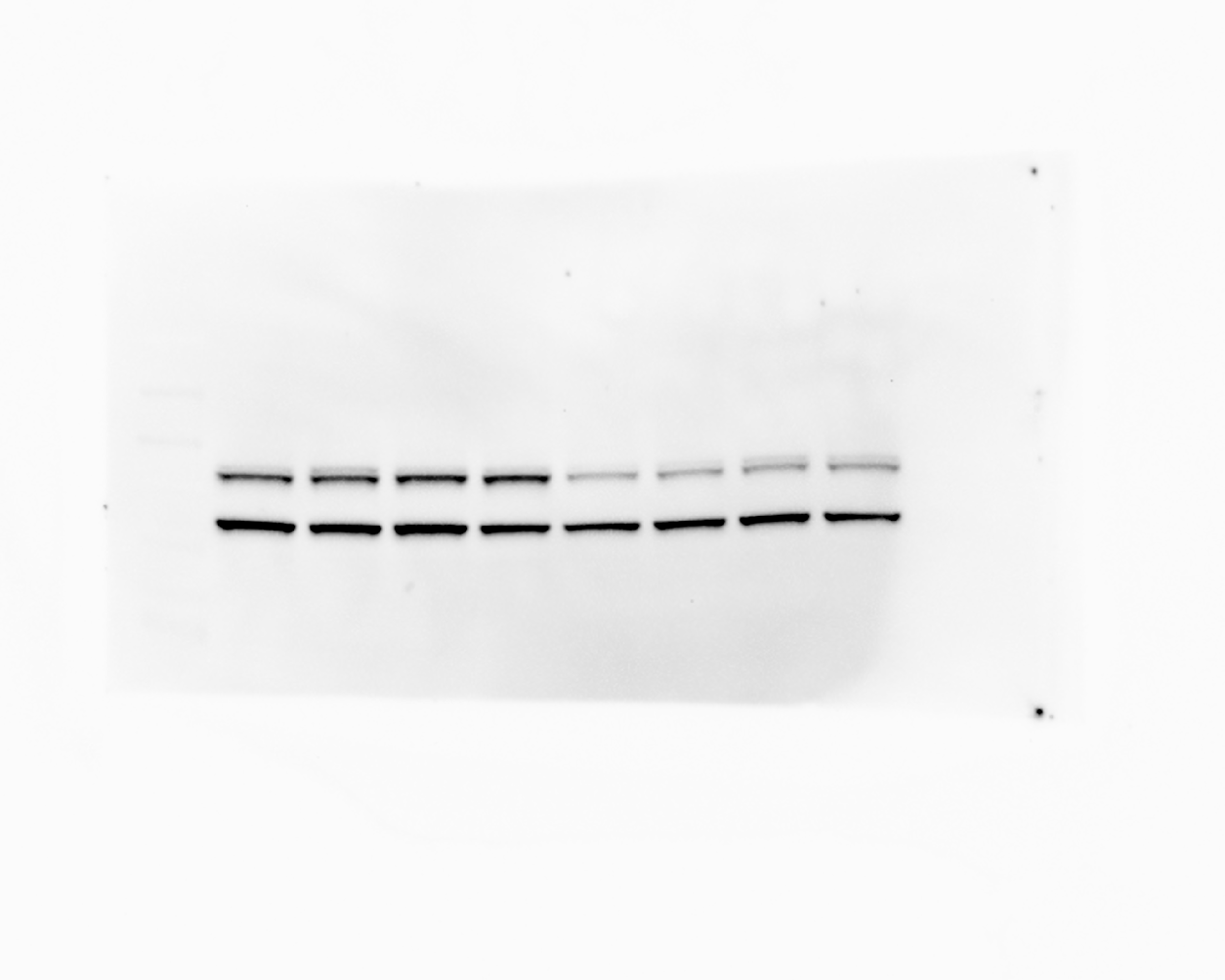

### 20241114_phosphogel2_phosphogel3_paperrerun_gel4_stat5_7(Colorimetric).tif

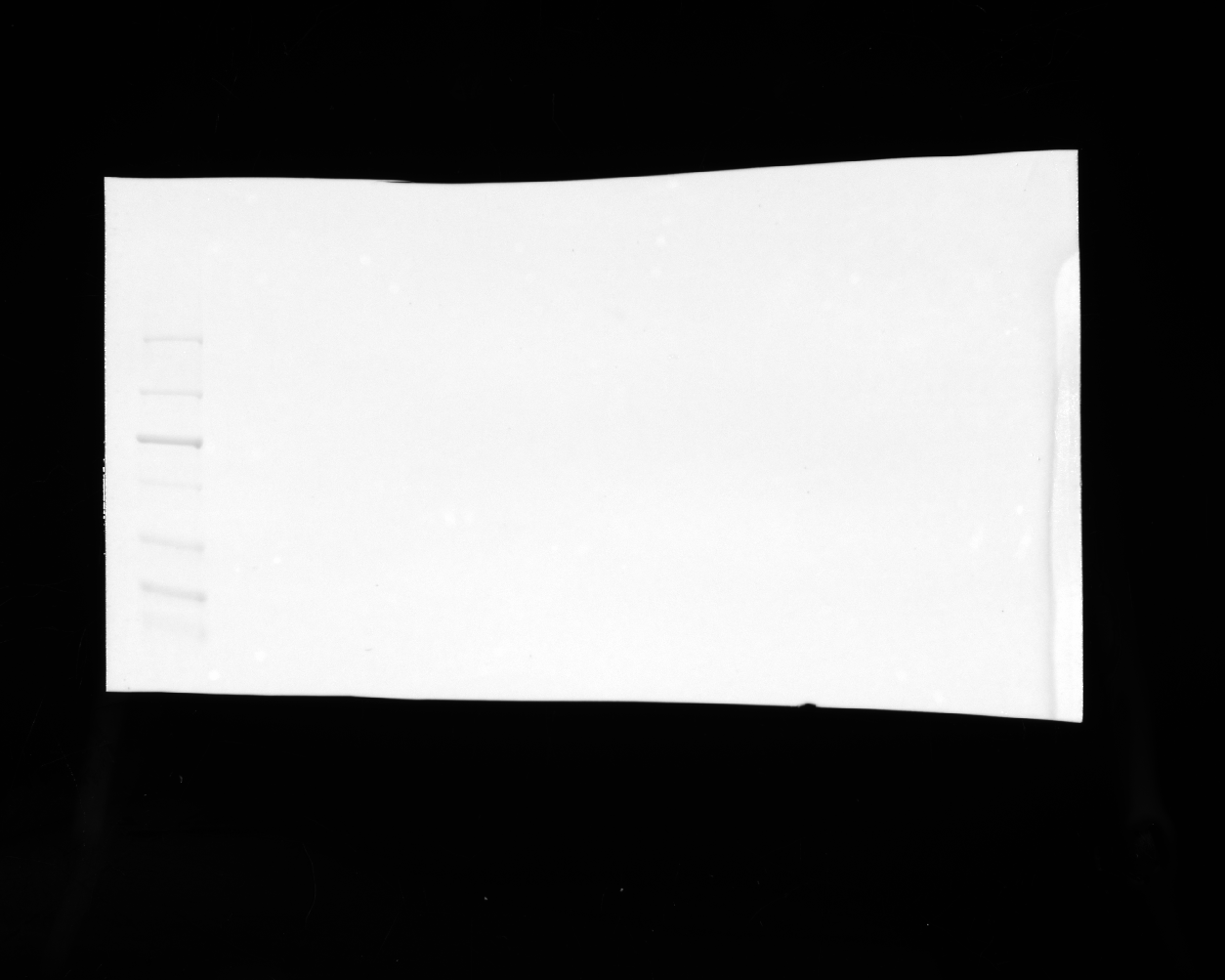

### 20241114_phosphogel2_phosphogel3_paperrerun_gel4_stat5_7(Composite).tif

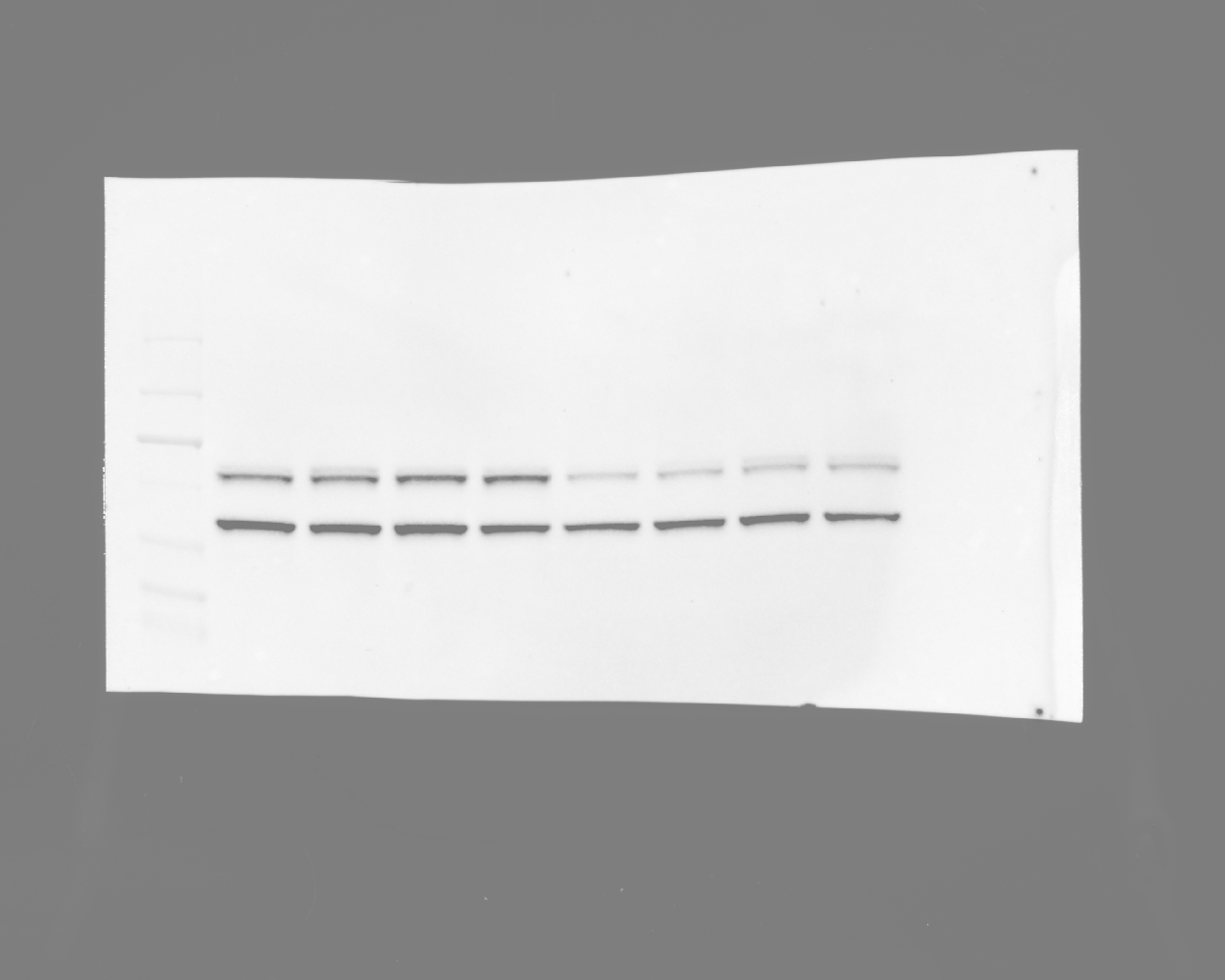

### 20241114_phosphogel2_phosphogel3_paperrerun_gel4_stat5_7_2.tif

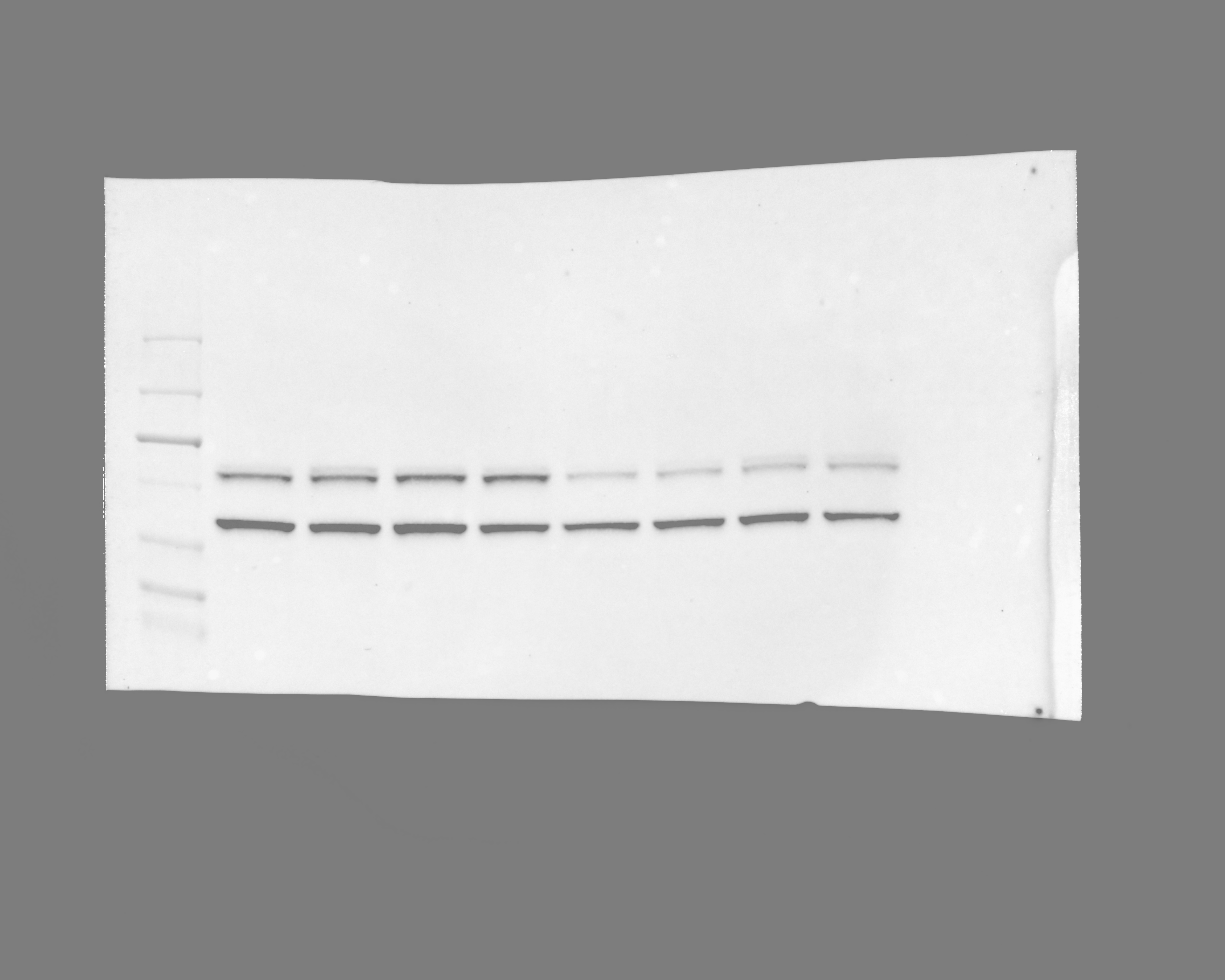

### 20241114_phosphogel2_phosphogel3_paperrerun_gel5_hh3_4 (Multichannel)_final.tif

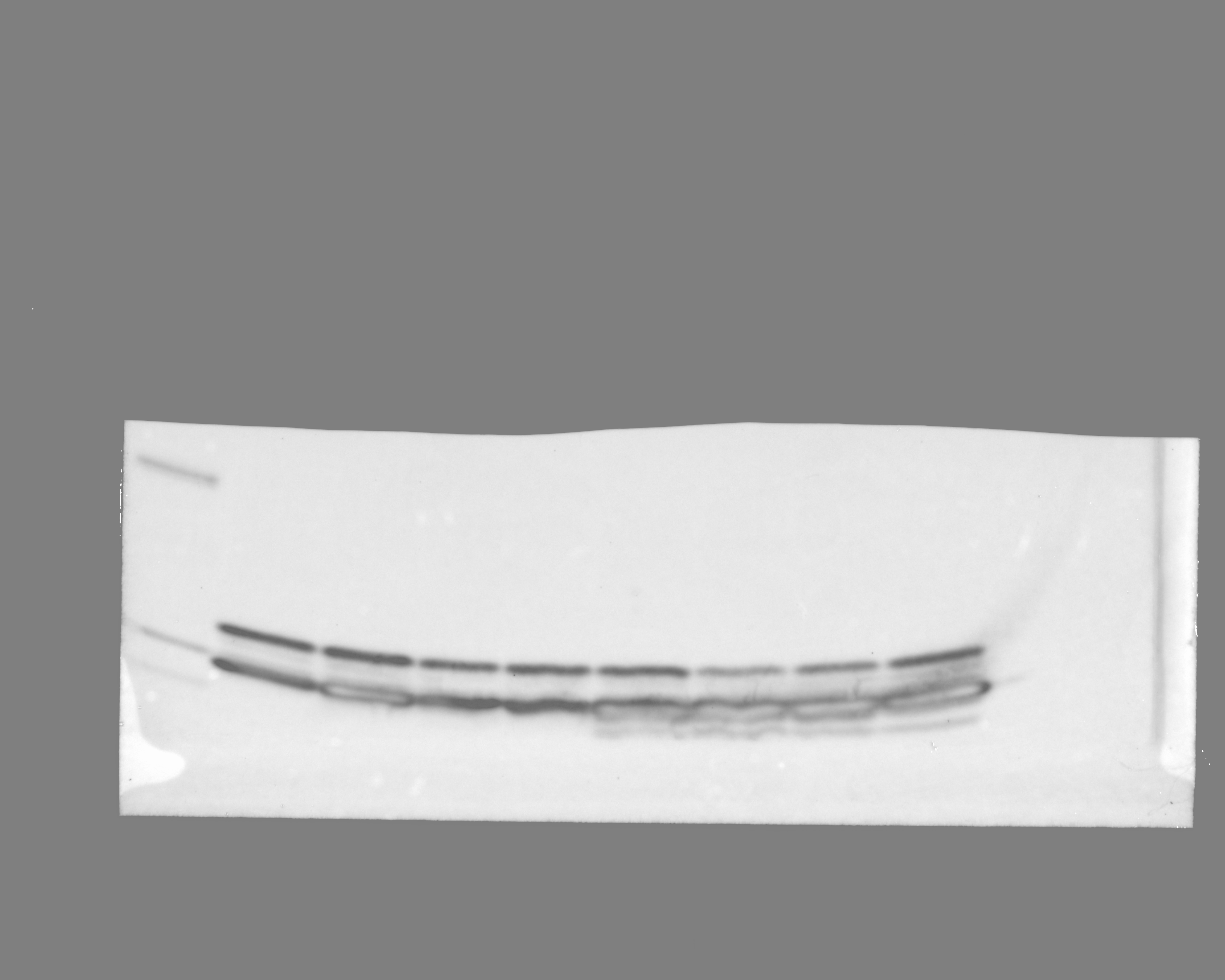

### 20241114_phosphogel2_phosphogel3_paperrerun_gel5_hh3_9(Chemiluminescence).tif

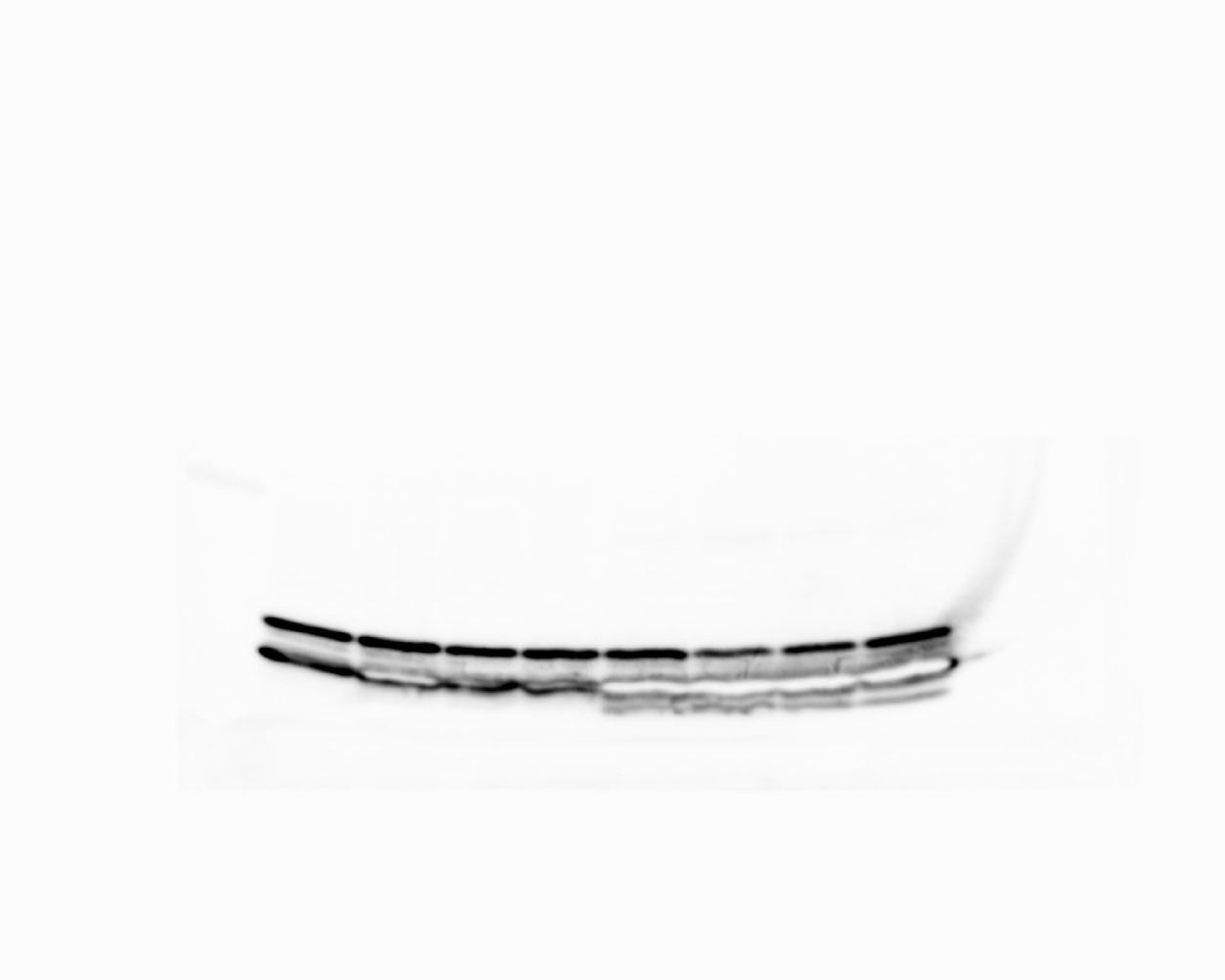

### 20241114_phosphogel2_phosphogel3_paperrerun_gel5_hh3_9(Colorimetric).tif

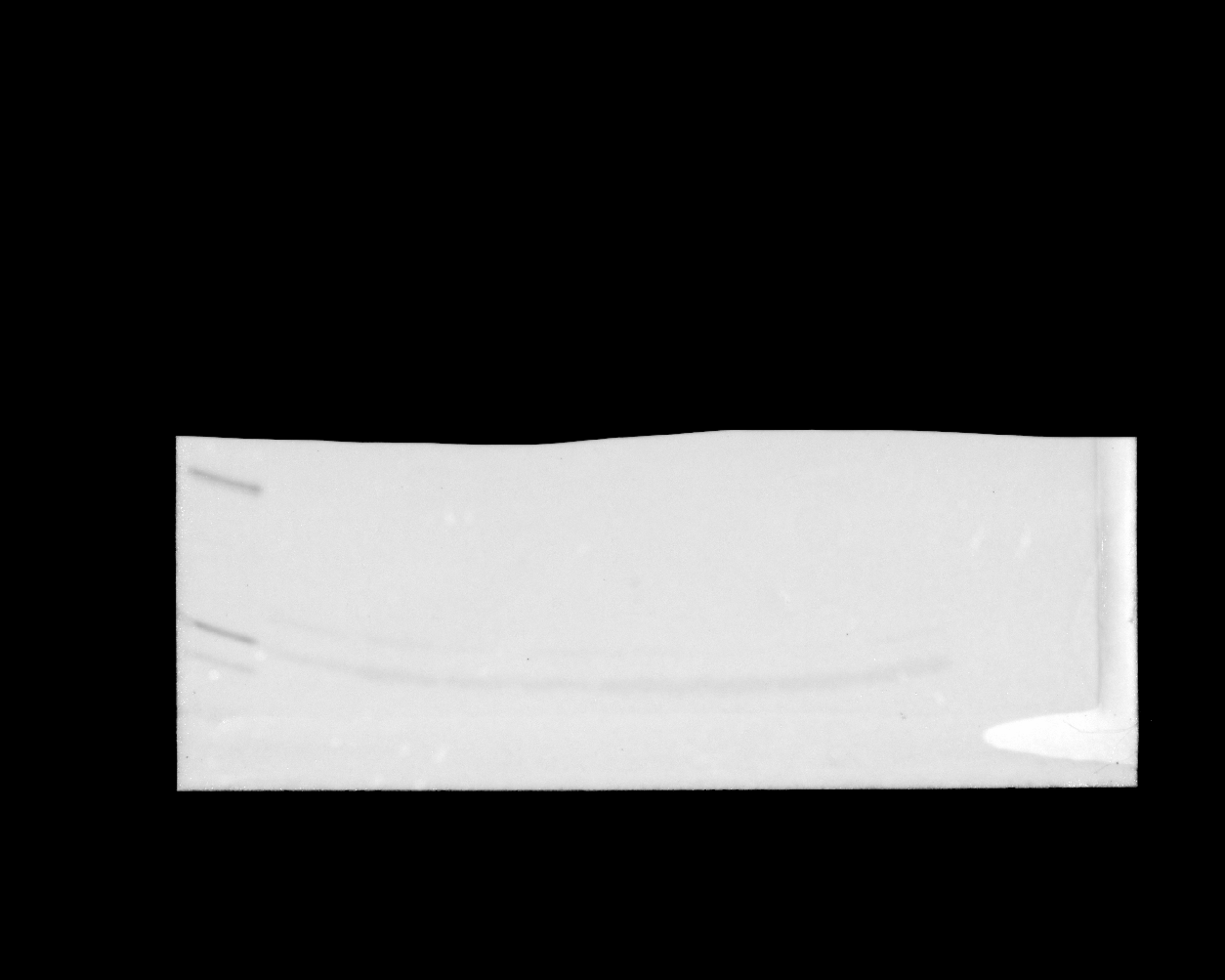

### 20241114_phosphogel2_phosphogel3_paperrerun_gel5_hh3_9(Composite).tif

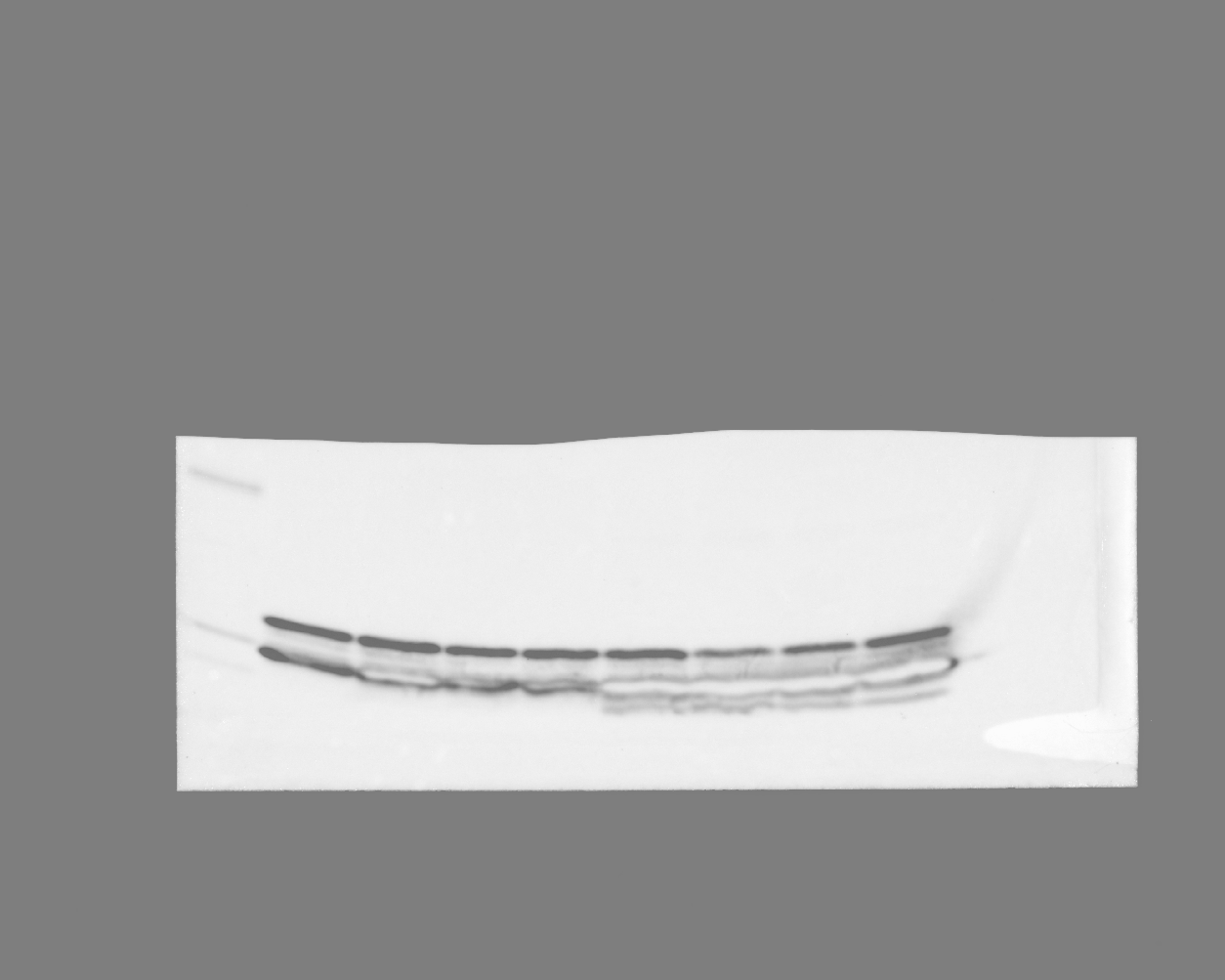

### 20241114_phosphogel2_phosphogel3_paperrerun_gel5_reprobe_stat5_phospho_12(Chemiluminescence).tif

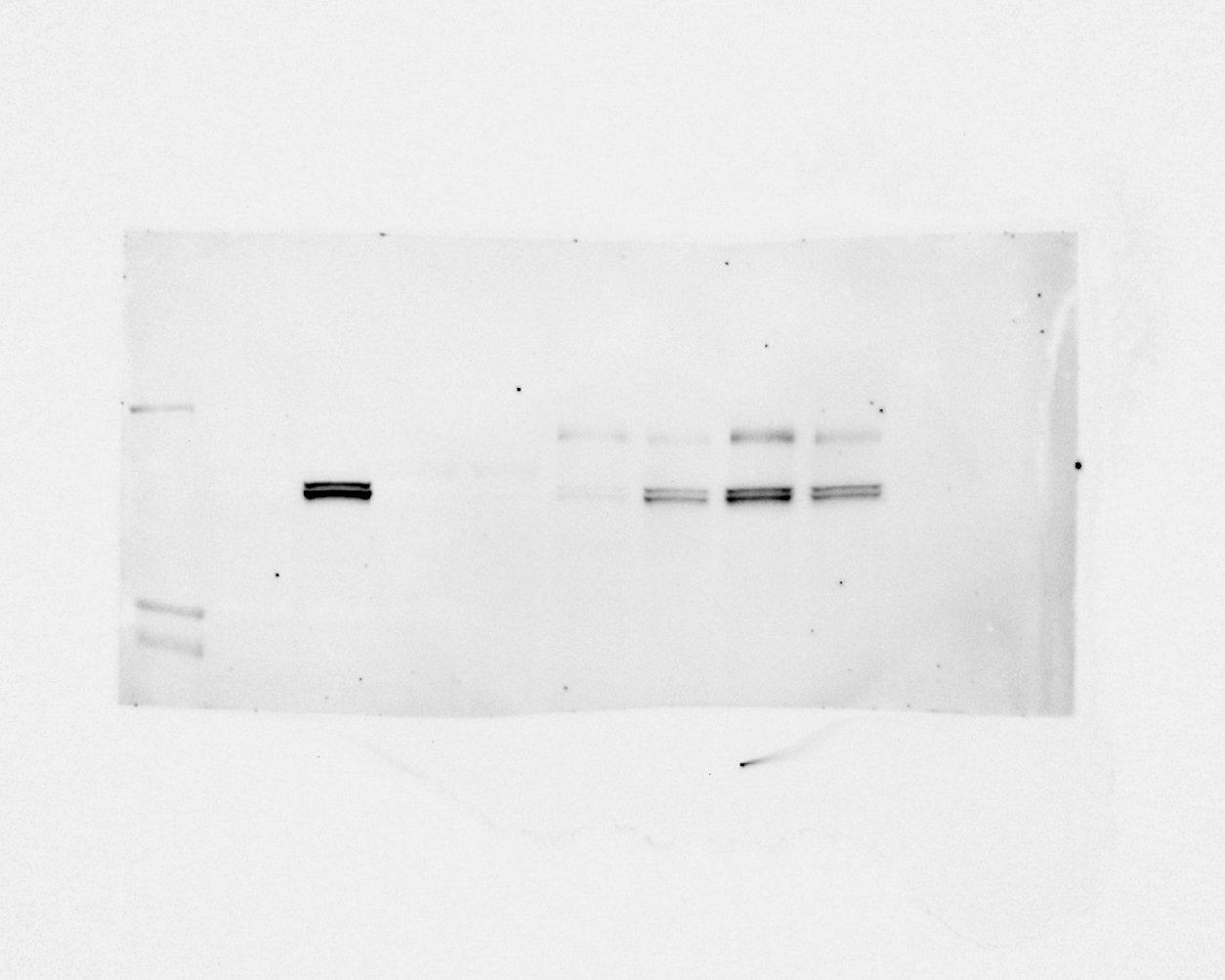

### 20241114_phosphogel2_phosphogel3_paperrerun_gel5_reprobe_stat5_phospho_12(Colorimetric).tif

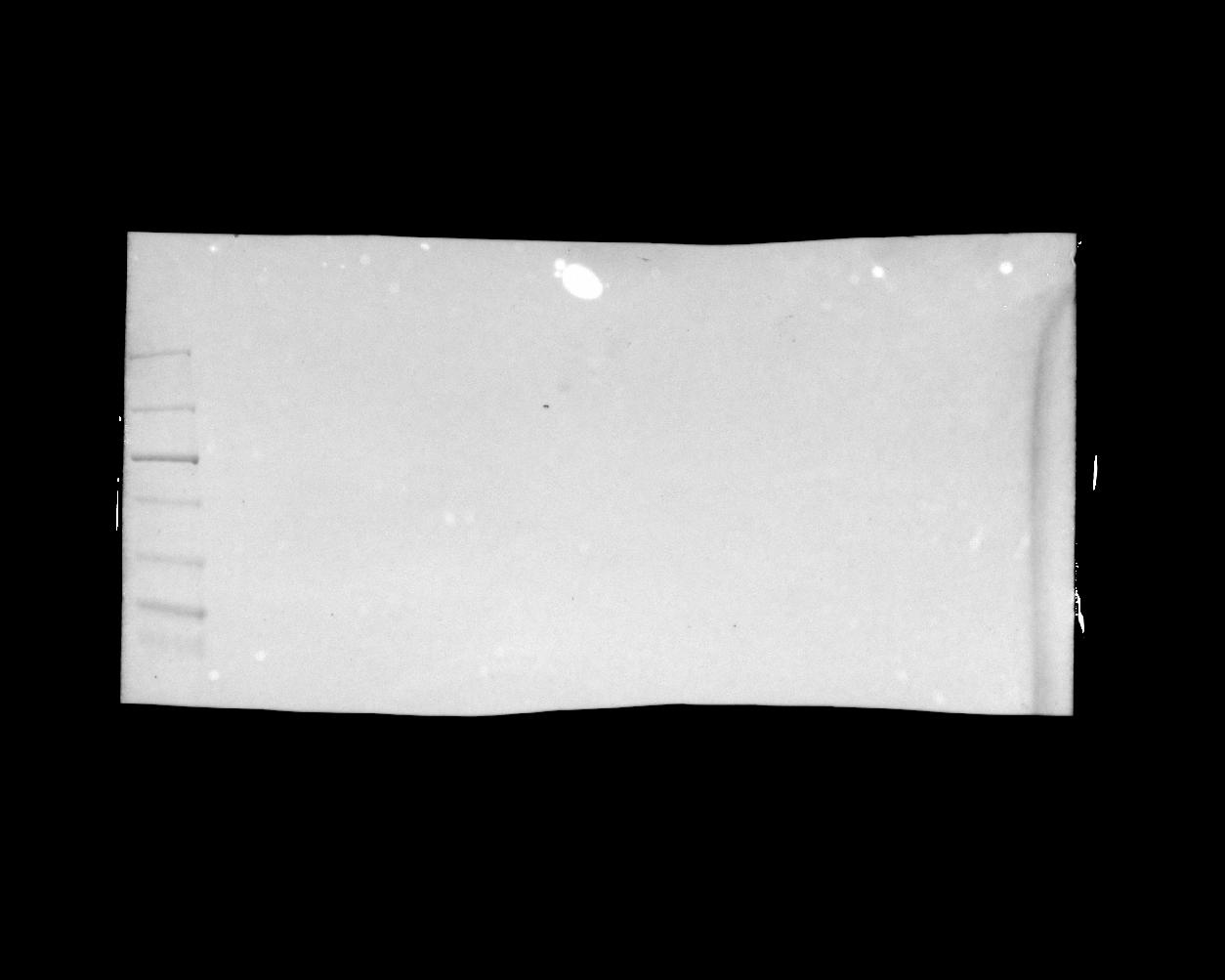

### 20241114_phosphogel2_phosphogel3_paperrerun_gel5_reprobe_stat5_phospho_12(Composite).tif

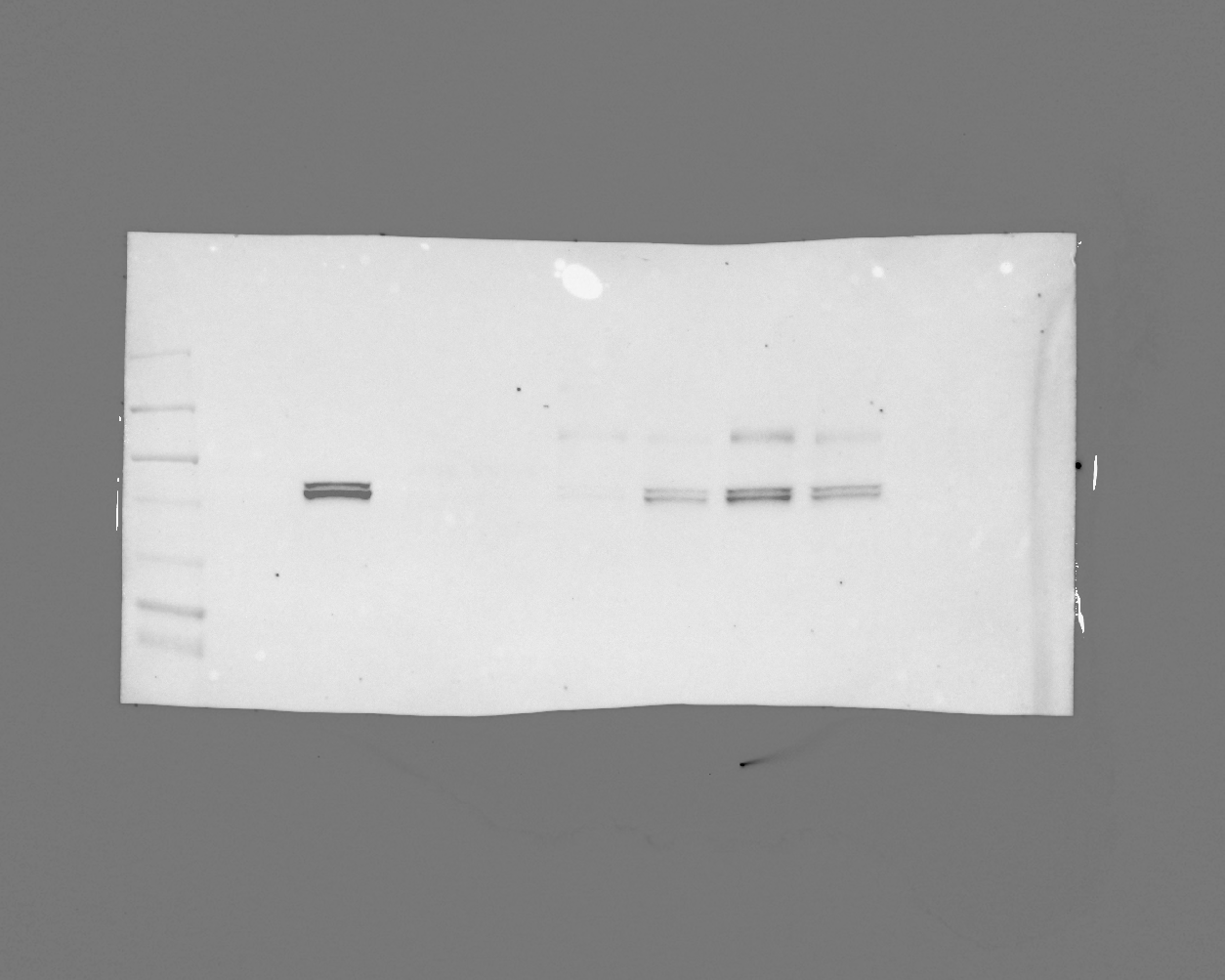

### 20241119_ut7variantgel_gel1_hh3_7(Chemiluminescence).tif

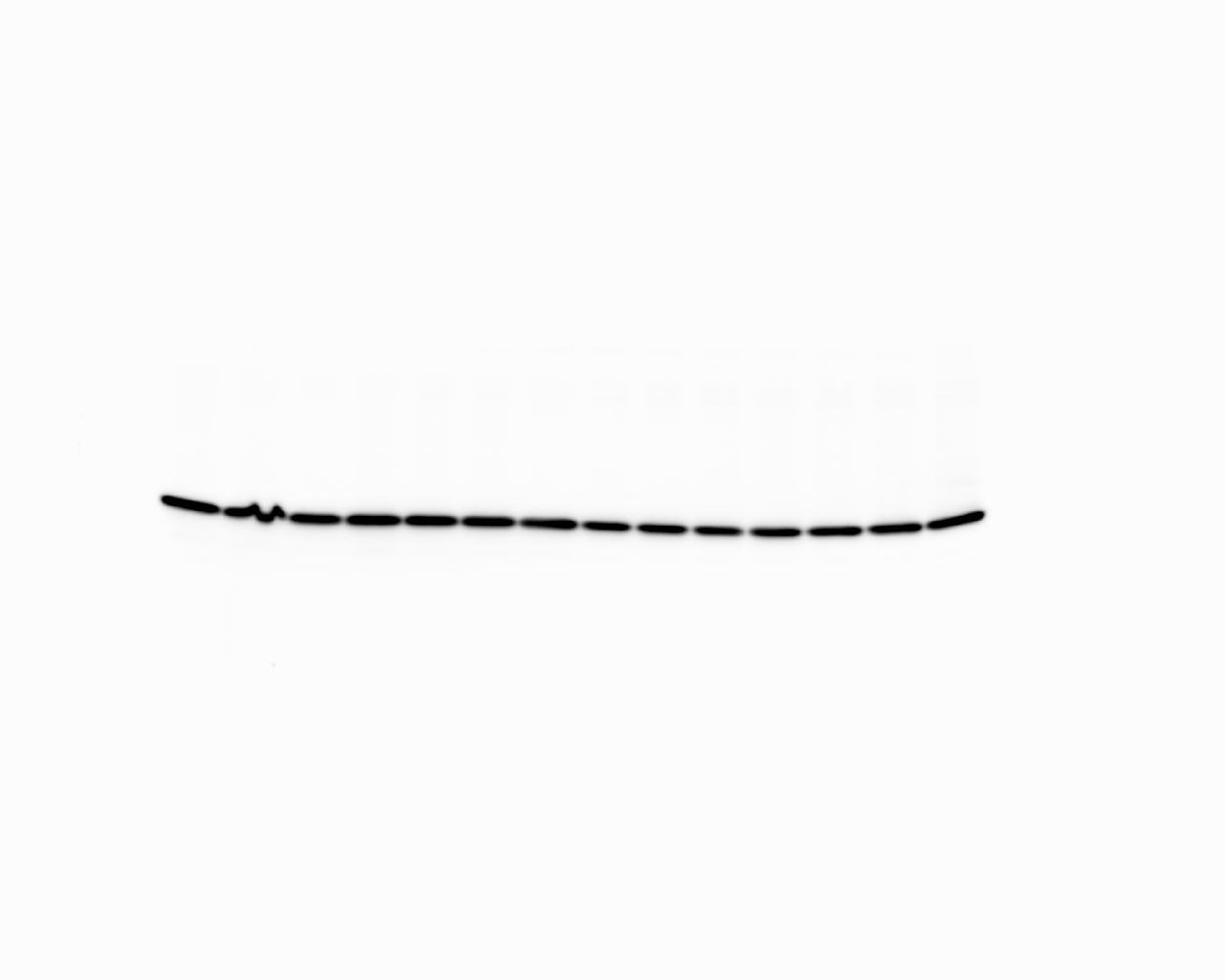

### 20241119_ut7variantgel_gel1_hh3_7(Colorimetric).tif

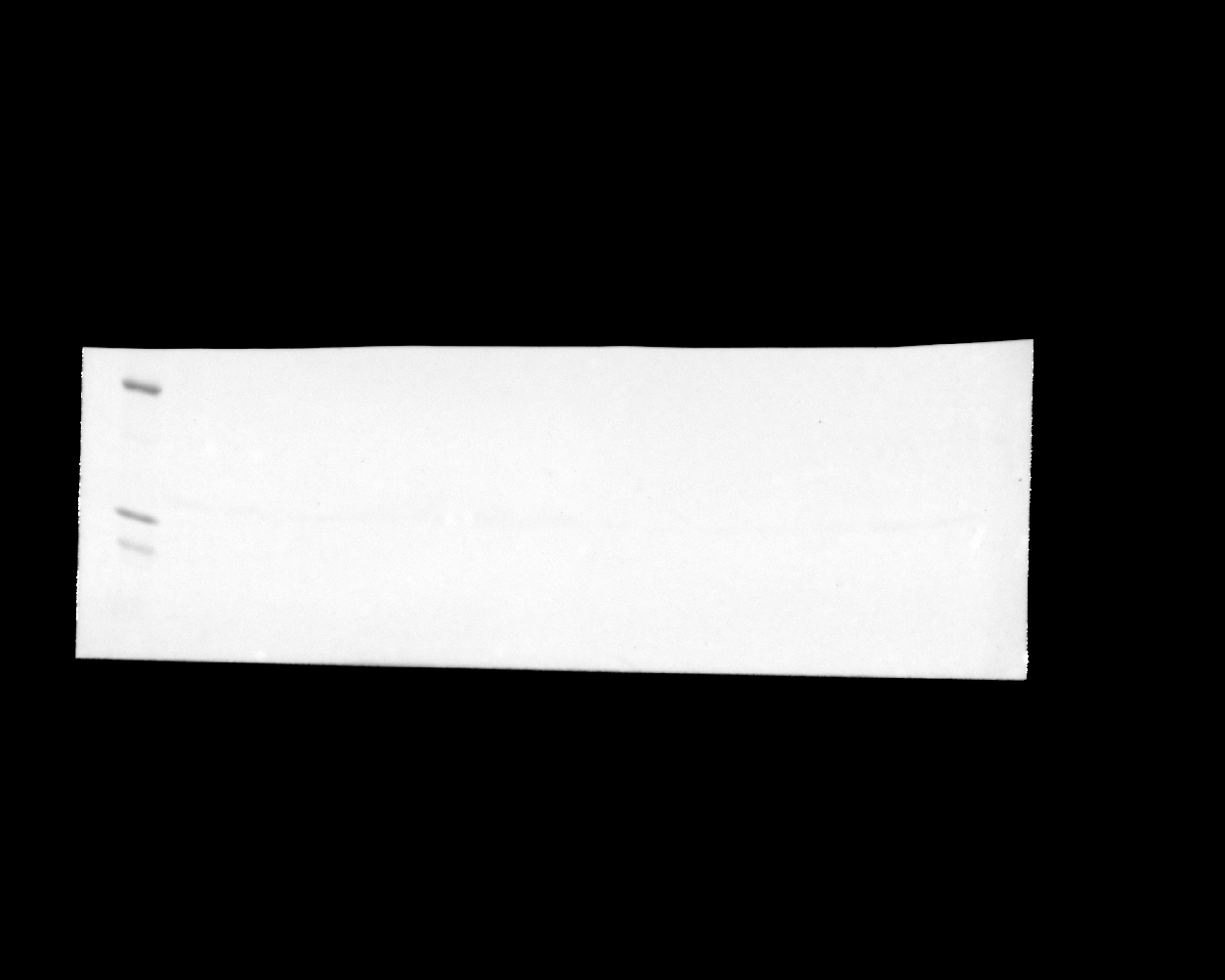

### 20241119_ut7variantgel_gel1_hh3_7(Composite).tif

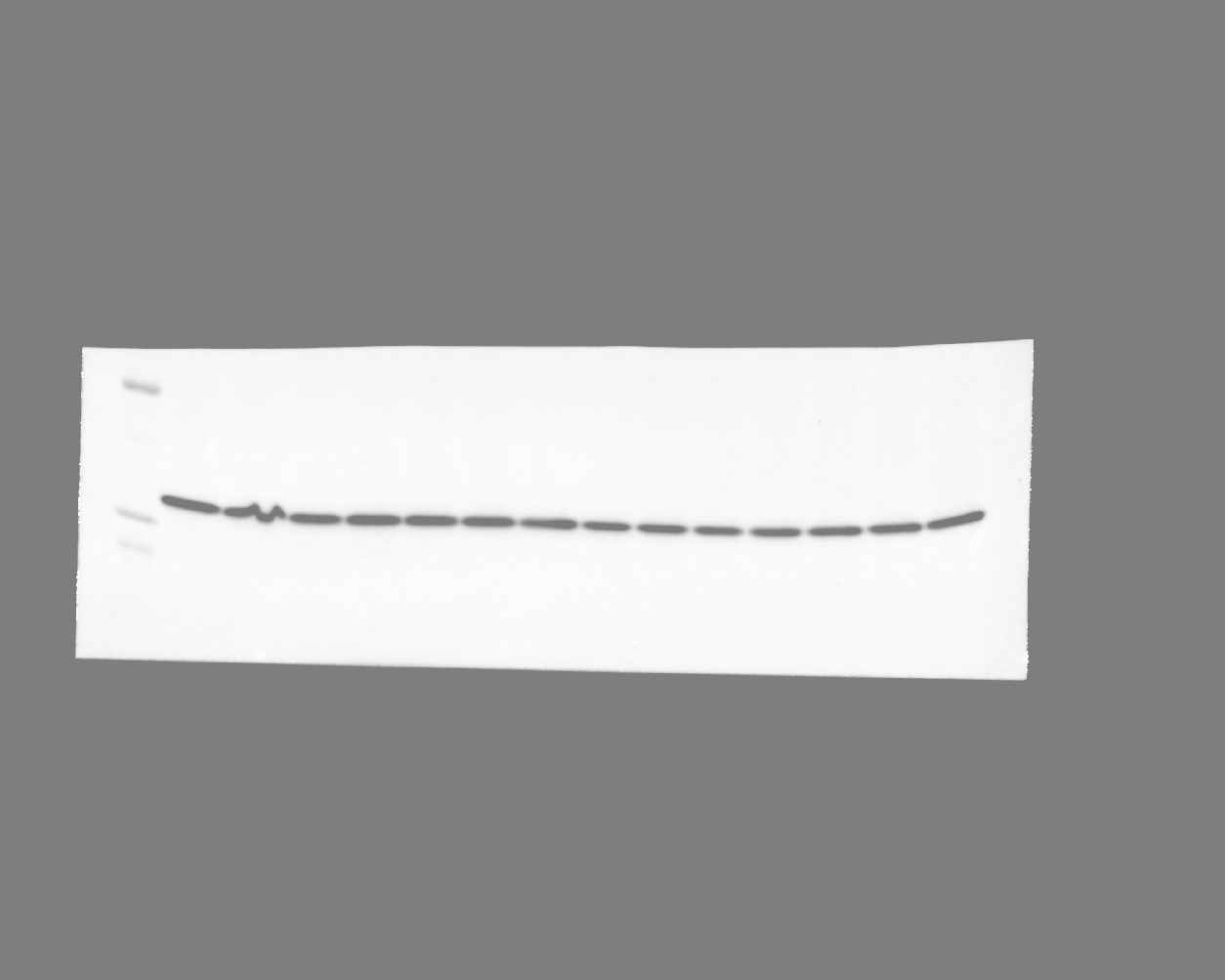

### 20241119_ut7variantgel_gel1_stat5_phospho_13(Chemiluminescence).tif

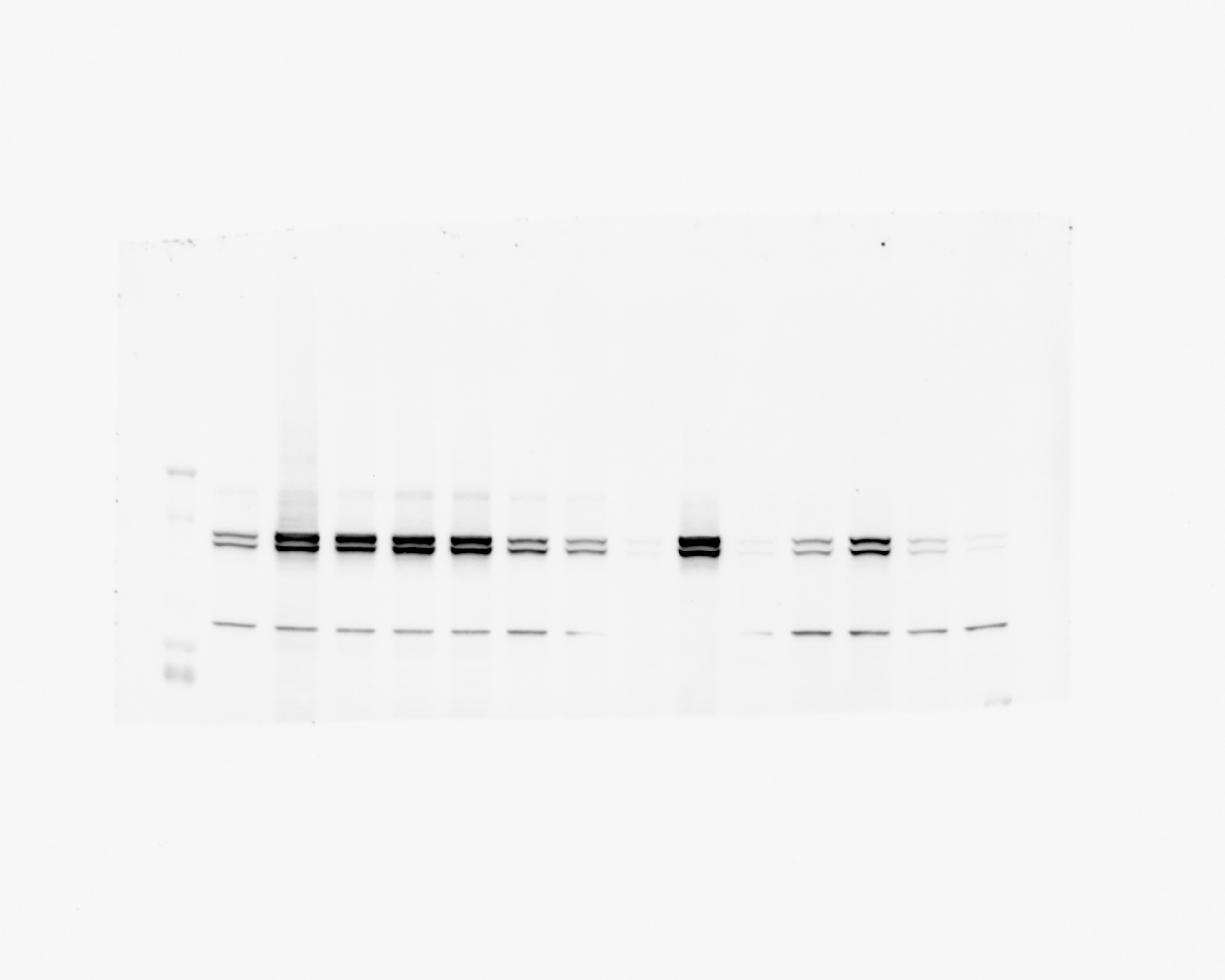

### 20241119_ut7variantgel_gel1_stat5_phospho_13(Colorimetric).tif

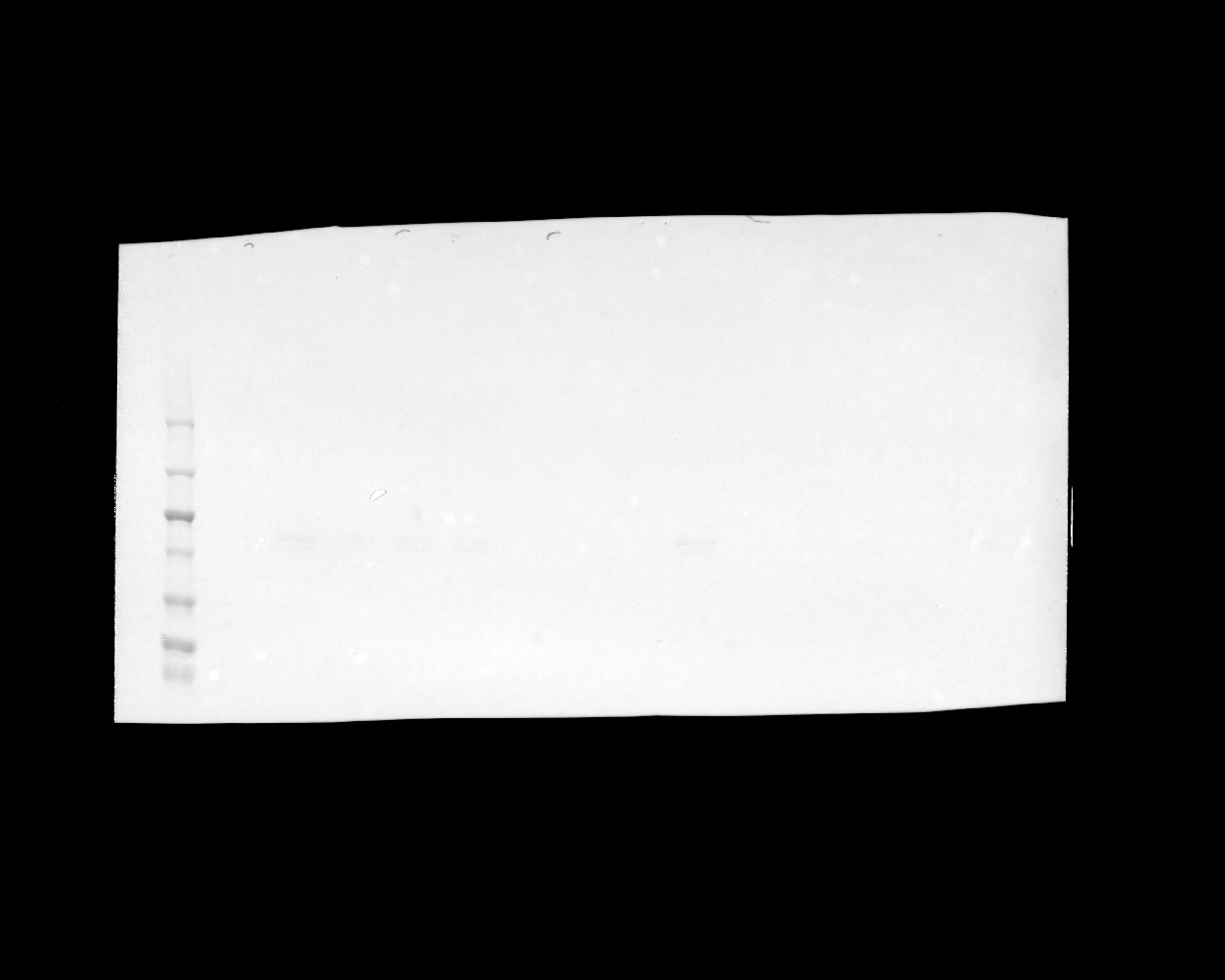

### 20241119_ut7variantgel_gel1_stat5_phospho_13(Composite).tif

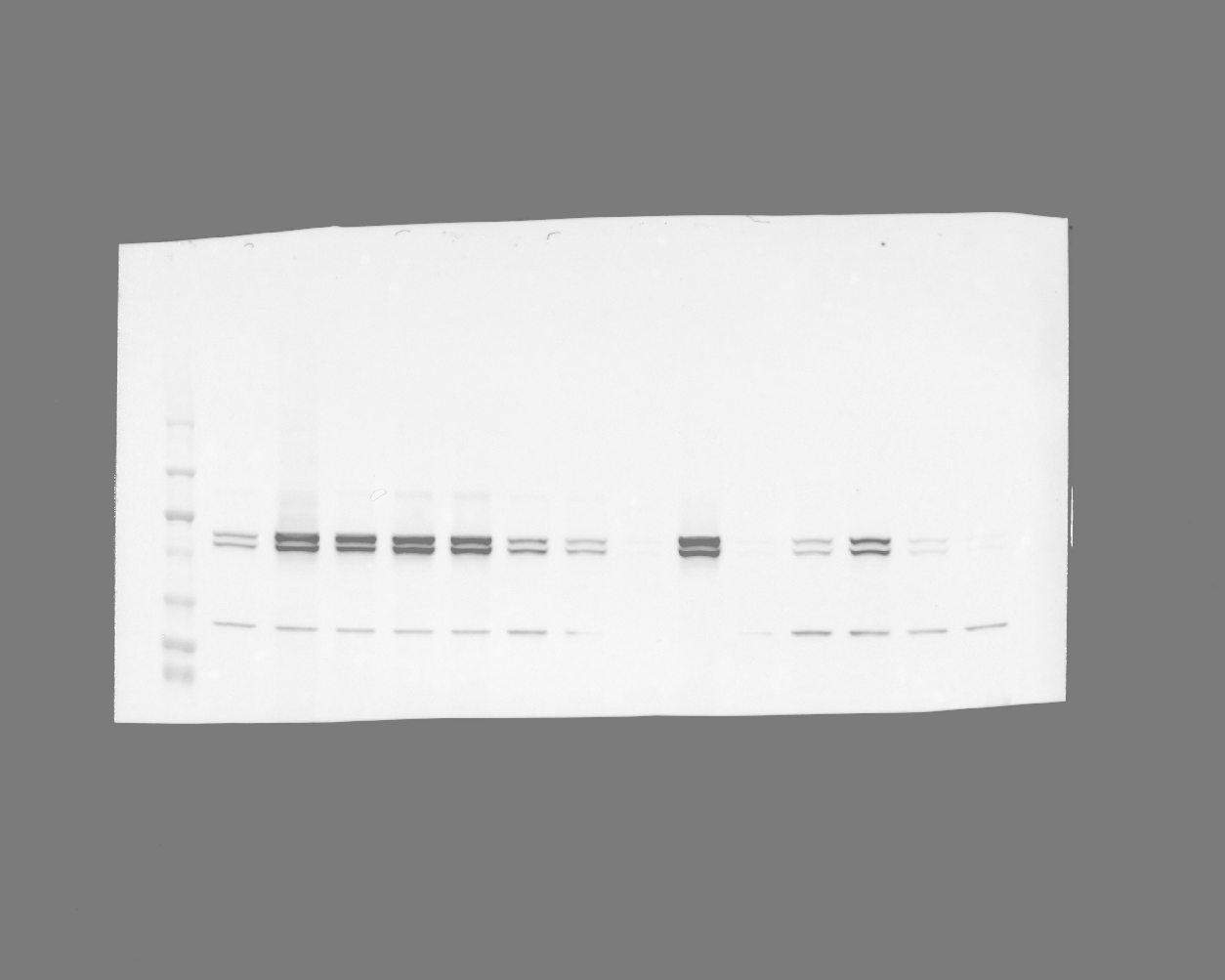

### 20241119_ut7variantgel_gel2_hh3_7(Chemiluminescence).tif

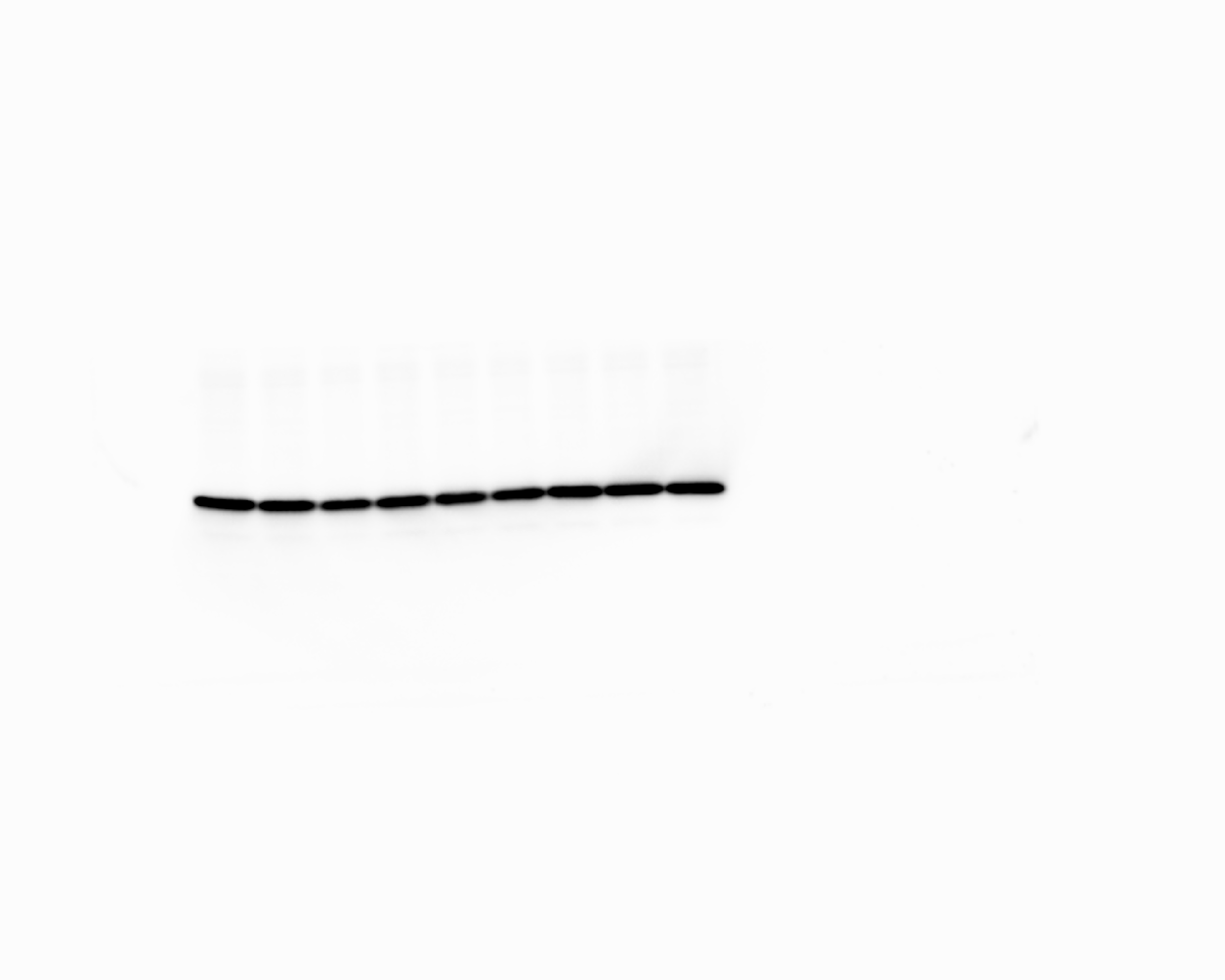

### 20241119_ut7variantgel_gel3_stat5_7 (Multichannel).tif

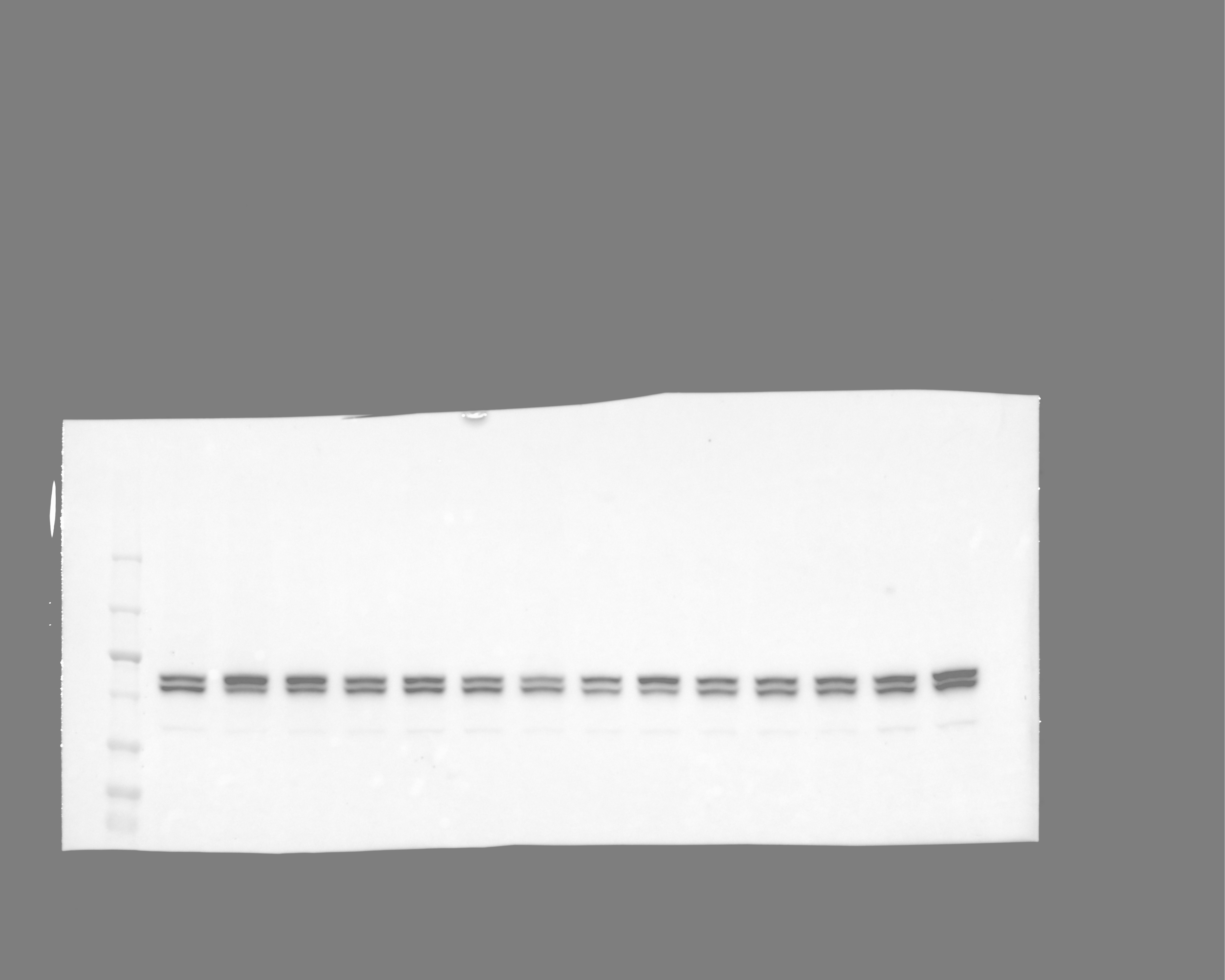

### 20241119_ut7variantgel_gel3_stat5_7(Chemiluminescence).tif

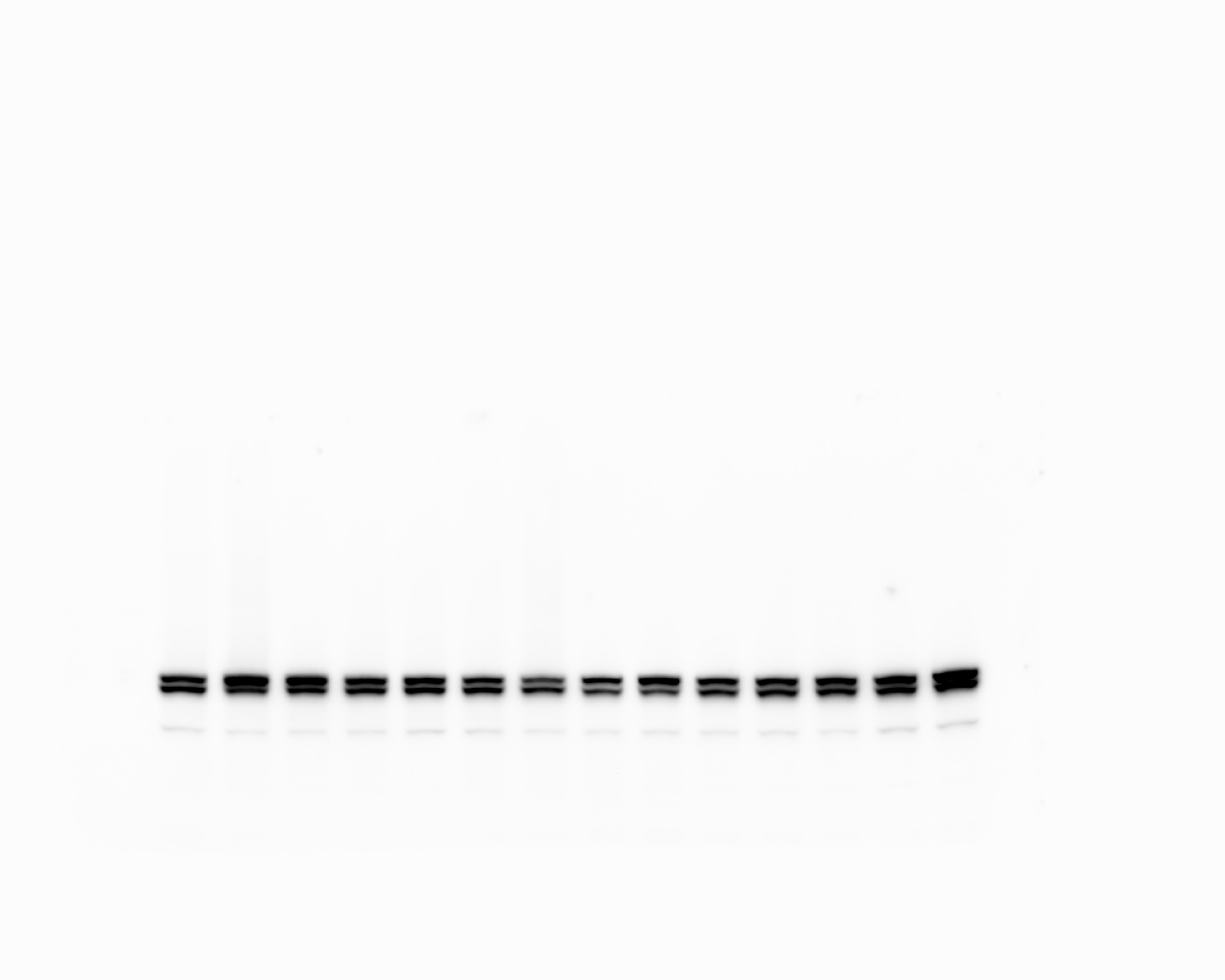

### 20241119_ut7variantgel_gel3_stat5_7(Colorimetric).tif

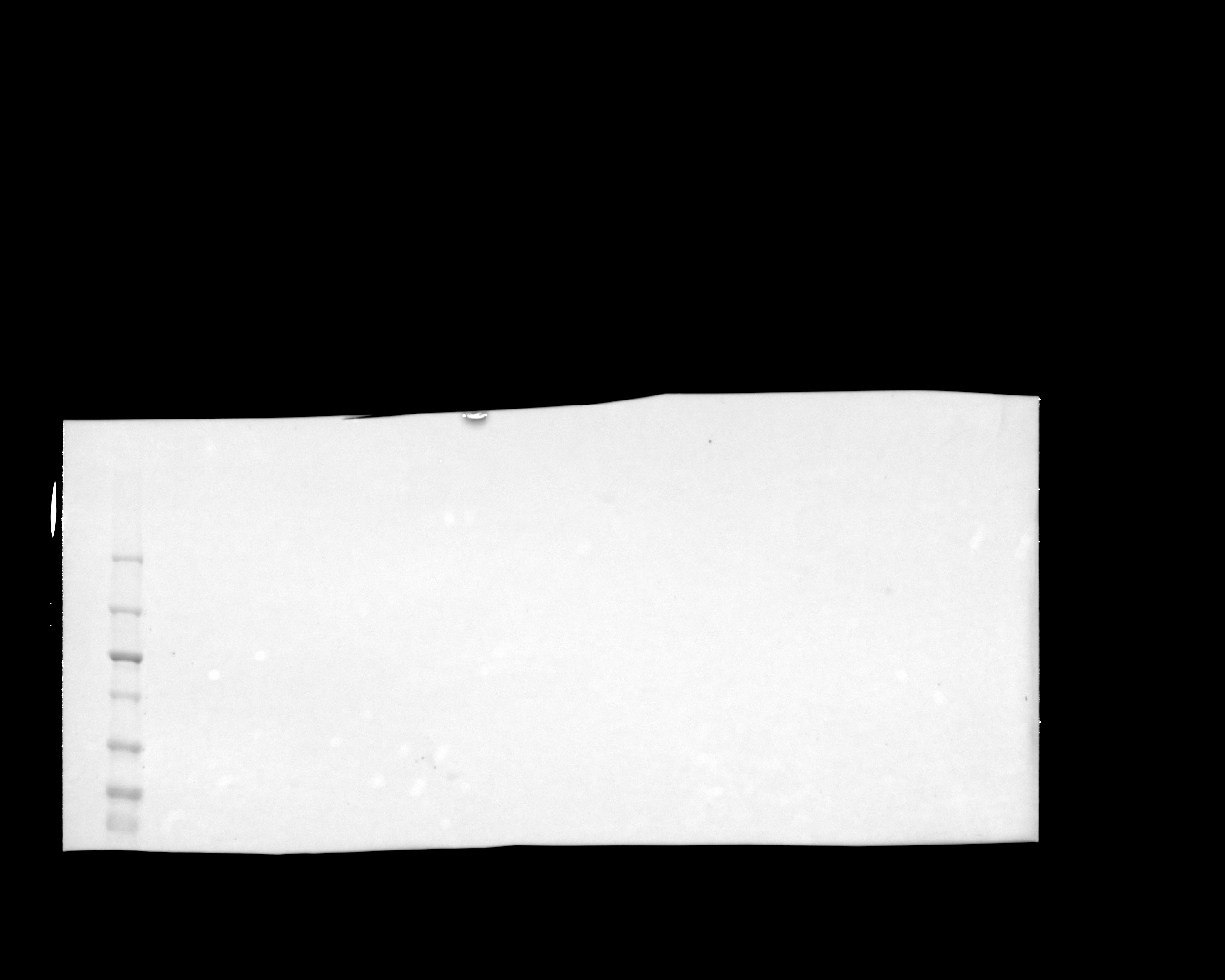

### final_acn_cd131_morecontrast.png

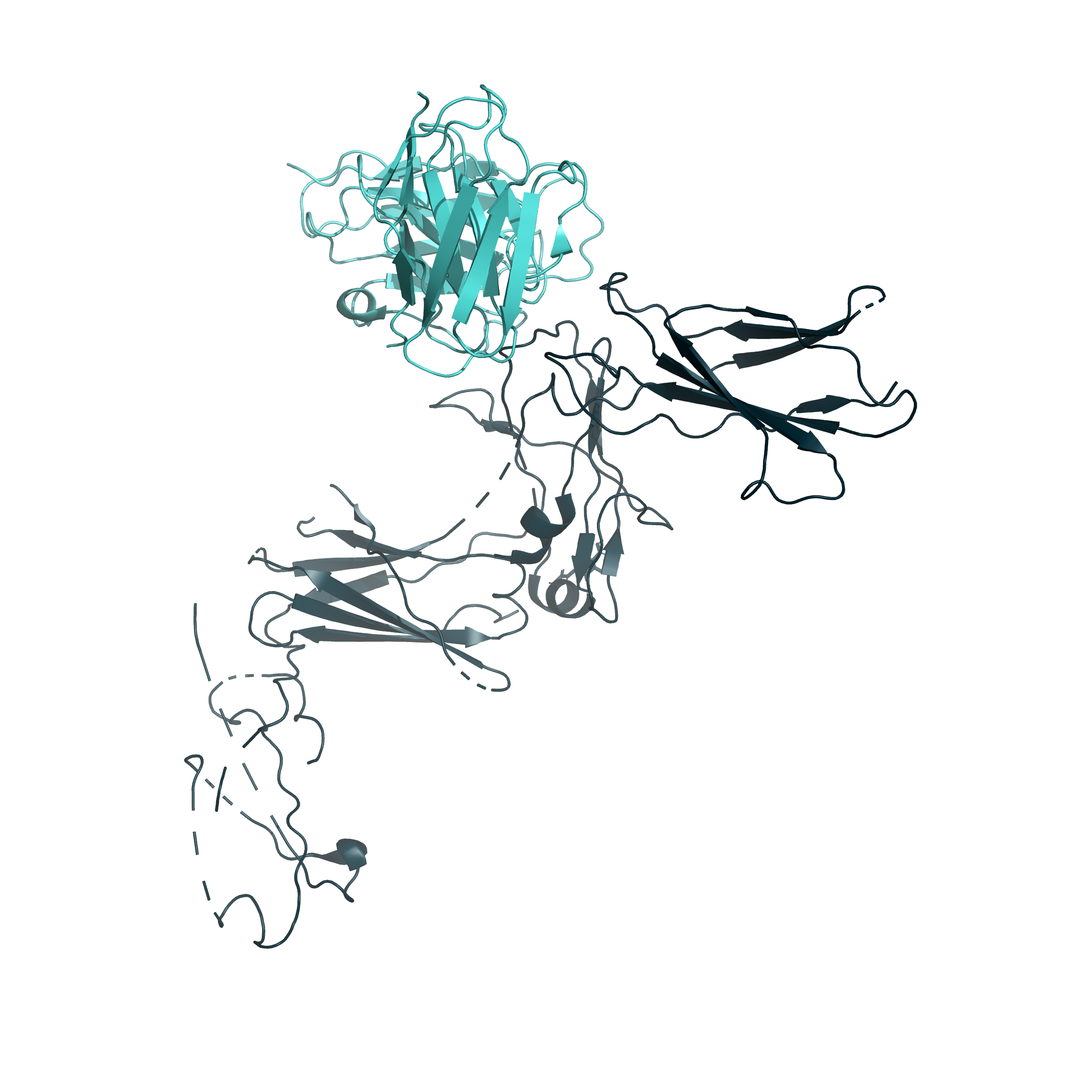

### final_da330_epor_dualcolor_lighter_FF955C_ FF3506.png

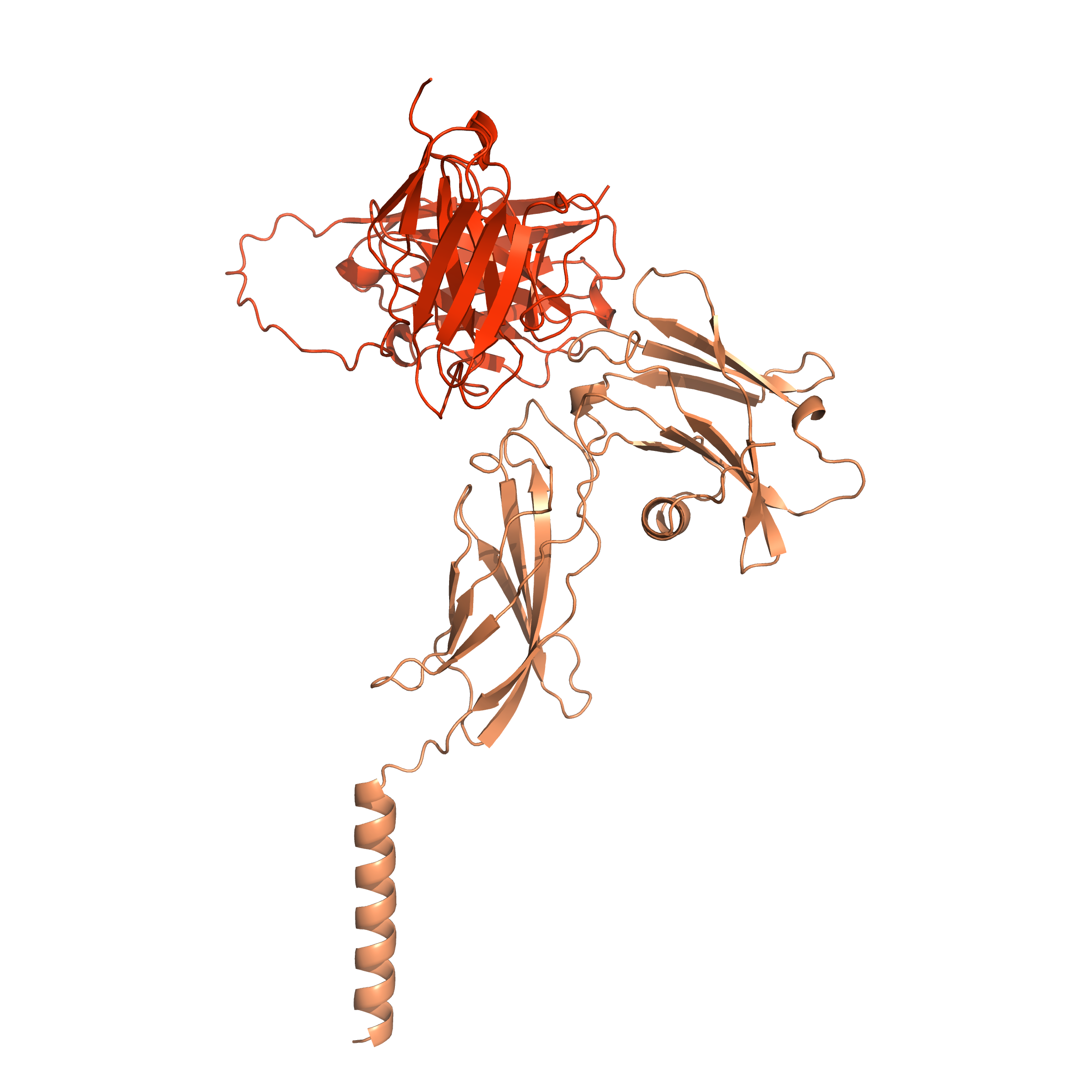

### final_labeled_blueorange_alphafoldoverlay.png

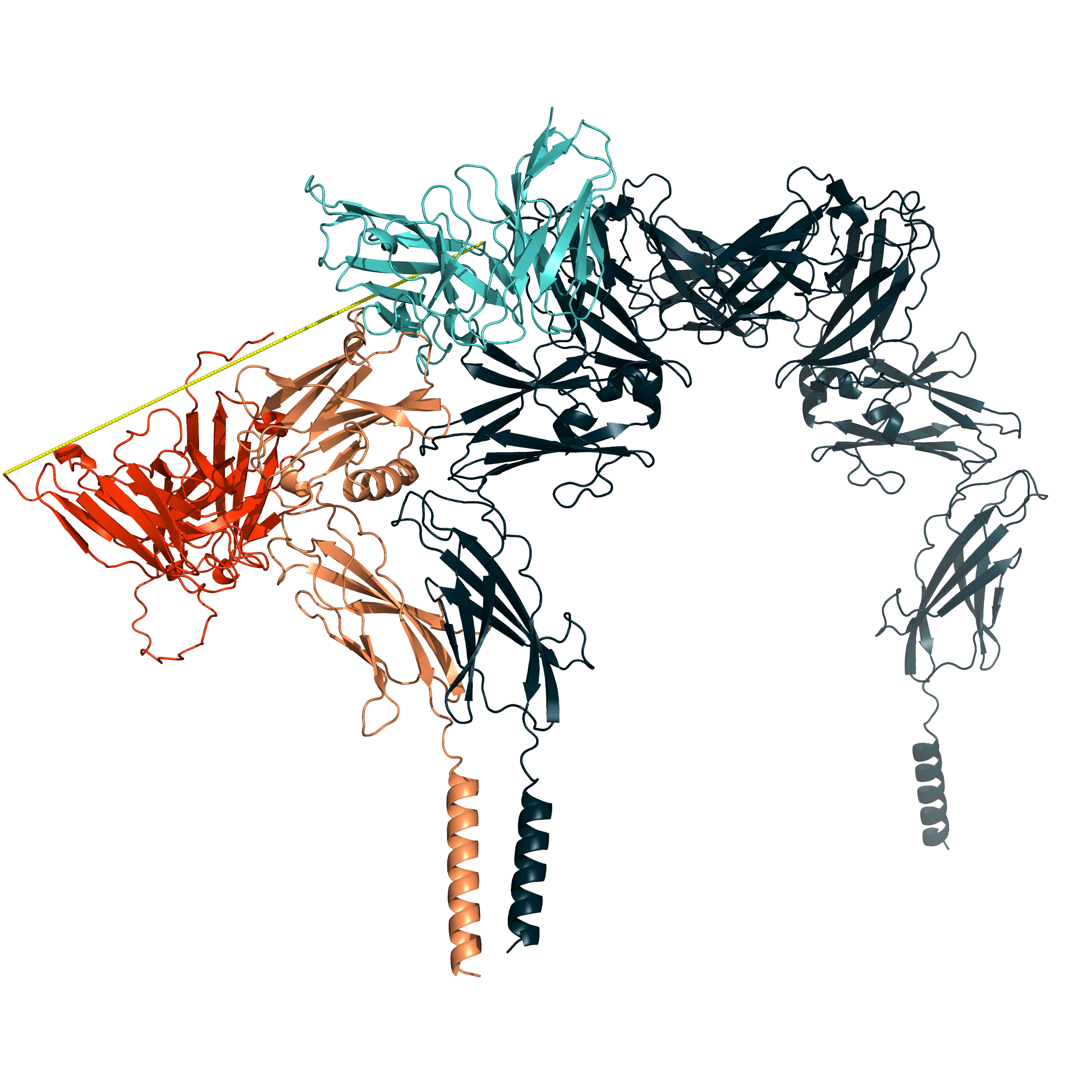
